# A closed-loop optogenetic screen for neurons controlling feeding in *Drosophila*

**DOI:** 10.1101/2020.08.21.261701

**Authors:** Celia KS Lau, Michael D Gordon

## Abstract

Feeding is an essential part of animal life that is impacted greatly by the sense of taste. Although the characterization of taste-detection at the periphery is extensive, higher-order taste and feeding circuits are still being elucidated. Here, we use an automated closed-loop optogenetic activation screen for novel taste and feeding neurons in *Drosophila melanogaster*. Out of 122 Janelia FlyLight Project GAL4 lines preselected based on expression pattern, we identify six lines that acutely promote feeding and 35 lines that inhibit it. As proof of principle, we follow up on the *R70C07-GAL4* neuron population, which strongly inhibits feeding. Using split-GAL4 lines to isolate subsets of the *R70C07-GAL4* population, we find both appetitive and aversive neurons. We also show that *R70C07-GAL4* labels a population of putative second-order taste interneurons that contact both sweet and bitter sensory neurons. These results serve as a resource for further functional dissection of fly feeding circuits.

## Introduction

Gustation is a primary sense conserved across the animal kingdom. It contributes to individual fitness, allowing animals to assess and distinguish between potentially nutritious foods and those that may be toxic. In general, sweet foods (an indicator of energy) promote consumption, whereas bitter foods (an indicator of toxicity) trigger rejection behavior (Yarmolinsky *et al*. 2009). However, the details of how gustatory information is relayed through the brain to evoke the corresponding behaviors remains unclear. On-going research efforts in this area are fuelled by studies conducted in the *Drosophila melanogaster* model, owing to their reliability of innate feeding behaviors and the accessibility of powerful genetic tools.

Similar to mammalian taste receptor cells, fruit flies have gustatory receptor neurons (GRNs) that are capable of detecting basic taste qualities, including sweet, bitter, salt and acids, responsible for triggering food acceptance or rejection (Scott 2018; Chen and Dahanukar 2019). The tuning of sweet and bitter GRNs are dictated by gustatory receptors (GRs). Namely, Gr5a-positive neurons respond broadly to sweetness and those expressing Gr66a respond to bitter (Thorne *et al*. 2004; Wang *et al*. 2004; Marella *et al*. 2006; Dahanukar *et al*. 2007). GRNs send projections to the brain, with arborizations terminating in the subesophageal zone (SEZ). This compartment of the brain is referred to as the primary taste center, acting as the first point of taste signal relay (Rajashekhar and Singh 1994; Thorne *et al*. 2004; Wang *et al*. 2004; Ito *et al*. 2014). GRN projection patterns in the SEZ are roughly localized based on the taste modality it is encoding and the origin of the GRN— from the pharynx, proboscis or legs (Wang *et al*. 2004; Kwon *et al*. 2014).

While GRNs are well characterized, only a handful of studies have identified downstream neurons. These include the sweet gustatory projection neurons (sGPNs), which make contact with Gr5a projections, project to the antennal mechanosensory and motor center (AMMC), and evoke PER (Kain and Dahanukar 2015). Another pair of second-order neurons identified were denoted the gustatory second-order neurons (G2N-1s)(Miyazaki *et al*. 2015). These also receive synaptic input from Gr5a GRNs, but unlike sGPNs, the G2N-1s locally arborize and terminate within the ventral SEZ. A third set of distinct second-order neurons activated by sucrose are specific to pharyngeal inputs (Yapici *et al*. 2016). These cholinergic neurons were named the ingestion neurons (IN1), and their activation is sufficient to prolong ingestion. More recently, a single pair of bilaterally symmetrical interneurons called bitter gustatory local neurons (bGLNs) was characterized to be activated by bitter tastants and is sufficient in inhibiting attractive behavior upon receiving signals from bitter GRNs (Bohra *et al*. 2018). Aside from second-order taste neurons that reside locally in the SEZ, long-range taste projection neurons (TPNs) also exist to relay taste input to regions of the higher brain (Kim *et al*. 2017). Notably, the valence of signals appears to remain consistent and segregated at the level of second-order taste neurons. Ultimately, the discovery of these second-order taste neurons brings us closer to understanding the pathways by which peripheral taste can be translated into the proper motor response. However, a global picture of taste circuitry remains obscure, suggesting the need for identifying more higher-order taste and feeding neurons.

Recently, we developed the Sip-Triggered Optogenetic Behavior Enclosure (STROBE) for closed-loop optogenetic activation of fly neurons during feeding (Jaeger *et al*. 2018; Musso *et al*. 2019). The STROBE temporally couples LED activation to interactions between a fly and one of two food sources in a small arena. In combination with targeted expression of light-gated channels, this system effectively activates peripheral and central neurons, allowing real-time modulation of the fly’s sensory experience or motor patterns during feeding (Musso *et al*. 2019). The preference of the fly for the light-triggering food compared to the non-light-triggering food indicates the appetitive or aversive valence of the neurons undergoing optogenetic activation.

In this study, 122 Janelia FlyLight Project enhancer-GAL4 lines (Jenett *et al*. 2012) were crossed with *UAS-CsChrimson* and subjected to testing in the STROBE. We found six lines that produced a preference for the light-triggering food and 35 lines that drove preference for the non-light-triggering food. One line in particular, *R70C07-GAL4*, was chosen for further characterization of its role in feeding inhibition. A GAL4 hemidriver version of *R70C07* (*R70C07-p65*) was combined with five different GAL4.DBD hemidrivers to generate five split-GAL4s labeling subsets of an SEZ interneuron population that is prominent within the *R70C07-GAL4* expression pattern. Unexpectedly, while one split-GAL4 line phenocopied the aversion seen with *R70C07-GAL4* activation, another drove the opposite effect (attraction), and three produced little or no significant effect. Nonetheless, GFP reconstitution across synaptic partners (GRASP) revealed that neurons within this population make contacts with both sweet and bitter GRNs, suggesting a role in taste processing. The results presented here demonstrate the feasibility of using the STROBE to identify novel candidate taste and feeding neurons, and provide a resource of lines for finer-scale dissection in the future.

## Materials and methods

### Drosophila stocks and crosses

Fly stocks were raised on standard cornmeal-dextrose fly food at 25°C in 70% humidity. *20XUAS-IVS-CsChrimson*.*mVenus* (BDSC, stock number: 55135) was used for optogenetic activation. See S1 for the full list of enhancer-GAL4 lines of the Janelia FlyLight Project (http://flweb.janelia.org/) that were used for the optogenetic activation screen. The following split-GAL4 lines were created by combining selected hemidrivers with *R70C07-p65*.*AD* (stock 71122): *SEZ1-GAL4* (*R37H08-GAL4*.*DBD* stock 68786); *SEZ2-GAL4* (*R53C05-GAL4*.*DBD* stock 69451); *SEZ3-GAL4* (*VT044519-GAL4*.*DBD* stock 75123); *SEZ4-GAL4* (*R38E08-GAL4*.*DBD* stock 69427); and *SEZ5-GAL4* (*R10E08-GAL4*.*DBD* stock 69792). For GRASP experiments we used: *Gr5a-LexA::VP16, UAS-CD4::spGFP1-10, LexAop-CD4::spGFP11, UAS-CD8::dsRed* (Gordon and Scott 2009) and *Gr66a-LexA::VP16* (Thistle *et al*. 2012).

### Fly preparation and STROBE experiments

Flies were maintained at 25°C in 70% humidity. Females (2-5 d after eclosion) were collected and allowed to recover in fresh vials containing standard medium for at least 1 d before being transferred to covered vials that contained 1 ml standard medium with either 1 mM of all-*trans*-retinal and 1% ethanol as a vehicle (retinal-fed group) or ethanol vehicle alone (non-retinal-fed controls) for 2 days. Flies were then starved for 20-24 hrs in similar conditions, except the standard medium was replaced with 1% agar. As such, flies were maintained on either + or – all-*trans*-retinal diets throughout the 3 d that preceded testing. To promote food interaction during testing, flies were water-deprived for 1 hour.

Both channels of the STROBE chambers were loaded with 4 μl of 1% agar for the activation screen. Some follow-up experiments were performed with 100 mM sucrose in 1% agar to further promote food interactions. To commence each experiment, the acquisition on the STROBE software was initiated before flies were individually placed in each arena via mouth aspiration. Experiments were continued for 1 hour and the preference indices were calculated as: (Interactions with Food 1 – Interactions with Food 2)/ (Interactions with Food 1 + Interactions with Food 2). The red LED is always associated to the left channel, with Food 1. Details of the STROBE system, including the design and programming were previously described (Musso *et al*. 2019).

### Immunohistochemistry

Brain immunohistochemistry was performed as previously described (Chu *et al*. 2014). To stain flies with the *UAS-CsChrimson*.*mVenus* transgene, the following primary antibodies were used: mouse anti-brp (1:50, Developmental Studies Hybridoma Bank #nc82) and rabbit anti-GFP (1:1000, Invitrogen), with secondary antibodies: goat anti-rabbit Alexa-488 (A11008, Invitrogen) and goat anti-mouse Alexa-546 (A11030, Invitrogen). For GRASP experiments, the following were used as primary antibodies: mouse anti-GFP (1:100, Sigma-Aldrich, G6539), rat anti-DN-cadherin (1:25, DSHB DNEX#8), and rabbit anti-DsRed (1:1000, Clontech #632496) with secondary antibodies: goat anti-mouse Alexa-488 (A11029, Invitrogen), goat anti-rat Alexa-568 (A11077, Invitrogen) and goat anti-rabbit Alexa-647 (A21245, Invitrogen).

All images were acquired using a Leica SP5 II Confocal microscope. Images taken at a magnification of 25x were with a water immersion objective with a Z-stack step size of 1 μm, while at 63x were with oil immersion with a step size of 0.5 μm.

### Statistics and data exclusion

The STROBE sometimes records very small or very large interaction numbers due to technical malfunctions. To account for this, trials from individual flies were removed under three conditions: (i) if no interactions were recorded from the light-triggering channel; (ii) if fewer than five interactions were recorded on the non-light-triggering channel; (iii) if the number of interactions was more than 2.68 standard deviations from the mean. The rationale for (i) and (ii) is that very aversive neurons can produce few interactions on the light-triggering channel, but flies will generally record more than five interactions on the other channel in a functional trial.

Statistical tests were performed using Graphpad Prism 6. T-tests were used to compare experimental (retinal fed) to control (same genotype not fed retinal). The purpose of these statistics is to evaluate the significance of effects within individual genotypes for the purposes of selecting lines for follow-up, rather than to minimize the overall false positive rate. Therefore, no correction was applied when combining the different genotypes in the summary graph shown in Figure 1.

**Figure 1.**
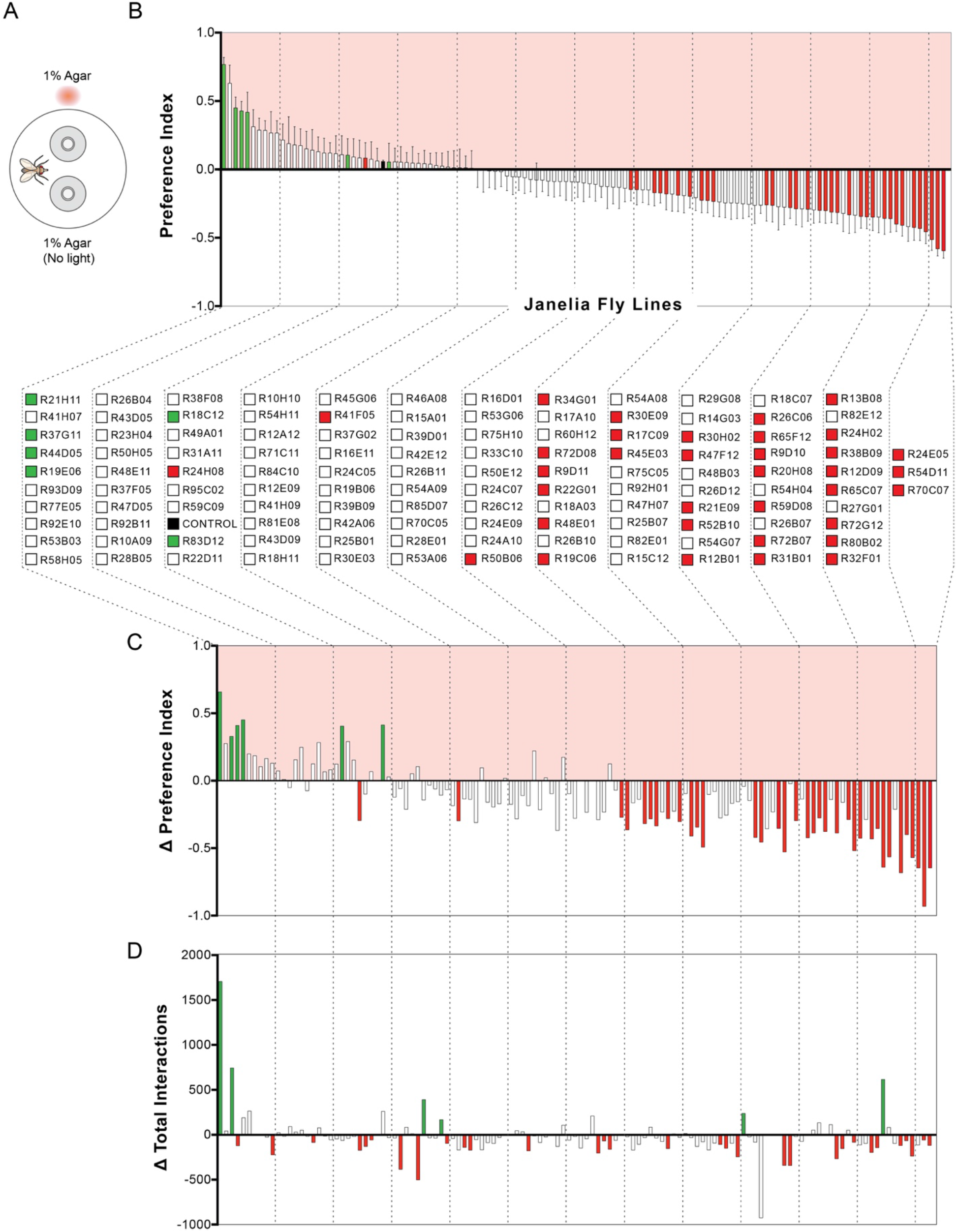
Summary of STROBE screen results. (**A**) Experimental setup: each STROBE arena contains two channels containing 1% agar, and only interactions with one of the channels triggers red light activation. (**B**) Mean preference indices of GAL4 driver lines that were tested in the STROBE. A positive PI indicates preference for the light-triggering food, while a negative PI indicates avoidance of the light-triggering food. Color code indicates significance compared to genetically identical, non-retinal-fed controls (control preferences for each line not shown): green bars denote significantly more positive preference than controls and red bars denote lines that generated a significantly more negative preference than controls; the dark grey bar shows the aggregate responses of the non-retinal controls across all experiments. Values represent mean ± SEM. n = 8–37, except for the aggregate control where n = 1978 (this aggregate control was not used for any statistical testing). (**C**) Difference in preference index between experimental and control groups for each line. Bars are color coded as in (B). (**D**) Difference in average total interaction numbers across both channels (light and non-light) for each line. Color code indicates significance compared to non-retinal controls for each line: green bars denote significantly elevated interactions compared to controls; red bars indicate significantly suppressed interactions compared to controls. Detailed results for all lines are depicted graphically in File S1 and raw data is presented in Files S2 and S3.

### Data and reagent availability

All raw numerical data is available for download in supplemental files. Reagents are available upon request.

## Results & Discussion

### Optogenetic screening of driver lines with the STROBE

The Janelia FlyLight Project has generated more than 8000 transgenic GAL4 lines, providing a vast resource for manipulating specific neurons in the fly brain (Pfeiffer *et al*. 2008; Jenett *et al*. 2012). We selected 122 GAL4 driver lines from the collection based on the following criteria: having relatively sparse labeling, having split-GAL4 versions available, and labeling neurons that have not yet been implicated in taste. Importantly, the first two criteria predetermine the feasibility of further neural population refinement since sparseness allows the systematic selection of different neuron populations to isolate via split-GAL4 combinations. Flies expressing CsChrimson under control of the selected GAL4 drivers were fed with all-*trans-*retinal 3 days prior to the experiment to make CsChrimson functional, whereas control flies of the same genotype were not fed all-*trans-*retinal. Flies were individually mouth-aspirated into STROBE arenas containing two choices of identical plain agar (1%), where interactions with one of the choices triggers a red LED light to excite neurons with functional CsChrimson (Figure 1A). Flies that choose both options equally would have a near-0 preference index (PI), while flies that interacted relatively more or less with the light-paired option are represented by positive and negative PIs, respectively.

Our screen identified six GAL4 lines that produced a significantly positive preference in the STROBE compared to their matched isogenetic non-retinal controls, and 35 lines that produced a significantly negative preference (Figure 1B). Since food interactions measured in the STROBE correlate with food consumption (Itskov *et al*. 2014; Musso *et al*. 2019), we can interpret these lines as containing neurons that acutely impact feeding behavior. Notably, there are examples where a significant difference from controls was observed, despite an absolute preference near zero. This is because each line was compared to its own set of controls, and there are cases where the control group displayed a preference that deviated from neutrality, despite the fact that pooling all the controls revealed the expected preference near zero (Figure 1B). The difference between experimental and control preferences for each line is displayed in Figure 1C. We also identified lines that produced a change in total sip number across both food choices, which may or may not be associated with a change in preference (Figure 1D). These lines are good candidates for containing neurons that exert persistent modulation of feeding that lasts beyond the time period of individual feeding events and therefore affects interactions with the non-light-triggering food as well. Detailed data for each line, including expression, time curves and interaction numbers for each replicate is presented in graphical form (File S1) and raw data is available for download (Files S2 and S3) in the supplementary materials.

By pitting the choice of agar against agar paired with neuronal activation, we were able to efficiently identify driver lines labeling neurons that impact feeding in either a positive or negative direction. One question is why we observed many more lines producing a negative feeding preference. We speculate that this is because there are many ways to decrease feeding – paralysis, inducing a behavior that interrupts feeding, or producing any kind of negative percept. One the other hand, we expect effects that increase feeding to be relatively more specific to taste or feeding themselves.

### Subsets of the *R70C07-GAL4* neuron population drive opposing feeding behaviors

We identified *R70C07-GAL4* to be a driver line of interest, as it showed the strongest feeding aversion upon neuronal activation (Figure 1B). Immunofluorescence of brains and ventral nerve cords (VNCs) from *R70C07-GAL4>CsChrimson* flies revealed a prominent set of 15 strongly labelled cell bodies on each side of the SEZ, with dense arborization across the medial and lateral SEZ (Figure 2A). Weaker and sparser projections were also observed in the antennal lobes and superior medial protocerebrum. Despite the absence of labellar and pharyngeal taste projections in this driver, stereotypical leg GRN projections were observed in the SEZ, as well as weak VNC processes, which could be contributing to the aversive feeding behavior observed in the STROBE (Stocker 1994). We retested *R70C07>CsChrimson* flies in the STROBE with the addition of 100 mM sucrose to both 1% agar options, which we have previously shown to enhance negative effects by increasing overall interaction numbers (Musso *et al*. 2019). This revealed intense aversion to the light-triggering side reminiscent of bitter GRN activation in the STROBE (Figure 2B,C)(Jaeger *et al*. 2018; Musso *et al*. 2019).

**Figure 2.**
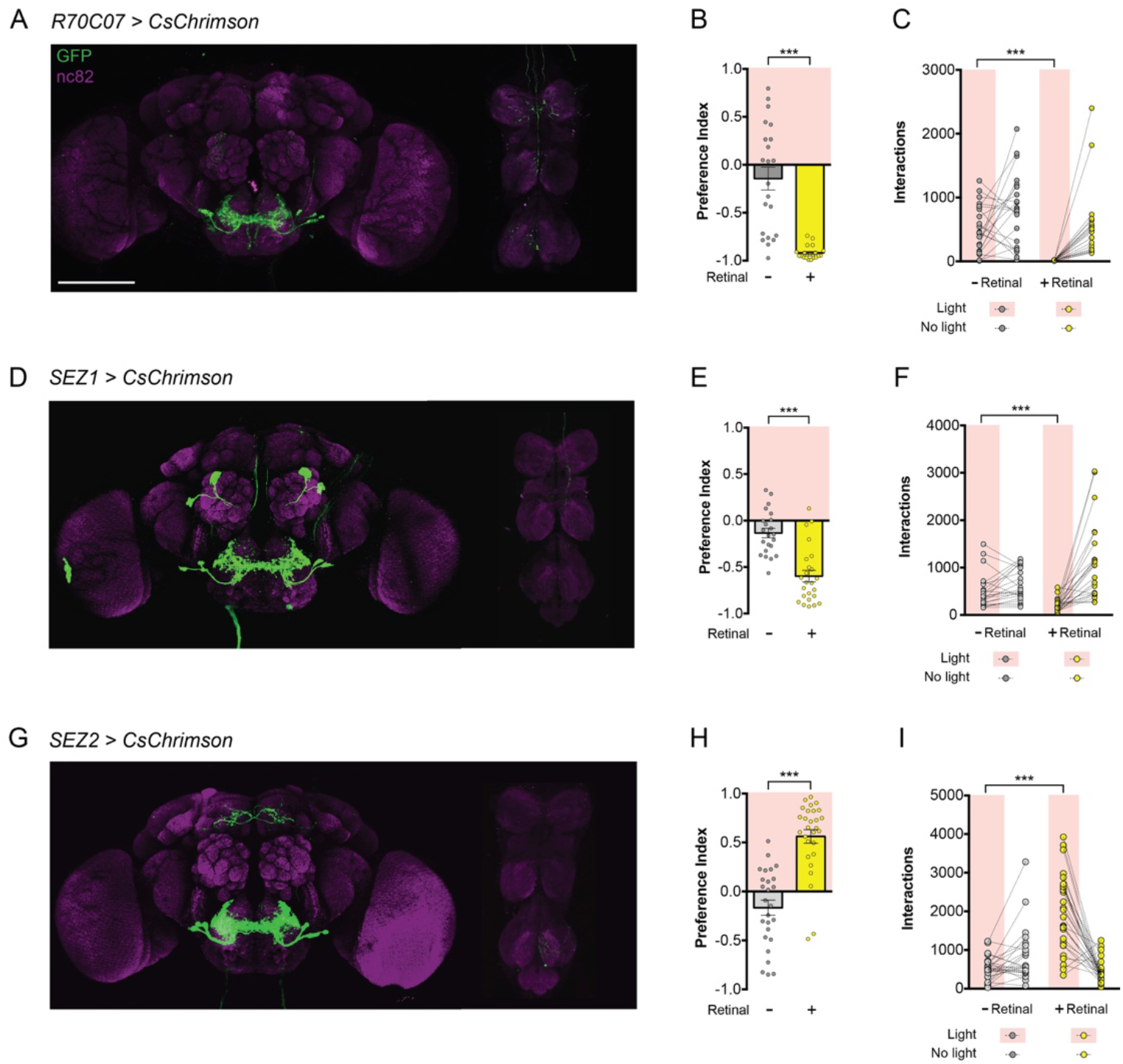
Subsets of the *R70C07-GAL4* population drive opposing feeding behaviors. (**A-C**) *R70C07-GAL4, UAS-CsChrimson*.*mVenus* expression in the brain and VNC (A), preference index in the STROBE containing 100 mM sucrose in 1% agar in both channels (B), and individual interaction numbers from the same experiment (C). (**D-F**) *SEZ1-GAL4* expression (D), preference index in the STROBE containing 100 mM sucrose in 1% agar in both channels (E), and interaction numbers (F). (**G-I**) *SEZ2-GAL4* expression (G), preference index in the STROBE containing 100 mM sucrose in 1% agar in both channels (H), and interaction numbers (I). For immunofluorescence (A,D,G) brains and VNC are stained for GFP (green) and nc82 (magenta). Scale bars, 100 µm. For preference indices (B,E,H), yellow bars represent the retinal fed group and grey bars represent the no retinal controls of the same genotype. Values are mean +/- SEM. For individual interaction numbers (C,F,I), lines connect values for individual flies. n = 21-32. Statistical tests: *t*-test, ***p < 0.001. Raw data is available in File S4.

To identify the specific neurons involved in producing feeding aversion, split-GAL4 lines (Luan *et al*.; Tirian and Dickson 2017; Dionne *et al*. 2018) were created by combining the *R70C07-p65*.*AD* hemidriver with various GAL4 DNA binding domain (DBD) hemidrivers that were selected based on putative expression in the SEZ neuron population (Figures 2 and 3). Leg projections and most VNC projections were successfully eliminated in *R70C07-p65*.*AD; R37H08-GAL4*.*DBD* (combination called *SEZ1-GAL4*) and *R70C07-p65*.*AD; R53C05-GAL4*.*DBD* (*SEZ2-GAL4*)(Figure 2D,G). Additionally, this intersectional refinement reduced the number of SEZ neurons from 15 per side in the original *R70C07-GAL4* driver to subsets of 3 and 7 in *SEZ1-GAL4* and *SEZ2-GAL4*, respectively. Light-activation of *SEZ1>CsChrimson* flies in the STROBE inhibited feeding similar to *R70C07-GAL4* (Figure 2E,F). The magnitude was slightly reduced, which could be explained by the elimination of either the leg inputs or subsets of SEZ neurons. Surprisingly, light-activation of *SEZ2-GAL4* produced the opposite effect by strongly promoting feeding (Figure H,I). We also identified one split-GAL4 combination (*SEZ3-GAL4*) that produced mild but significant feeding inhibition, and two (*SEZ4-GAL4*, and *SEZ5-GAL4*) that labeled subsets of the *R70C07* SEZ neurons but produced no significant behavioral effect in the STROBE, although both trended in the negative direction (Figure 3).

**Figure 3.**
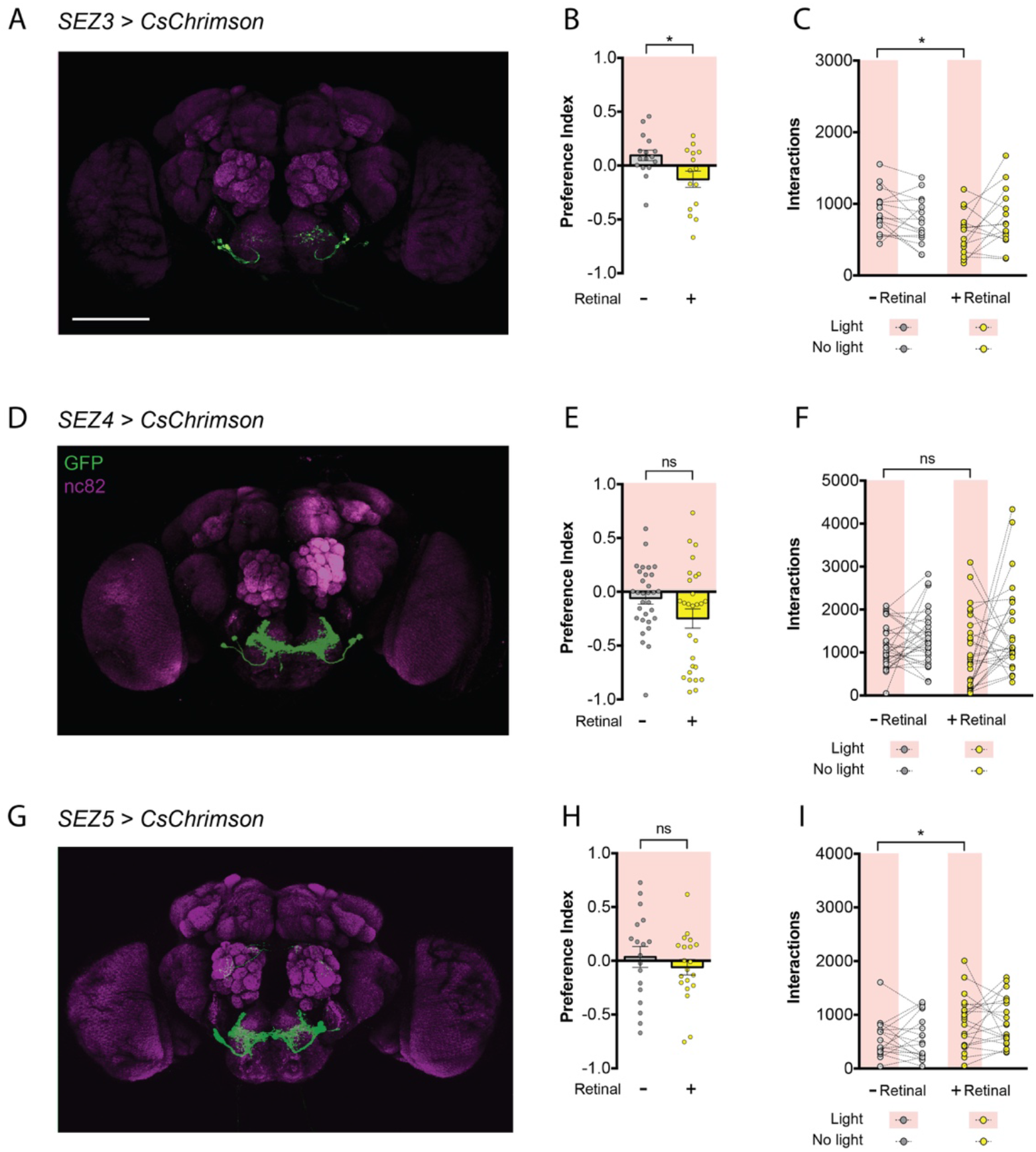
Not all *R70C07-GAL4* SEZ neurons are sufficient to alter feeding. (**A-C**) *SEZ3-GAL4* expression in the brain and VNC (A), preference index in the STROBE containing 100 mM sucrose in 1% agar in both channels (B), and individual interaction numbers from the same experiment (C). (**D-F**) *SEZ4-GAL4* expression (D), preference index in the STROBE containing 100 mM sucrose in 1% agar in both channels (E), and interaction numbers (F). (**G-I**) *SEZ5-GAL4* expression (G), preference index in the STROBE containing 100 mM sucrose in 1% agar in both channels (H), and interaction numbers (I). For immunofluorescence (A,D,G) brains and VNC are stained for GFP (green) and nc82 (magenta). Scale bars, 100 µm. For preference indices (B,E,H), yellow bars represent the retinal fed group and grey bars represent the no retinal controls of the same genotype. Values are mean +/- SEM. For individual interaction numbers (C,F,I), lines connect values for individual flies. n = 15-20. Statistical tests: *t*-test, *p < 0.05. Raw data is available in File S4.

There are two possible broad explanations for the phenotypes observed following split-GAL4 refinement. First, it is possible that the *R70C07* SEZ population comprises multiple neuron types with different behavioral effects. Perhaps *SEZ1-GAL4* isolated a predominantly negative set, while *SEZ2-GAL4* isolated a subset that was predominantly positive. This theory can be extended to suggest that the three split-GAL4 lines producing little or no effect labeled both positive and negative SEZ neurons that effectively cancelled each other out. The second possibility is that neurons outside the SEZ population affected preference in one or more of the split-GAL4 populations. Additional split-GAL4 lines that completely eliminate all expression outside the SEZ would be required to tease apart these possibilities. It is notable that both *SEZ1-GAL4* and *SEZ2-GAL4* labeled 1-2 neurons that were not clearly visible in the original *R70C07* driver. Whether this reflects different relative expression levels or differences between *R70C07-GAL4* and *R70C07-p65*.*AD* is unclear.

### Two distinct neuronal clusters make up the *R70C07* SEZ population

Closer examination of *R70C07-GAL4* revealed that the SEZ cluster is actually composed of two distinct clusters. One cluster of eight neurons on each side arborize medially and laterally within the SEZ. The other cluster is comprised of seven neurons with more anterior cell bodies and processes that project close to the antennal nerve into the posterior SEZ, where the arbors remain mostly lateral (Figure 4A). *SEZ1-, SEZ2-, SEZ4-*, and *SEZ5-GAL4* all label neurons from within the first cluster, while *SEZ3-GAL4* labels all seven of the neurons in the second cluster. (Figure 4B-D). Based on the mild phenotype from *SEZ3-GAL4* activation we suspect that the neurons in the more posterior-projecting cluster are not the primary drivers of *R70C07*-mediated feeding inhibition. However, we cannot rule out the possibility that lower expression levels in *SEZ3-GAL4* also contribute to its lesser effect.

**Figure 4.**
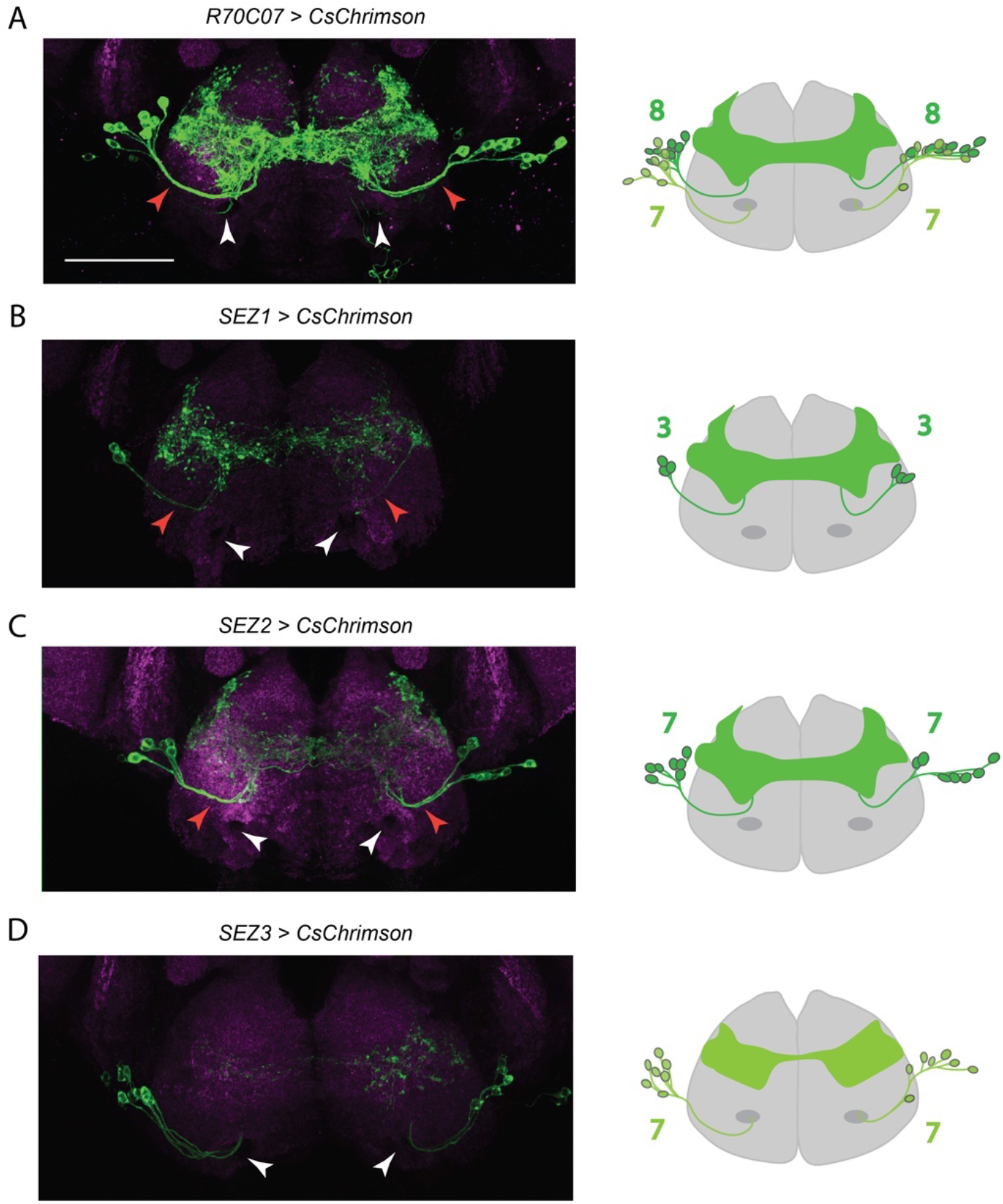
Two distinct neuronal clusters make up the *R70C07* SEZ population. Immunofluorescent detection of *UAS-CsChrimson*.*mVenus* (green) driven by *R70C07-GAL4* (A), *SEZ1-GAL4* (B), *SEZ2-GAL4* (C), and *SEZ3-GAL4* (D) in the SEZ with schematics on the right showing the number of neurons labelled by each split-GAL4. *SEZ1-GAL4* and *SEZ2-GAL4* label SEZ neurons that follow the same tract (red arrowhead). *SEZ3-GAL4* labels a distinct group of SEZ neurons that project more ventrally near the the labellar nerve tracts (white arrows). Counterstain is nc82 (magenta). All scale bar is 50 µm.

### Bitter and sweet sensory neurons GRASP with lateral SEZ neurons

We next wondered whether insight into the opposing behavioral effects of *SEZ1-GAL4* and *SEZ2-GAL4* could be gleaned from examining the synaptic inputs to these neurons. Thus, we used GFP-reconstitution across synaptic partners (GRASP) to test for contacts with sweet and bitter GRNs. One half of the split-GFP reporter (*lexAop-spGFP11*) was targeted to either the bitter- or sweet-sensitive GRNs using *Gr66a-LexA* or *Gr5a-LexA* as a driver; and the other half of the split-GFP reporter (*UAS-spGFP1-10*) was targeted to the lateral SEZ neurons with either *SEZ1-* (Figure 5A-D) or *SEZ2-GAL4* (Figure 5E-H). Unexpectedly, bitter and sweet GRASP signals were detected for both split-GAL4 lines, suggesting that bitter and sweet GRNs interact with at least one of the neurons labelled by each line. Notably, since Gr5a is not expressed in the pharyngeal sense organs, the strong GRASP signal with Gr5a GRNs suggests an interaction with inputs from the labellum, emphasizing the distinction between these neurons and the previously characterized and morphologically similar IN1 neurons (ref). Further analysis with calcium imaging will be necessary to determine the functional interactions between GRNs and the SEZ neurons.

**Figure 5.**
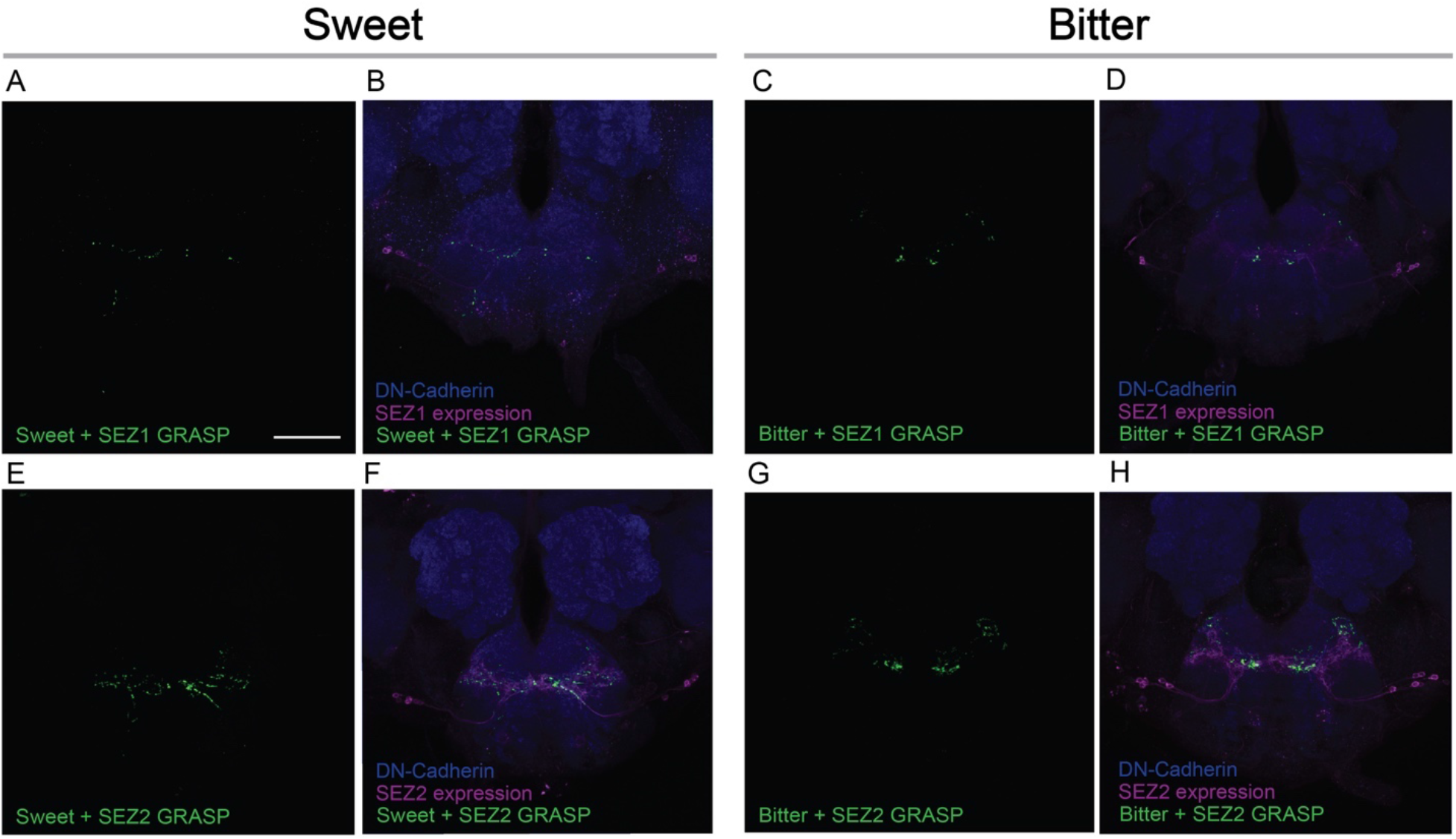
The *SEZ1-GAL4* and *SEZ2-GAL4* populations both contact sweet and bitter GRNs. (**A-D**) GRASP between *SEZ1-GAL4* and sweet (A,B) or bitter (C,D) GRNs. (**E-H**) GRASP between *SEZ2-GAL4* and sweet (E,F) or bitter (G,H) GRNs. Antibodies: anti-DN-Cadherin for counterstain (blue), anti-DsRed for GAL4 expression (magenta), anti-GFP used for GRASP signal (green). Scale bar is 50 µm.

Although the evidence that the *R70C07* SEZ represent bona fide second-order taste neurons is incomplete, neurons that appear very similar were previously identified as postsynaptic to sweet GRNs using the trans-synaptic tracer trans-Tango (Talay *et al*. 2017). The split-GAL4 lines identified in our study should greatly aid in more fully characterizing the functional properties of these neurons, including their inputs and post-synaptic targets. We also anticipate that the other lines identified in our behavioral screen will serve as useful starting points in the long-term prospect of more fully understanding the neural control of feeding behavior in flies.

## Acknowledgements

We thank Pierre Junca and Pierre-Yves Musso for help in identifying split-GAL4 hemidrivers, and members of the Gordon lab for comments on the manuscript. We also thank the Bloomington stock center for flies and the Janelia Flylight project for generating the lines screened and for the expression patterns shown in supplementary file S1. This work was funded by the Canadian Institutes of Health Research (CIHR) operating grant FDN-148424, with infrastructure funded by the Canadian Foundation for Innovation (CFI) grant 27290. M.D.G. is a Michael Smith Foundation for Health Research Scholar.

## Supplementary materials

**File S1. Full screen dataset in graphical form**. For each line tested in the screen, the following data is displayed: genotype and expression pattern taken from FlyLight database (Jenett *et al*. 2012); endpoint and time course values for the preference index; interaction numbers displayed for individual flies across each channel, as means for each channel, as pooled data across both channels, and as cumulative average over time. For all graphs, yellow indicates the experimental (retinal fed) group and grey indicates the control (no retinal). Bar graphs and time curves represent mean +/- SEM. Red shading denotes light-triggering channel for preference indices, time curves, and interaction numbers. For total interaction numbers, the data from individual flies are connected by lines. A key to interpreting the data is presented on the first page.

**File S2. Raw interaction numbers from screen**. Each tab holds the data for an individual genotype. Raw interaction counts are shown from the endpoint of each experiment (summary table) and as cumulative counts over time for each replicate.

**File S3. Full dataset preference indices from screen**. Each tab holds the data for an individual genotype. Preference indices are shown from the endpoint of each experiment (summary table) and as cumulative preference over time for each replicate.

**File S4. Raw data for figures 2 and 3**.

