## Supplementary material for "A closed-loop optogenetic screen for neurons controlling feeding in *Drosophila*": File S1

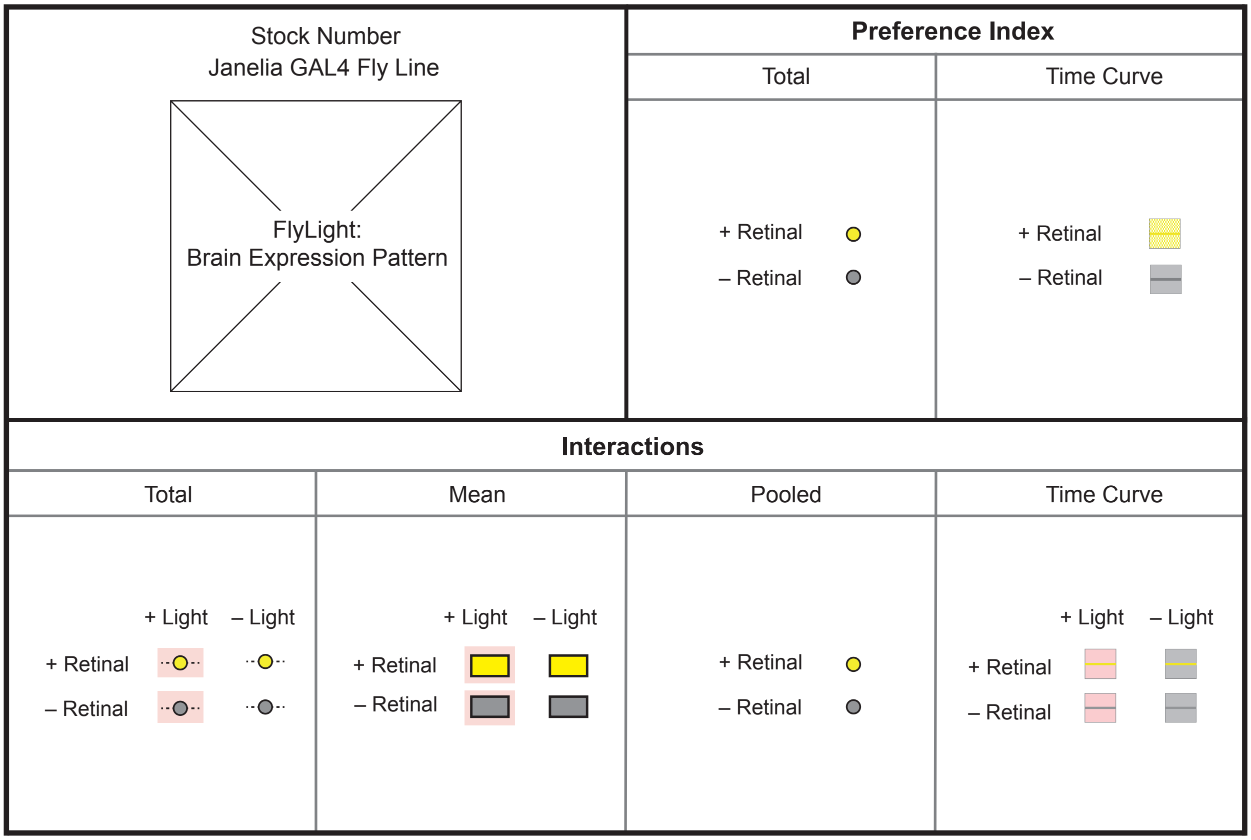

Stock Number
Janelia GAL4 Fly Line
FlyLight:
Brain Expression Pattern
Preference Index
Total
Time Curve
+ Retinal
– Retinal
+ Retinal
– Retinal
Interactions
Total
Mean
Pooled
Time Curve
+ Light
– Light
+ Retinal
– Retinal
+ Light
– Light
+ Retinal
– Retinal
+ Retinal
– Retinal
+ Light
– Light
+ Retinal
– Retinal

40730

w[1118];P{y[+t7.7] w[+mC]=**GMR9D10-GAL4**}attP2

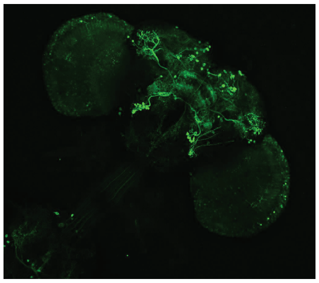

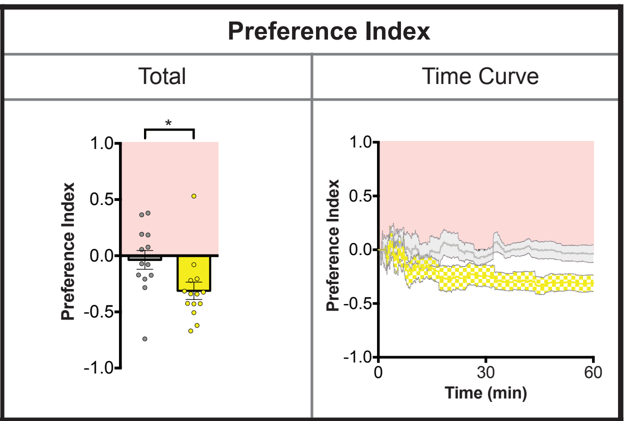


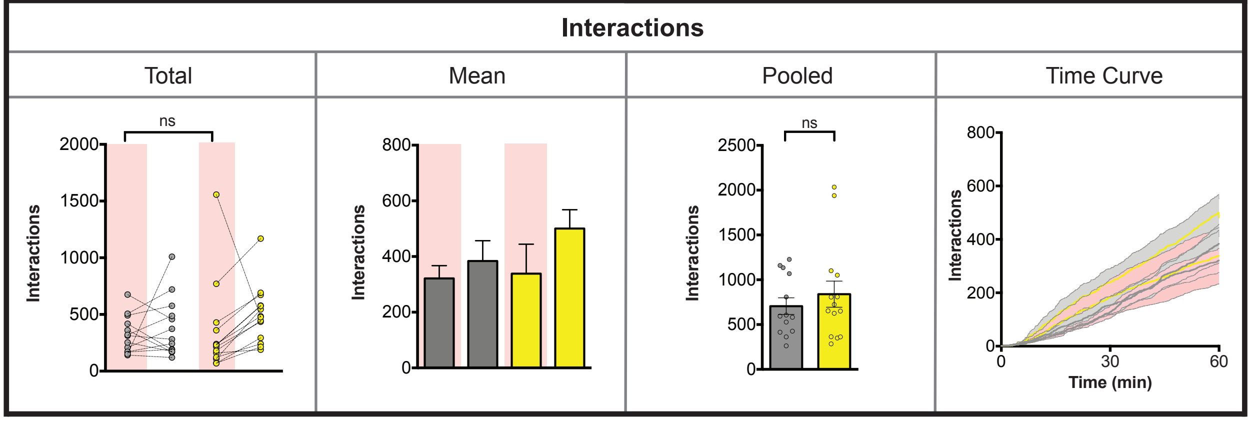


40731

w[1118]; P{y[+t7.7] w[+mC]=**GMR9D11-GAL4**}attP2

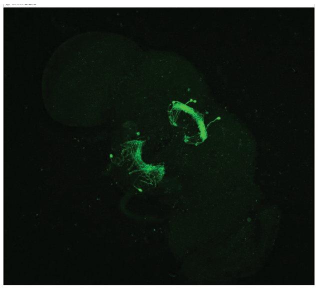

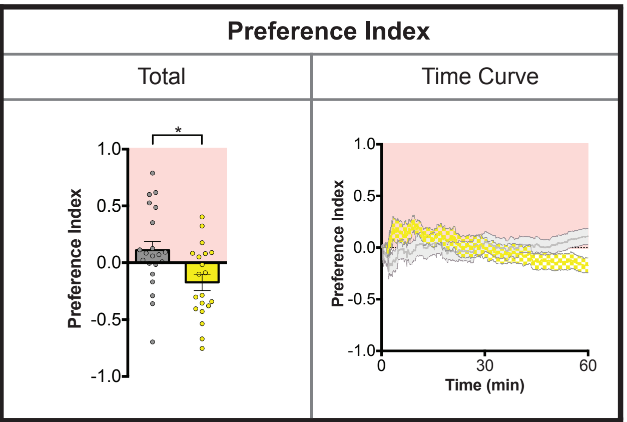


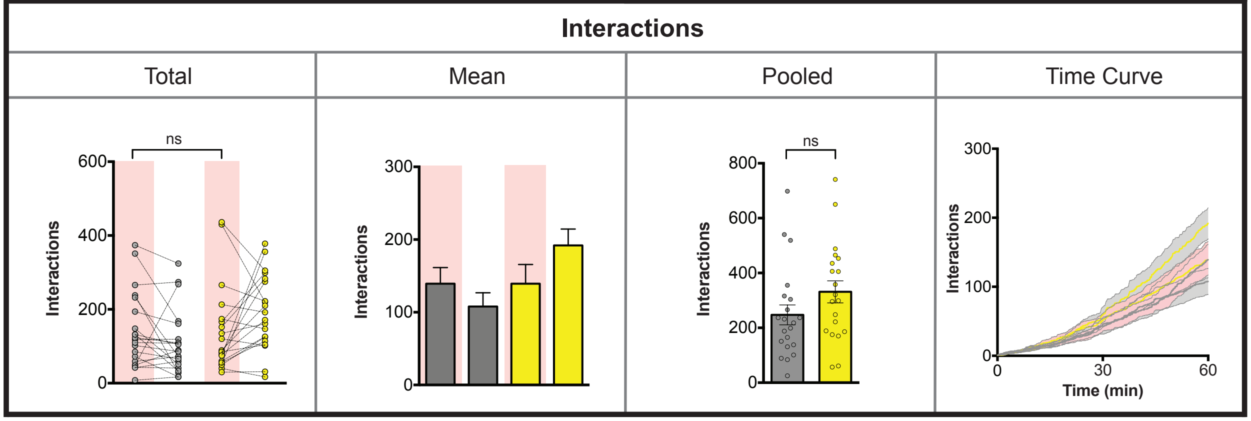

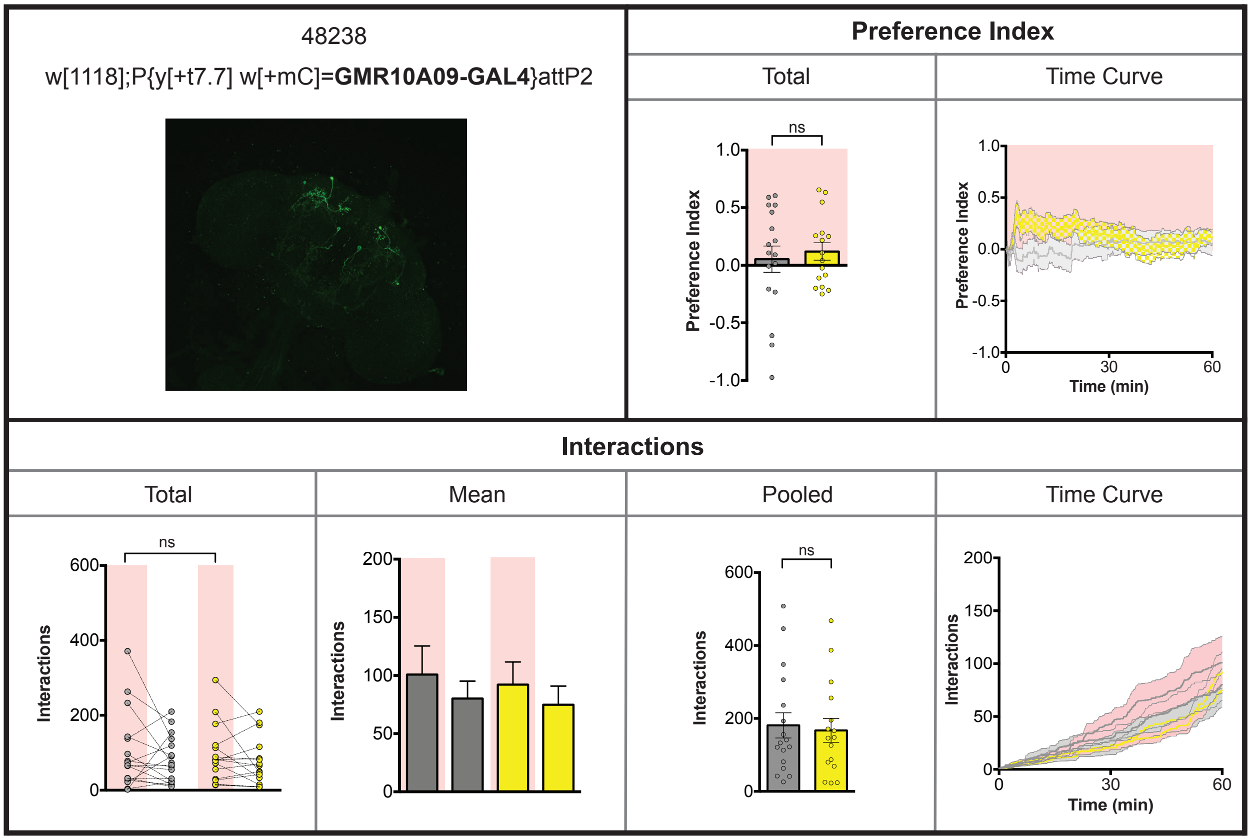
48238
w[1118];P{y[+t7.7] w[+mC]=**GMR10A09-GAL4**}attP2
Preference Index
Total
Time Curve
ns
Preference Index
1.0
0.5
0.0
-0.5
-1.0
Time (min)
0
30
60
Interactions
Total
Mean
Pooled
Time Curve
ns
Interactions
600
400
200
0
200
150
100
50
0
ns
600
400
200
0
200
150
100
50
0
Time (min)
0
30
60

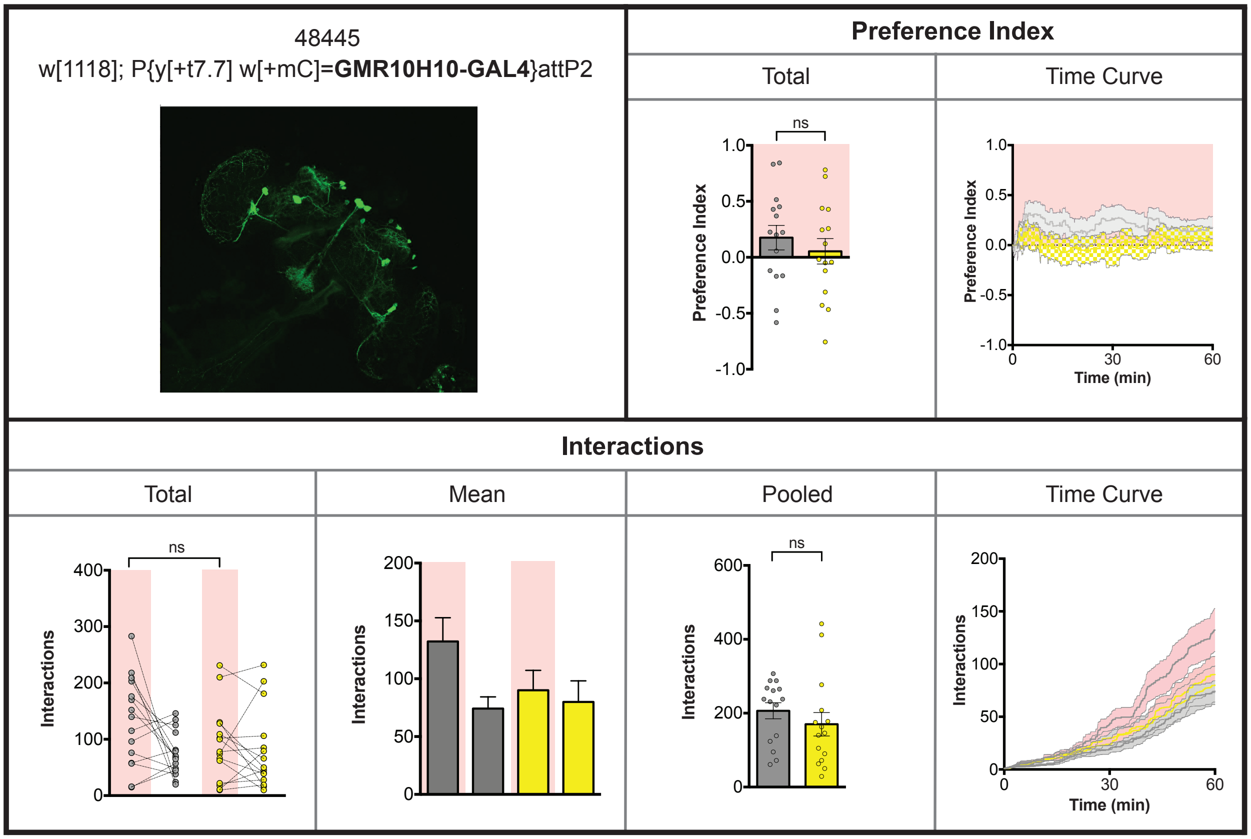
48445
w[1118]; P{y[+t7.7] w[+mC]=**GMR10H10-GAL4**}attP2
Preference Index
Total
Time Curve
ns
Preference Index
1.0
0.5
0.0
-0.5
-1.0
Time (min)
0
30
60
Interactions
Total
Mean
Pooled
Time Curve
ns
Interactions
400
300
200
100
0
200
150
100
50
0
ns
600
400
200
0
200
150
100
50
0
Time (min)
0
30
60

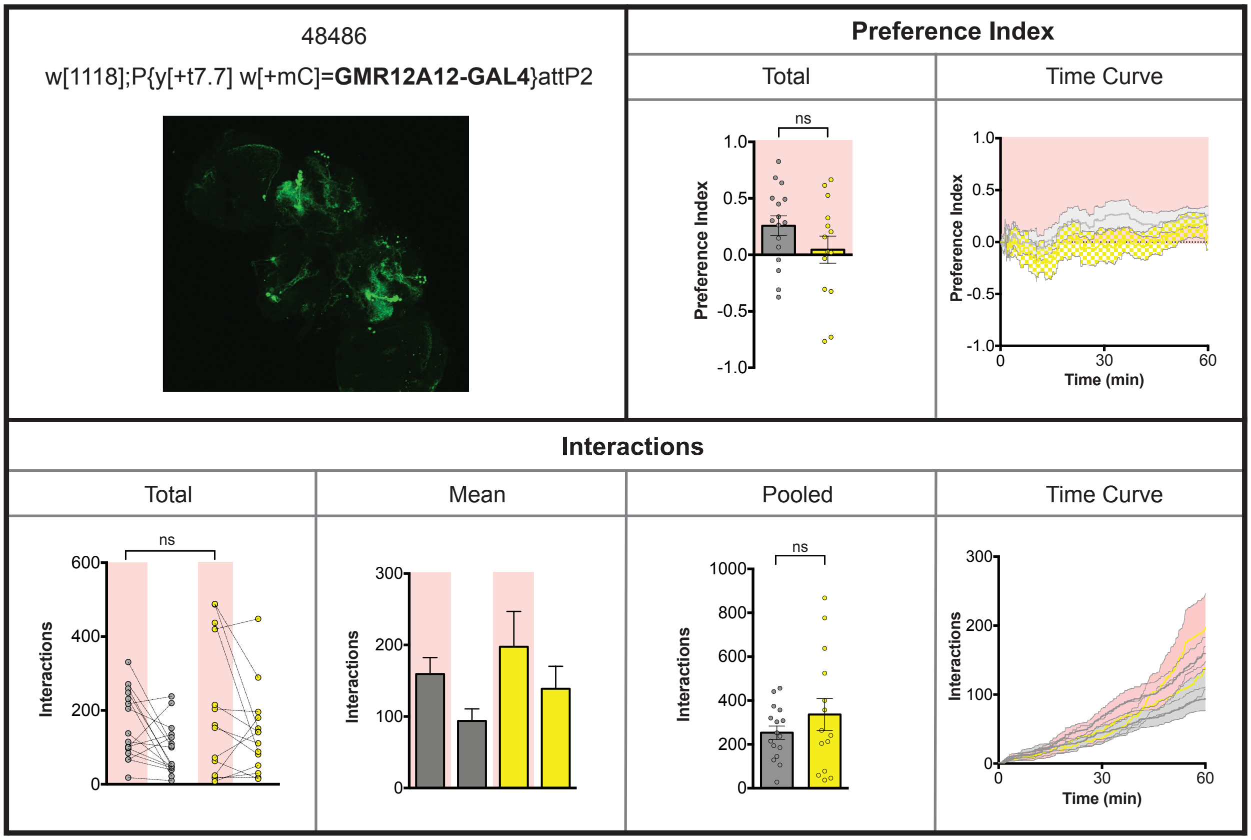
48486
w[1118];P{y[+t7.7] w[+mC]=**GMR12A12-GAL4**}attP2
Preference Index
Total
Time Curve
ns
Preference Index
1.0
0.5
0.0
-0.5
-1.0
0
30
60
Time (min)
Interactions
Total
Mean
Pooled
Time Curve
ns
Interactions
600
400
200
0
300
200
100
0
ns
1000
800
600
400
200
0
300
200
100
0
30
60
Time (min)

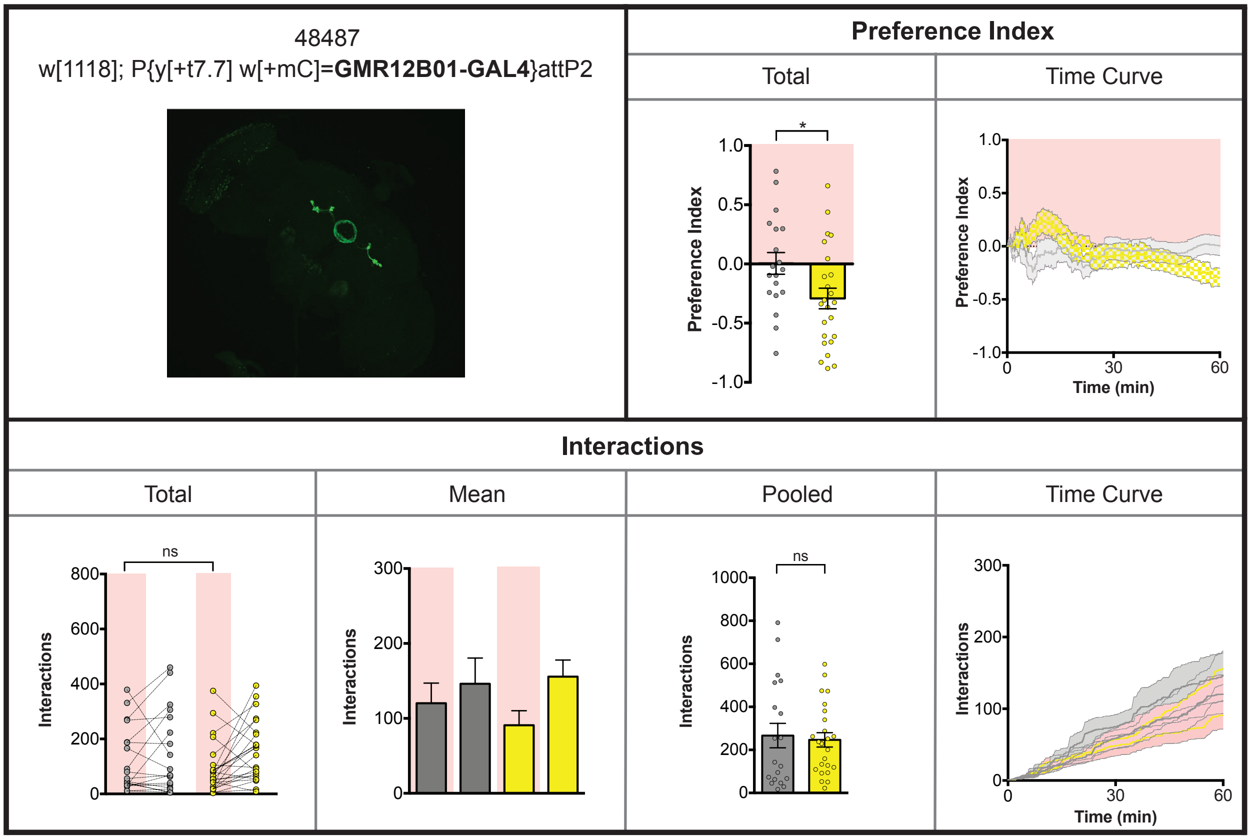
48487
w[1118]; P{y[+t7.7] w[+mC]=**GMR12B01-GAL4**}attP2
Preference Index
Total
Time Curve
*
Preference Index
1.0
0.5
0.0
-0.5
-1.0
0
30
60
Time (min)
Interactions
Total
Mean
Pooled
Time Curve
ns
Interactions
800
600
400
200
0
300
200
100
0
ns
1000
800
600
400
200
0
300
200
100
0
30
60
Time (min)

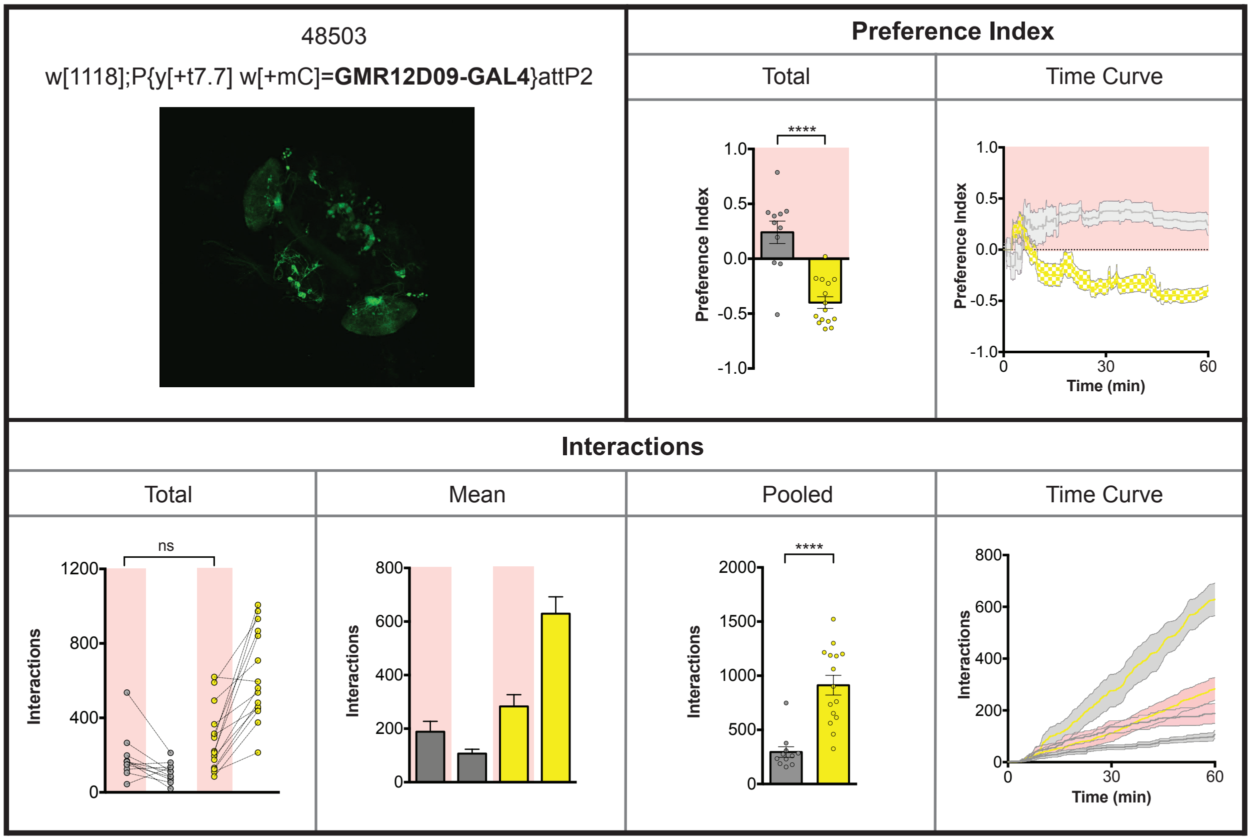
48503
w[1118];P{y[+t7.7] w[+mC]=**GMR12D09-GAL4**}attP2
Preference Index
Total
Time Curve
****
Preference Index
1.0
0.5
0.0
-0.5
-1.0
0
30
60
Time (min)
Interactions
Total
Mean
Pooled
Time Curve
ns
1200
800
400
0
600
200
****
2000
1500
1000
500
Interactions

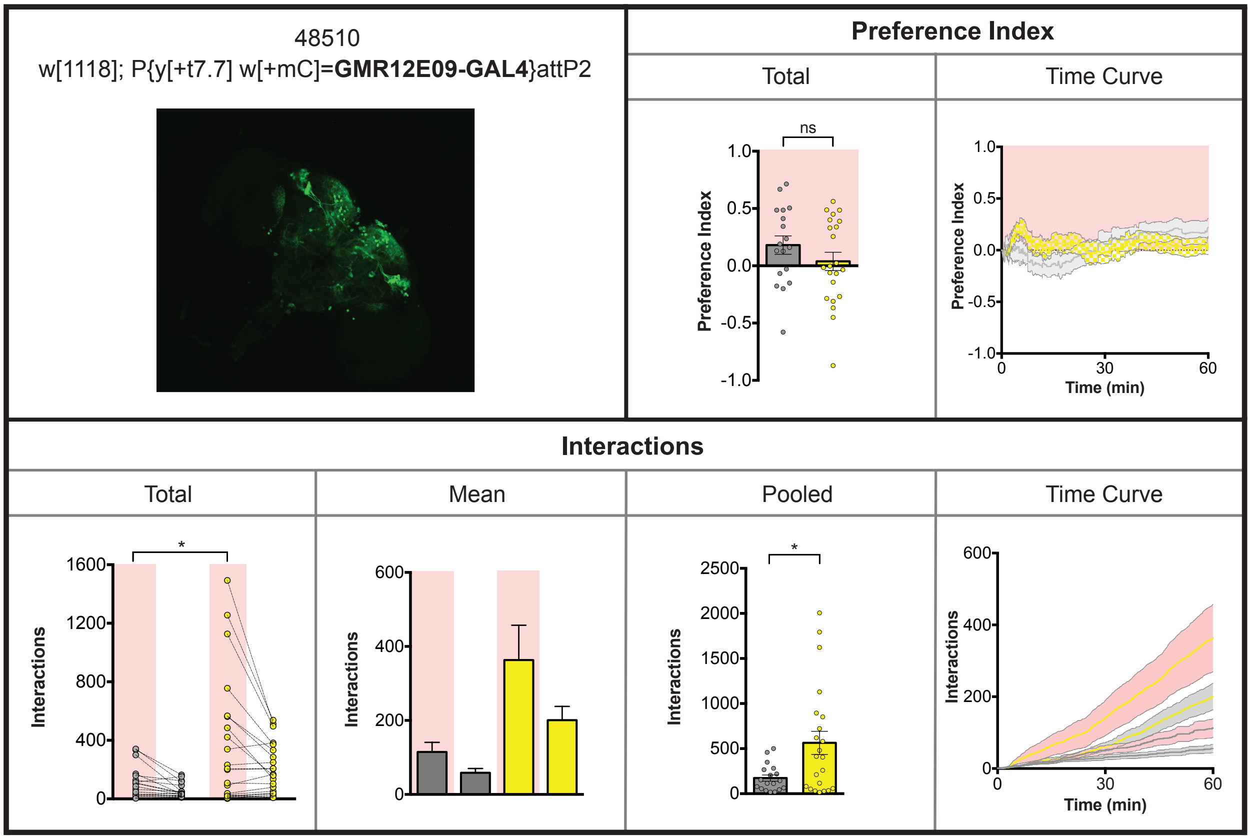
48510
w[1118]; P{y[+t7.7] w[+mC]=**GMR12E09-GAL4**}attP2
Preference Index
Total
Time Curve
ns
Preference Index
1.0
0.5
0.0
-0.5
-1.0
0
30
60
Time (min)
Interactions
Total
Mean
Pooled
Time Curve
*
1600
1200
800
400
0
600
200
2500
2000
1500
1000
500
Interactions

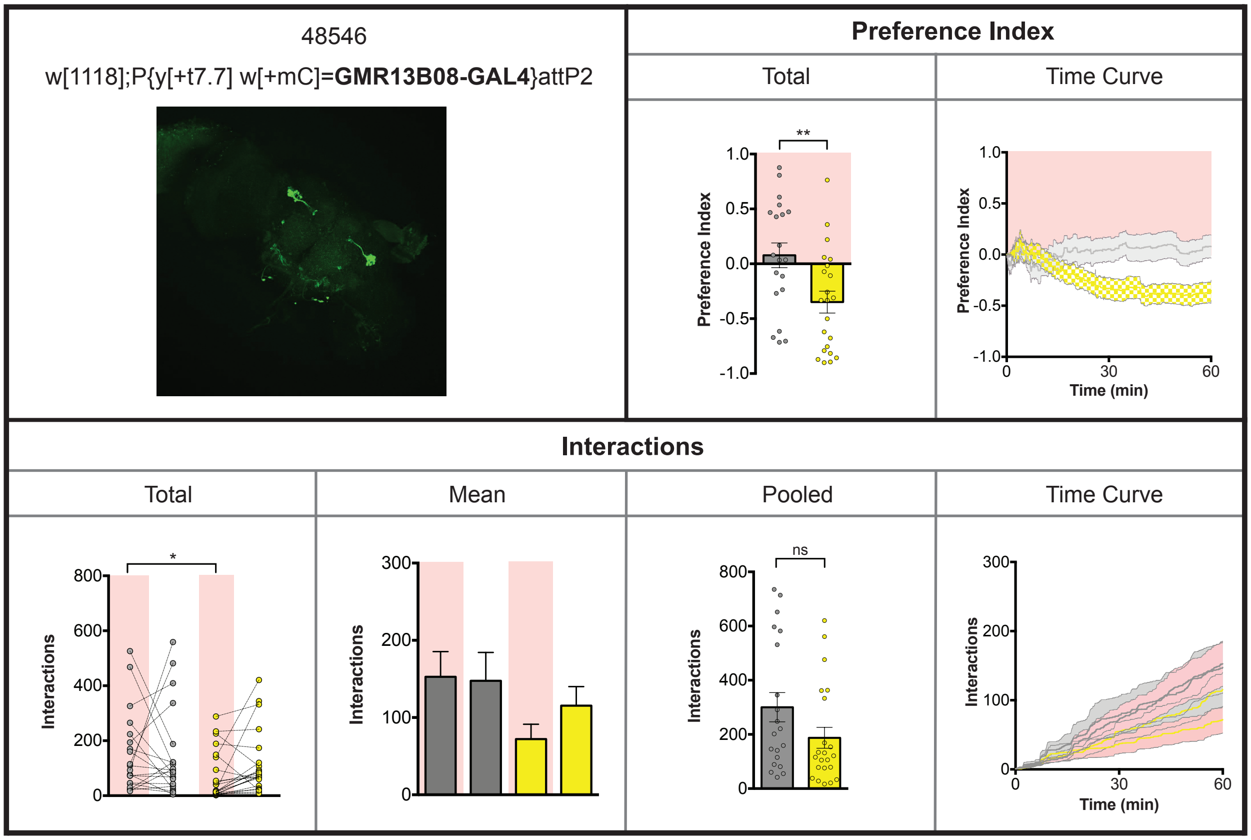
48546
w[1118];P{y[+t7.7] w[+mC]=GMR13B08-GAL4}attP2
Preference Index
Total
Time Curve
**
Preference Index
1.0
0.5
0.0
-0.5
-1.0
0
30
60
Time (min)
Interactions
Total
Mean
Pooled
Time Curve
*
ns
800
600
400
200
0
300
200
100
Interactions

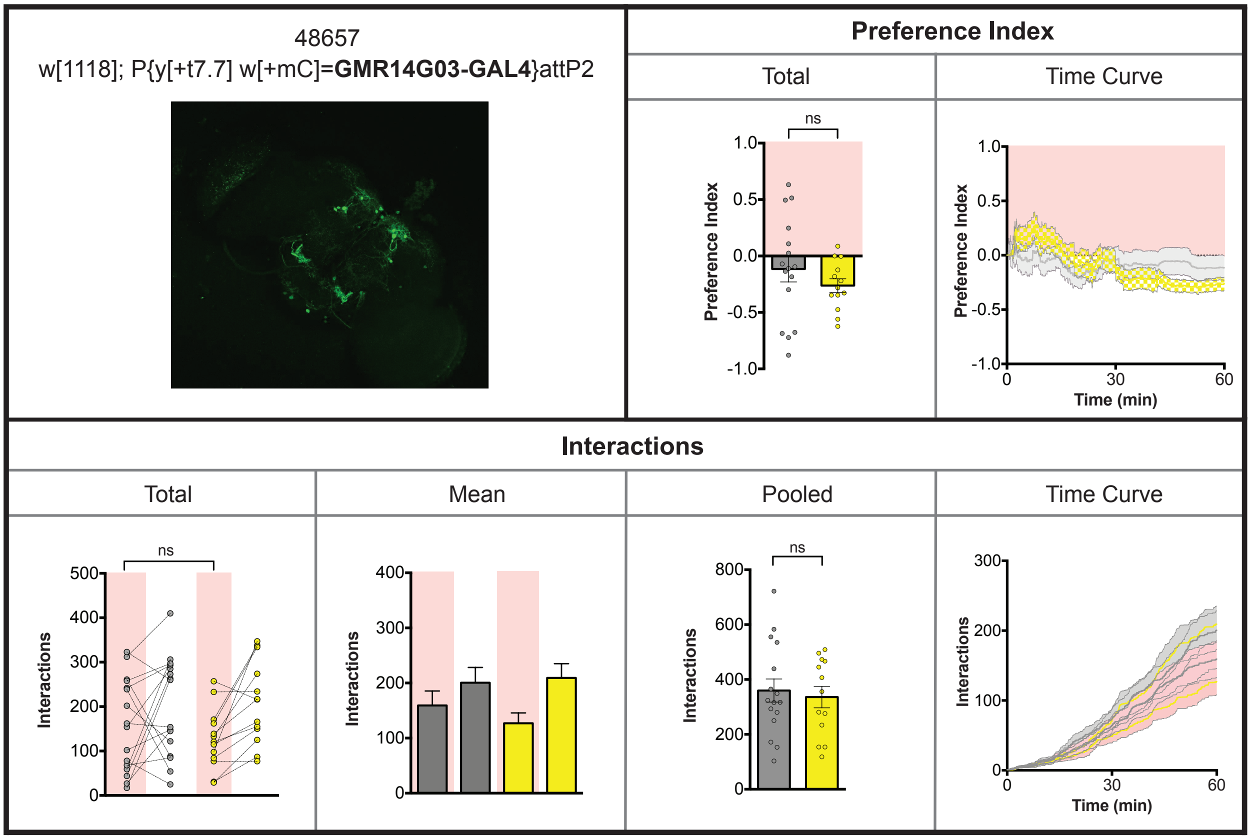
48657
w[1118]; P{y[+t7.7] w[+mC]=GMR14G03-GAL4}attP2
Preference Index
Total
Time Curve
ns
Preference Index
1.0
0.5
0.0
-0.5
-1.0
0
30
60
Time (min)
Interactions
Total
Mean
Pooled
Time Curve
ns
ns
500
400
300
200
100
0
800
600
Interactions

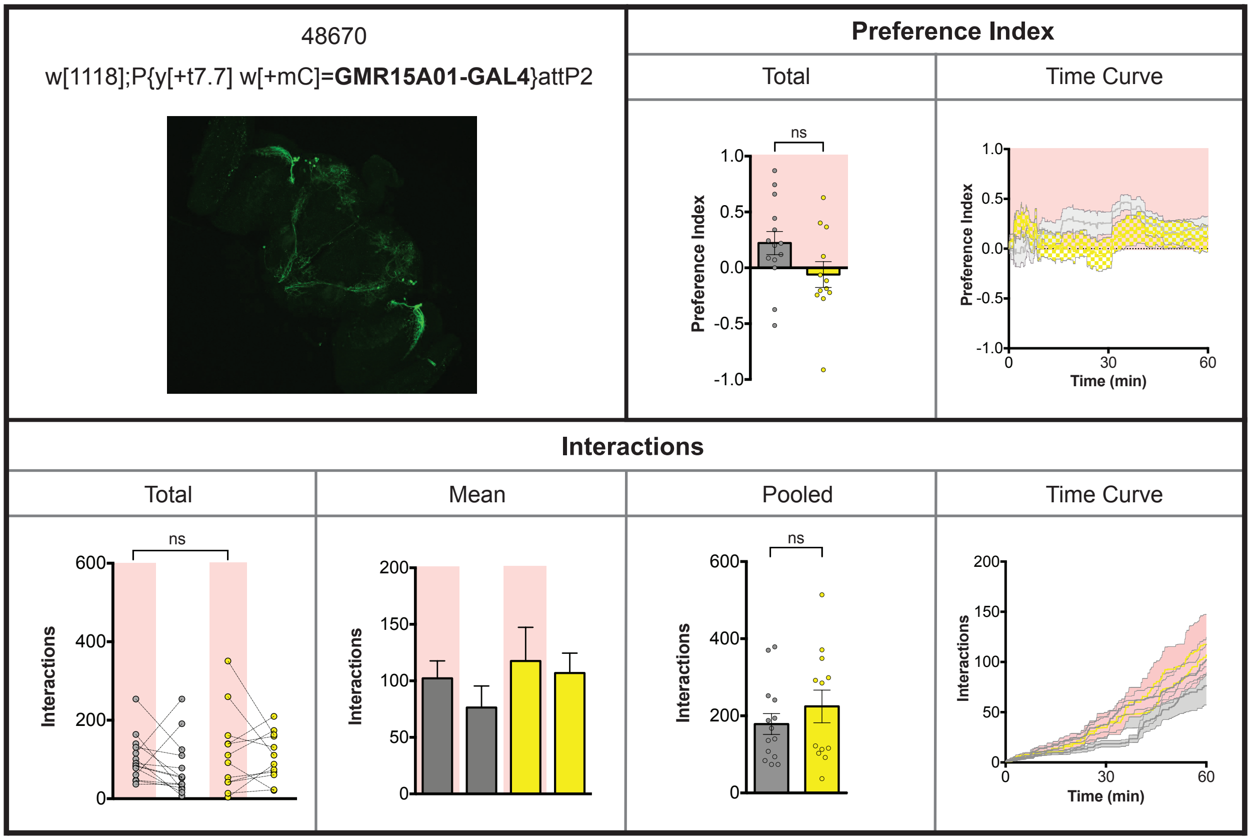
48670
w[1118];P{y[+t7.7] w[+mC]=**GMR15A01-GAL4**}attP2
Preference Index
Total
Time Curve
ns
Preference Index
Time (min)
Interactions
Total
Mean
Pooled
Time Curve
ns
ns
Interactions
Time (min)

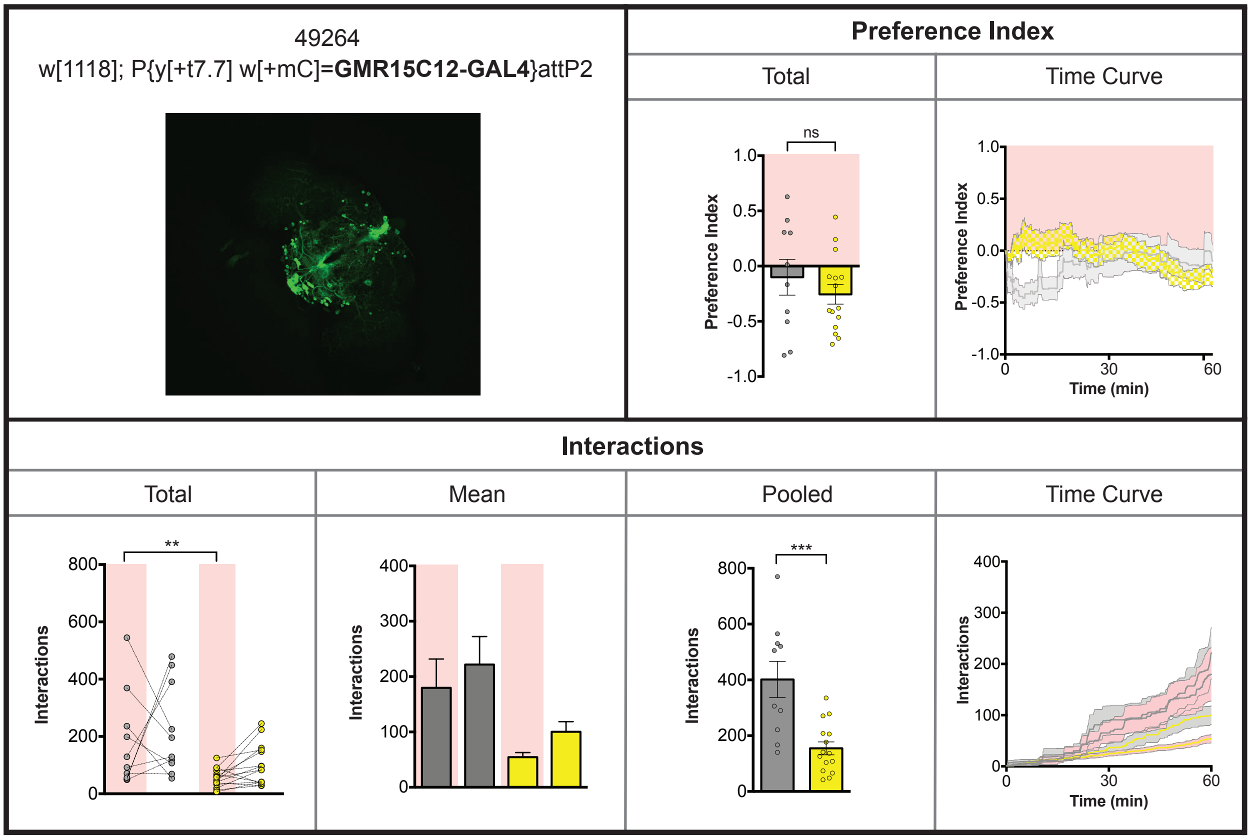
49264
w[1118]; P{y[+t7.7] w[+mC]=**GMR15C12-GAL4**}attP2
Preference Index
Total
Time Curve
ns
Preference Index
Time (min)
Interactions
Total
Mean
Pooled
Time Curve
**
***
Interactions
Time (min)

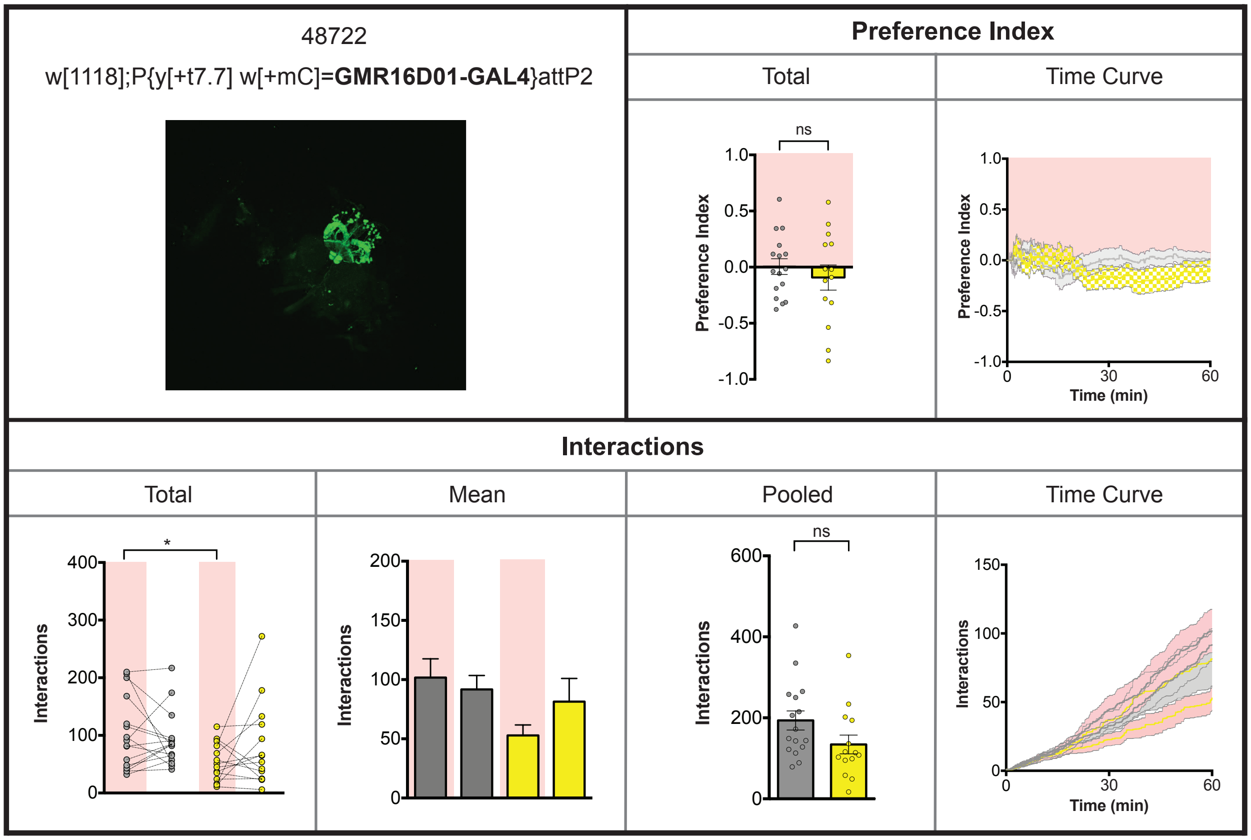
48722
w[1118];P{y[+t7.7] w[+mC]=GMR16D01-GAL4}attP2
Preference Index
Total
Time Curve
ns
Preference Index
Time (min)
Interactions
Total
Mean
Pooled
Time Curve
*
ns
Interactions
Time (min)

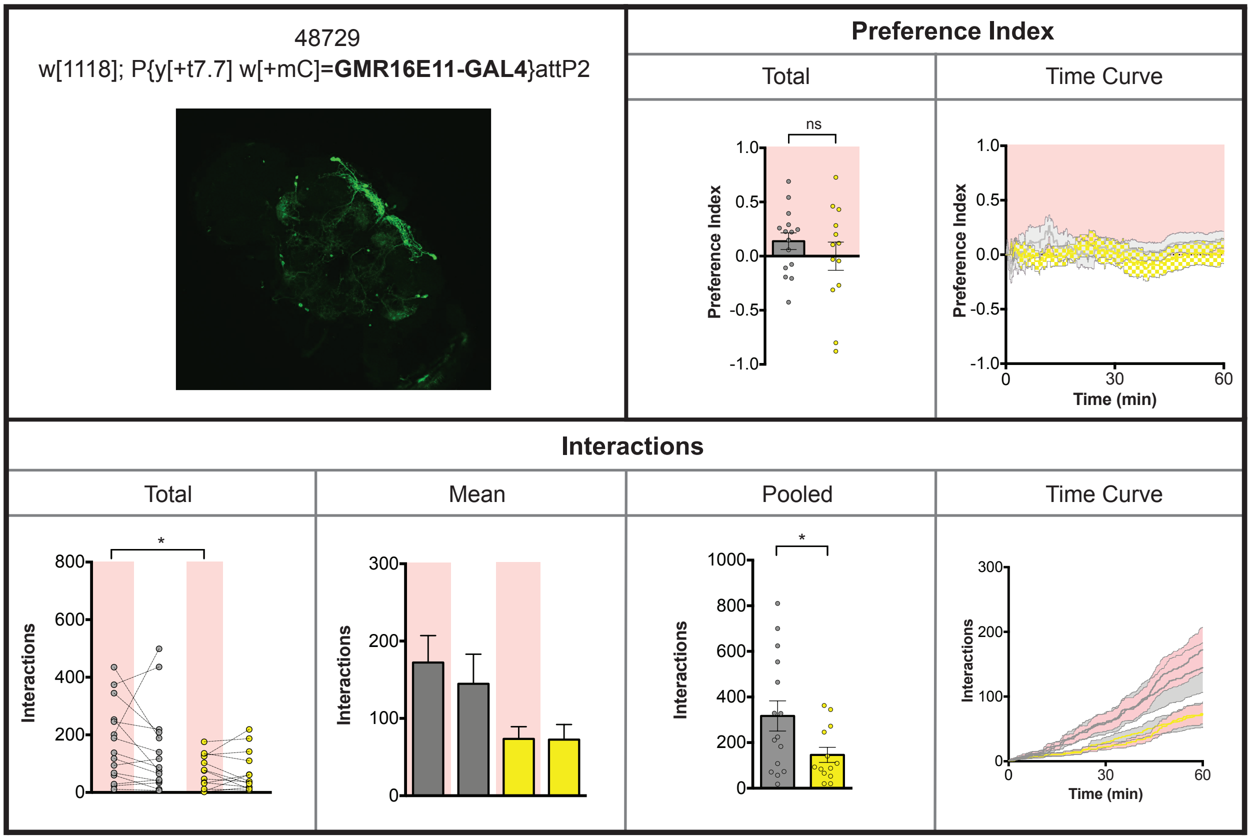
48729
w[1118]; P{y[+t7.7] w[+mC]=GMR16E11-GAL4}attP2
Preference Index
Total
Time Curve
ns
Preference Index
Time (min)
Interactions
Total
Mean
Pooled
Time Curve
*
*
Interactions
Time (min)

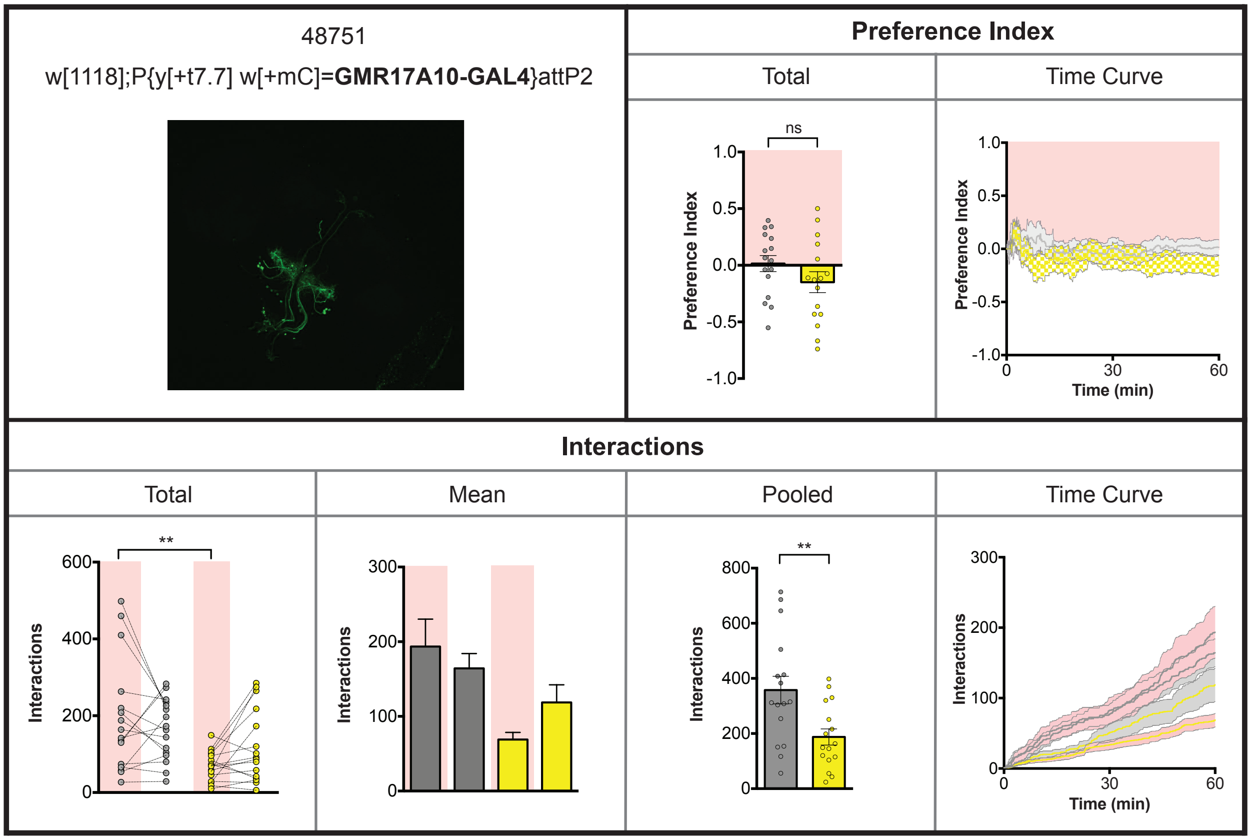
48751
w[1118];P{y[+t7.7] w[+mC]=GMR17A10-GAL4}attP2
Preference Index
Total
Time Curve
ns
Preference Index
1.0
0.5
0.0
-0.5
-1.0
Time (min)
0
30
60
Interactions
Total
Mean
Pooled
Time Curve
**
Interactions
600
400
200
0
300
200
100
0
800
600
400
200
0
**
Time (min)

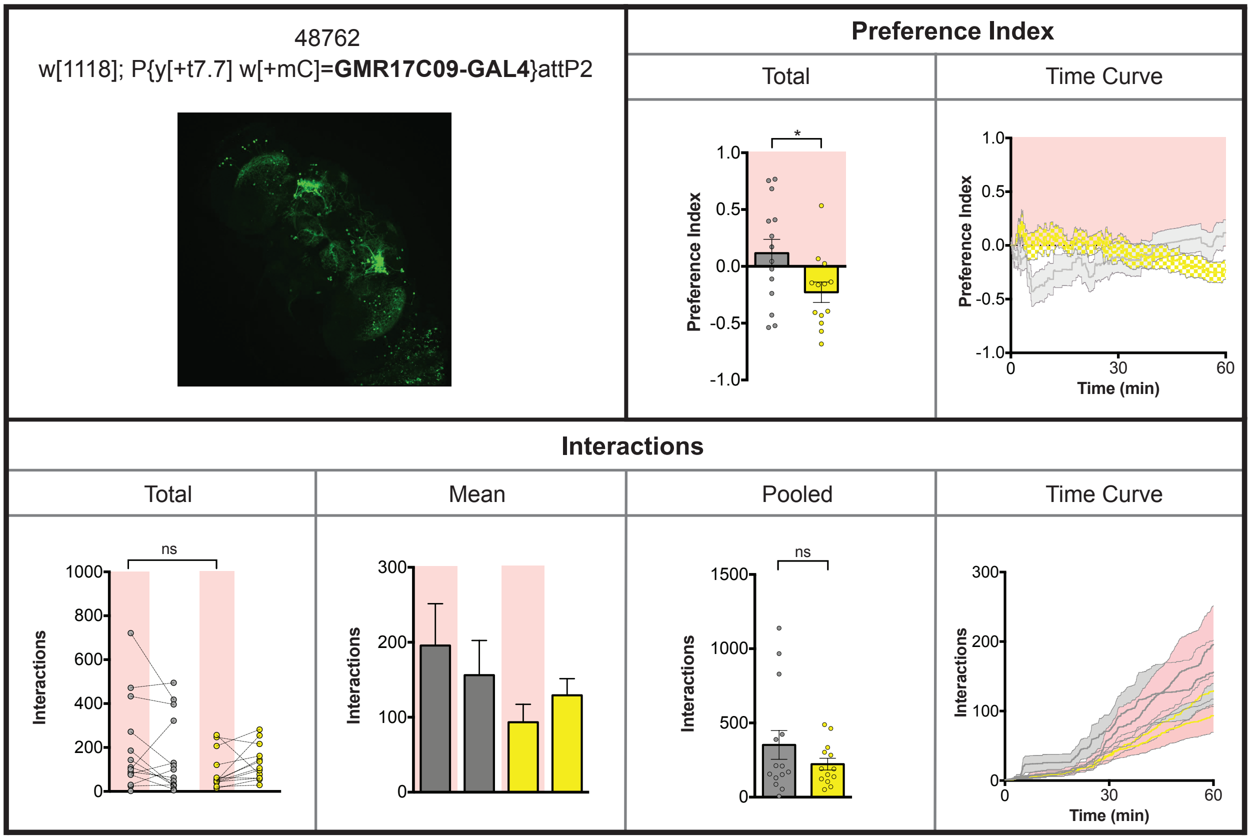
48762
w[1118]; P{y[+t7.7] w[+mC]=GMR17C09-GAL4}attP2
Preference Index
Total
Time Curve
*
Preference Index
1.0
0.5
0.0
-0.5
-1.0
Time (min)
0
30
60
Interactions
Total
Mean
Pooled
Time Curve
ns
Interactions
1000
800
600
400
200
0
300
200
100
0
1500
1000
500
0
ns
Time (min)

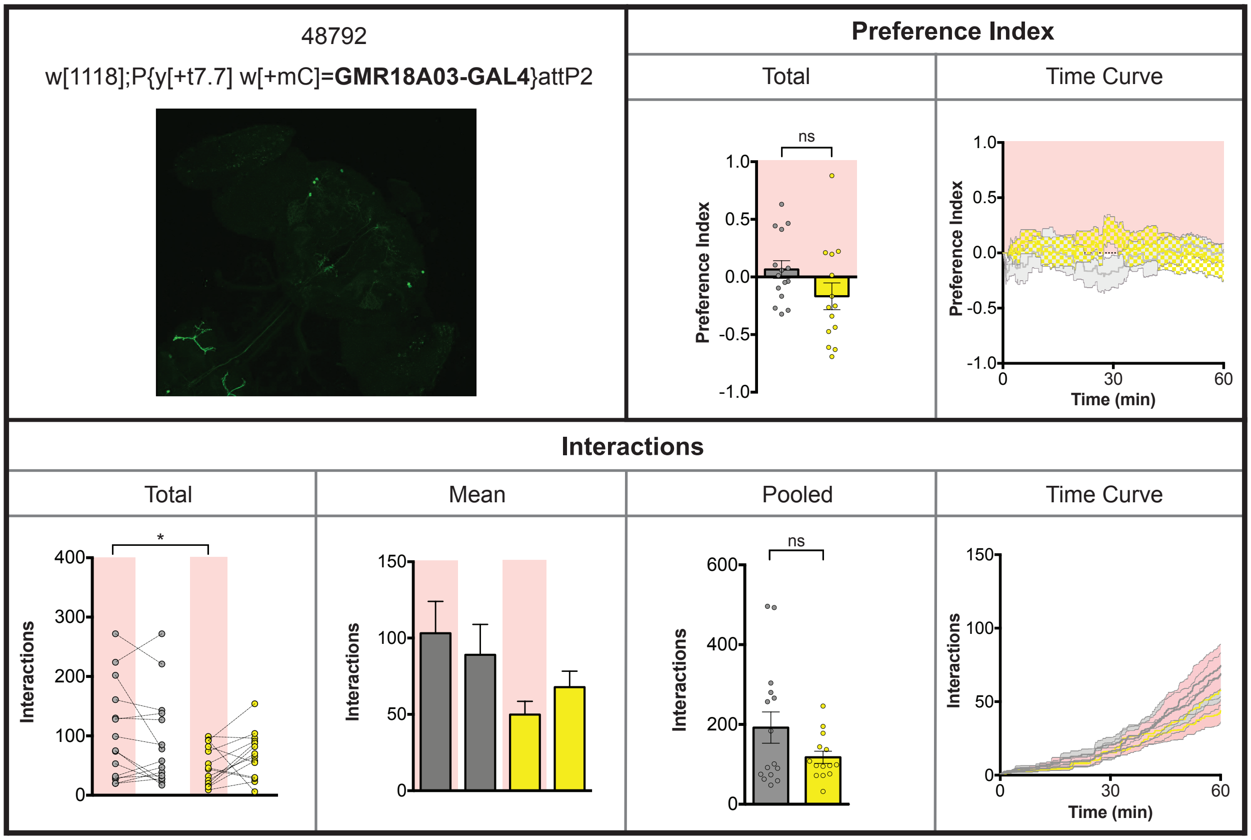
48792
w[1118];P{y[+t7.7] w[+mC]=GMR18A03-GAL4}attP2
Preference Index
Total
Time Curve
ns
Preference Index
Time (min)
Interactions
Total
Mean
Pooled
Time Curve
*
ns
Interactions
Time (min)

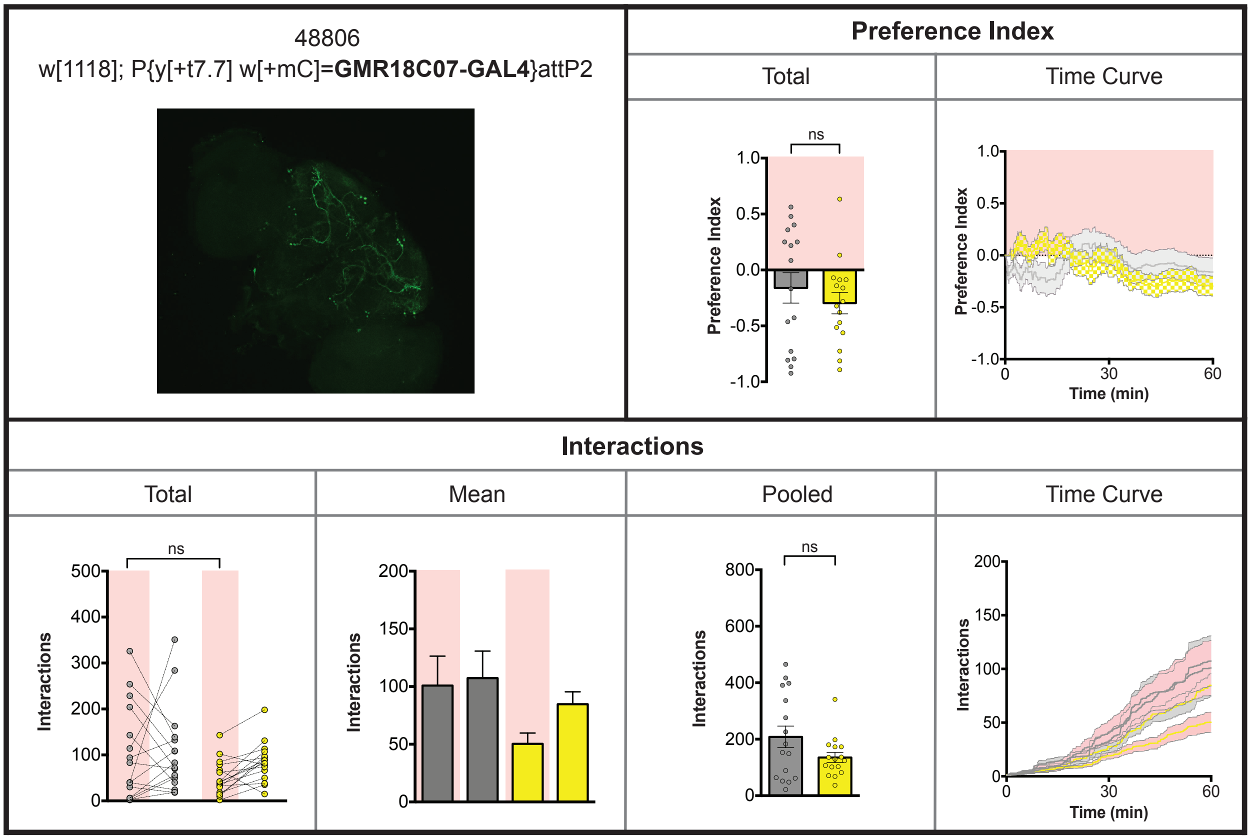
48806
w[1118]; P{y[+t7.7] w[+mC]=GMR18C07-GAL4}attP2
Preference Index
Total
Time Curve
ns
Preference Index
Time (min)
Interactions
Total
Mean
Pooled
Time Curve
ns
ns
Interactions
Time (min)

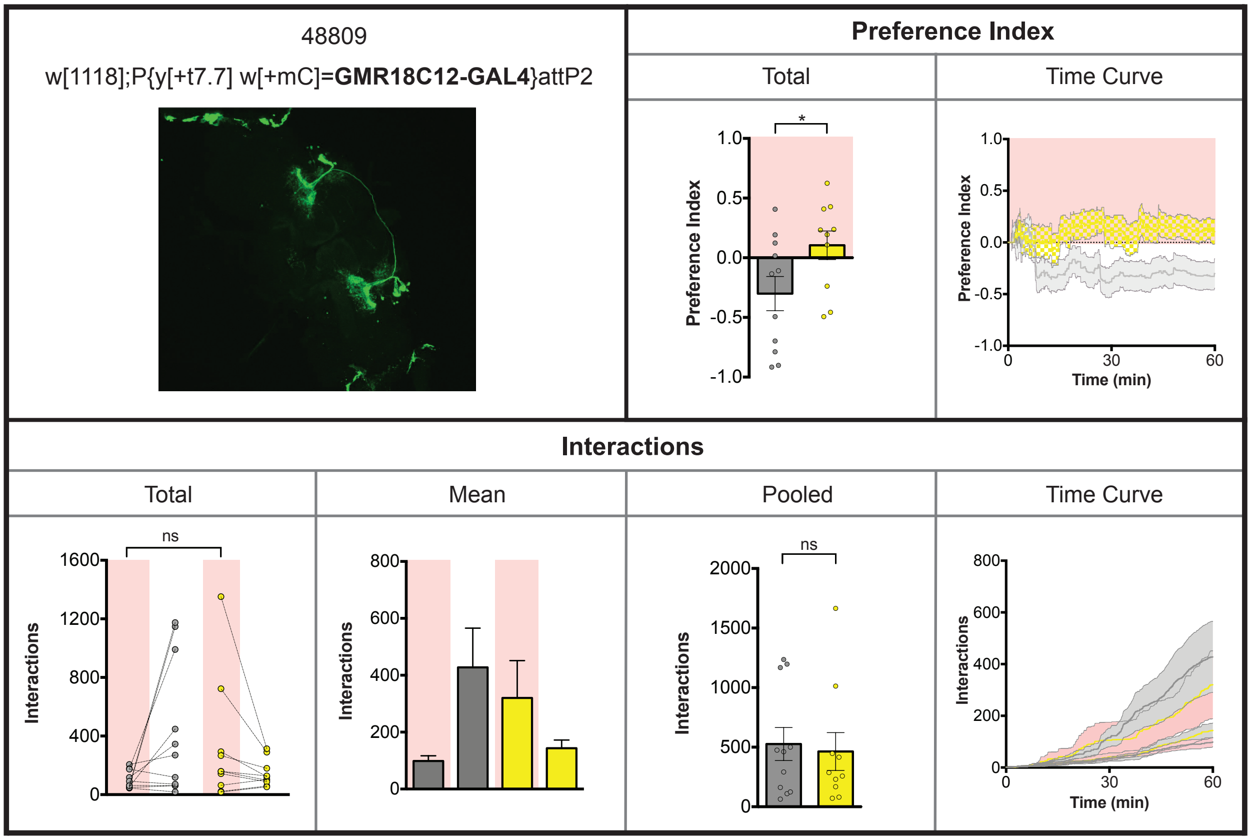
48809
w[1118];P{y[+t7.7] w[+mC]=GMR18C12-GAL4}attP2
Preference Index
Total
Time Curve
Preference Index
Time (min)
*
Interactions
Total
Mean
Pooled
Time Curve
ns
ns
Interactions
Time (min)

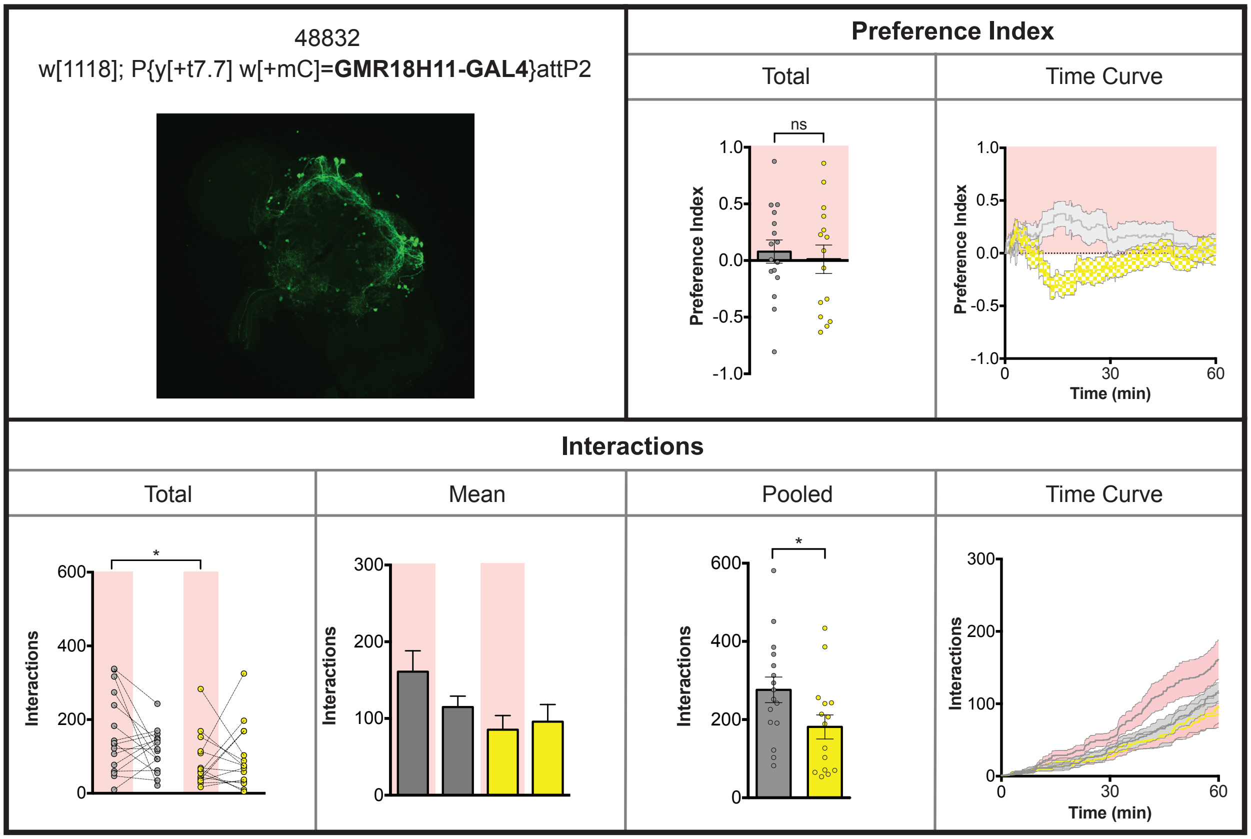
48832
w[1118]; P{y[+t7.7] w[+mC]=GMR18H11-GAL4}attP2
Preference Index
Total
Time Curve
ns
Preference Index
Time (min)
Interactions
Total
Mean
Pooled
Time Curve
*
*
Interactions
Time (min)

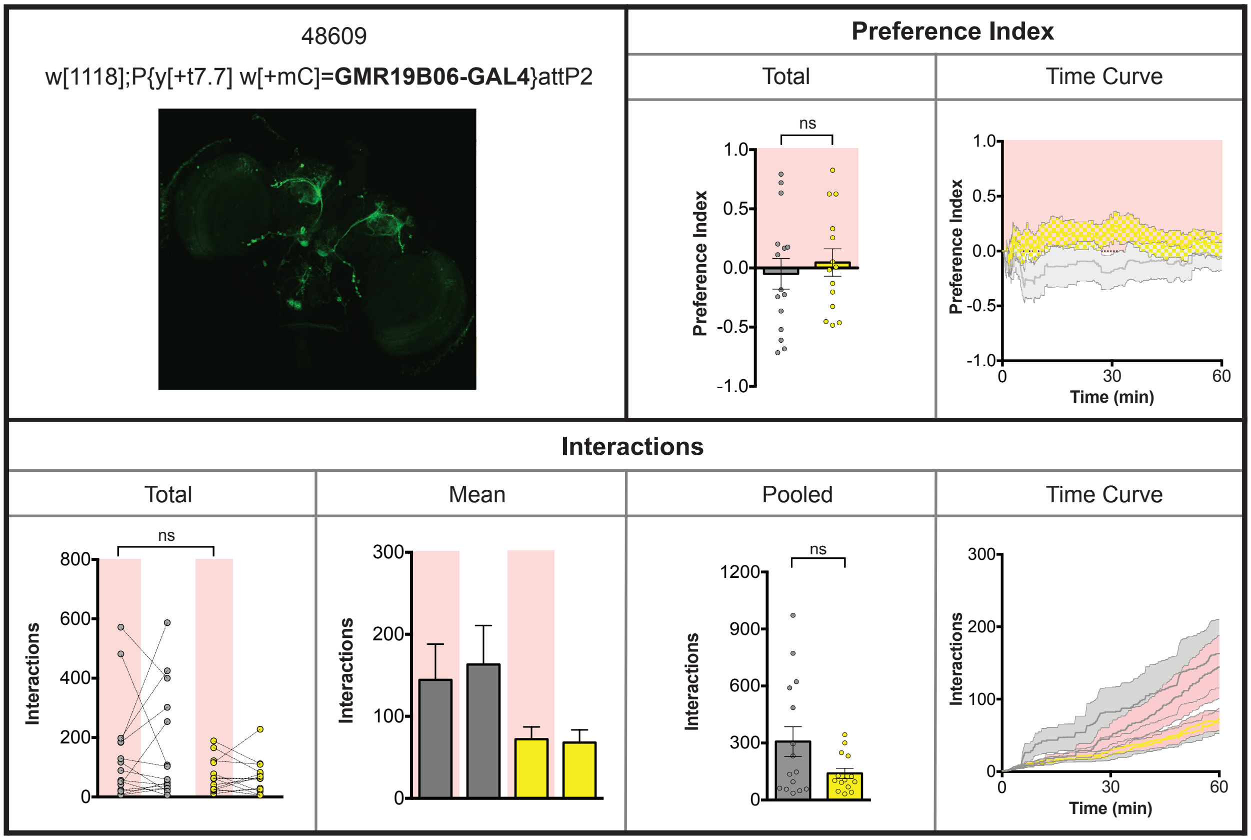
48609
w[1118];P{y[+t7.7] w[+mC]=**GMR19B06-GAL4**}attP2
Preference Index
Total
Time Curve
ns
Preference Index
1.0
0.5
0.0
-0.5
-1.0
Time (min)
0
30
60
Interactions
Total
Mean
Pooled
Time Curve
ns
800
600
400
200
0
300
200
100
1200
900
600
300
Time (min)

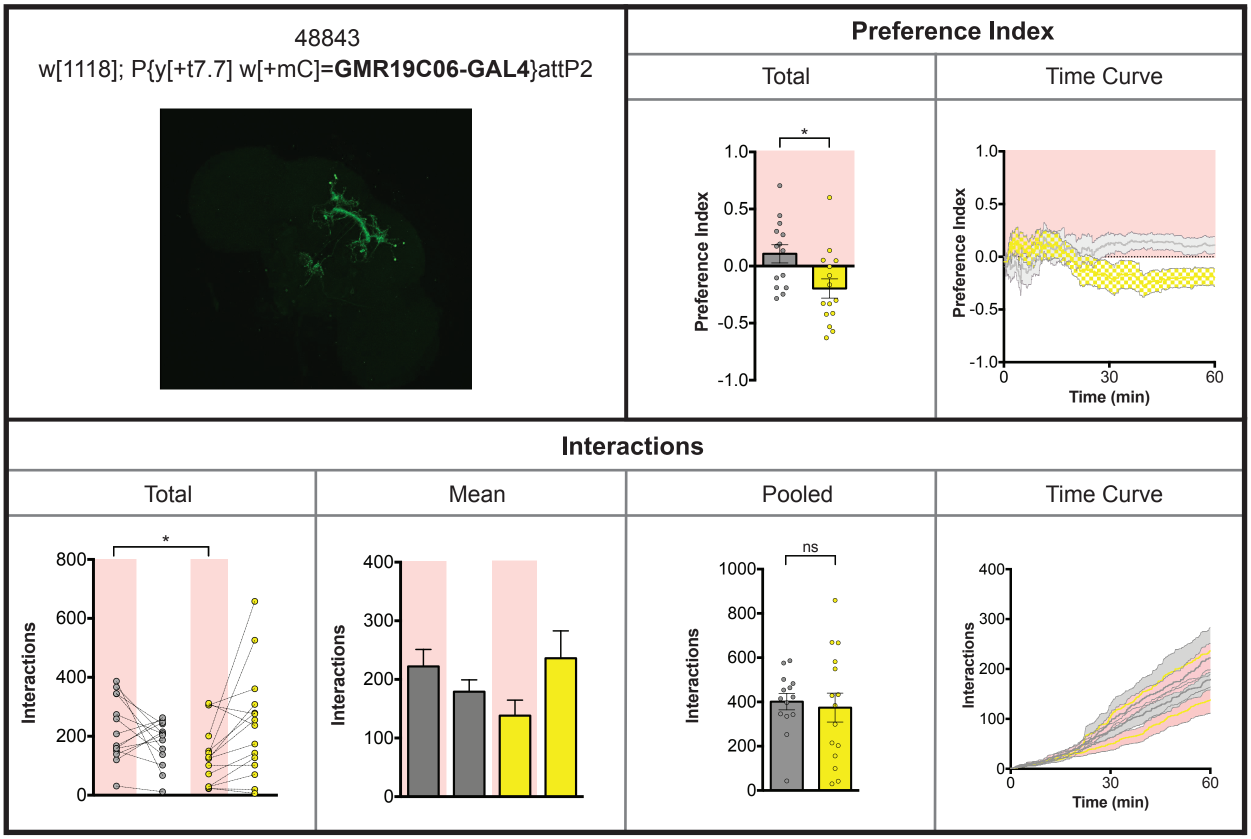
48843
w[1118]; P{y[+t7.7] w[+mC]=**GMR19C06-GAL4**}attP2
Preference Index
Total
Time Curve
*
Preference Index
1.0
0.5
0.0
-0.5
-1.0
Time (min)
0
30
60
Interactions
Total
Mean
Pooled
Time Curve
*
ns
800
600
400
200
0
300
100
1000
Time (min)

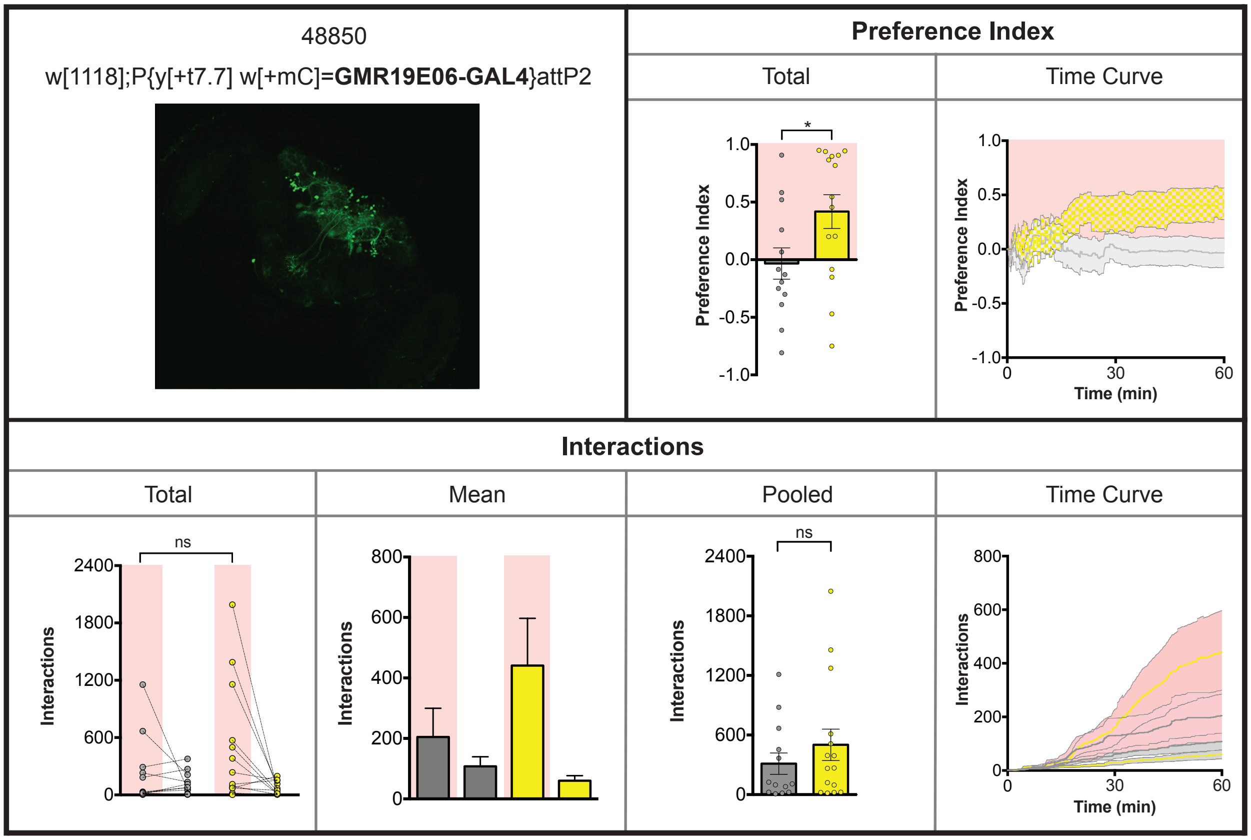
48850
w[1118];P{y[+t7.7] w[+mC]=GMR19E06-GAL4}attP2
Preference Index
Total
Time Curve
Preference Index
Time (min)
Interactions
Total
Mean
Pooled
Time Curve
ns
*
Time (min)

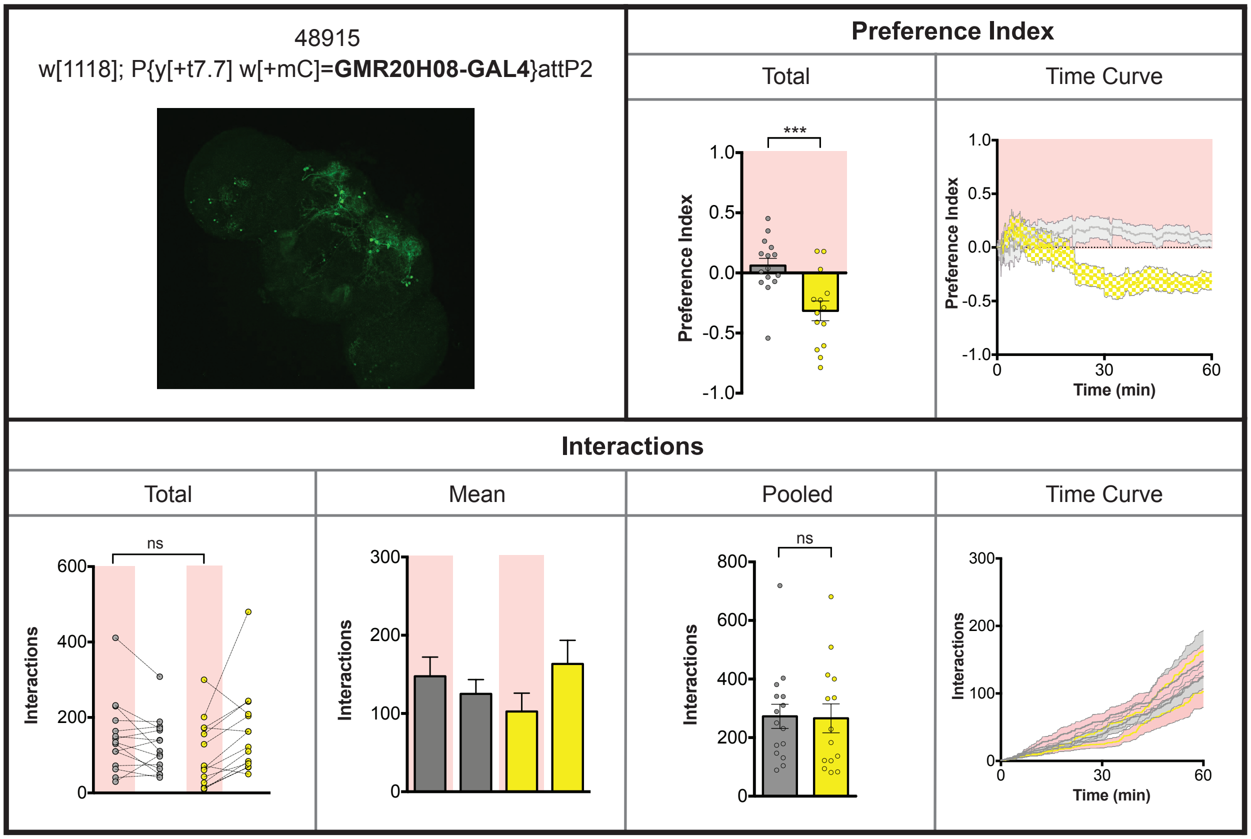
48915
w[1118]; P{y[+t7.7] w[+mC]=GMR20H08-GAL4}attP2
Preference Index
Total
Time Curve
Preference Index
Time (min)
Interactions
Total
Mean
Pooled
Time Curve
ns
***
Time (min)

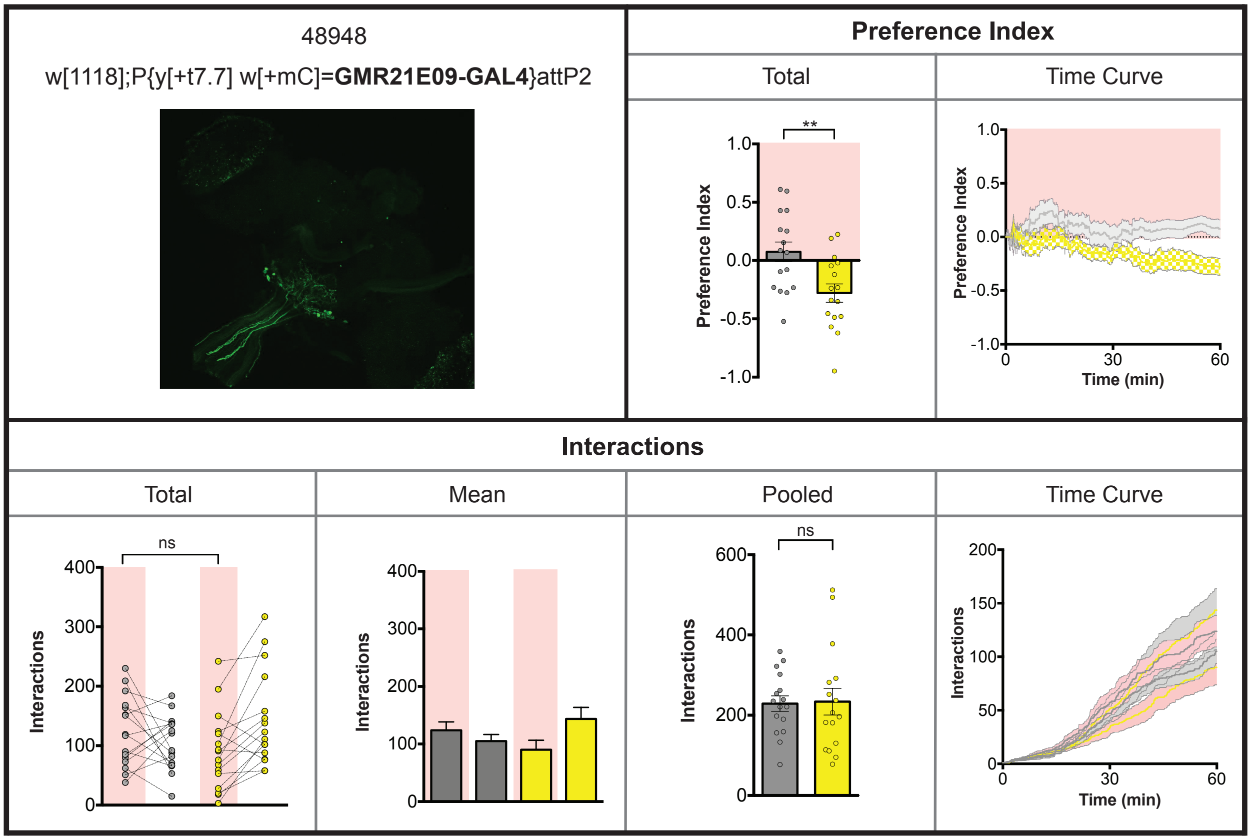
48948
w[1118];P{y[+t7.7] w[+mC]=**GMR21E09-GAL4**}attP2
Preference Index
Total
Time Curve
Preference Index
Time (min)
**
Interactions
Total
Mean
Pooled
Time Curve
ns
ns
Interactions
Time (min)

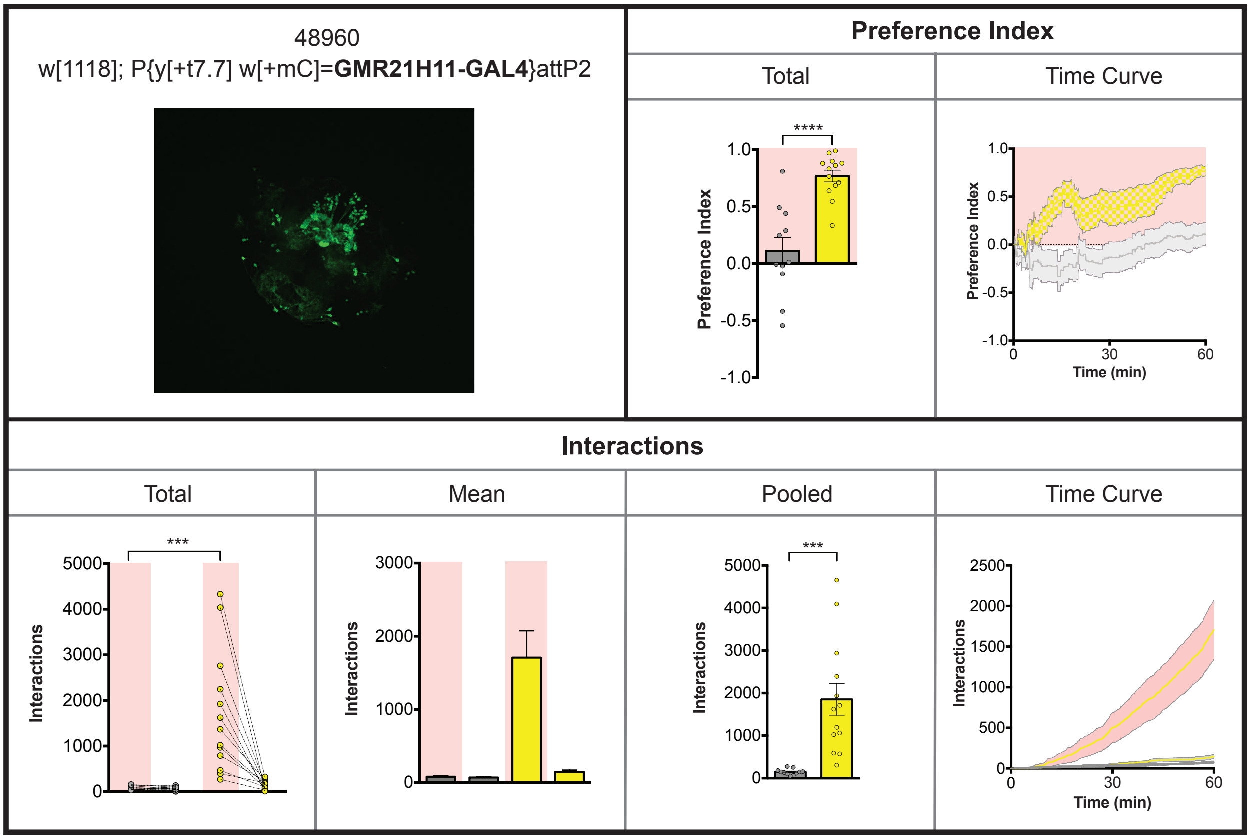
48960
w[1118]; P{y[+t7.7] w[+mC]=**GMR21H11-GAL4**}attP2
Preference Index
Total
Time Curve
Preference Index
Time (min)
****
Interactions
Total
Mean
Pooled
Time Curve
***
***
Interactions
Time (min)

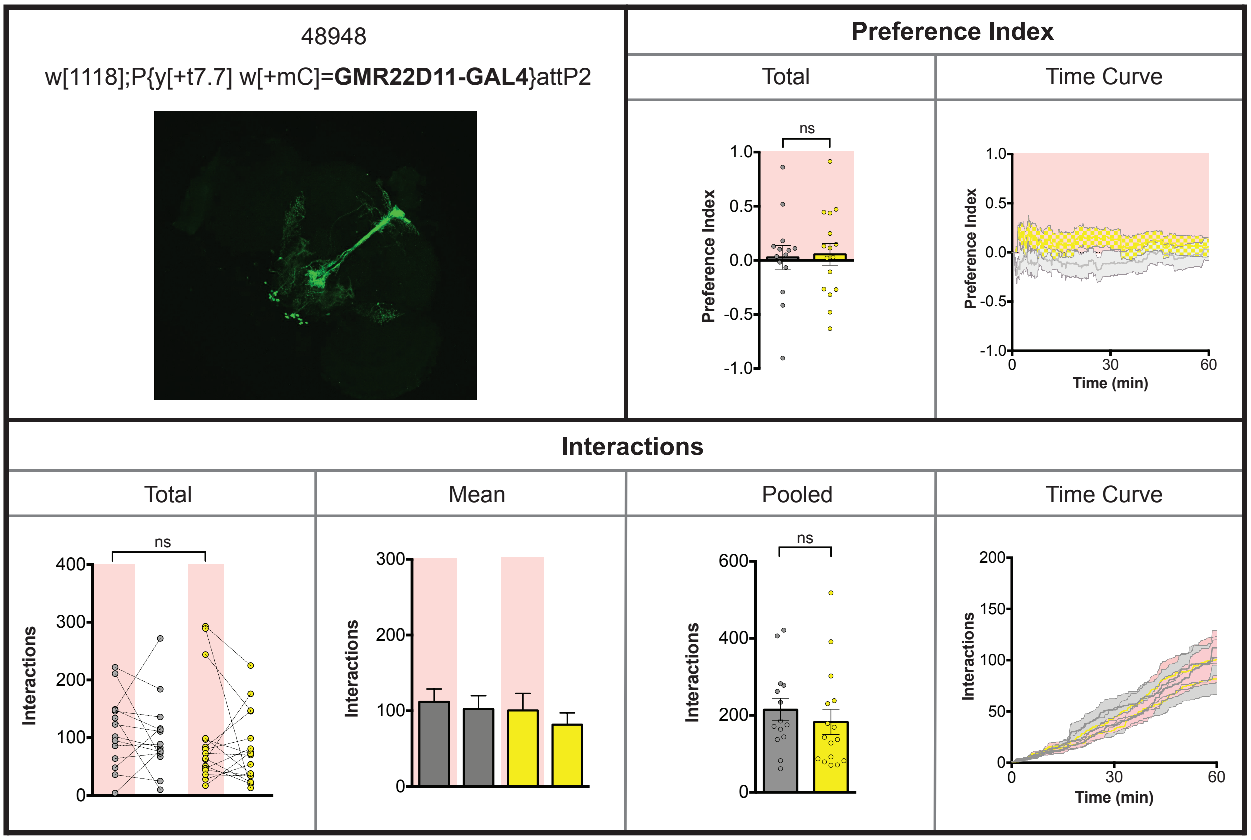
48948
w[1118];P{y[+t7.7] w[+mC]=**GMR22D11-GAL4**}attP2
Preference Index
Total
Time Curve
ns
Preference Index
Time (min)
Interactions
Total
Mean
Pooled
Time Curve
ns
Interactions
Time (min)

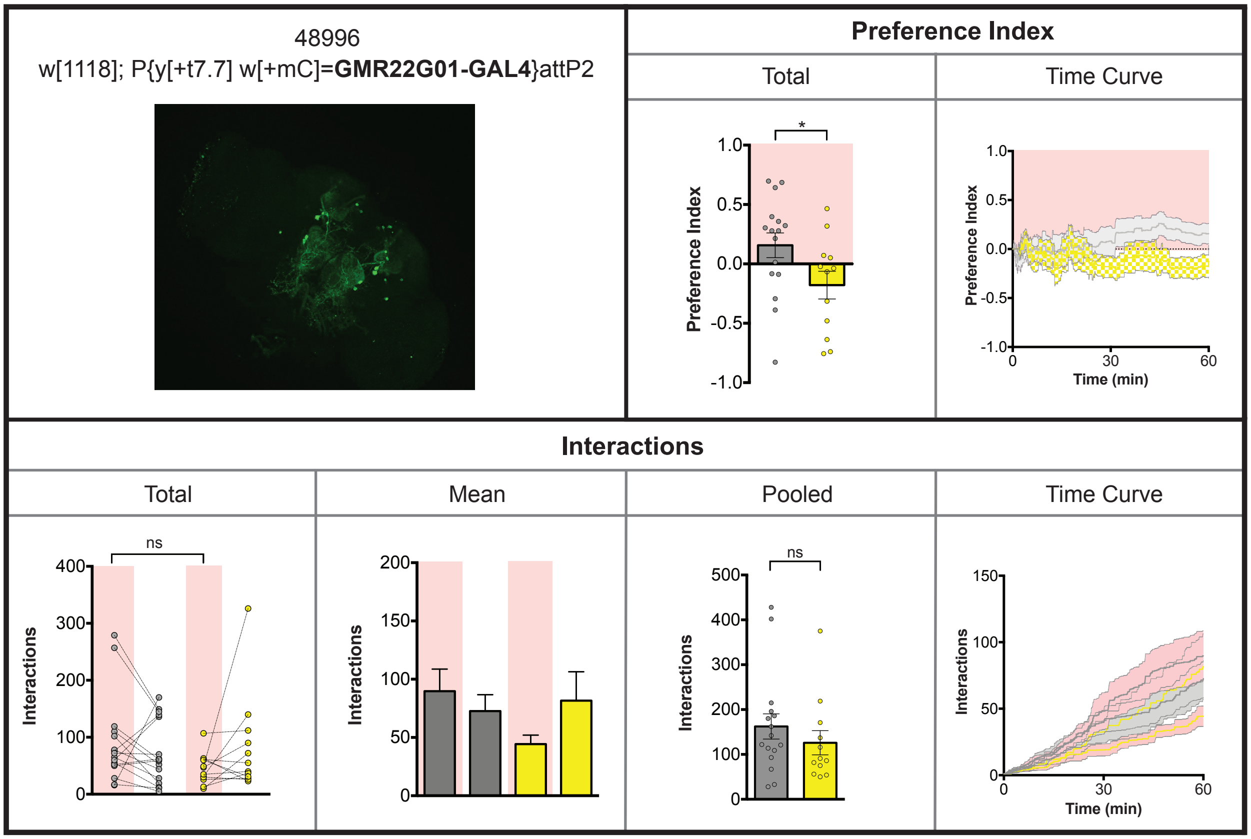
48996
w[1118]; P{y[+t7.7] w[+mC]=**GMR22G01-GAL4**}attP2
Preference Index
Total
Time Curve
*
Preference Index
Time (min)
Interactions
Total
Mean
Pooled
Time Curve
ns
Interactions
Time (min)

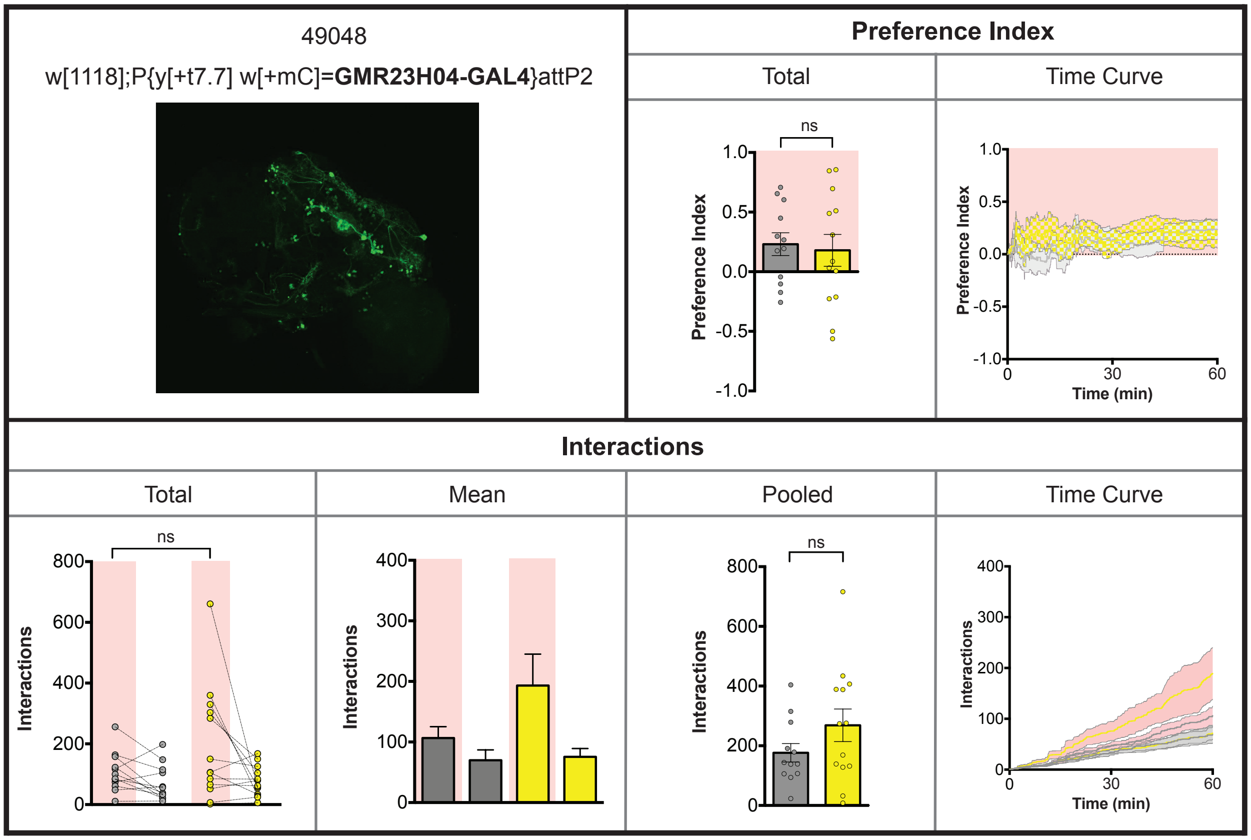

49048
w[1118];P{y[+t7.7] w[+mC]=**GMR23H04-GAL4**}attP2
Preference Index
Total
Time Curve
ns
1.0
0.5
0.0
-0.5
-1.0
Preference Index
0
30
60
Time (min)
Interactions
Total
Mean
Pooled
Time Curve
ns
800
600
400
200
0
Interactions
400
300
200
100
0
Interactions
ns
800
600
400
200
0
Interactions
400
300
200
100
0
Interactions
0
30
60
Time (min)


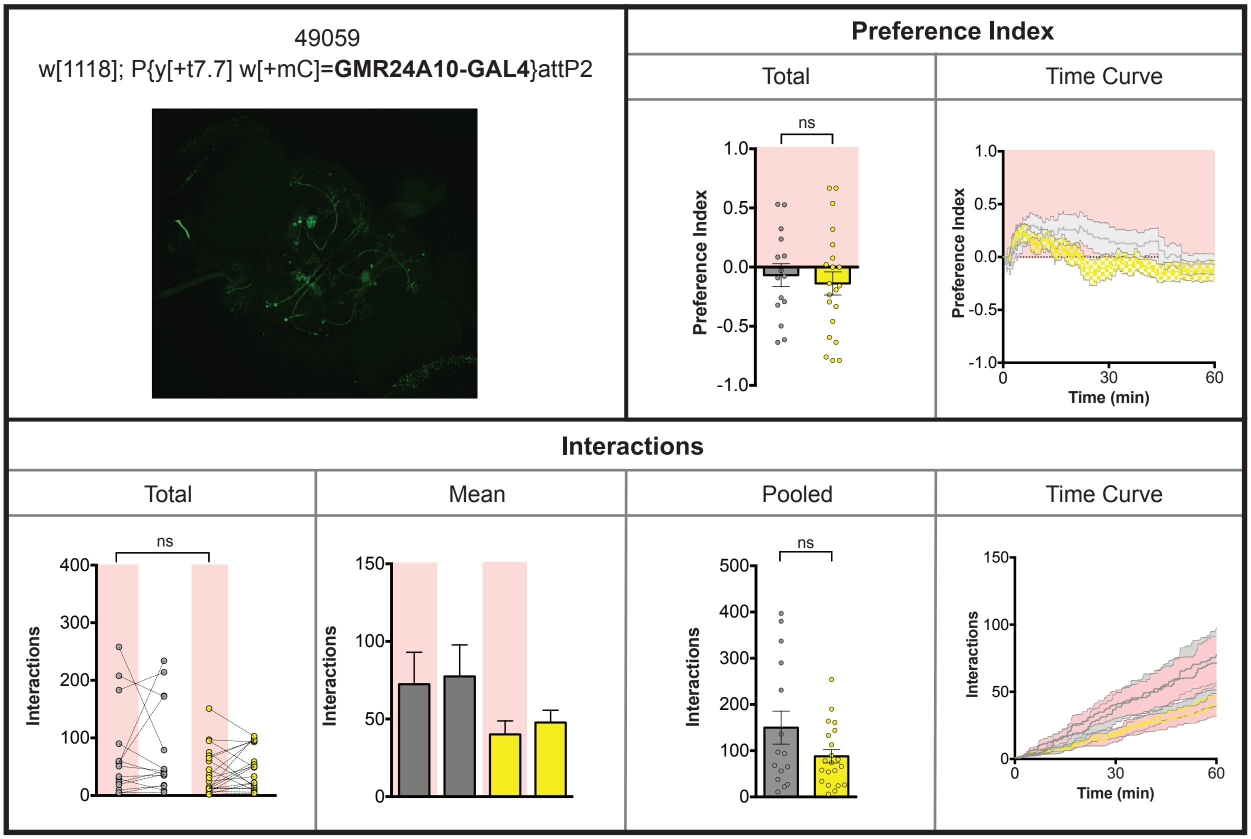

49059
w[1118]; P{y[+t7.7] w[+mC]=**GMR24A10-GAL4**}attP2
Preference Index
Total
Time Curve
ns
1.0
0.5
0.0
-0.5
-1.0
Preference Index
0
30
60
Time (min)
Interactions
Total
Mean
Pooled
Time Curve
ns
400
300
200
100
0
Interactions
150
100
50
0
Interactions
ns
500
400
300
200
100
0
Interactions
150
100
50
0
Interactions
0
30
60
Time (min)

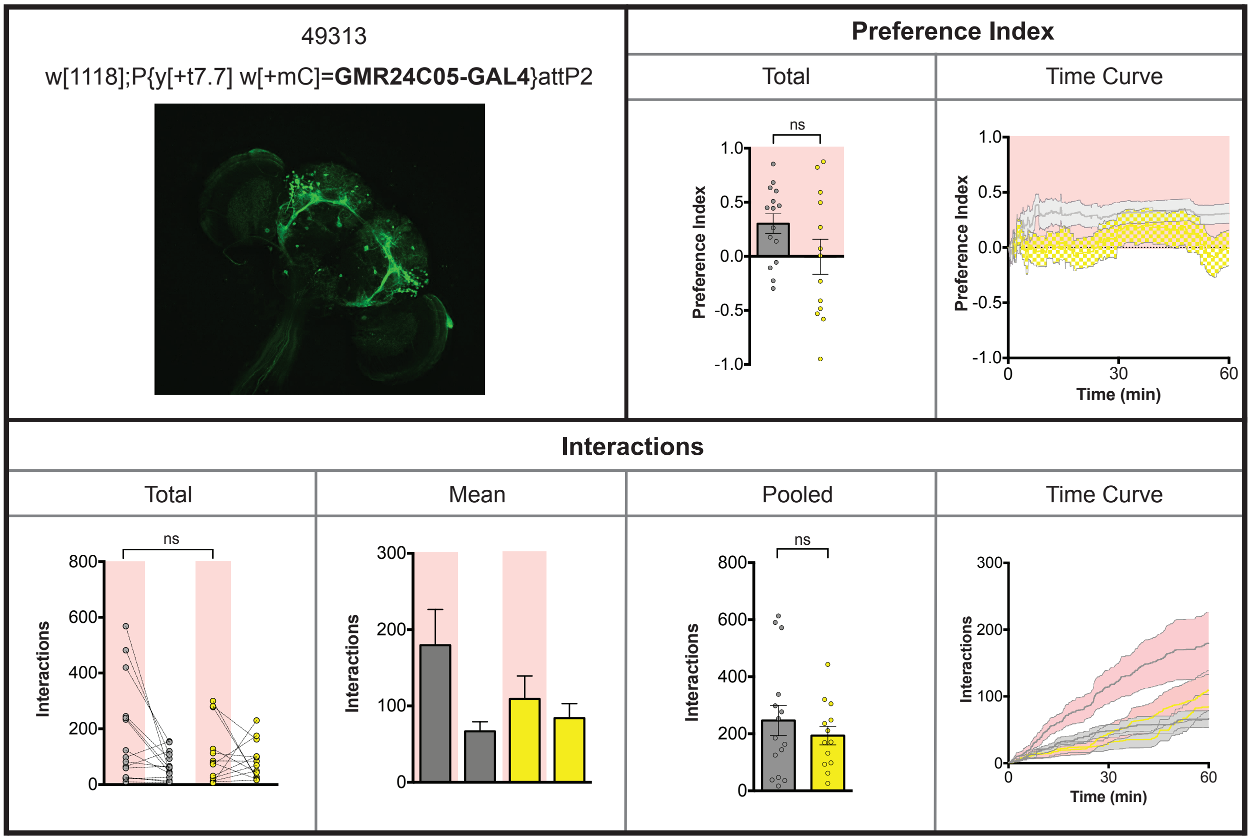
49313
w[1118];P{y[+t7.7] w[+mC]=**GMR24C05-GAL4**}attP2
Preference Index
Total
Time Curve
ns
Preference Index
Time (min)
Interactions
Total
Mean
Pooled
Time Curve
ns
Interactions
Time (min)

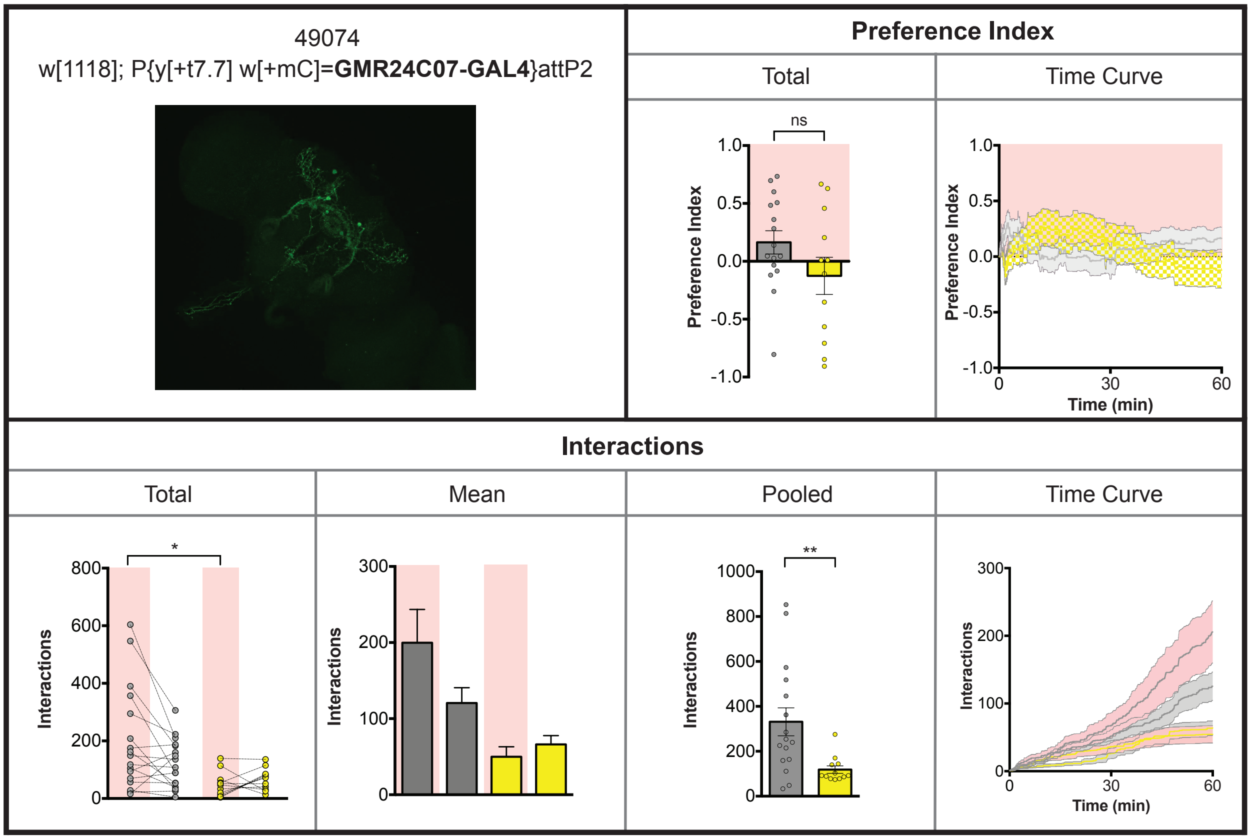
49074
w[1118]; P{y[+t7.7] w[+mC]=**GMR24C07-GAL4**}attP2
Preference Index
Total
Time Curve
ns
Preference Index
Time (min)
Interactions
Total
Mean
Pooled
Time Curve
*
**
Interactions
Time (min)

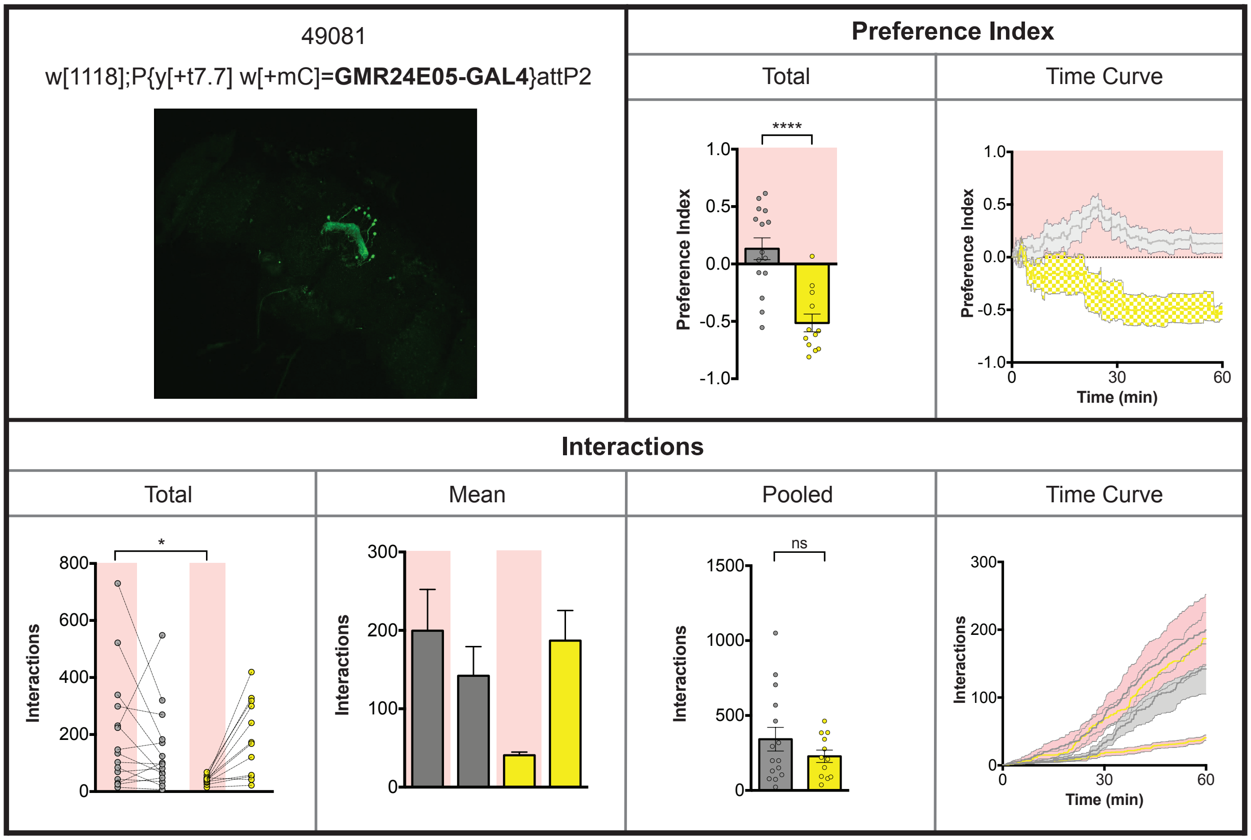
49081
w[1118];P{y[+t7.7] w[+mC]=**GMR24E05-GAL4**}attP2
Preference Index
Total
Time Curve
****
1.0
0.5
0.0
-0.5
-1.0
Preference Index
0
30
60
Time (min)
Interactions
Total
Mean
Pooled
Time Curve
*
800
600
400
200
0
Interactions
300
200
100
0
ns
1500
1000
500
0
Time (min)

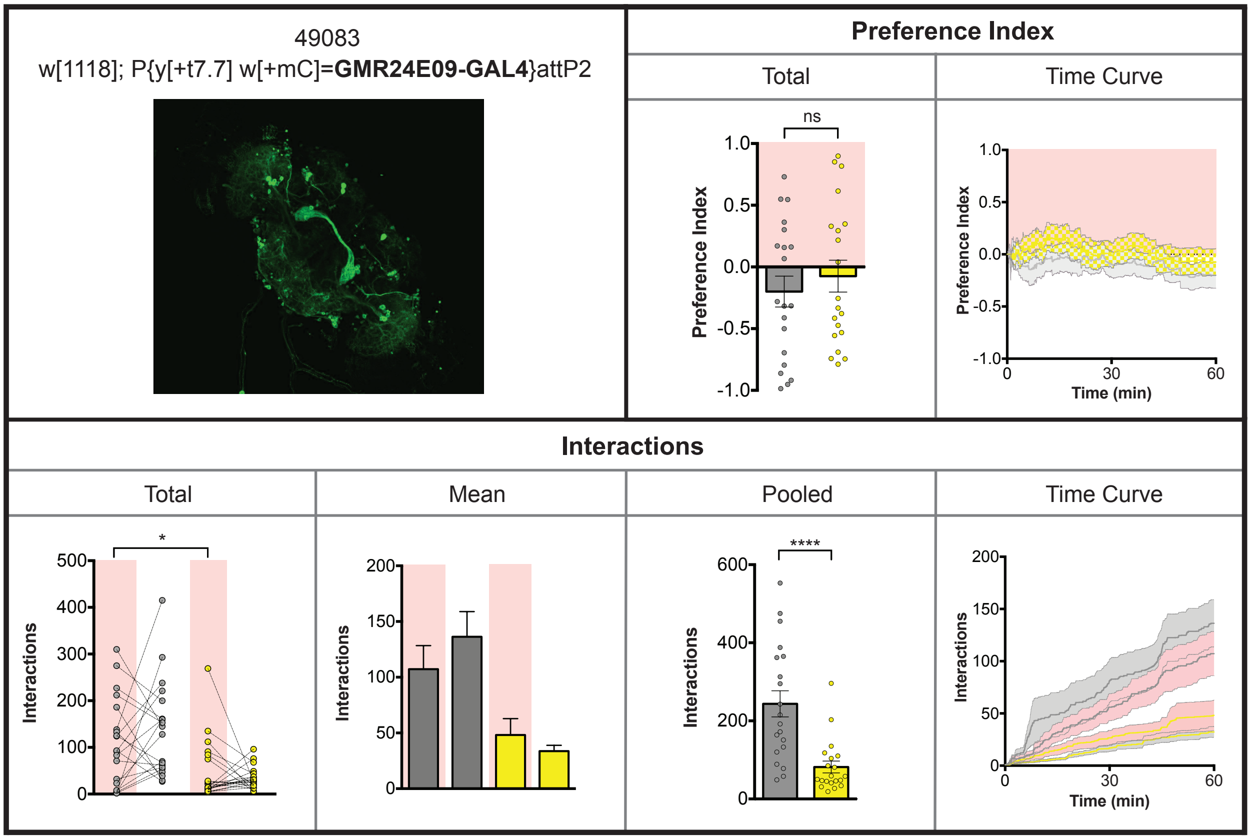
49083
w[1118]; P{y[+t7.7] w[+mC]=**GMR24E09-GAL4**}attP2
Preference Index
Total
Time Curve
ns
1.0
0.5
0.0
-0.5
-1.0
Preference Index
0
30
60
Time (min)
Interactions
Total
Mean
Pooled
Time Curve
*
500
400
300
200
100
0
Interactions
200
150
100
50
0
****
600
400
200
Time (min)

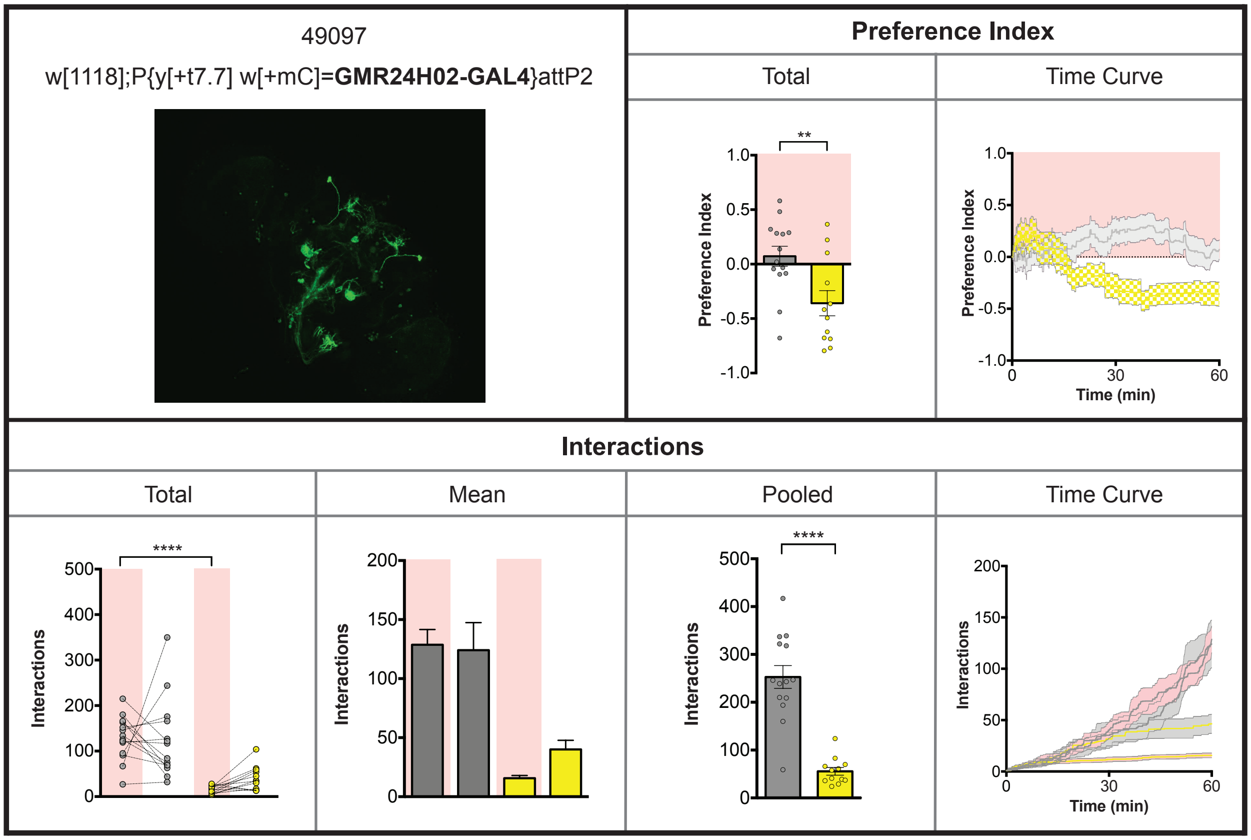
49097
w[1118];P{y[+t7.7] w[+mC]=GMR24H02-GAL4}attP2
Preference Index
Total
Time Curve
Preference Index
Time (min)
Interactions
Total
Mean
Pooled
Time Curve
Interactions
Time (min)

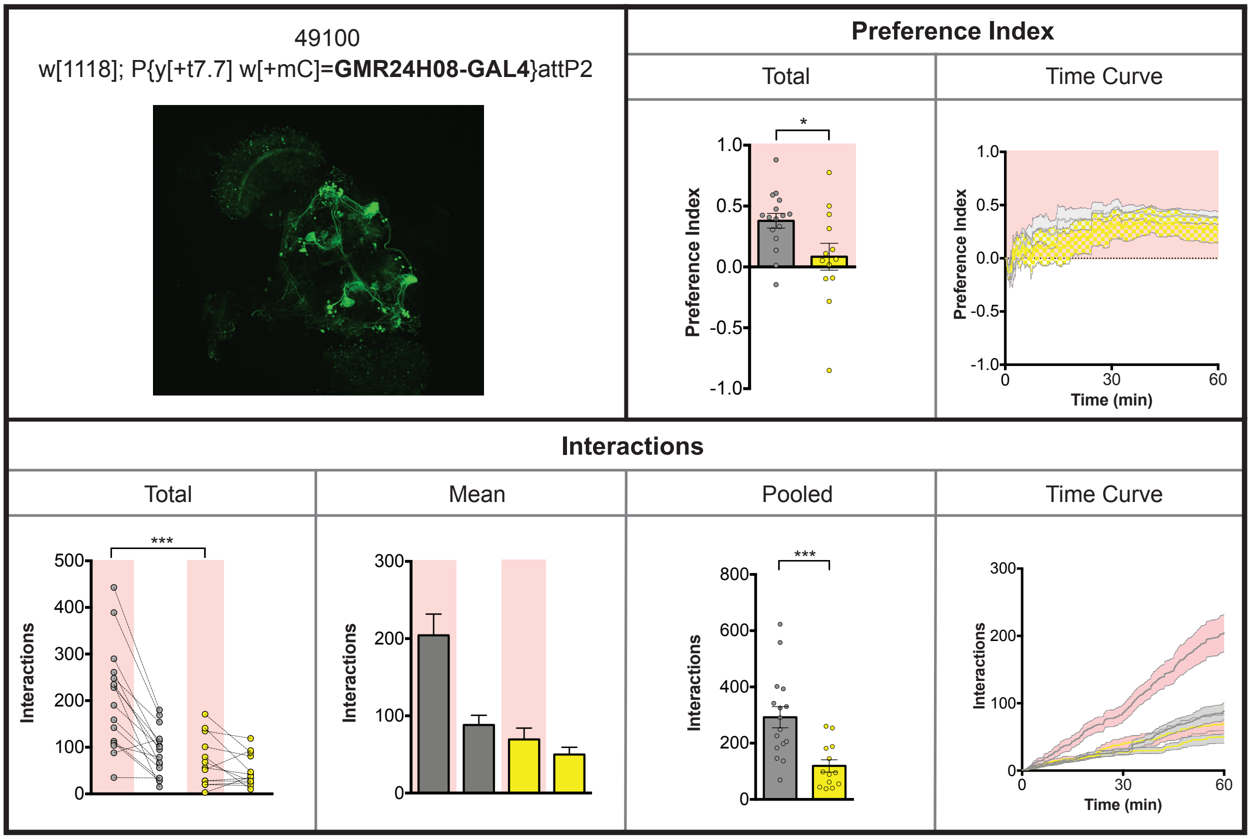
49100
w[1118]; P{y[+t7.7] w[+mC]=GMR24H08-GAL4}attP2
Preference Index
Total
Time Curve
Preference Index
Time (min)
Interactions
Total
Mean
Pooled
Time Curve
Interactions
Time (min)

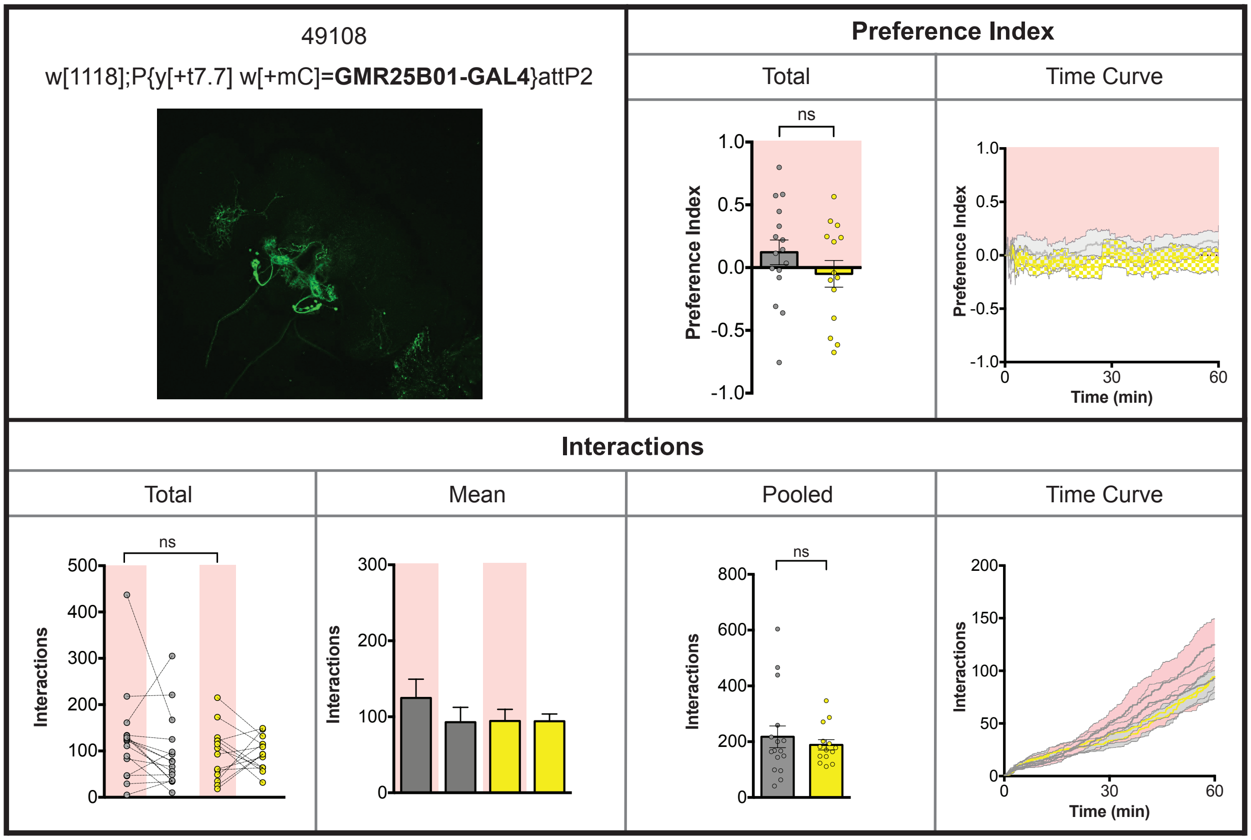
49108
w[1118];P{y[+t7.7] w[+mC]=GMR25B01-GAL4}attP2
Preference Index
Total
Time Curve
ns
Preference Index
1.0
0.5
0.0
-0.5
-1.0
Time (min)
0
30
60
Interactions
Total
Mean
Pooled
Time Curve
ns
500
400
300
200
100
0
300
200
100
800
600
400
200
150
50
Time (min)

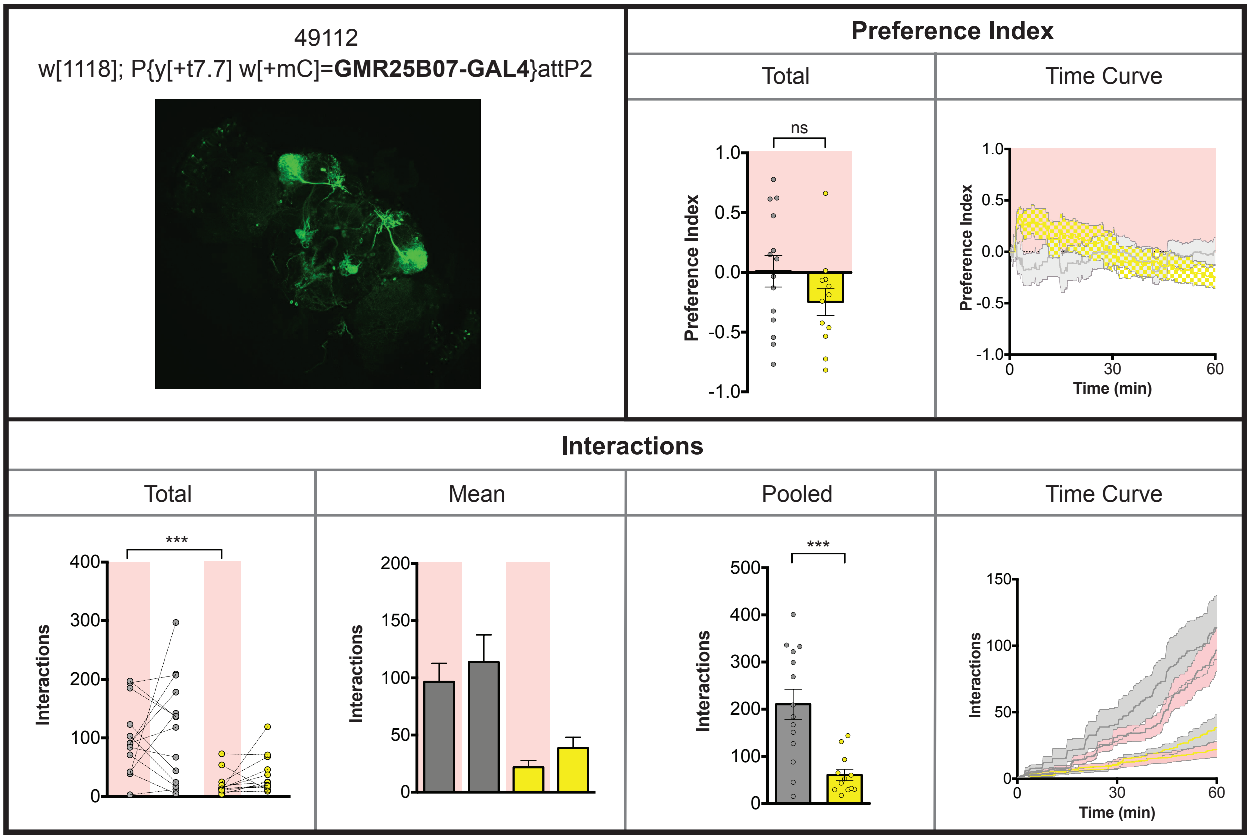
49112
w[1118]; P{y[+t7.7] w[+mC]=GMR25B07-GAL4}attP2
Preference Index
Total
Time Curve
ns
Preference Index
1.0
0.5
0.0
-0.5
-1.0
Time (min)
0
30
60
Interactions
Total
Mean
Pooled
Time Curve
***
400
300
200
100
0
150
50
500
Time (min)

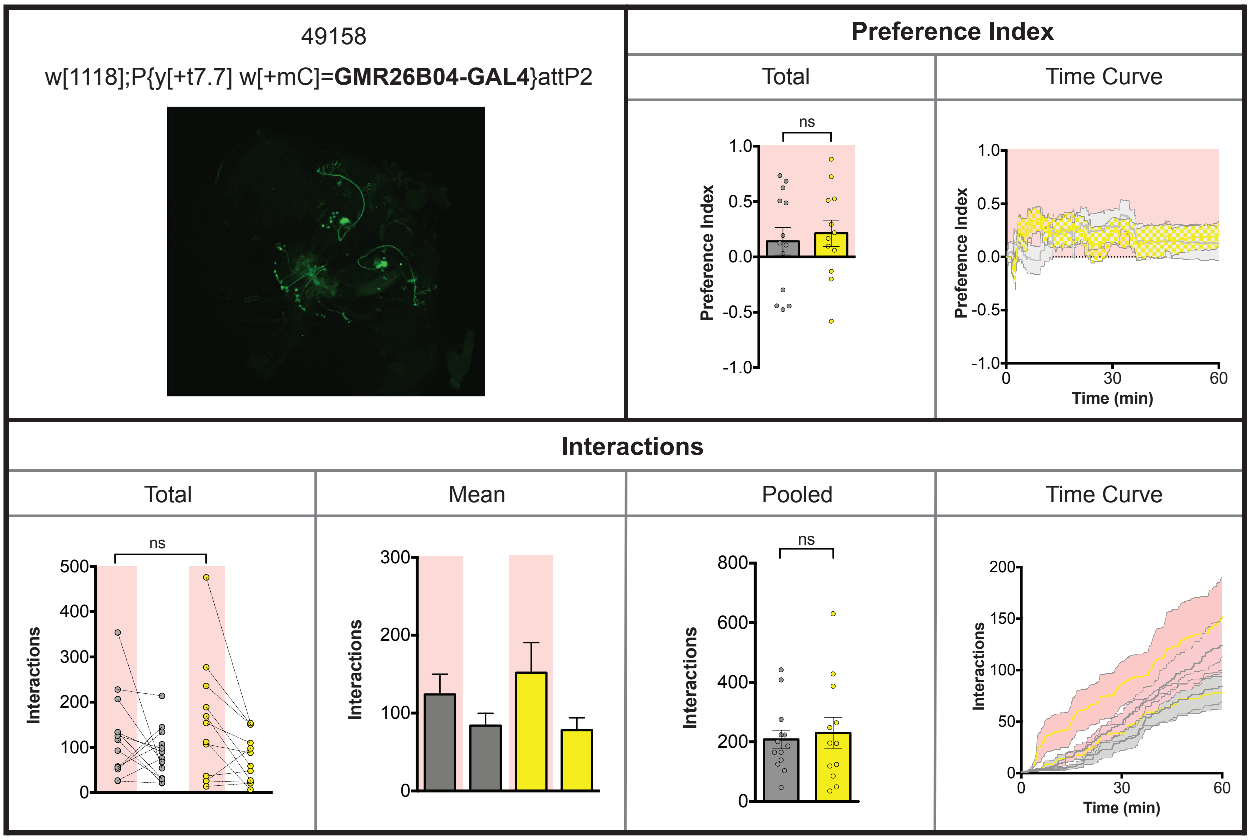
49158
w[1118];P{y[+t7.7] w[+mC]=GMR26B04-GAL4}attP2
Preference Index
Total
Time Curve
ns
Preference Index
1.0
0.5
0.0
-0.5
-1.0
0
30
60
Time (min)
Interactions
Total
Mean
Pooled
Time Curve
ns
500
400
300
200
100
0
300
200
100
0
800
600
400
200
0
150
50


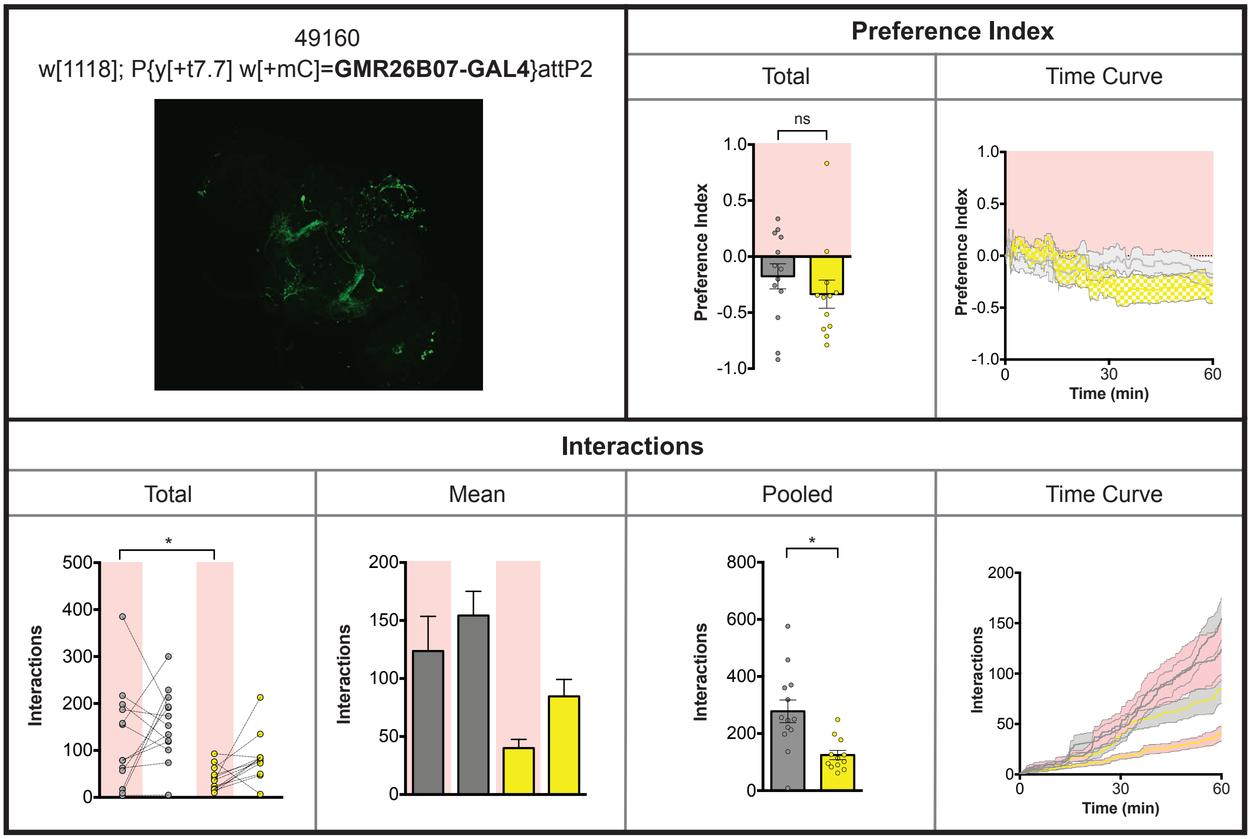
49160
w[1118]; P{y[+t7.7] w[+mC]=GMR26B07-GAL4}attP2
Preference Index
Total
Time Curve
ns
Preference Index
1.0
0.5
0.0
-0.5
-1.0
0
30
60
Time (min)
Interactions
Total
Mean
Pooled
Time Curve
*
500
400
300
200
100
0
150
50
800
600
400
200

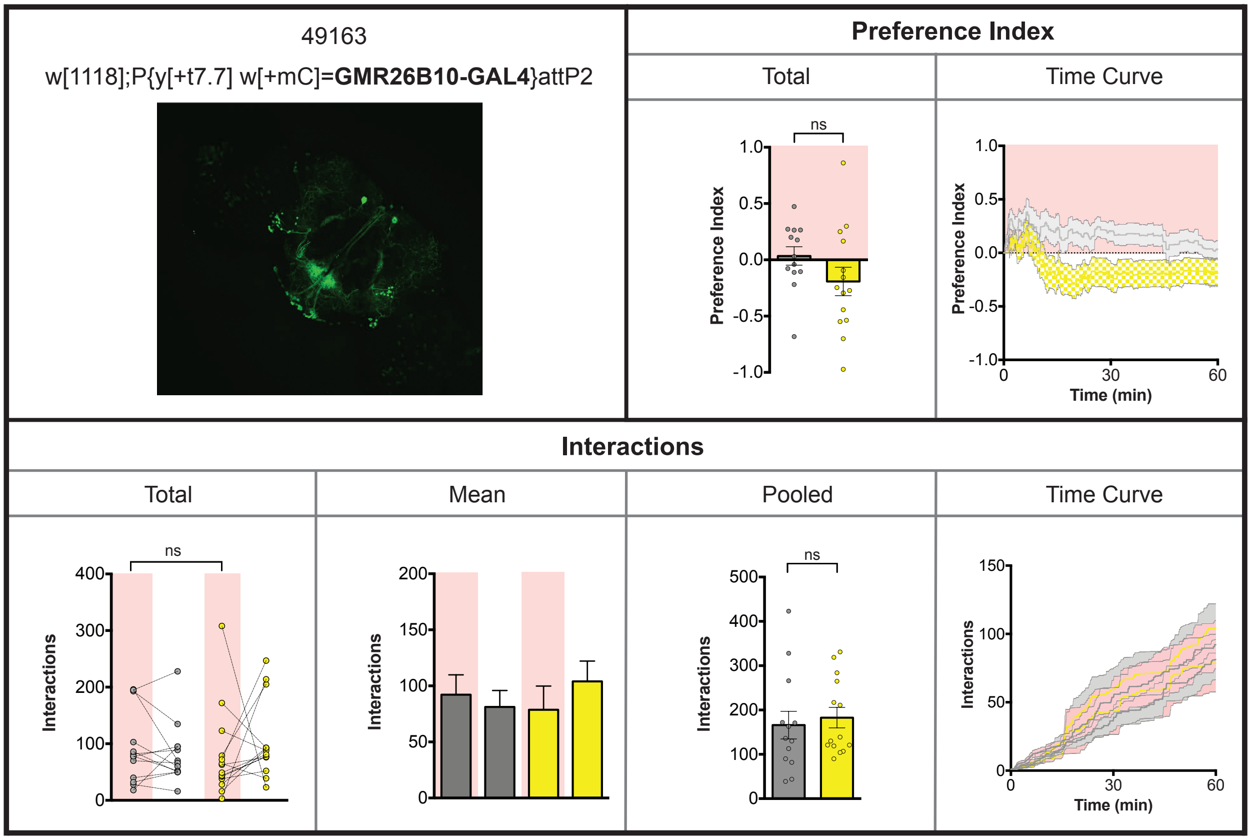
49163
w[1118];P{y[+t7.7] w[+mC]=**GMR26B10-GAL4**}attP2
Preference Index
Total
Time Curve
ns
Preference Index
1.0
0.5
0.0
-0.5
-1.0
0
30
60
Time (min)
Interactions
Total
Mean
Pooled
Time Curve
ns
Interactions
400
300
200
100
0
200
150
100
50
0
500
400
300
200
100
0
150
100
50
0
30
60
Time (min)

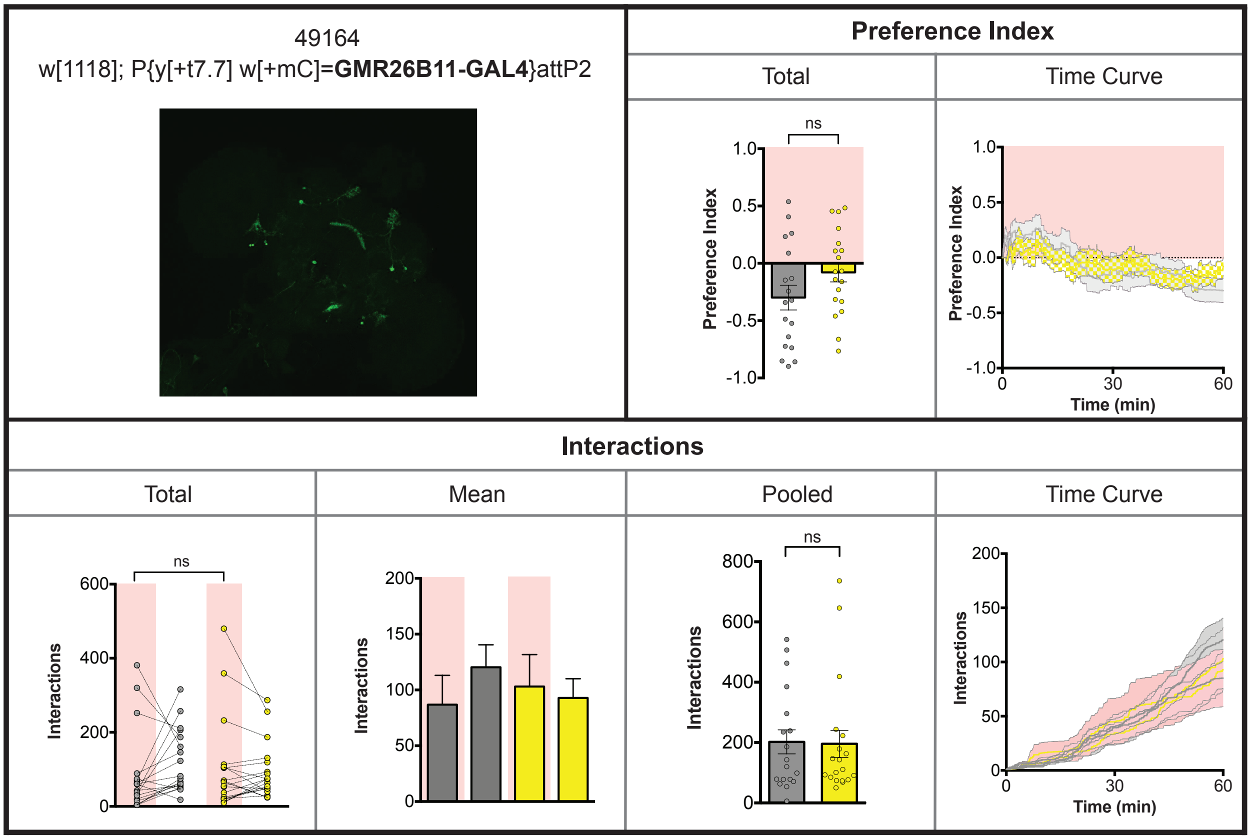
49164
w[1118]; P{y[+t7.7] w[+mC]=**GMR26B11-GAL4**}attP2
Preference Index
Total
Time Curve
ns
Preference Index
1.0
0.5
0.0
-0.5
-1.0
0
30
60
Time (min)
Interactions
Total
Mean
Pooled
Time Curve
ns
Interactions
600
400
200
0
200
150
100
50
0
800
600
400
200
0
200
150
100
50
0
30
60
Time (min)

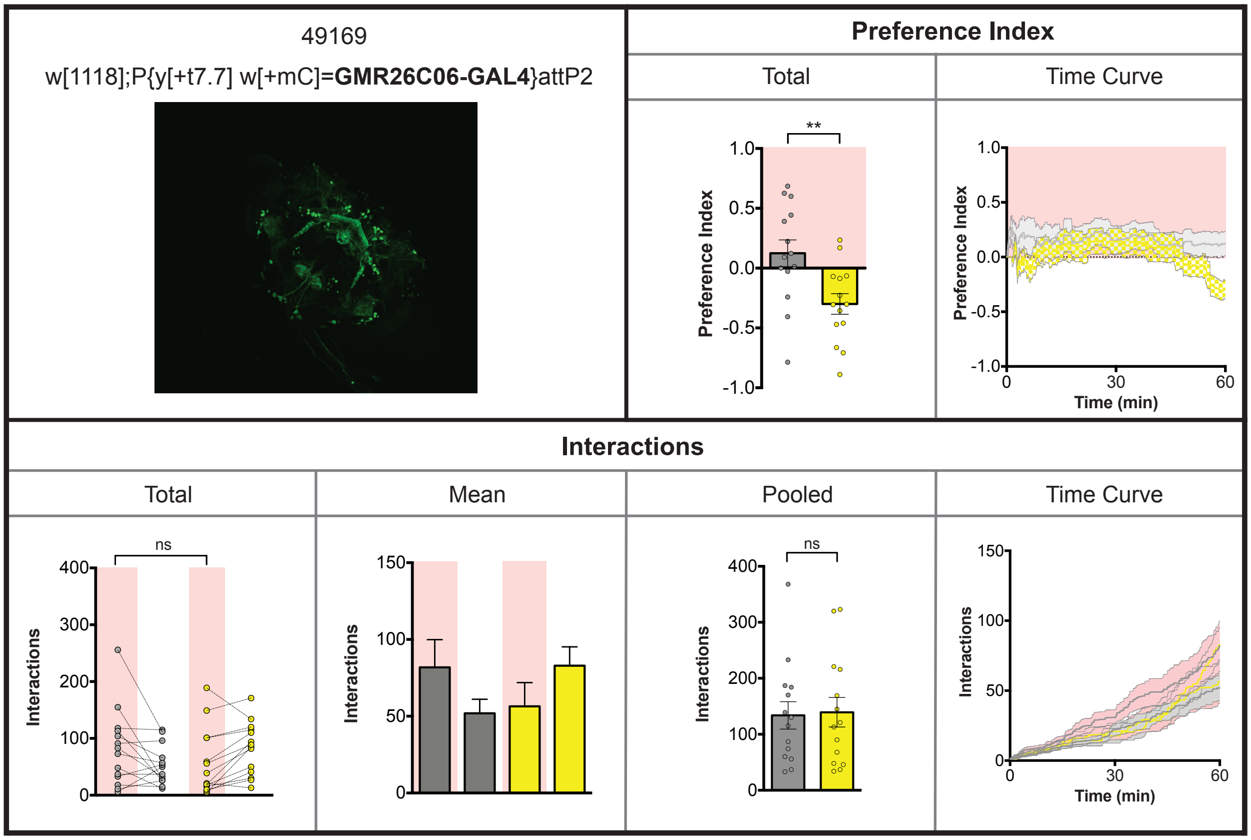
49169
w[1118];P{y[+t7.7] w[+mC]=GMR26C06-GAL4}attP2
Preference Index
Total
Time Curve
**
Preference Index
1.0
0.5
0.0
-0.5
-1.0
Time (min)
0
30
60
Interactions
Total
Mean
Pooled
Time Curve
ns
400
300
200
100
0
150
100
50
Interactions
Time (min)

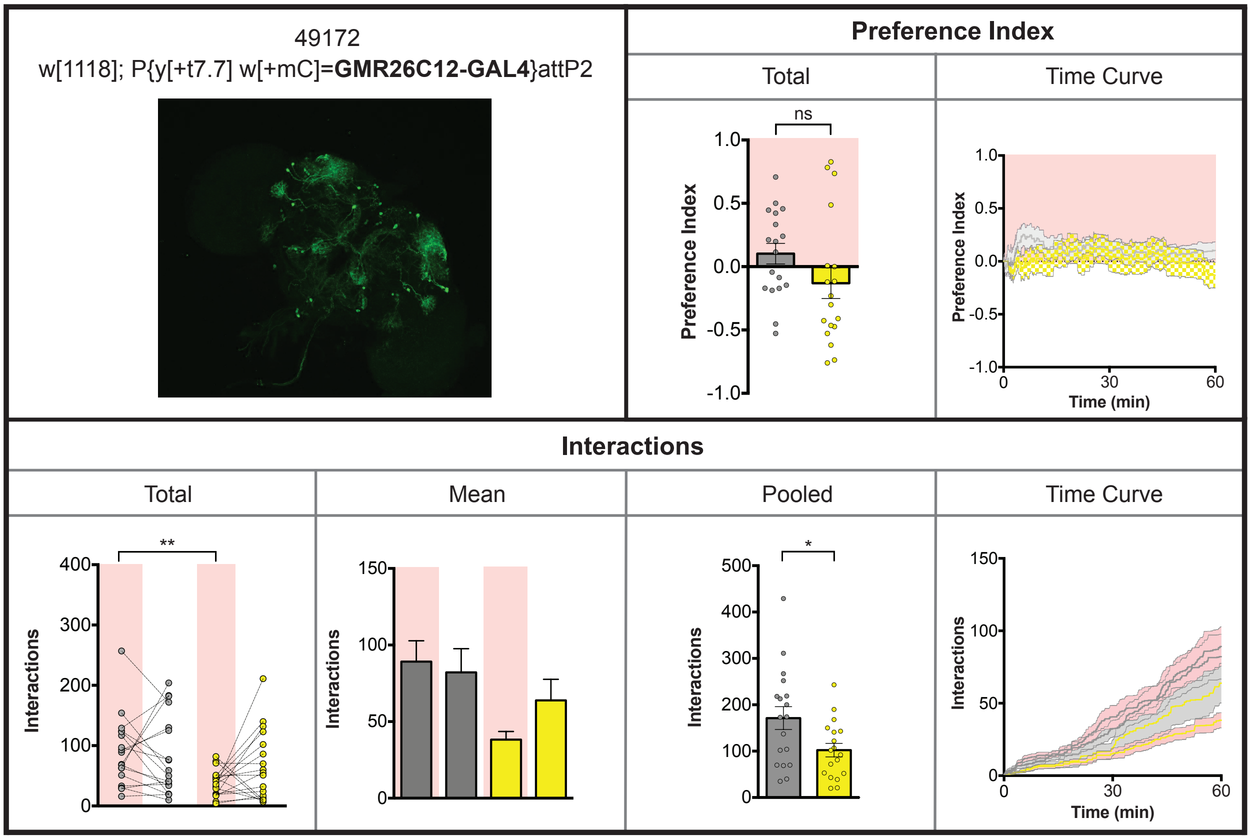
49172
w[1118]; P{y[+t7.7] w[+mC]=GMR26C12-GAL4}attP2
Preference Index
Total
Time Curve
ns
Preference Index
1.0
0.5
0.0
-0.5
-1.0
Time (min)
0
30
60
Interactions
Total
Mean
Pooled
Time Curve
**
*
500
400
300
200
100
0
150
100
50
Interactions
Time (min)

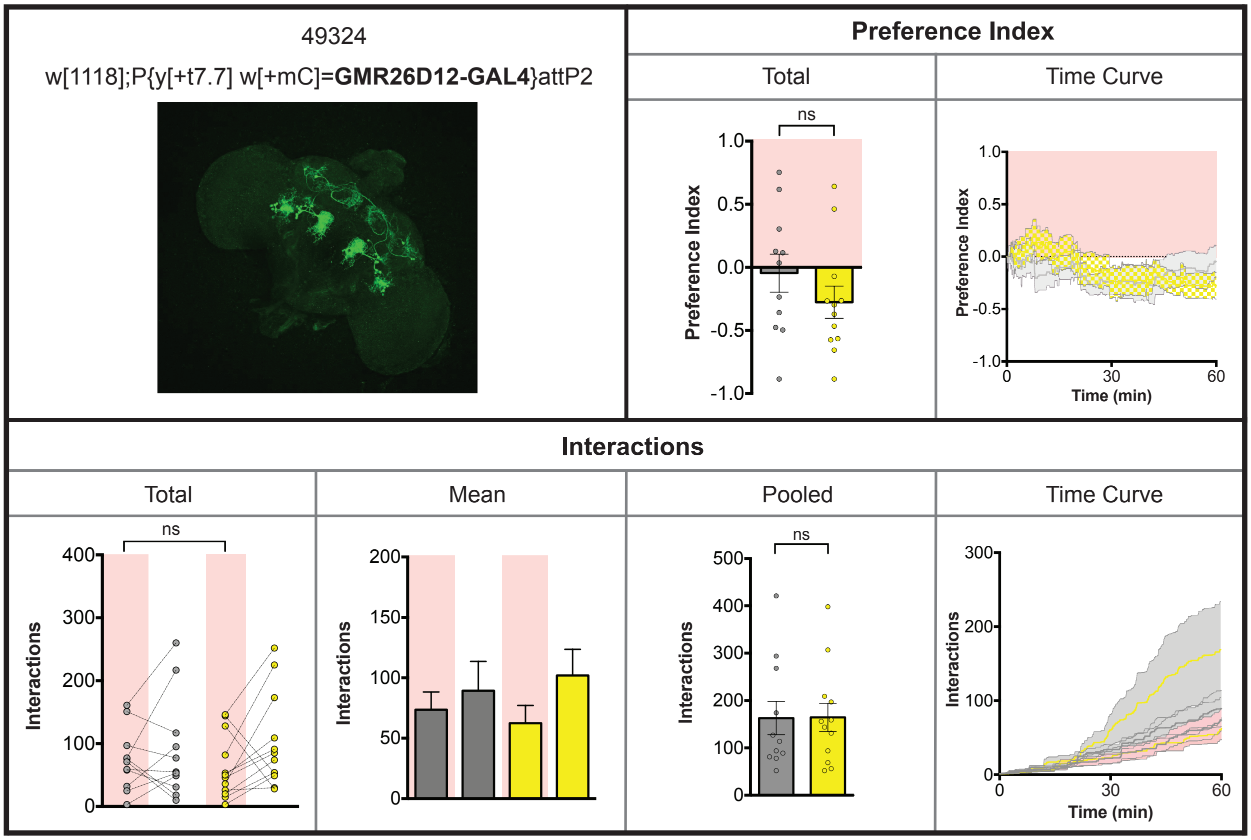
49324
w[1118];P{y[+t7.7] w[+mC]=**GMR26D12-GAL4**}attP2
Preference Index
Total
Time Curve
ns
Preference Index
1.0
0.5
0.0
-0.5
-1.0
Time (min)
0
30
60
Interactions
Total
Mean
Pooled
Time Curve
ns
Interactions
400
300
200
100
0
200
150
100
50
500
400
300
200
100
Time (min)

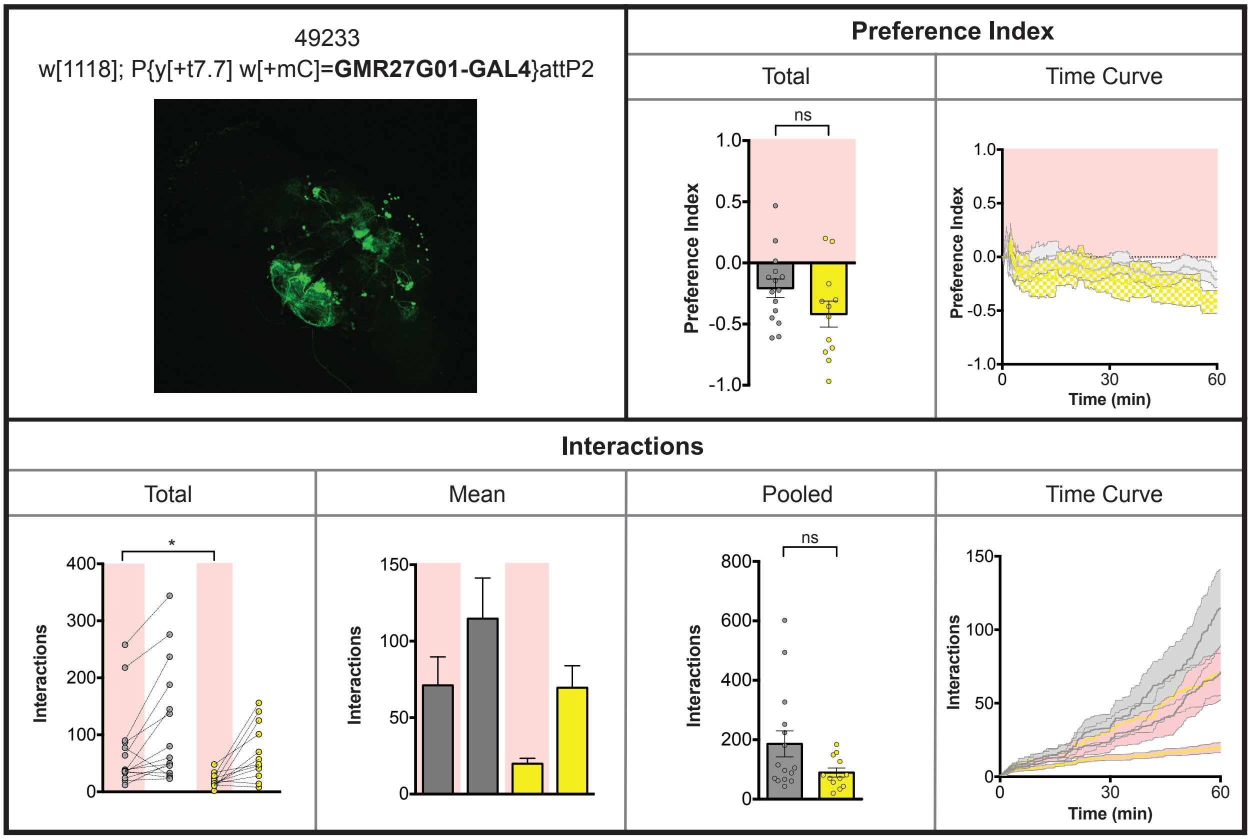
49233
w[1118]; P{y[+t7.7] w[+mC]=**GMR27G01-GAL4**}attP2
Preference Index
Total
Time Curve
ns
Preference Index
1.0
0.5
0.0
-0.5
-1.0
Time (min)
0
30
60
Interactions
Total
Mean
Pooled
Time Curve
*
ns
Interactions
400
300
200
100
0
150
100
50
800
600
400
200
Time (min)

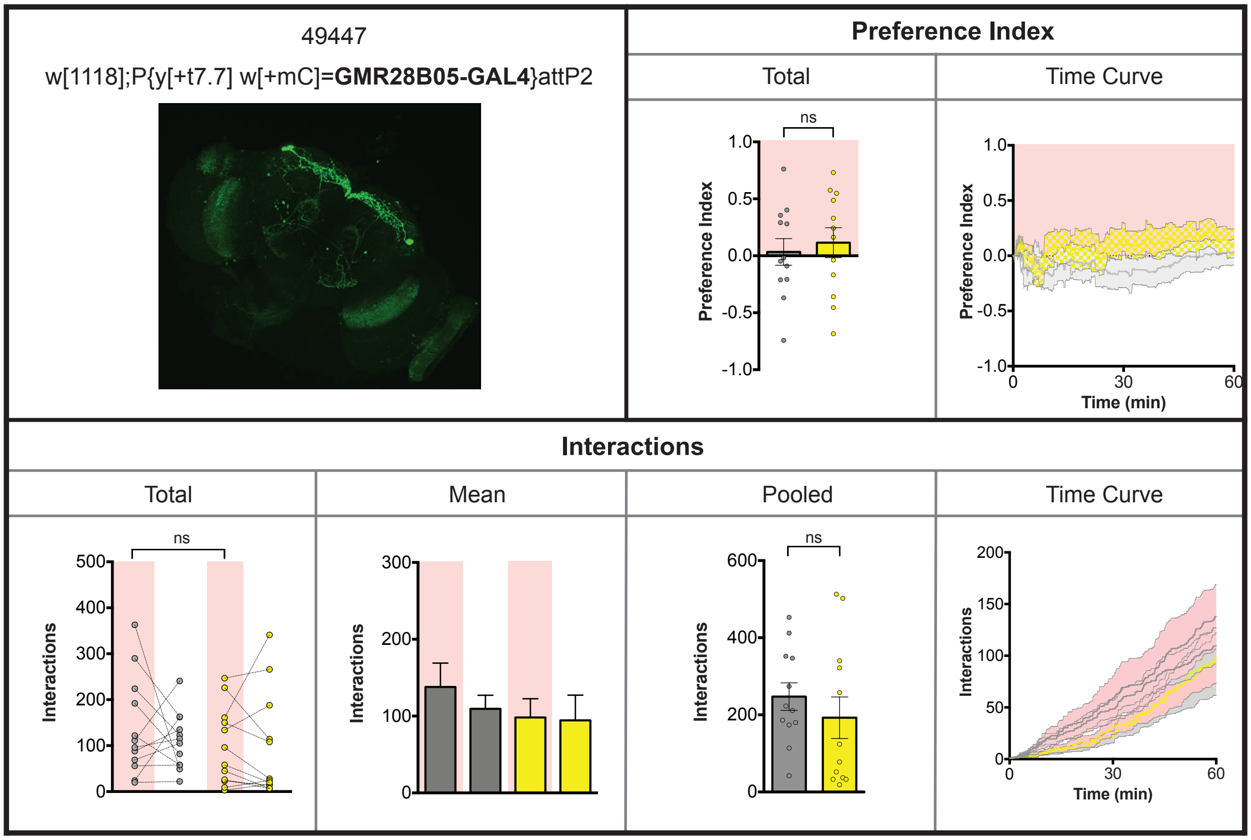
49447
w[1118];P{y[+t7.7] w[+mC]=**GMR28B05-GAL4**}attP2
Preference Index
Total
Time Curve
ns
Preference Index
1.0
0.5
0.0
-0.5
-1.0
0
30
60
Time (min)
Interactions
Total
Mean
Pooled
Time Curve
Interactions
500
400
300
200
100
600
150
50
0

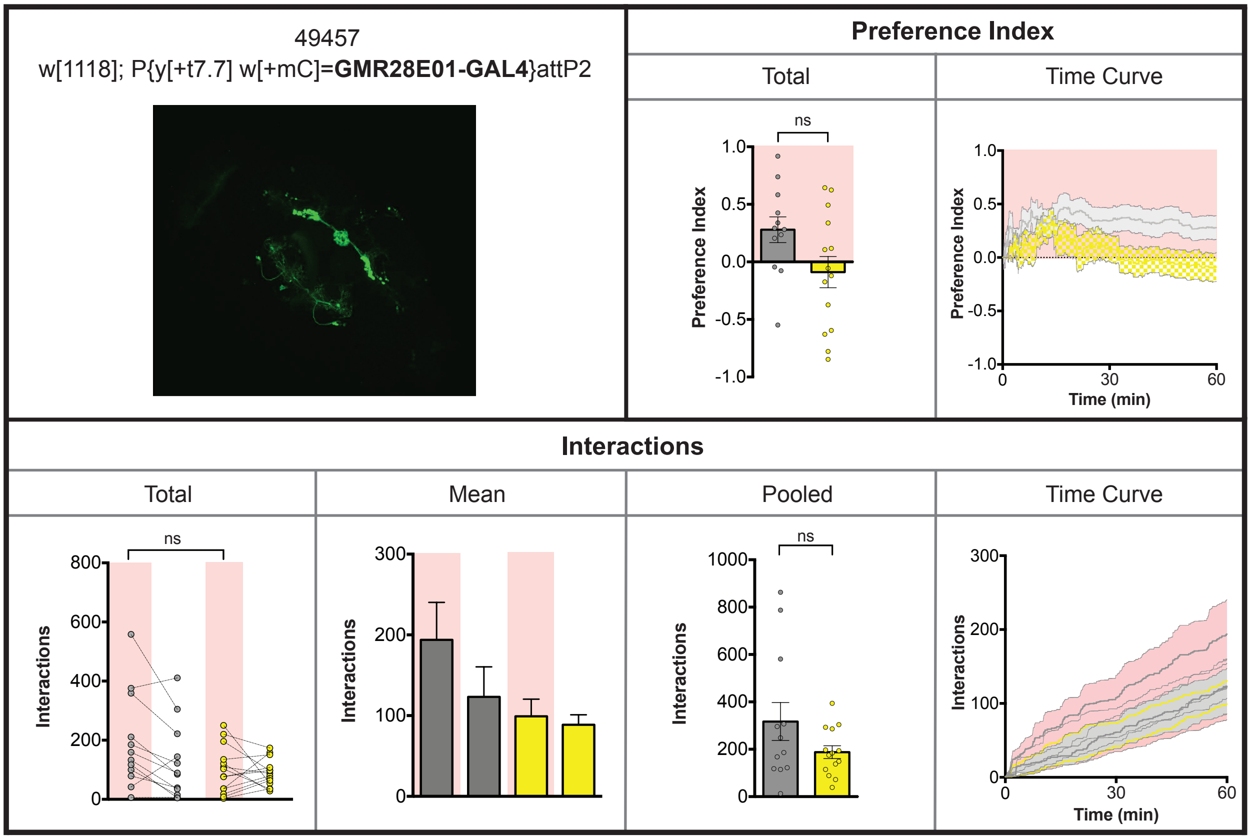
49457
w[1118]; P{y[+t7.7] w[+mC]=**GMR28E01-GAL4**}attP2
Preference Index
Total
Time Curve
ns
Preference Index
1.0
0.5
0.0
-0.5
-1.0
0
30
60
Time (min)
Interactions
Total
Mean
Pooled
Time Curve
Interactions
1000
800
600
400
300
200
100
0

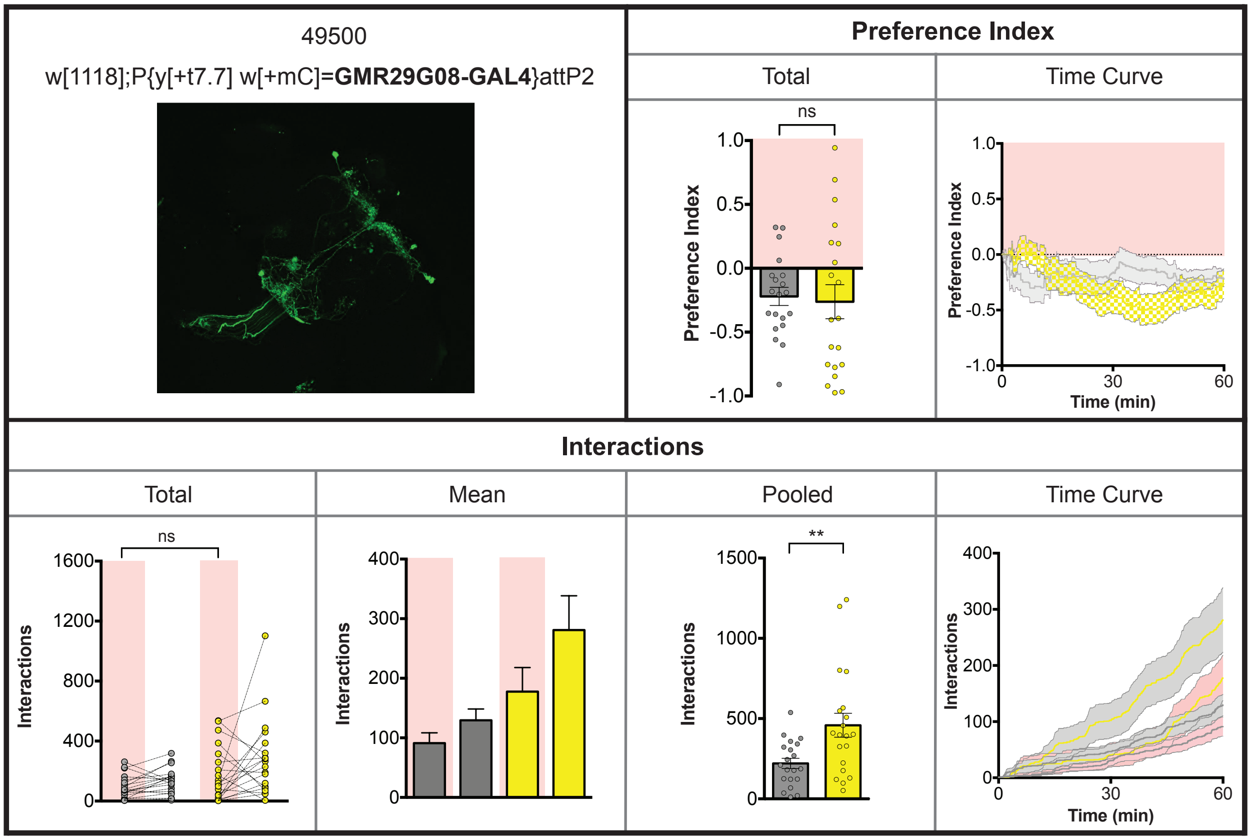
49500
w[1118];P{y[+t7.7] w[+mC]=**GMR29G08-GAL4**}attP2
Preference Index
Total
Time Curve
ns
Preference Index
Time (min)
Interactions
Total
Mean
Pooled
Time Curve
ns
**
Interactions
Time (min)

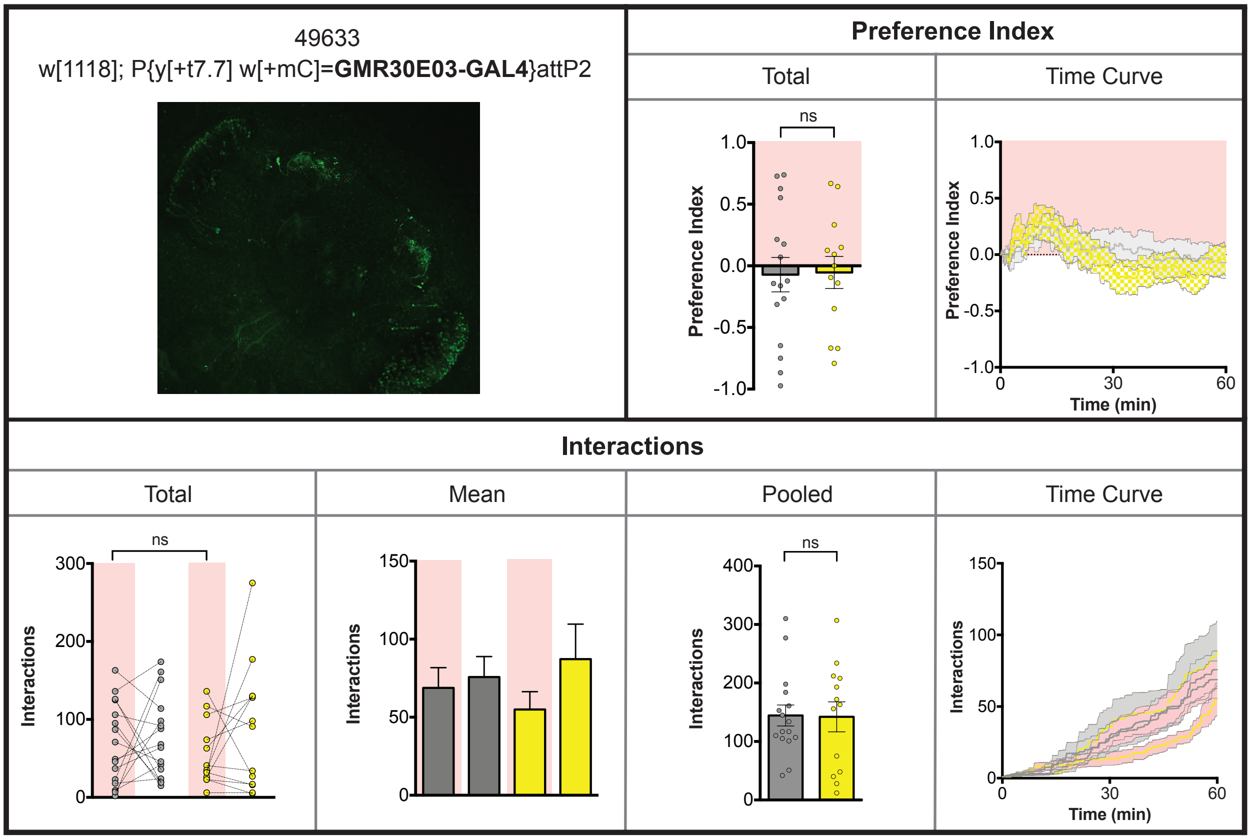
49633
w[1118]; P{y[+t7.7] w[+mC]=**GMR30E03-GAL4**}attP2
Preference Index
Total
Time Curve
ns
Preference Index
Time (min)
Interactions
Total
Mean
Pooled
Time Curve
ns
ns
Interactions
Time (min)

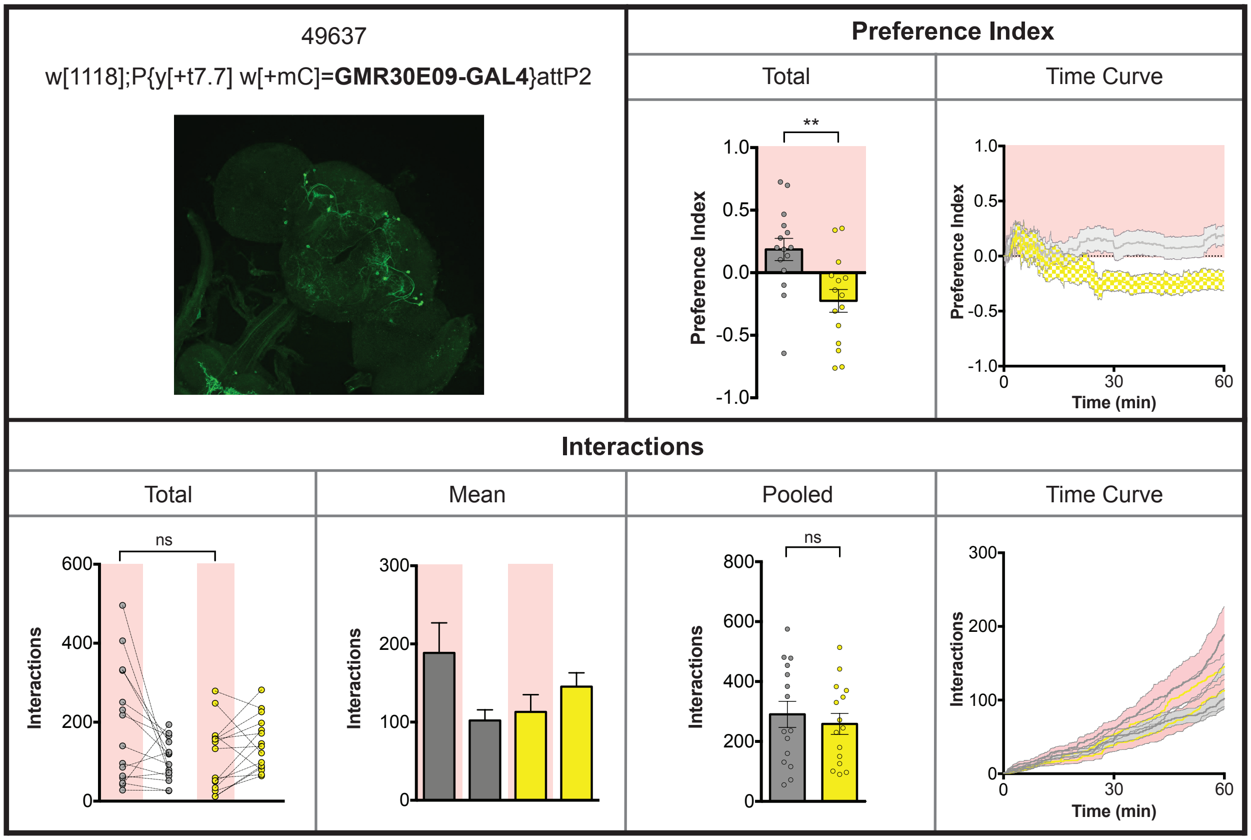
49637
w[1118];P{y[+t7.7] w[+mC]=GMR30E09-GAL4}attP2
Preference Index
Total
Time Curve
**
Preference Index
1.0
0.5
0.0
-0.5
-1.0
Time (min)
0
30
60
Interactions
Total
Mean
Pooled
Time Curve
ns
600
400
200
0
300
200
100
ns
800
Time (min)

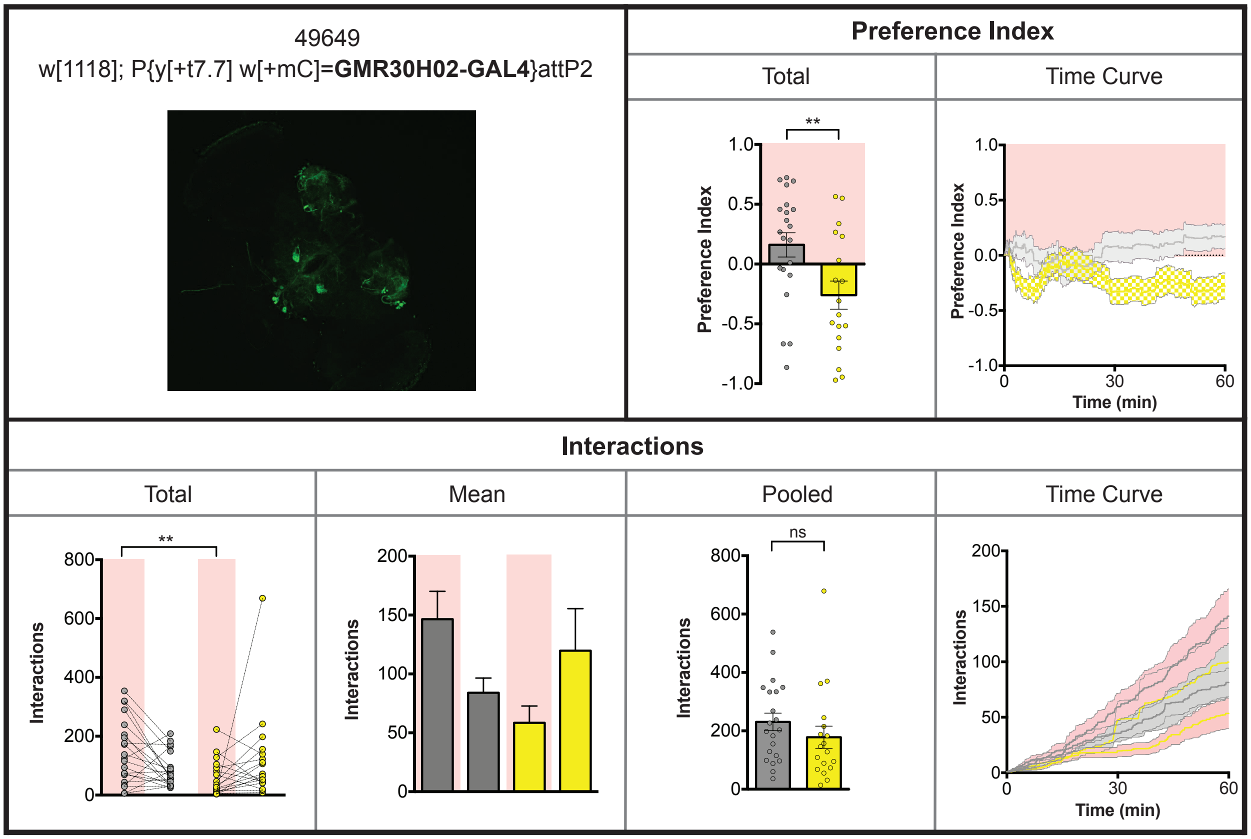
49649
w[1118]; P{y[+t7.7] w[+mC]=GMR30H02-GAL4}attP2
Preference Index
Total
Time Curve
**
Preference Index
1.0
0.5
0.0
-0.5
-1.0
Time (min)
0
30
60
Interactions
Total
Mean
Pooled
Time Curve
**
800
600
400
200
0
150
100
50
ns
Time (min)

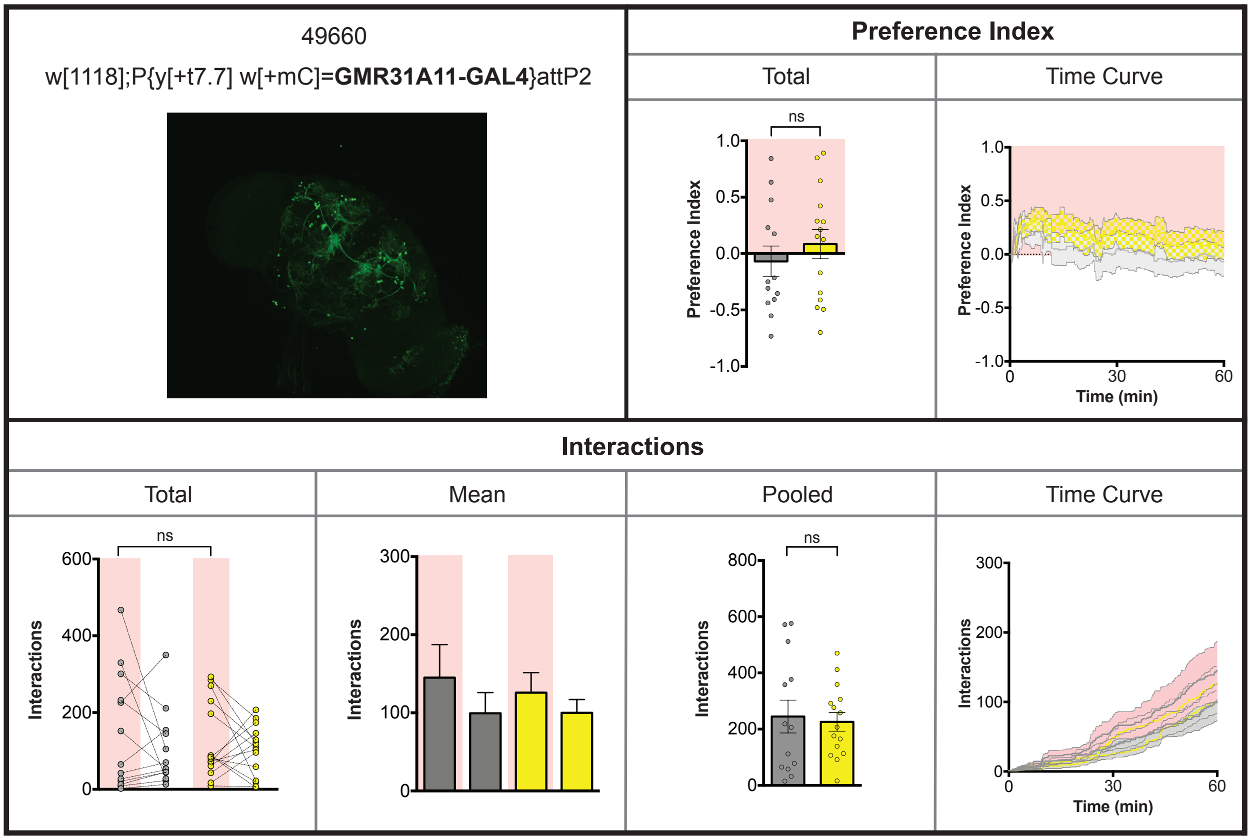
49660
w[1118];P{y[+t7.7] w[+mC]=GMR31A11-GAL4}attP2
Preference Index
Total
Time Curve
ns
Preference Index
Time (min)
Interactions
Total
Mean
Pooled
Time Curve
ns
ns
Interactions
Time (min)

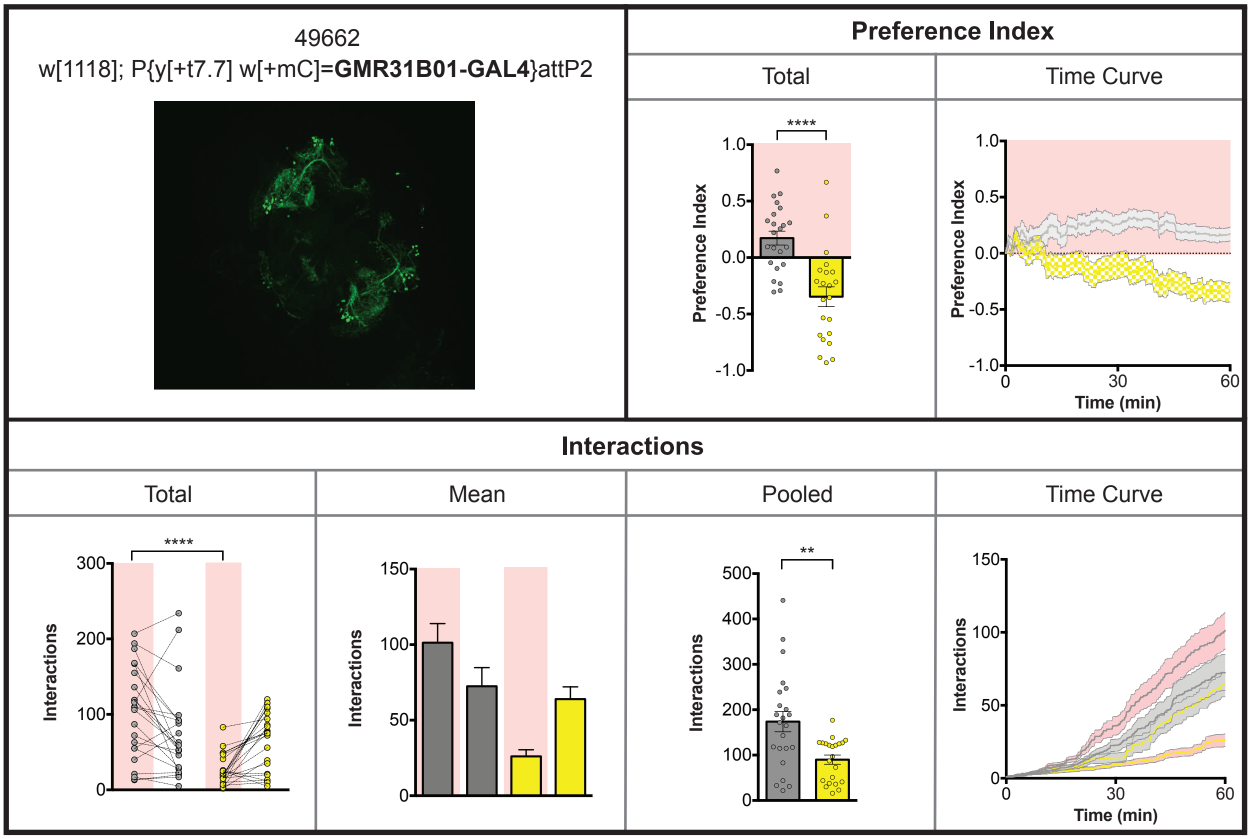
49662
w[1118]; P{y[+t7.7] w[+mC]=GMR31B01-GAL4}attP2
Preference Index
Total
Time Curve
****
Preference Index
Time (min)
Interactions
Total
Mean
Pooled
Time Curve
****
**
Interactions
Time (min)

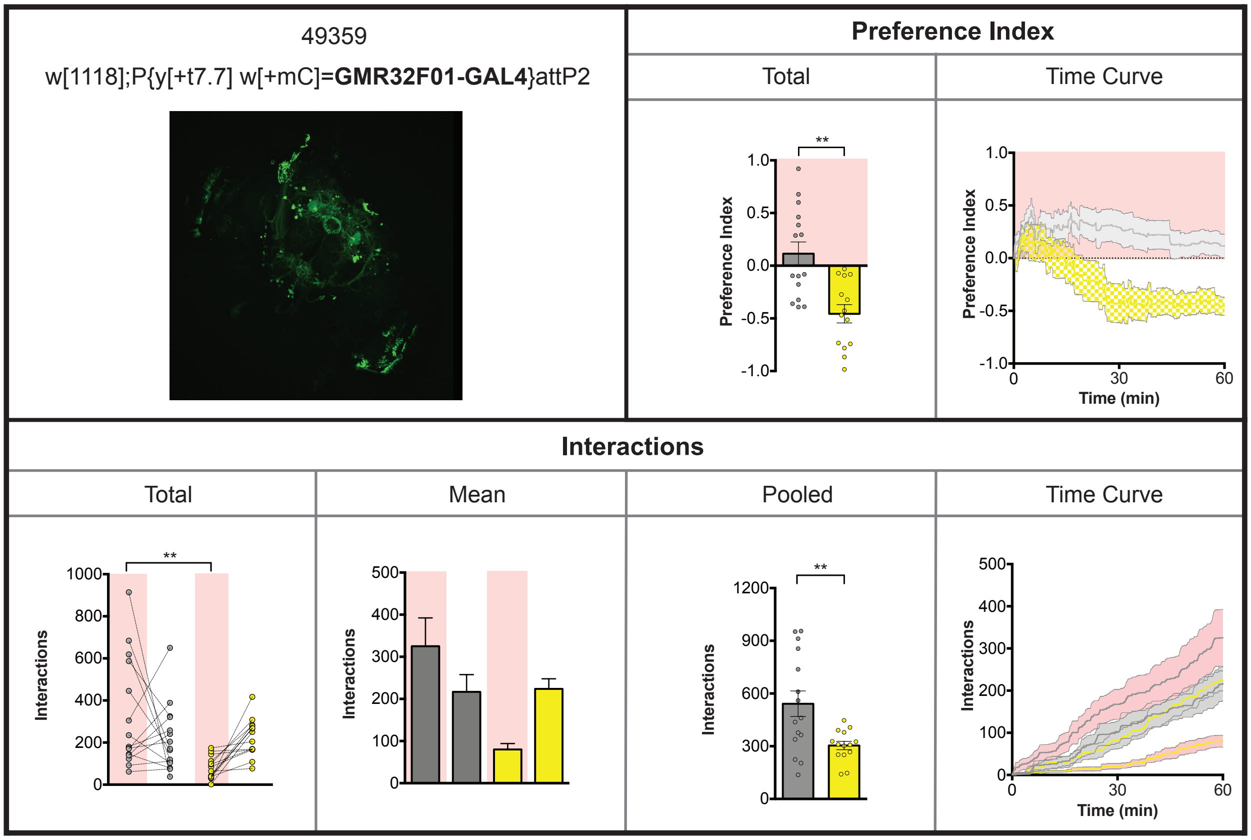
49359
w[1118];P{y[+t7.7] w[+mC]=**GMR32F01-GAL4**}attP2
Preference Index
Total
Time Curve
Preference Index
Time (min)
**
Interactions
Total
Mean
Pooled
Time Curve
Interactions
**
**
Time (min)

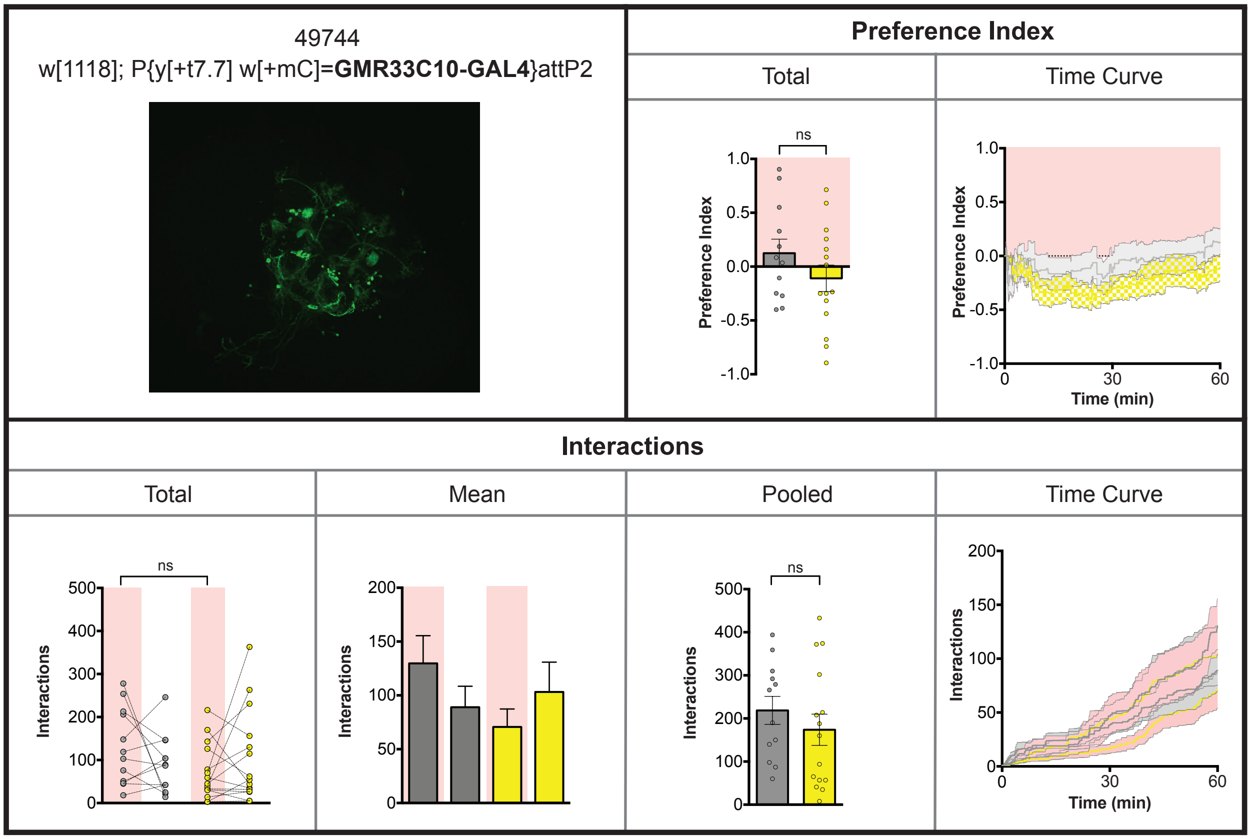
49744
w[1118]; P{y[+t7.7] w[+mC]=**GMR33C10-GAL4**}attP2
Preference Index
Total
Time Curve
Preference Index
Time (min)
ns
Interactions
Total
Mean
Pooled
Time Curve
Interactions
ns
ns
Time (min)

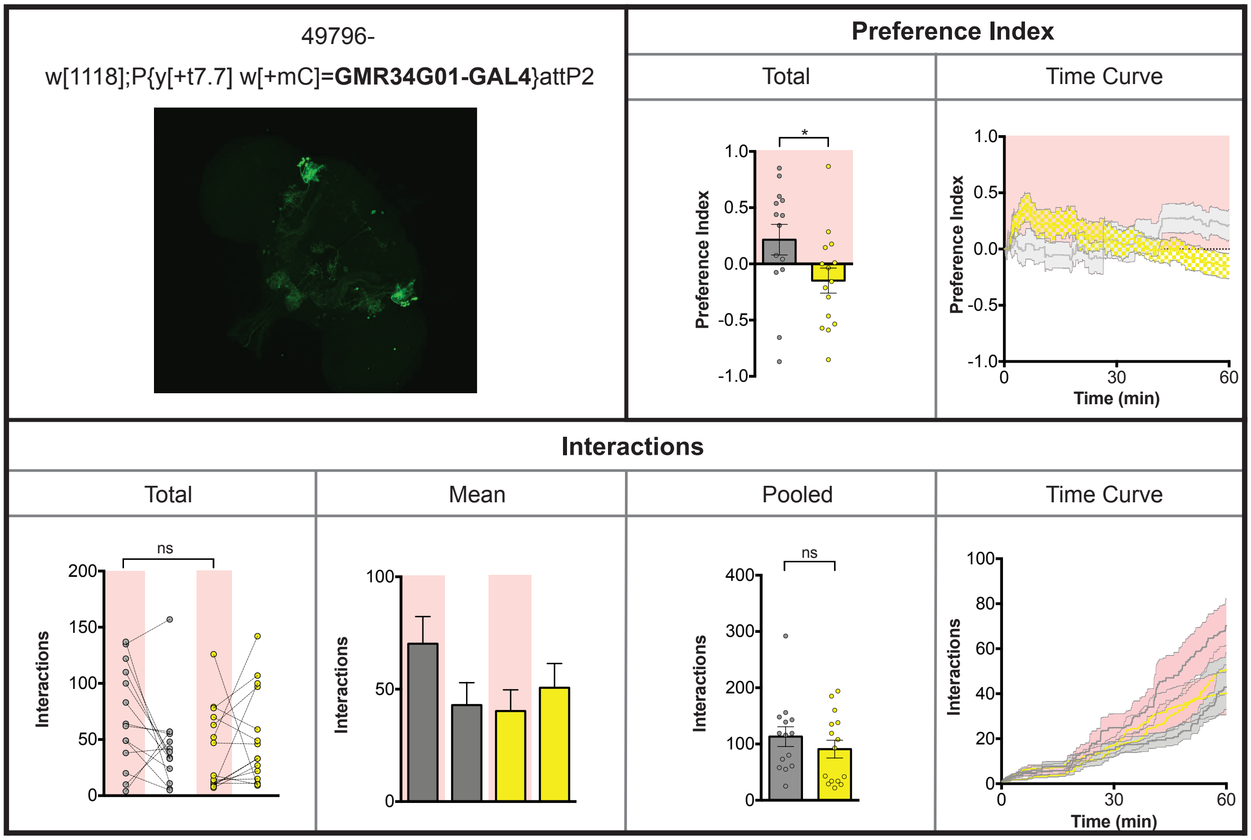
49796-
w[1118];P{y[+t7.7] w[+mC]=**GMR34G01-GAL4**}attP2
Preference Index
Total
Time Curve
Preference Index
*
1.0
0.5
0.0
-0.5
-1.0
Time (min)
0
30
60
Interactions
Total
Mean
Pooled
Time Curve
ns
200
150
100
50
0
400
300
Time (min)

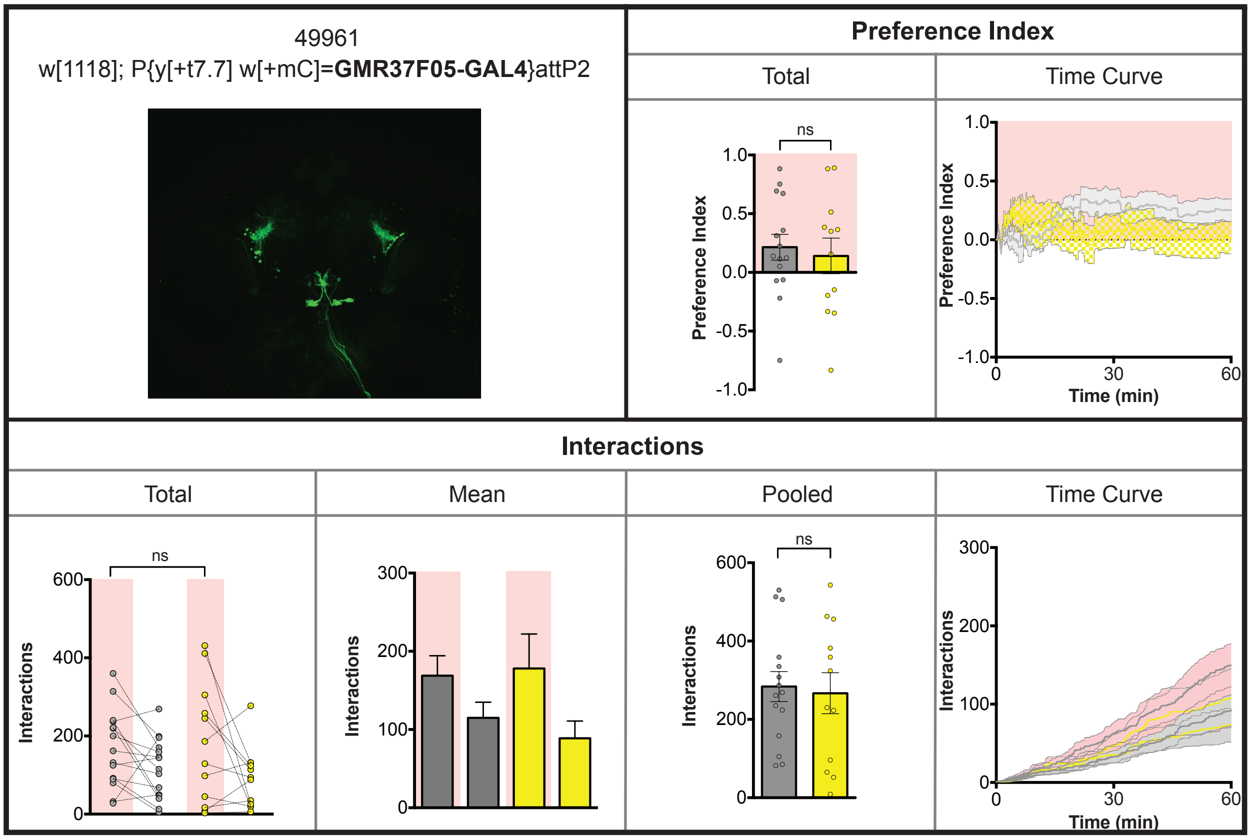
49961
w[1118]; P{y[+t7.7] w[+mC]=**GMR37F05-GAL4**}attP2
Preference Index
Total
Time Curve
Preference Index
ns
1.0
0.5
0.0
-0.5
-1.0
Time (min)
0
30
60
Interactions
Total
Mean
Pooled
Time Curve
ns
600
400
200
0
300
100
Time (min)

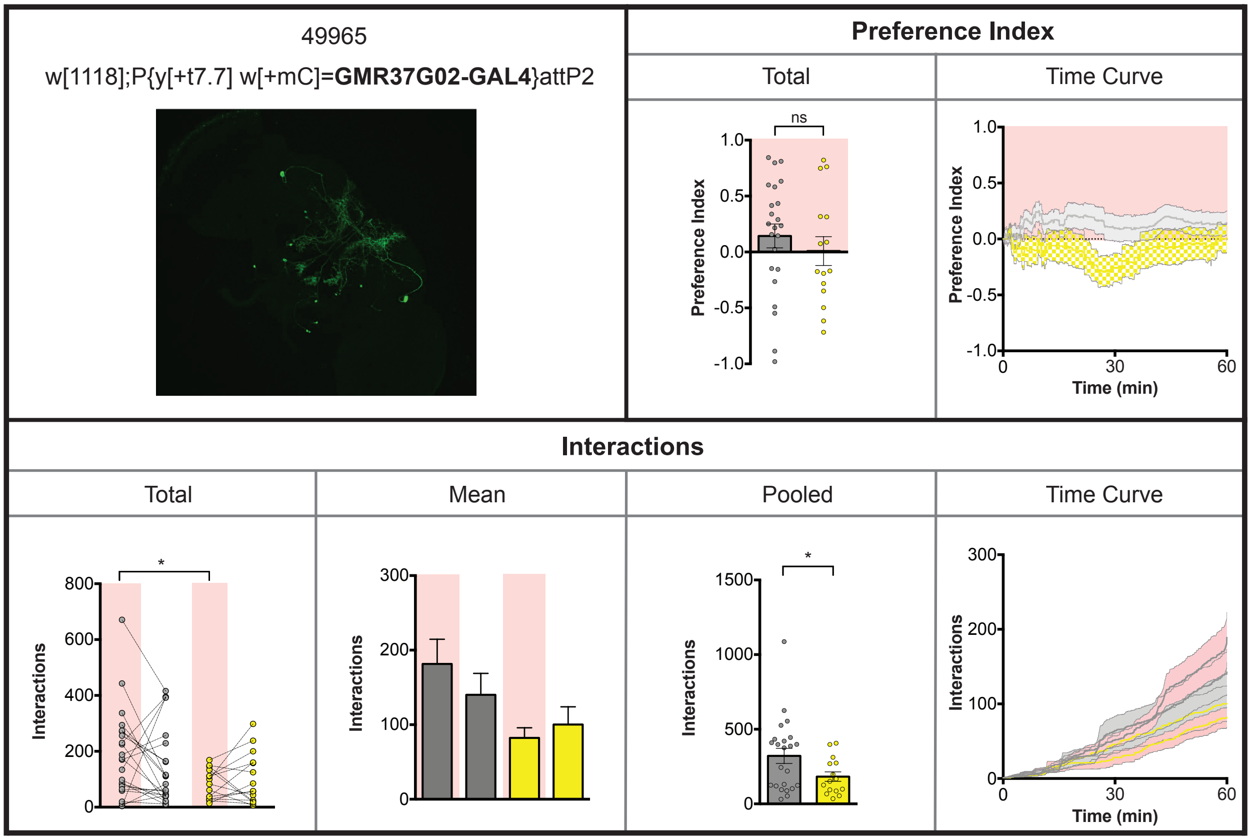
49965
w[1118];P{y[+t7.7] w[+mC]=GMR37G02-GAL4}attP2
Preference Index
Total
Time Curve
ns
Preference Index
1.0
0.5
0.0
-0.5
-1.0
0
30
60
Time (min)
Interactions
Total
Mean
Pooled
Time Curve
*
800
600
400
200
0
300
200
100
1500
1000
500
Interactions
*
Time (min)

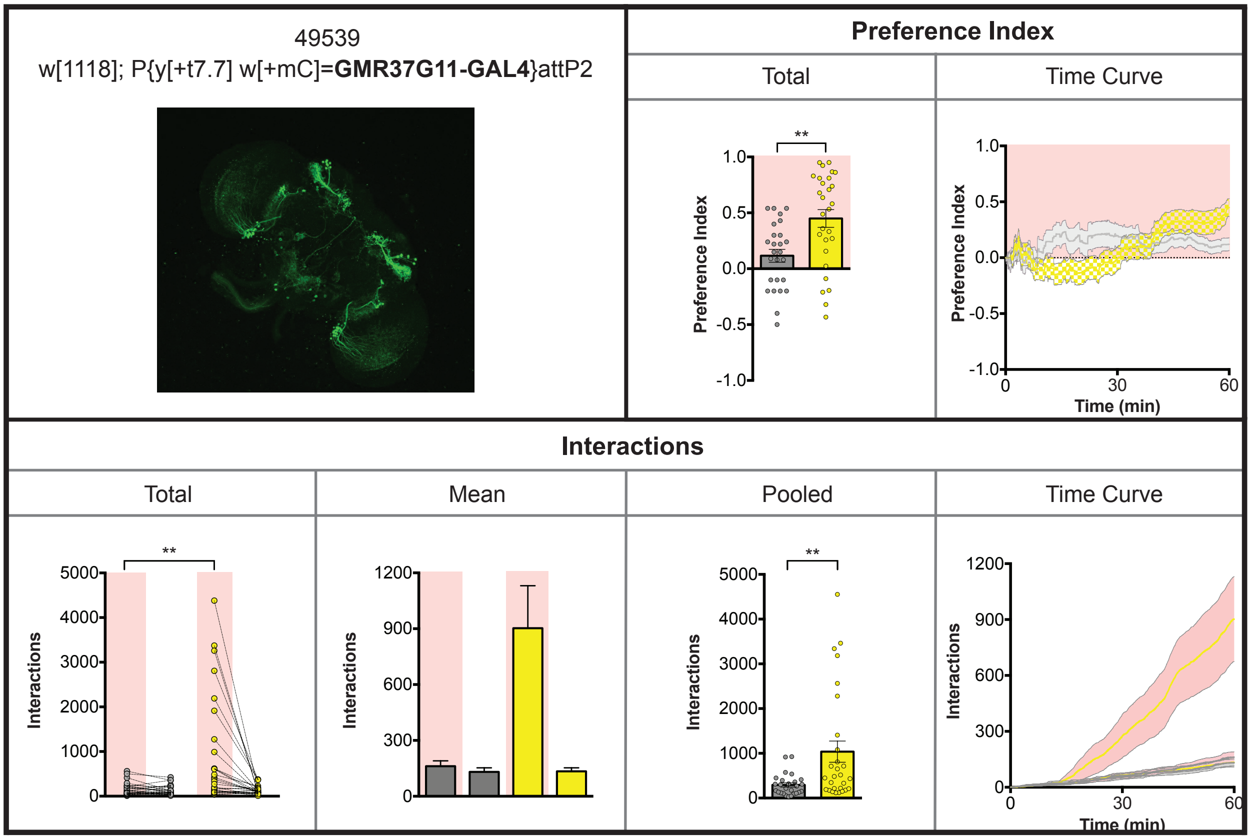
49539
w[1118]; P{y[+t7.7] w[+mC]=GMR37G11-GAL4}attP2
Preference Index
Total
Time Curve
**
Preference Index
1.0
0.5
0.0
-0.5
-1.0
0
30
60
Time (min)
Interactions
Total
Mean
Pooled
Time Curve
**
5000
4000
3000
2000
1000
0
1200
900
600
300
Interactions
**
Time (min)

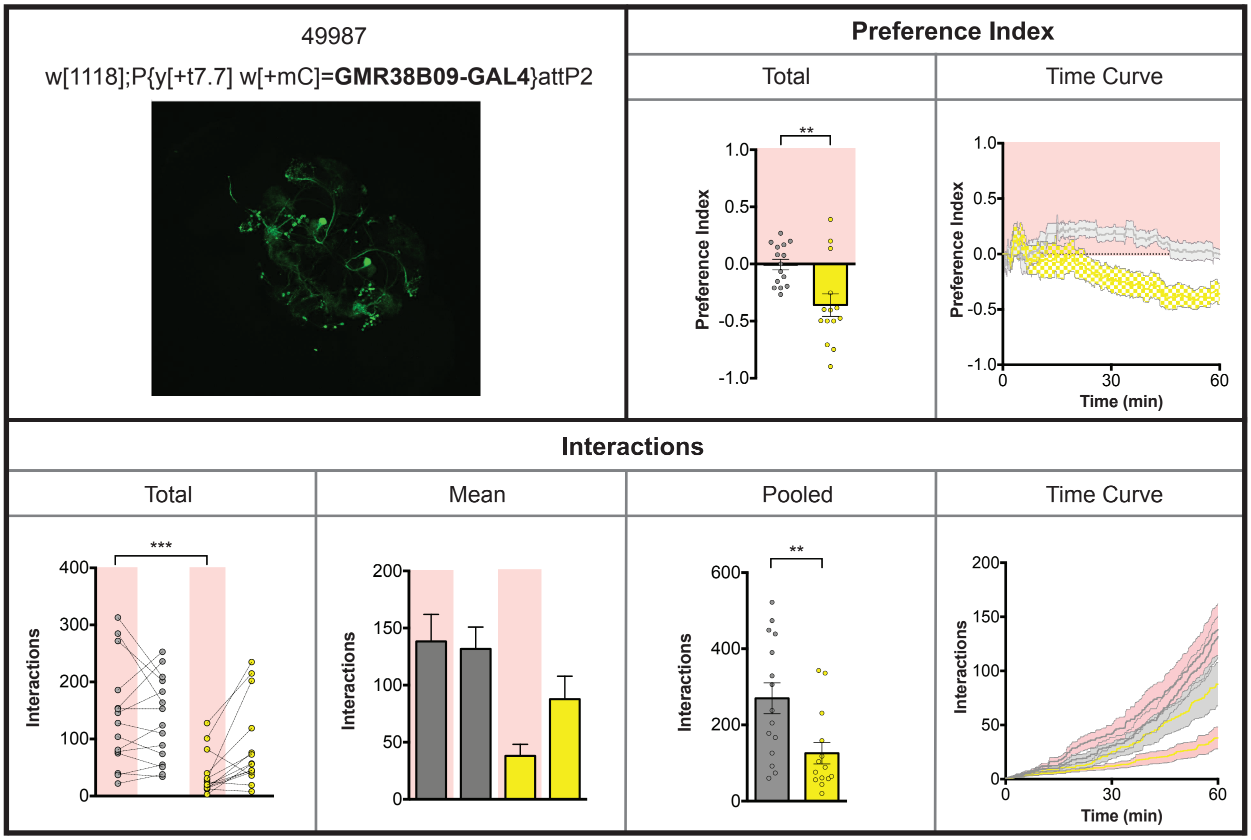
49987
w[1118];P{y[+t7.7] w[+mC]=GMR38B09-GAL4}attP2
Preference Index
Total
Time Curve
**
Preference Index
1.0
0.5
0.0
-0.5
-1.0
0
30
60
Time (min)
Interactions
Total
Mean
Pooled
Time Curve
***
Interactions
400
300
200
100
0
150
50
**
600
Time (min)

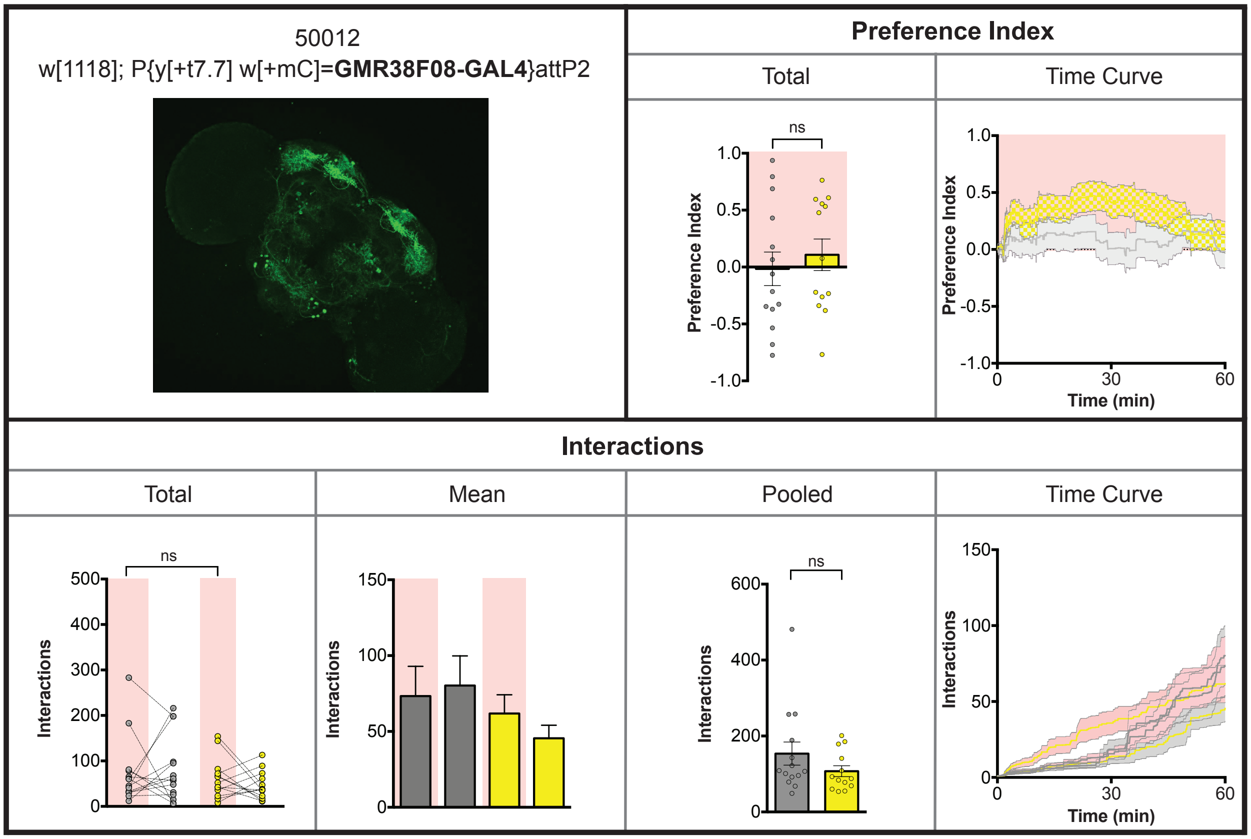
50012
w[1118]; P{y[+t7.7] w[+mC]=GMR38F08-GAL4}attP2
Preference Index
Total
Time Curve
ns
Preference Index
1.0
0.5
0.0
-0.5
-1.0
0
30
60
Time (min)
Interactions
Total
Mean
Pooled
Time Curve
ns
Interactions
500
400
300
200
100
0
150
50
600
Time (min)

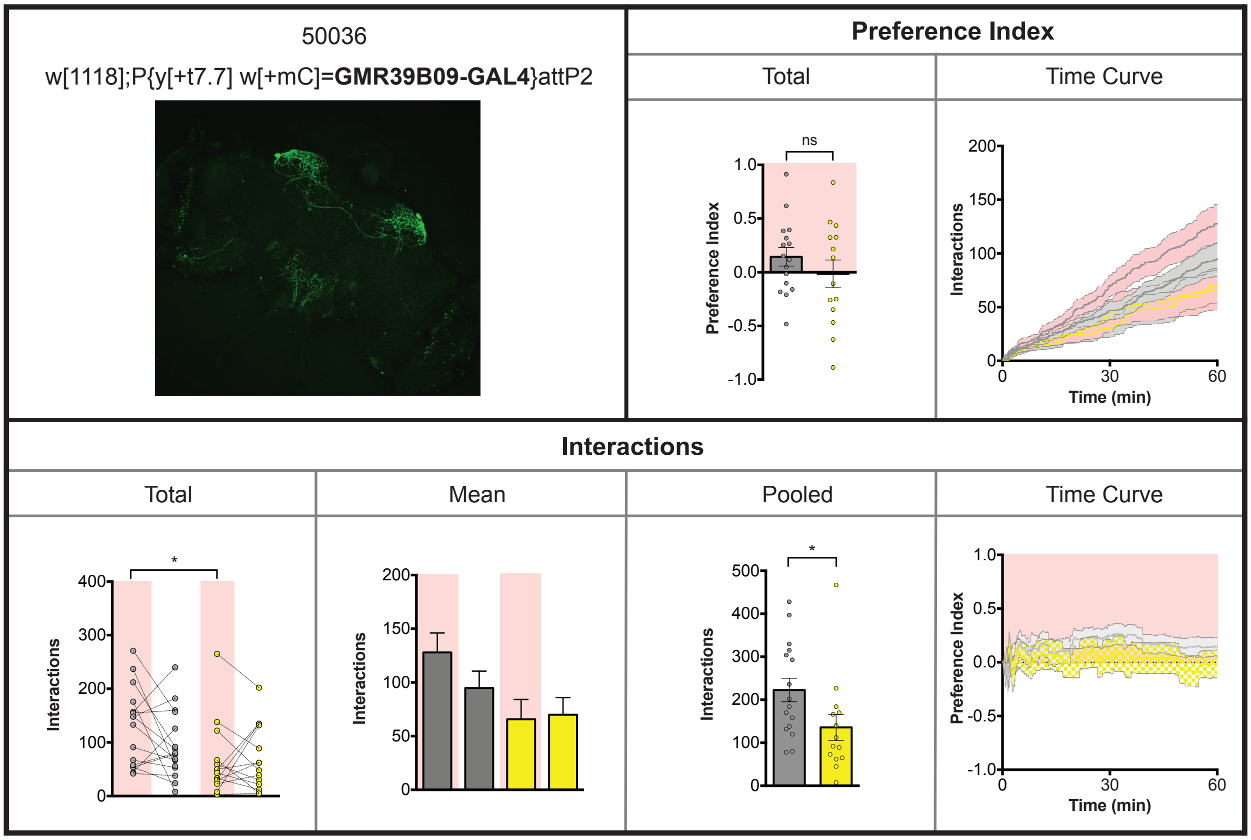
50036
w[1118];P{y[+t7.7] w[+mC]=GMR39B09-GAL4}attP2
Preference Index
Total
Time Curve
ns
Preference Index
1.0
0.5
0.0
-0.5
-1.0
Interactions
200
150
100
50
0
30
60
Time (min)
Interactions
Total
Mean
Pooled
Time Curve
*
400
300
200
100
Interactions
200
150
100
50
500
*
Preference Index

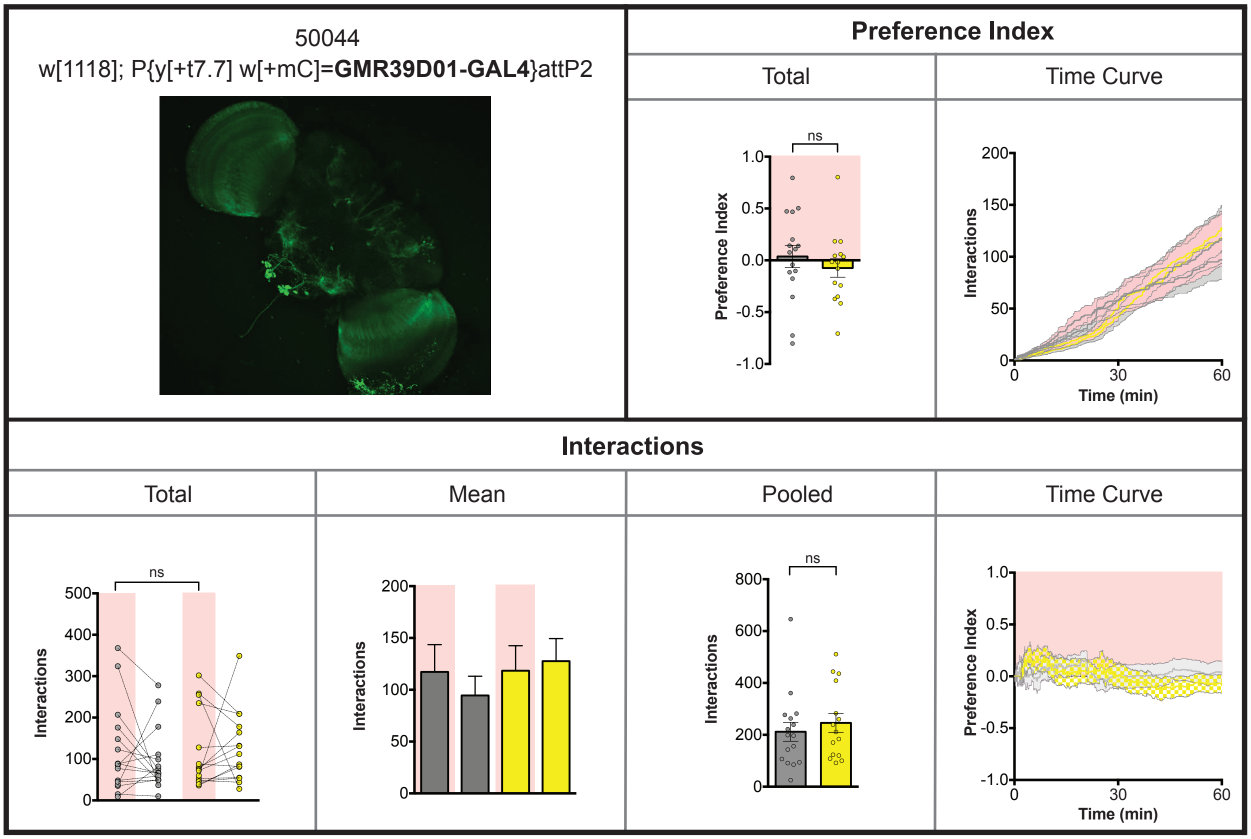
50044
w[1118]; P{y[+t7.7] w[+mC]=GMR39D01-GAL4}attP2
Preference Index
Total
Time Curve
ns
Preference Index
1.0
0.5
0.0
-0.5
-1.0
Interactions
200
150
100
50
0
30
60
Time (min)
Interactions
Total
Mean
Pooled
Time Curve
ns
500
400
300
200
100
Interactions
800
600
400
200
Preference Index

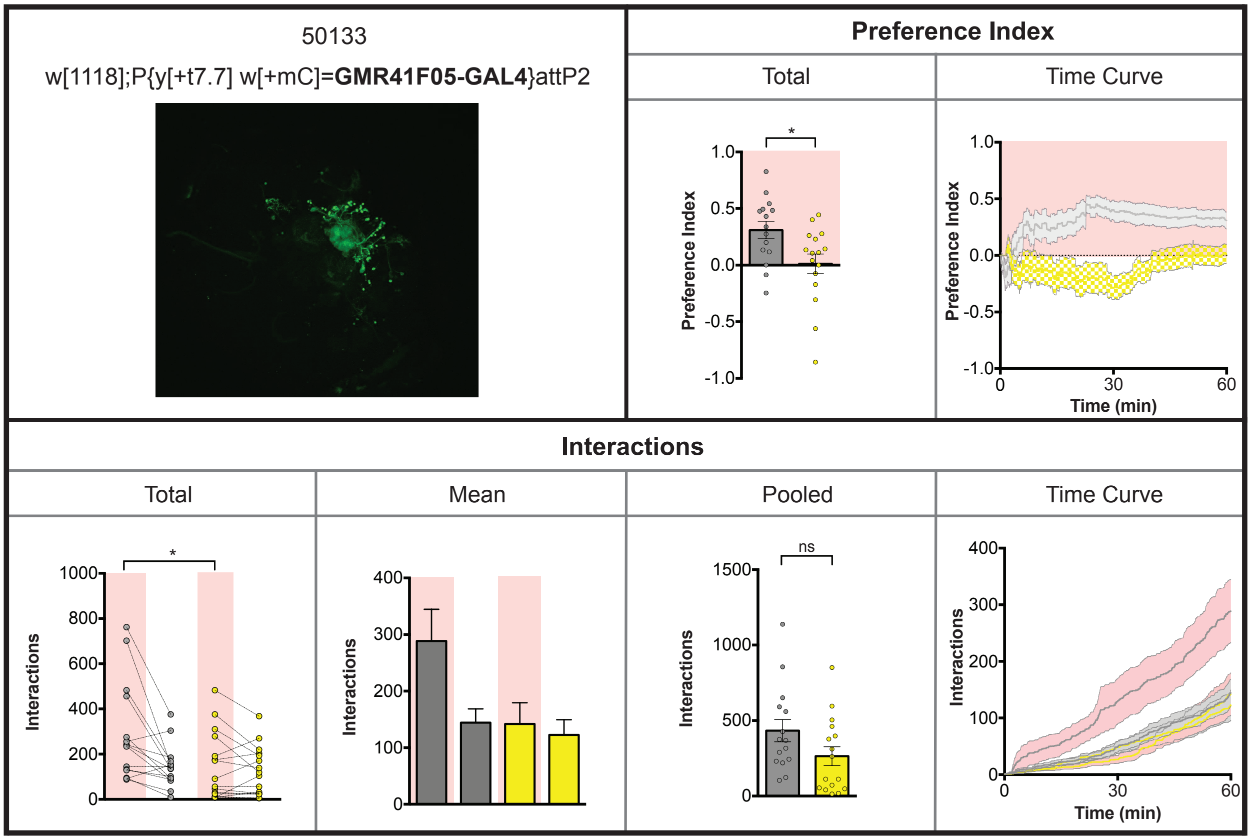
50133
w[1118];P{y[+t7.7] w[+mC]=GMR41F05-GAL4}attP2
Preference Index
Total
Time Curve
Preference Index
*
Time (min)
Interactions
Total
Mean
Pooled
Time Curve
Interactions
ns
Time (min)

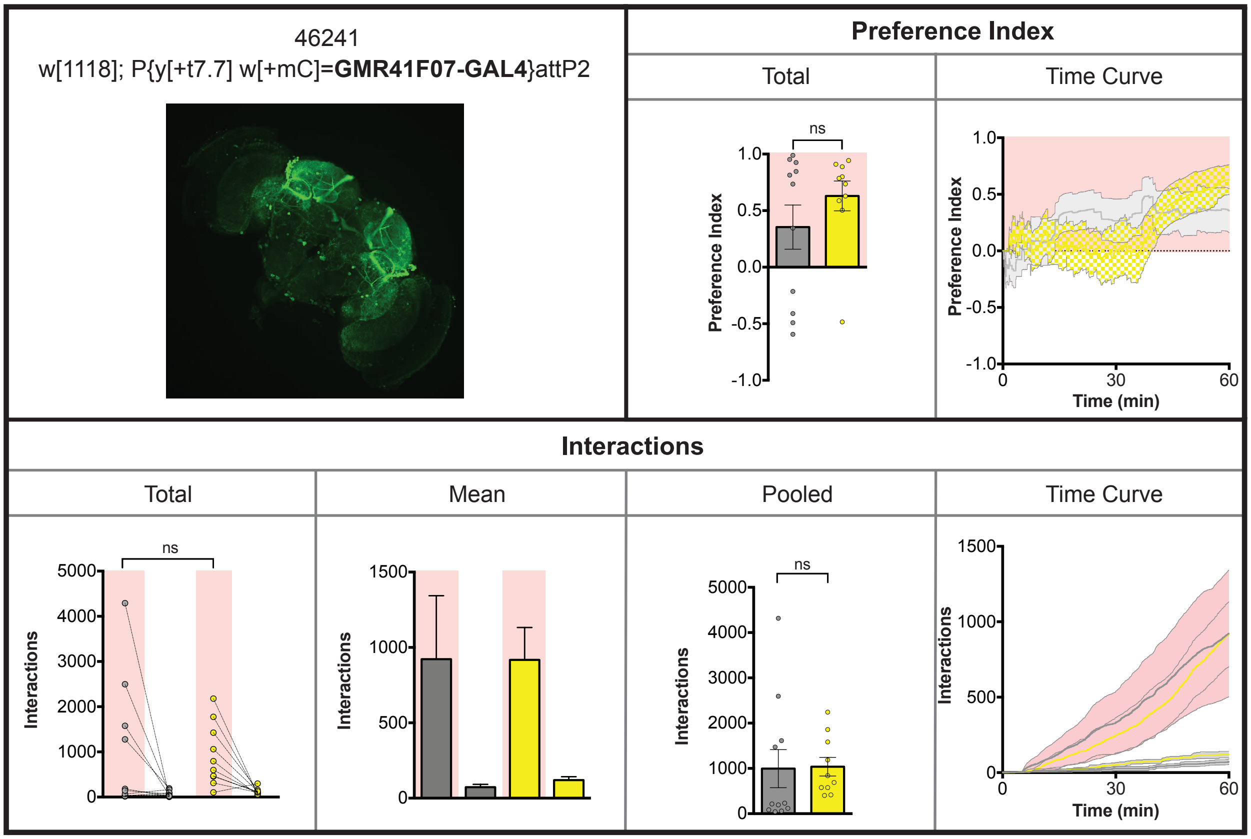
46241
w[1118]; P{y[+t7.7] w[+mC]=GMR41F07-GAL4}attP2
Preference Index
Total
Time Curve
Preference Index
ns
Time (min)
Interactions
Total
Mean
Pooled
Time Curve
Interactions
ns
Time (min)

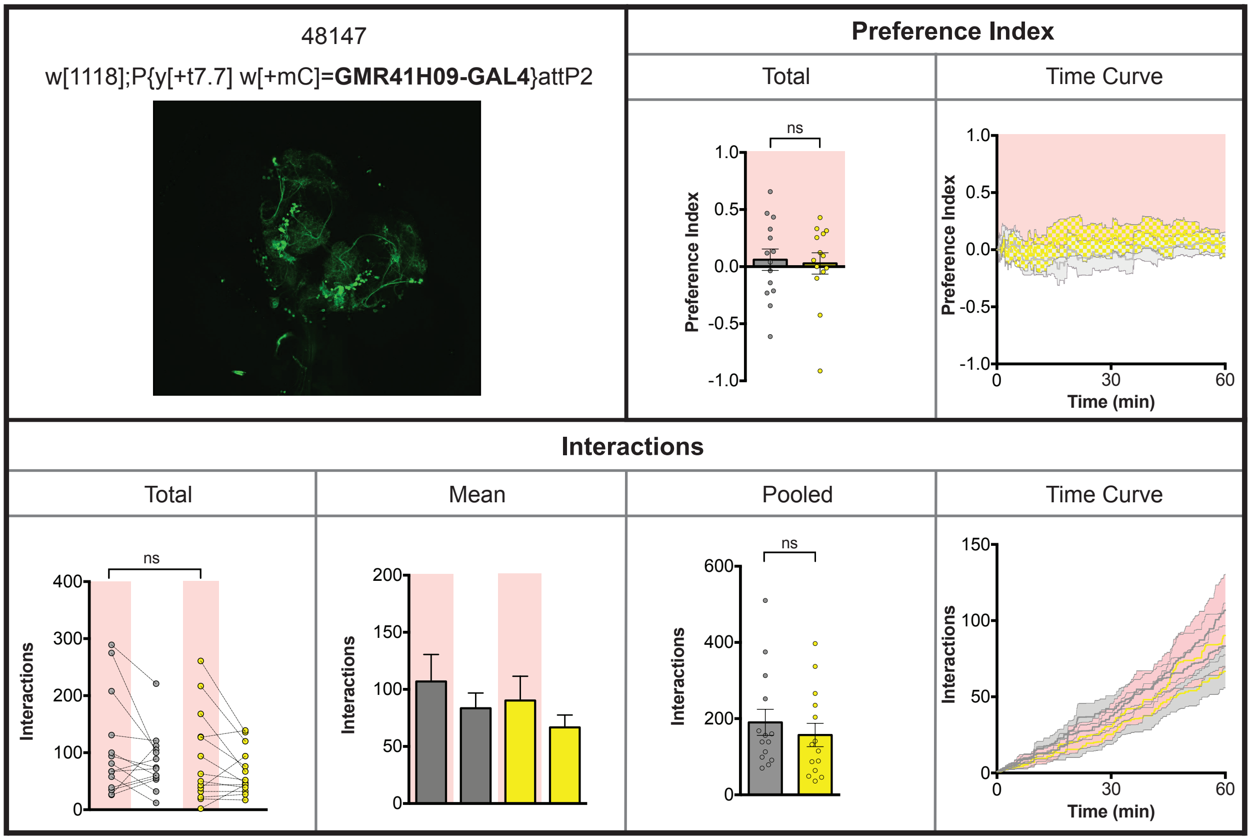
48147
w[1118];P{y[+t7.7] w[+mC]=**GMR41H09-GAL4**}attP2
Preference Index
Total
Time Curve
ns
Preference Index
1.0
0.5
0.0
-0.5
-1.0
Time (min)
0
30
60
Interactions
Total
Mean
Pooled
Time Curve
ns
Interactions
400
300
200
100
0
200
150
100
50
0
ns
600
400
200
0
150
100
50
0
Time (min)
0
30
60

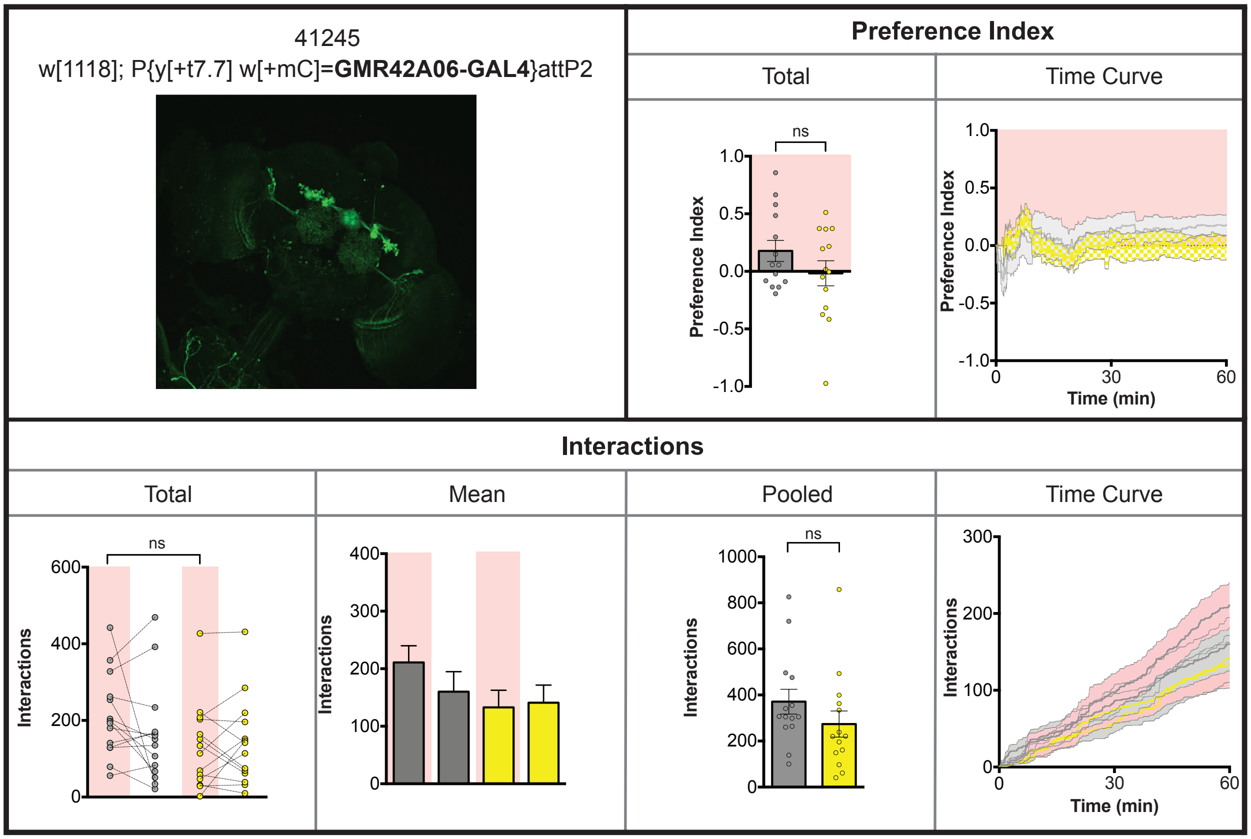
41245
w[1118]; P{y[+t7.7] w[+mC]=**GMR42A06-GAL4**}attP2
Preference Index
Total
Time Curve
ns
Preference Index
1.0
0.5
0.0
-0.5
-1.0
Time (min)
0
30
60
Interactions
Total
Mean
Pooled
Time Curve
ns
Interactions
600
400
200
0
400
300
200
100
0
ns
1000
800
600
400
200
0
300
200
100
0
Time (min)
0
30
60

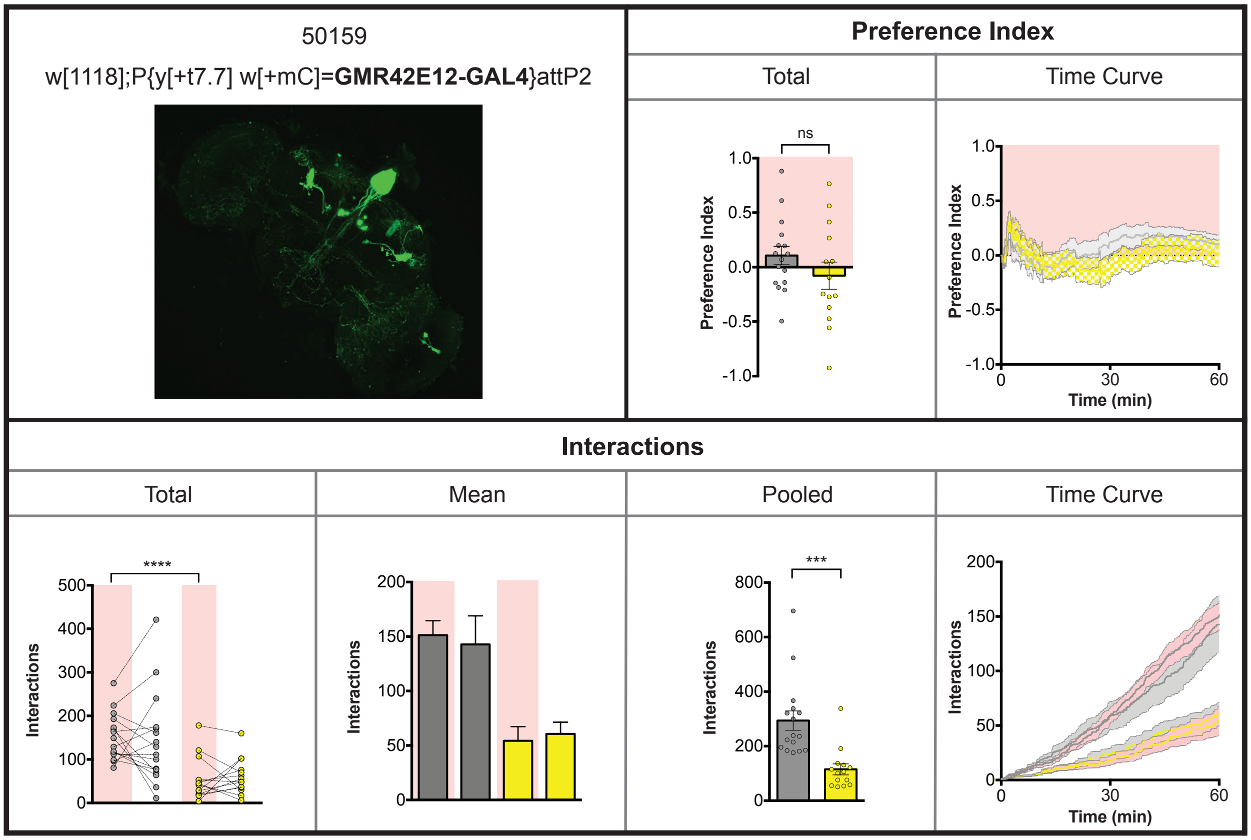
50159
w[1118];P{y[+t7.7] w[+mC]=**GMR42E12-GAL4**}attP2
Preference Index
Total
Time Curve
ns
Preference Index
1.0
0.5
0.0
-0.5
-1.0
Time (min)
0
30
60
Interactions
Total
Mean
Pooled
Time Curve
****
***
Interactions
500
400
300
200
100
0
150
50
800
600

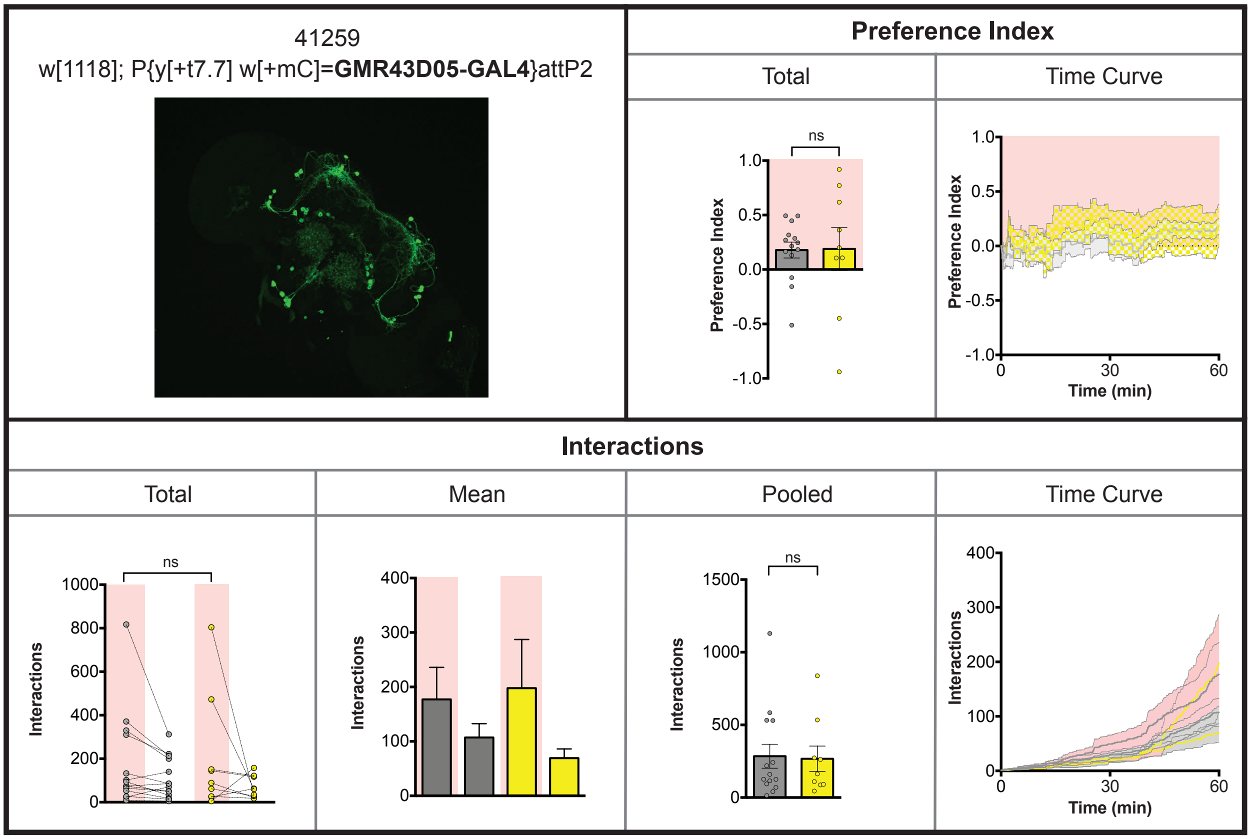
41259
w[1118]; P{y[+t7.7] w[+mC]=**GMR43D05-GAL4**}attP2
Preference Index
Total
Time Curve
ns
Preference Index
1.0
0.5
0.0
-0.5
-1.0
Time (min)
0
30
60
Interactions
Total
Mean
Pooled
Time Curve
ns
Interactions
1000
800
600
400
200
0
300
100
1500
500

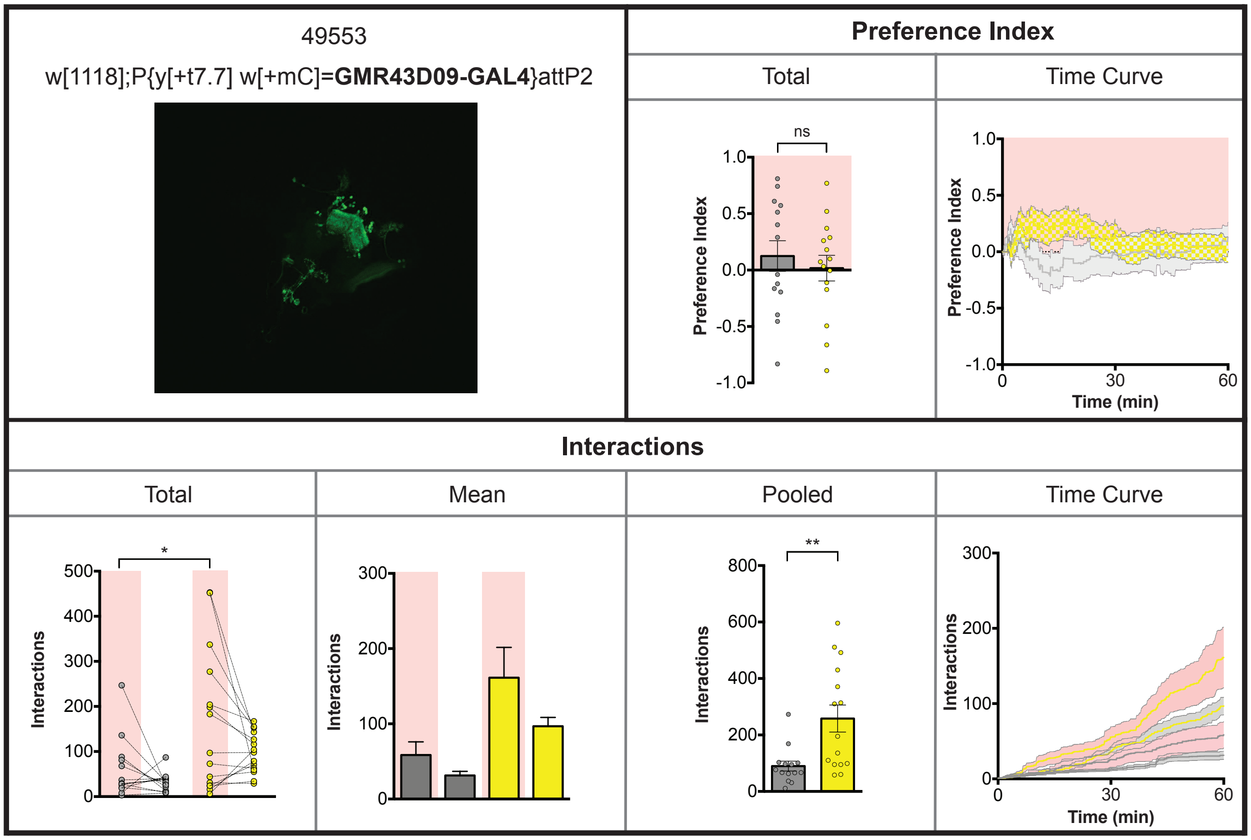
49553
w[1118];P{y[+t7.7] w[+mC]=GMR43D09-GAL4}attP2
Preference Index
Total
Time Curve
ns
Preference Index
1.0
0.5
0.0
-0.5
-1.0
0
30
60
Time (min)
Interactions
Total
Mean
Pooled
Time Curve
*
**
500
400
300
200
100
0
800
600
Interactions

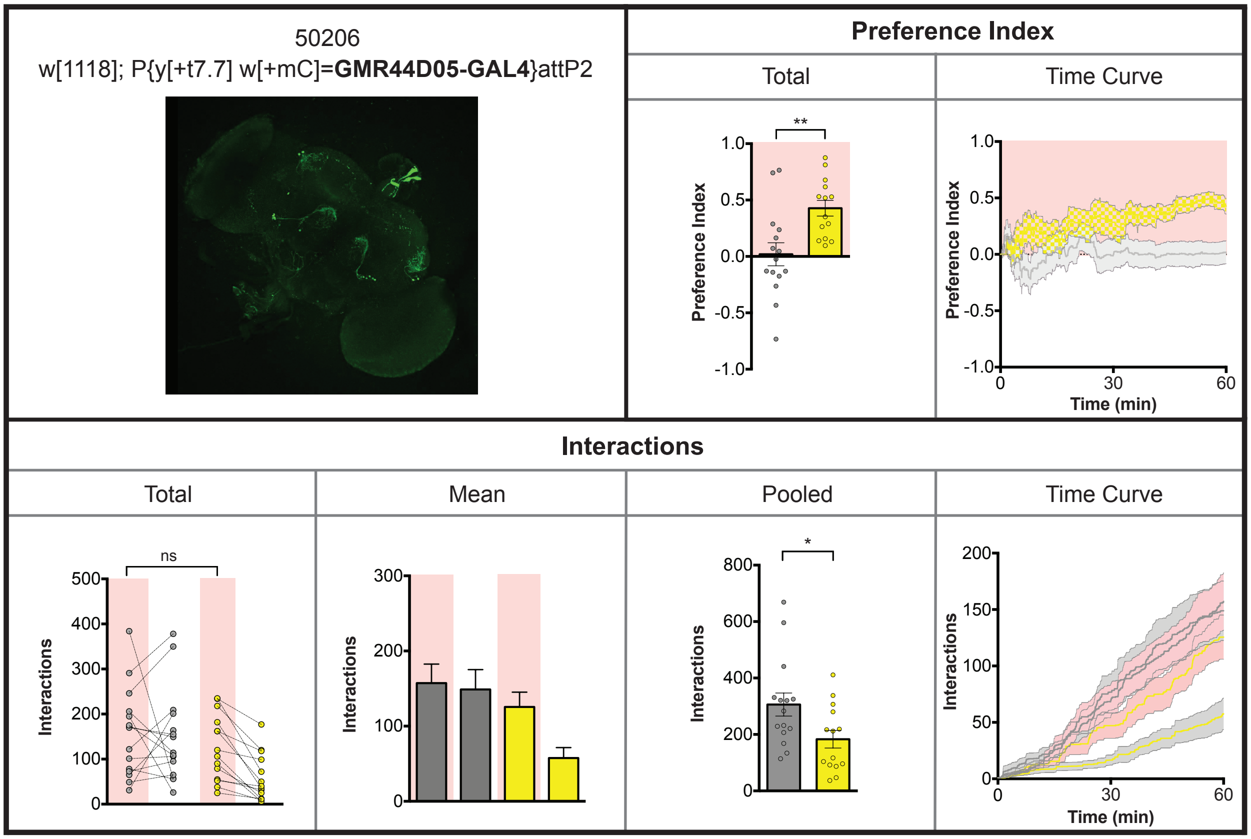
50206
w[1118]; P{y[+t7.7] w[+mC]=GMR44D05-GAL4}attP2
Preference Index
Total
Time Curve
**
Preference Index
1.0
0.5
0.0
-0.5
-1.0
0
30
60
Time (min)
Interactions
Total
Mean
Pooled
Time Curve
ns
*
500
400
300
200
150
100
50
0
800
600
Interactions

50229

w[1118];P{y[+t7.7] w[+mC]=**GMR45E03-GAL4**}attP2

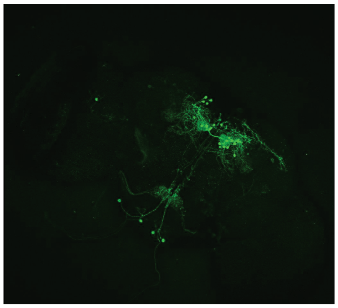

### Preference Index

Total

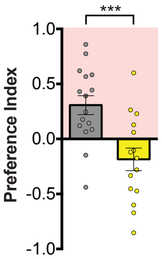


Time Curve

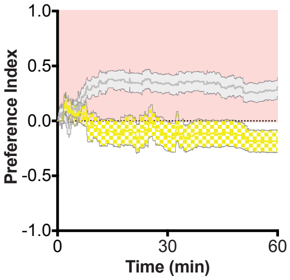


### Interactions

Total

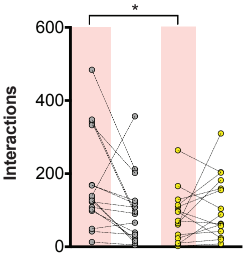


Mean

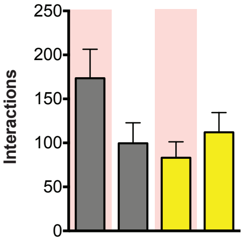


Pooled

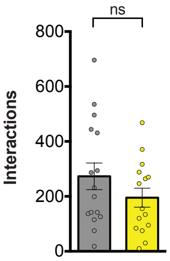


Time Curve

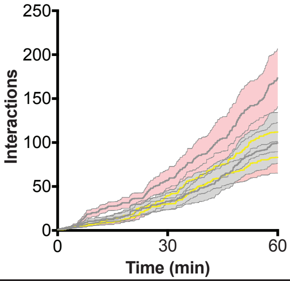


50244

w[1118]; P{y[+t7.7] w[+mC]=**GMR45G06-GAL4**}attP2

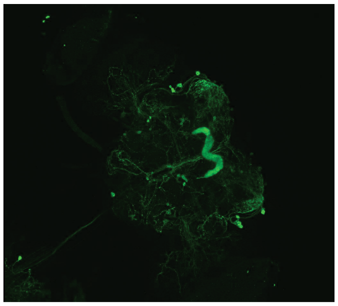

### Preference Index

Total

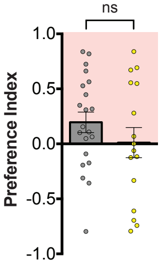


Time Curve

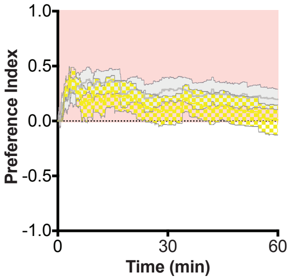


### Interactions

Total

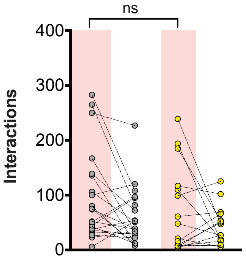


Mean

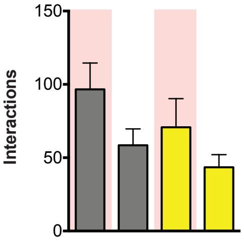


Pooled

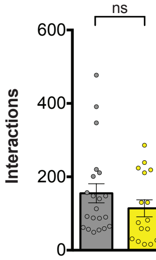


Time Curve

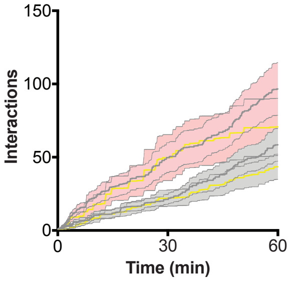

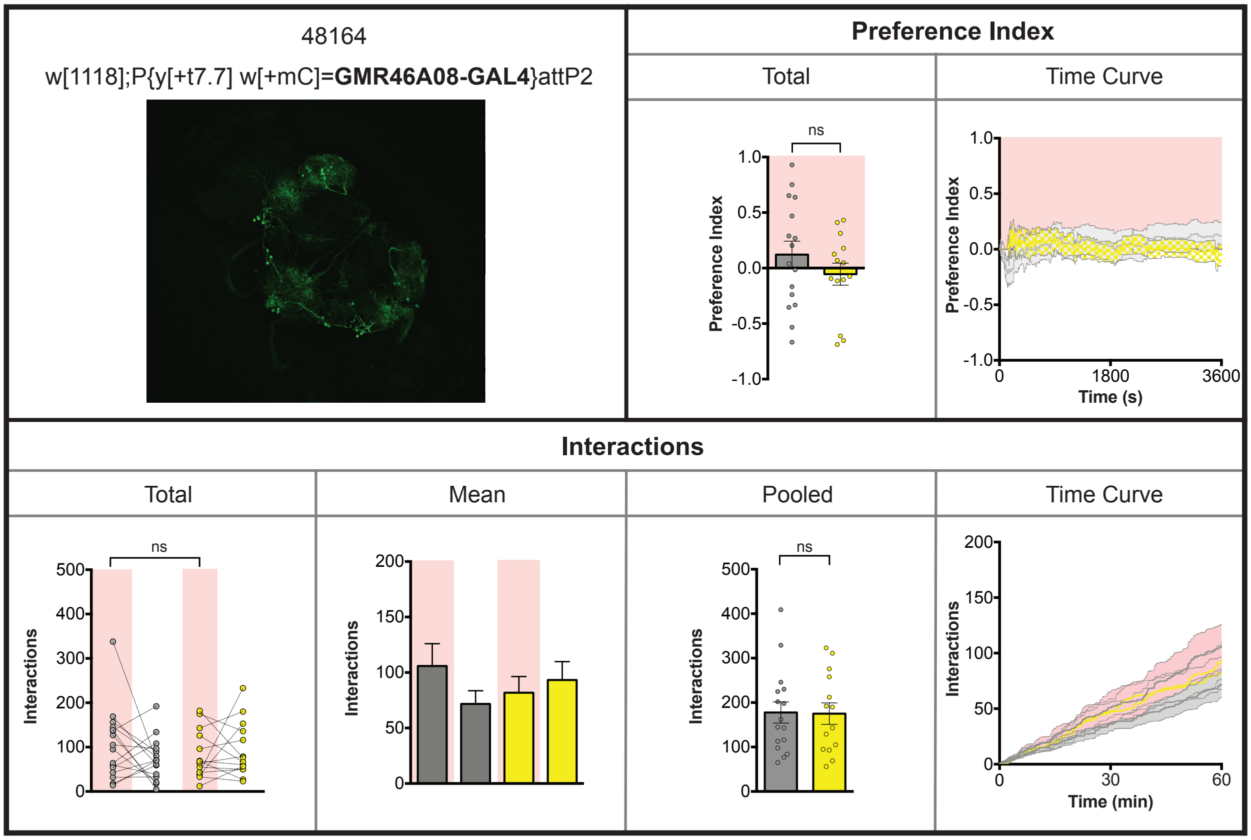
48164
w[1118];P{y[+t7.7] w[+mC]=GMR46A08-GAL4}attP2
Preference Index
Total
Time Curve
ns
Preference Index
Time (s)
Interactions
Total
Mean
Pooled
Time Curve
ns
Interactions
Time (min)

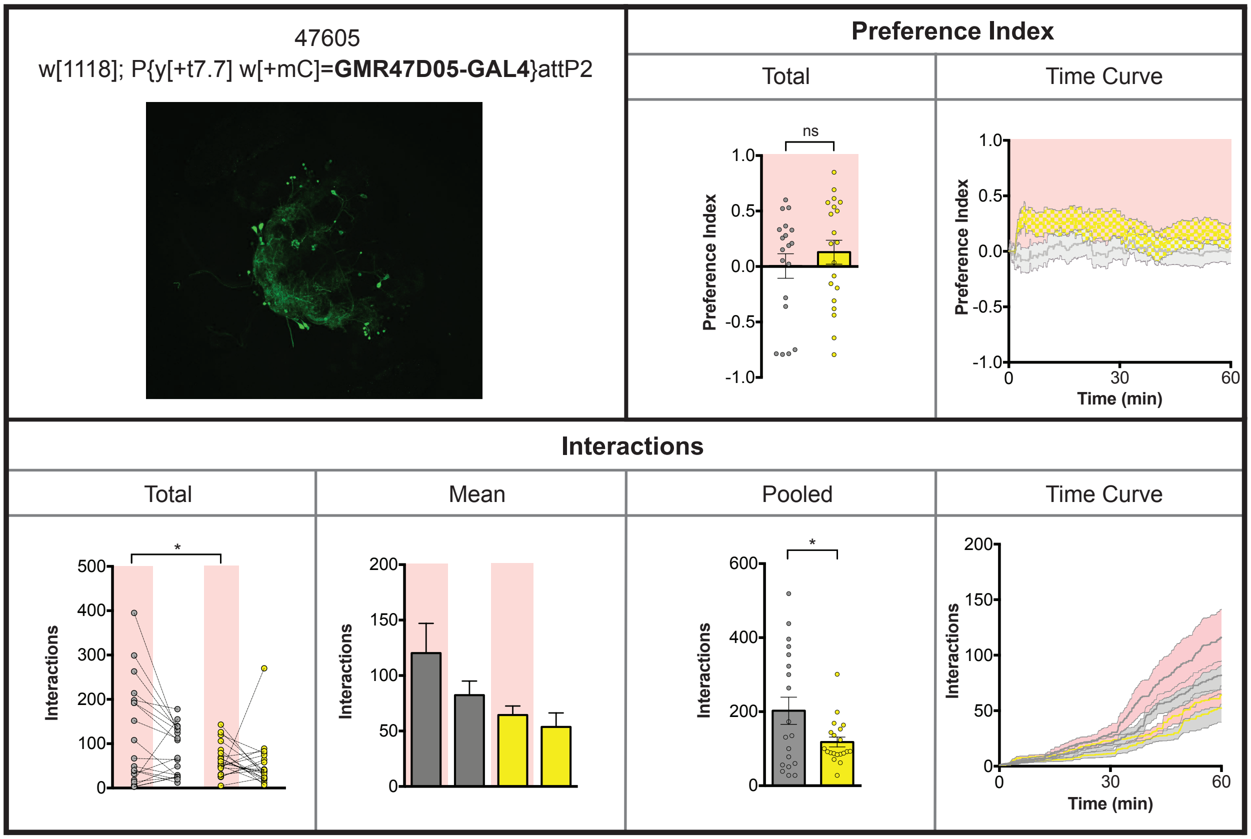
47605
w[1118]; P{y[+t7.7] w[+mC]=GMR47D05-GAL4}attP2
Preference Index
Total
Time Curve
ns
Preference Index
Time (min)
Interactions
Total
Mean
Pooled
Time Curve
*
Interactions
Time (min)

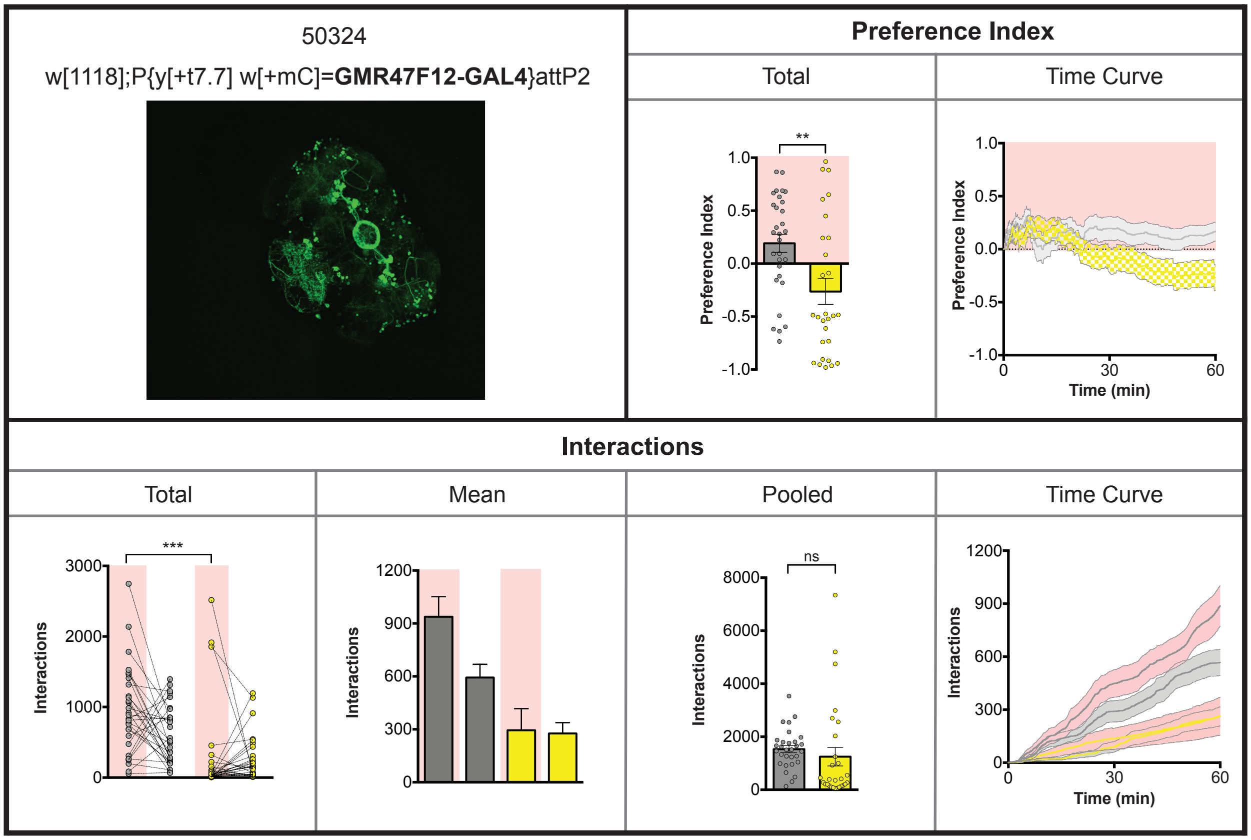
50324
w[1118];P{y[+t7.7] w[+mC]=GMR47F12-GAL4}attP2
Preference Index
Total
Time Curve
Preference Index
**
Time (min)
Interactions
Total
Mean
Pooled
Time Curve
Interactions
***
ns
Time (min)

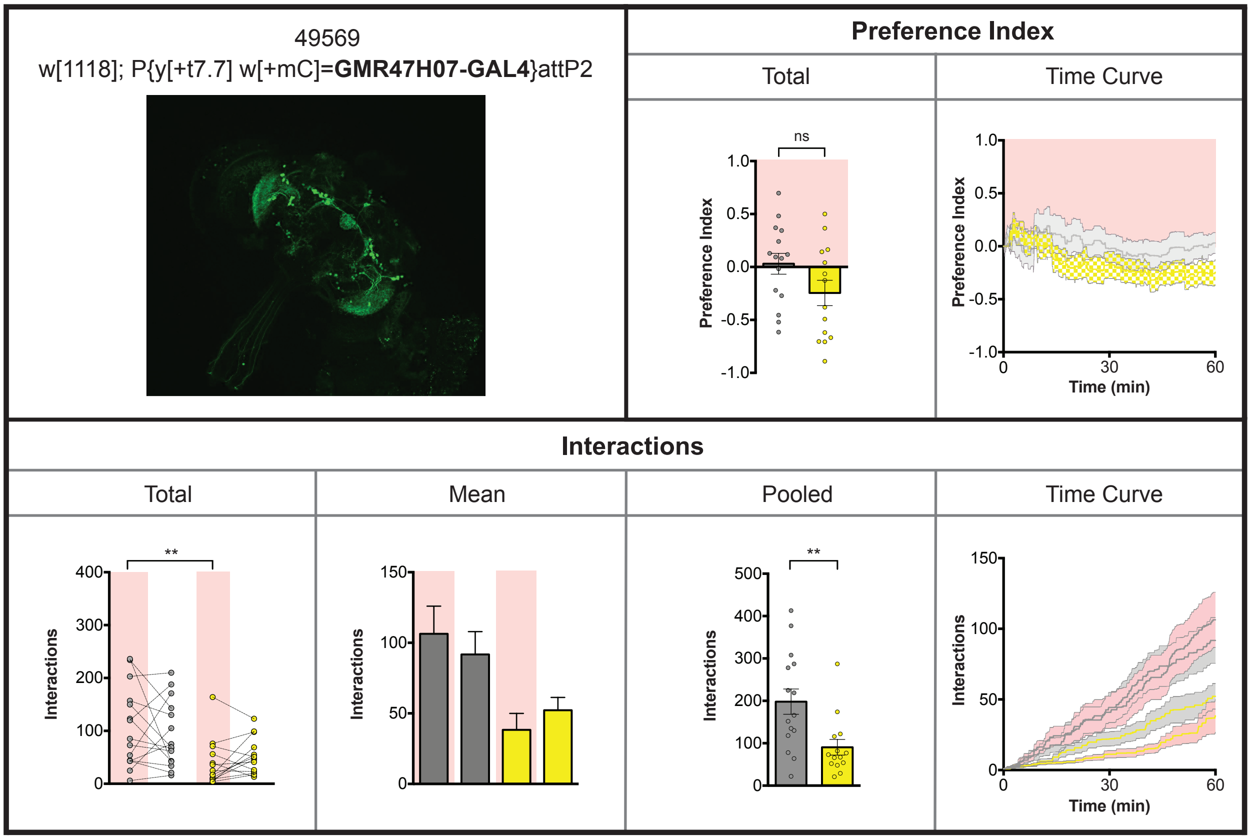
49569
w[1118]; P{y[+t7.7] w[+mC]=GMR47H07-GAL4}attP2
Preference Index
Total
Time Curve
Preference Index
ns
Time (min)
Interactions
Total
Mean
Pooled
Time Curve
Interactions
**
**
Time (min)

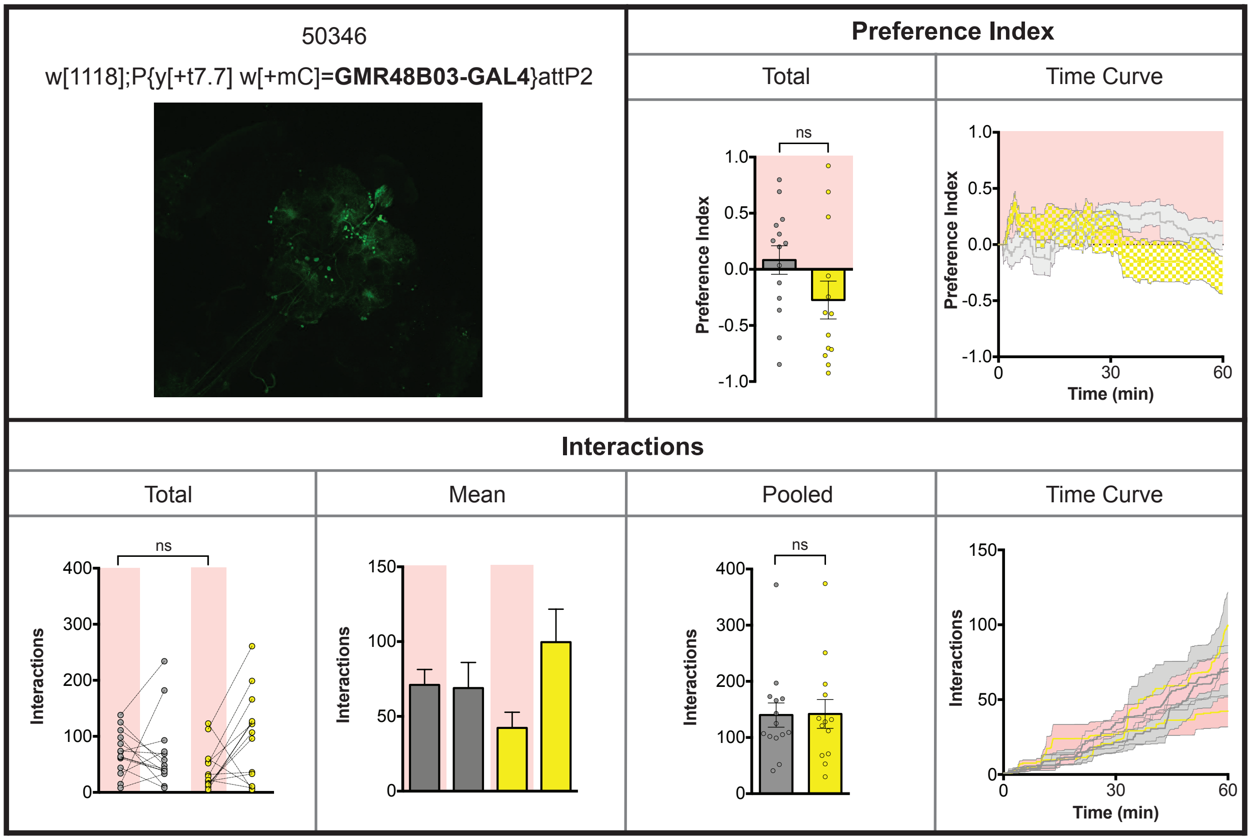
50346
w[1118];P{y[+t7.7] w[+mC]=GMR48B03-GAL4}attP2
Preference Index
Total
Time Curve
ns
Preference Index
Time (min)
Interactions
Total
Mean
Pooled
Time Curve
ns
ns
Interactions
Time (min)

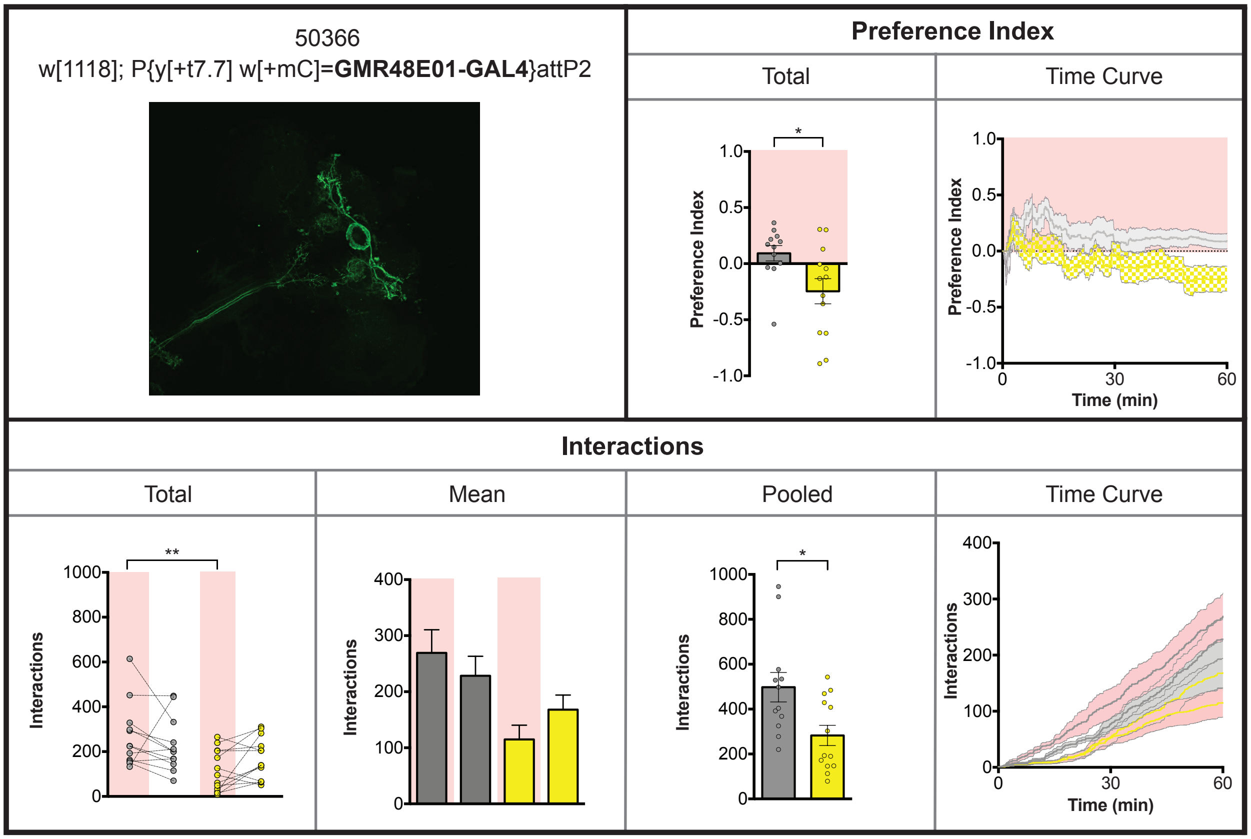
50366
w[1118]; P{y[+t7.7] w[+mC]=GMR48E01-GAL4}attP2
Preference Index
Total
Time Curve
*
Preference Index
Time (min)
Interactions
Total
Mean
Pooled
Time Curve
**
*
Interactions
Time (min)

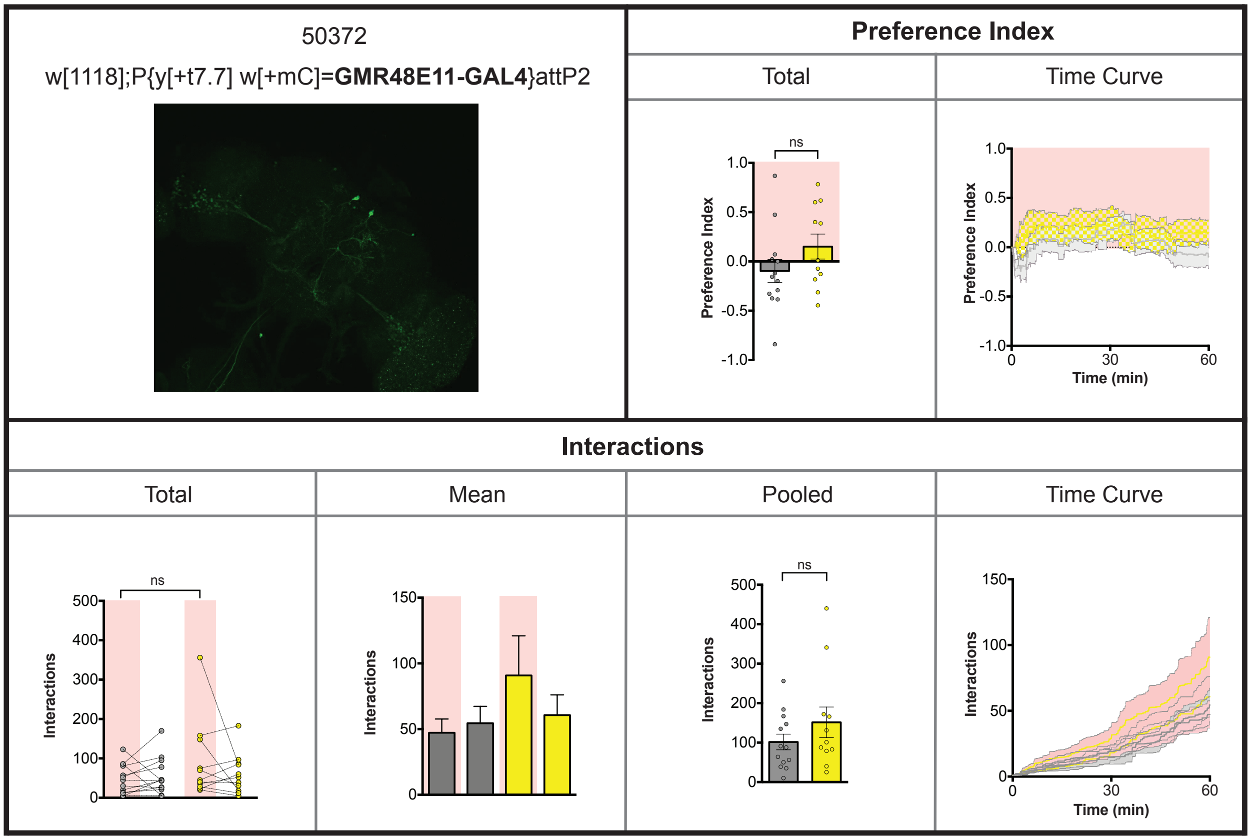
50372
w[1118];P{y[+t7.7] w[+mC]=**GMR48E11-GAL4**}attP2
Preference Index
Total
Time Curve
ns
Preference Index
Time (min)
Interactions
Total
Mean
Pooled
Time Curve
Interactions
Time (min)

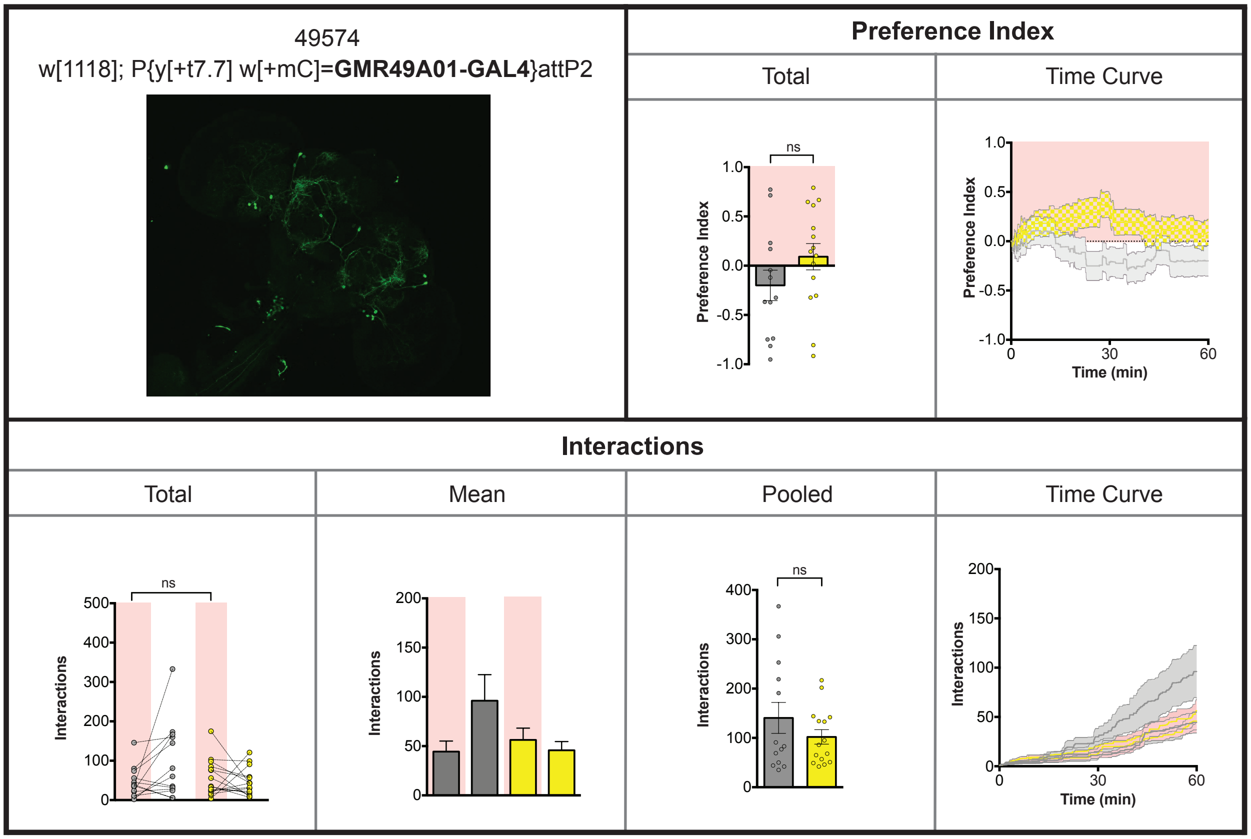
49574
w[1118]; P{y[+t7.7] w[+mC]=**GMR49A01-GAL4**}attP2
Preference Index
Total
Time Curve
ns
Preference Index
Time (min)
Interactions
Total
Mean
Pooled
Time Curve
Interactions
Time (min)

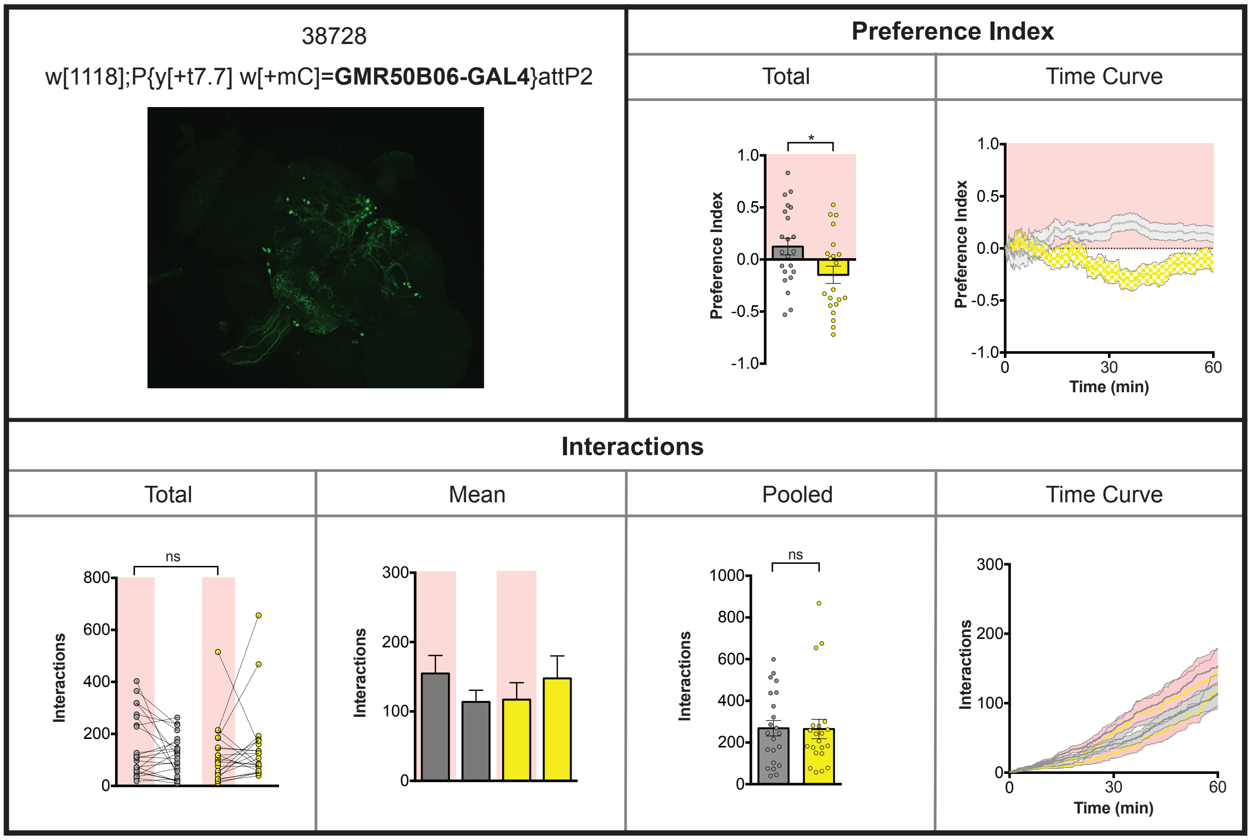
38728
w[1118];P{y[+t7.7] w[+mC]=GMR50B06-GAL4}attP2
Preference Index
Total
Time Curve
*
Preference Index
1.0
0.5
0.0
-0.5
-1.0
0
30
60
Time (min)
Interactions
Total
Mean
Pooled
Time Curve
ns
Interactions
800
600
400
200
0
300
200
100
0
1000
800
600
400
200
0
Time (min)

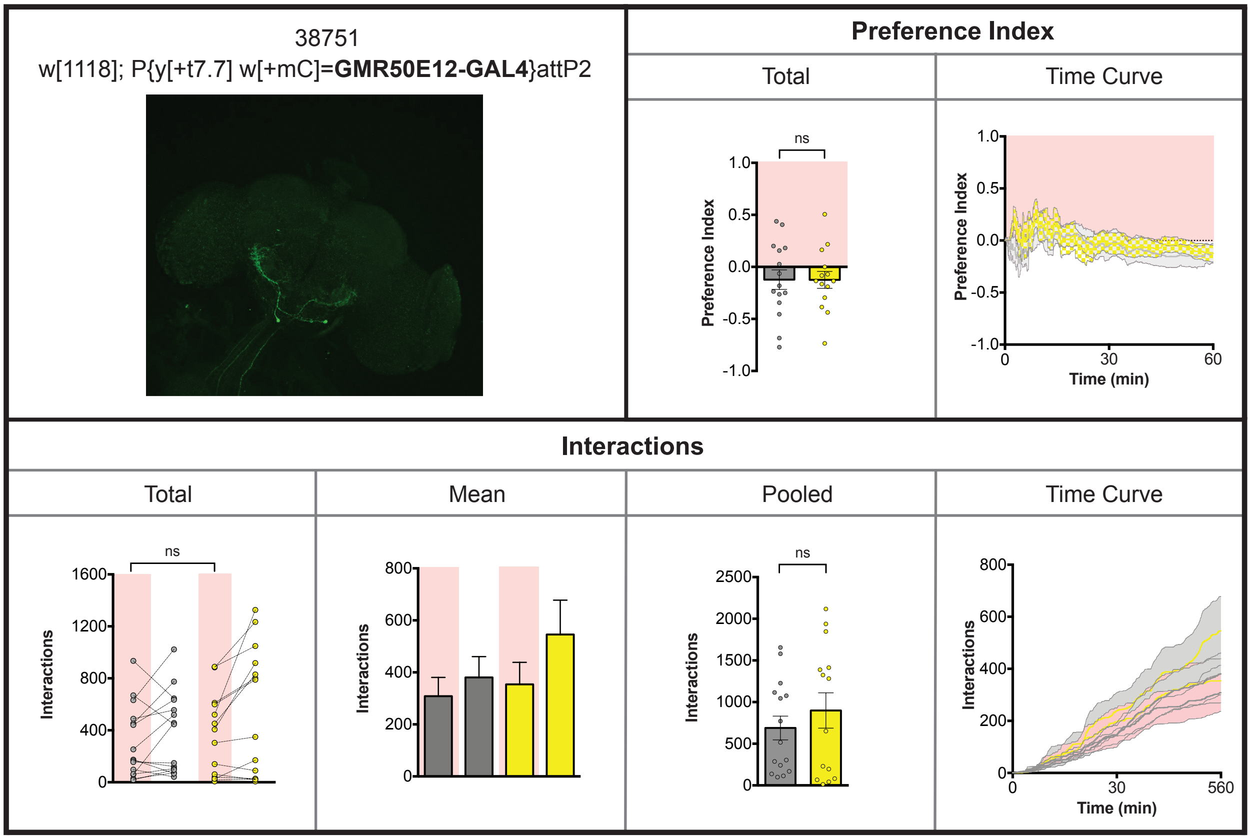
38751
w[1118]; P{y[+t7.7] w[+mC]=GMR50E12-GAL4}attP2
Preference Index
Total
Time Curve
ns
Preference Index
1.0
0.5
0.0
-0.5
-1.0
0
30
60
Time (min)
Interactions
Total
Mean
Pooled
Time Curve
ns
Interactions
1600
1200
800
400
0
600
200
2500
2000
1500
1000
500
0
30
560
Time (min)

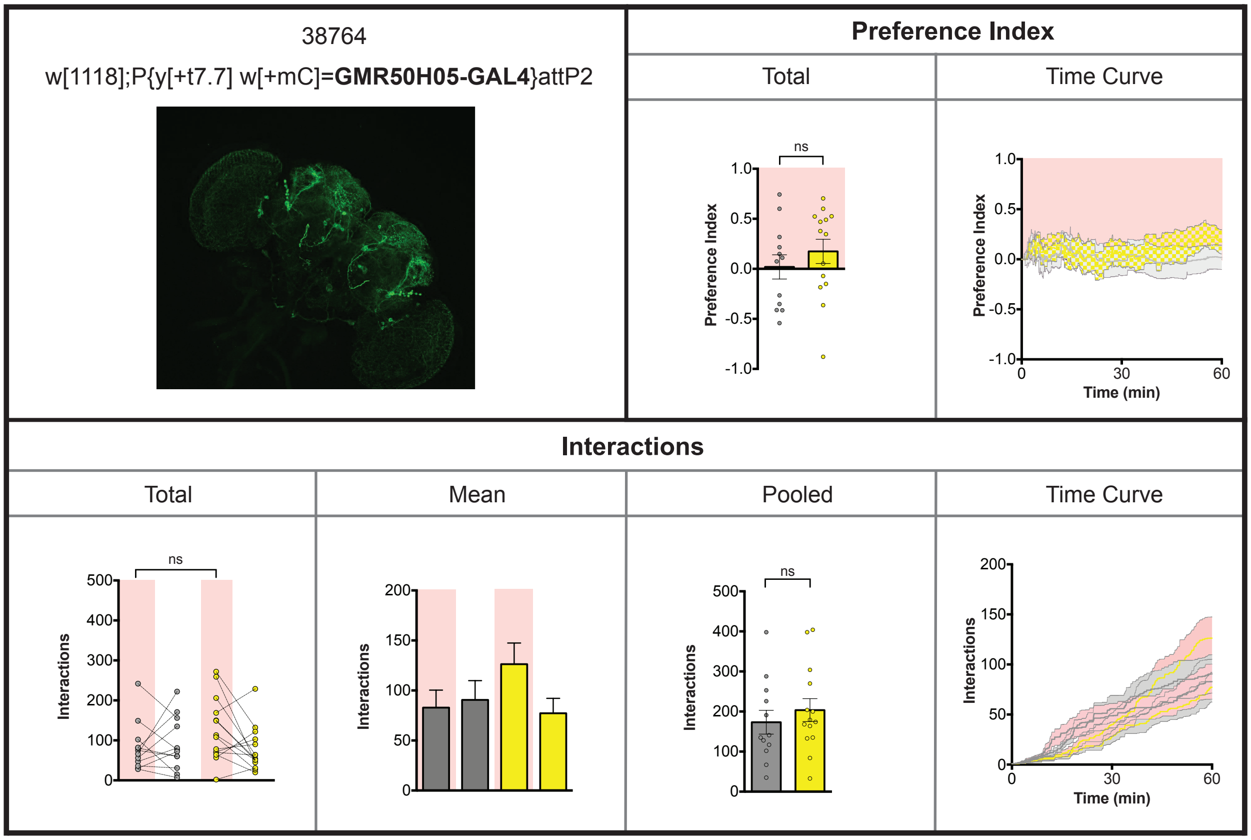
38764
w[1118];P{y[+t7.7] w[+mC]=GMR50H05-GAL4}attP2
Preference Index
Total
Time Curve
ns
Preference Index
1.0
0.5
0.0
-0.5
-1.0
Time (min)
0
30
60
Interactions
Total
Mean
Pooled
Time Curve
ns
Interactions
500
400
300
200
100
0
150
50
Time (min)

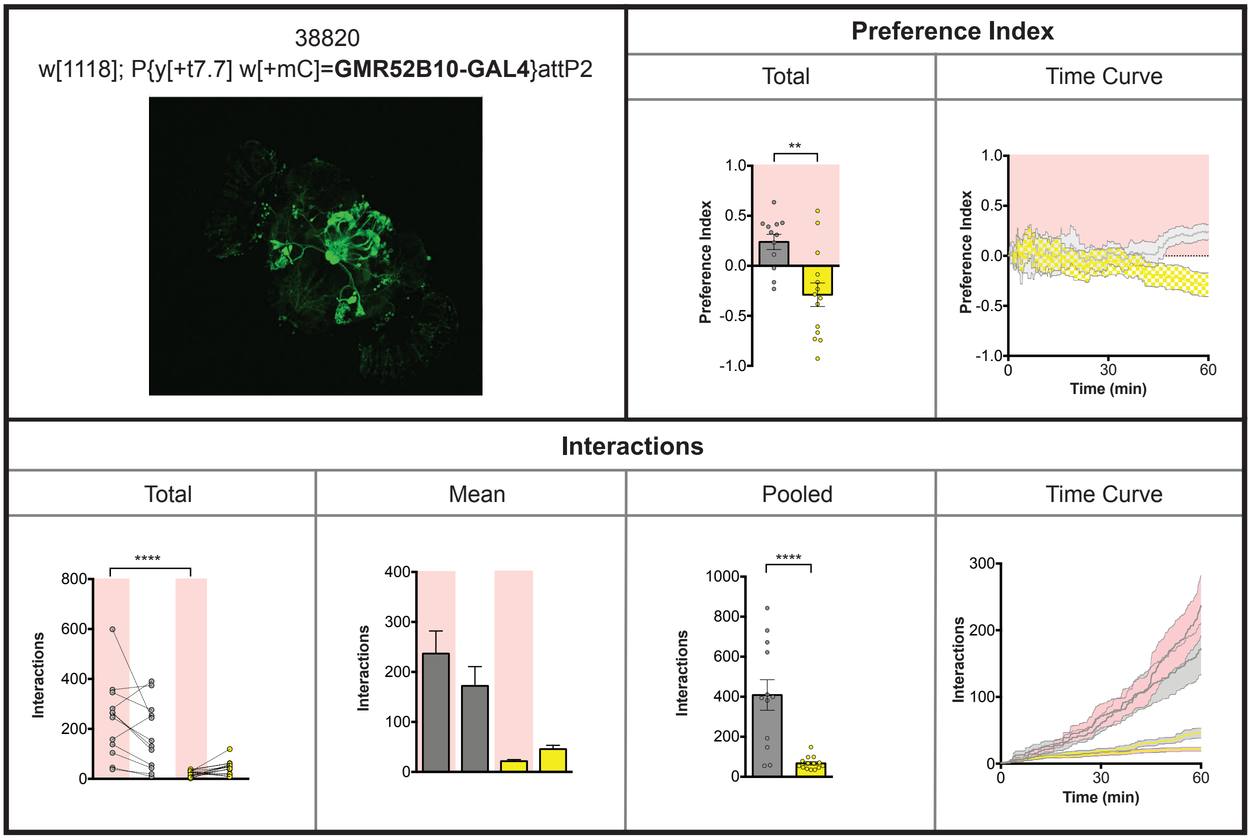
38820
w[1118]; P{y[+t7.7] w[+mC]=GMR52B10-GAL4}attP2
Preference Index
Total
Time Curve
**
Preference Index
1.0
0.5
0.0
-0.5
-1.0
Time (min)
0
30
60
Interactions
Total
Mean
Pooled
Time Curve
****
Interactions
1000
800
600
400
300
200
100
0
Time (min)

38860

w[1118];P{y[+t7.7] w[+mC]=**GMR53A06-GAL4**}attP2

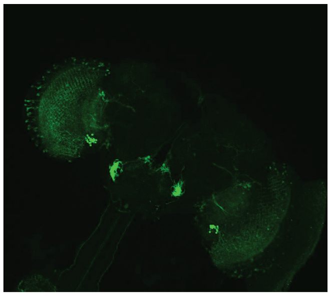

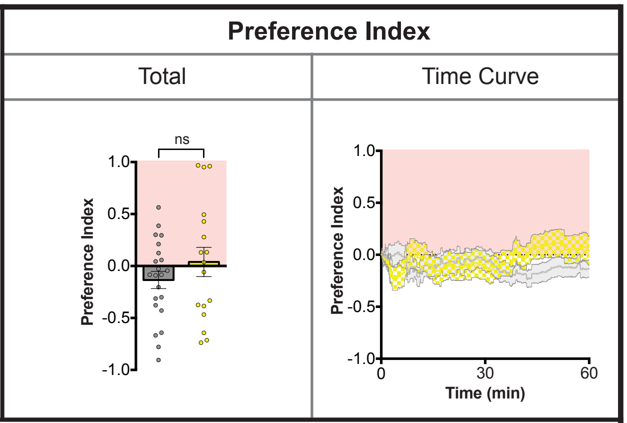


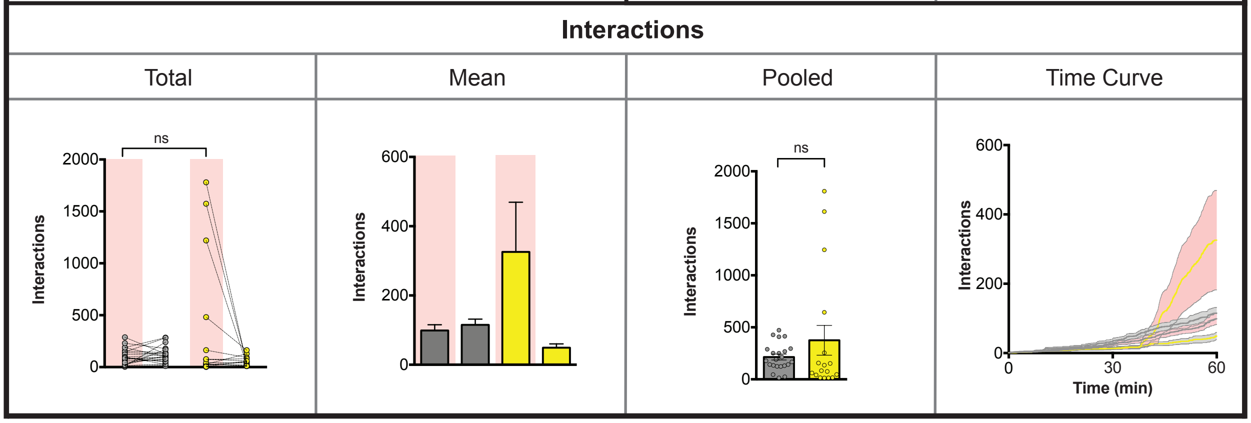


49583

w[1118]; P{y[+t7.7] w[+mC]=**GMR53B03-GAL4**}attP2

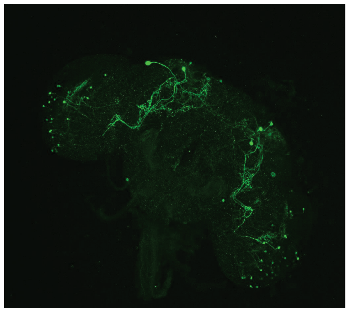

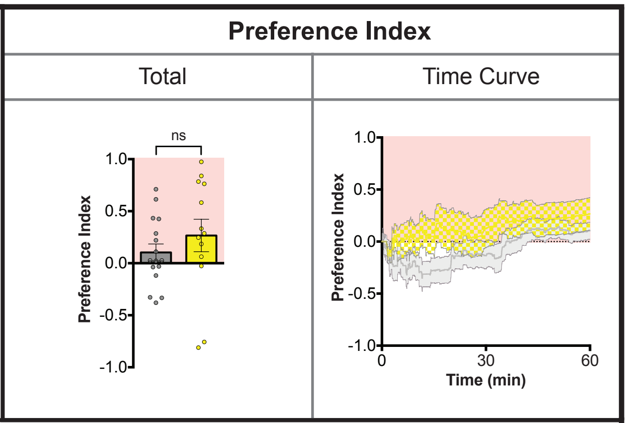


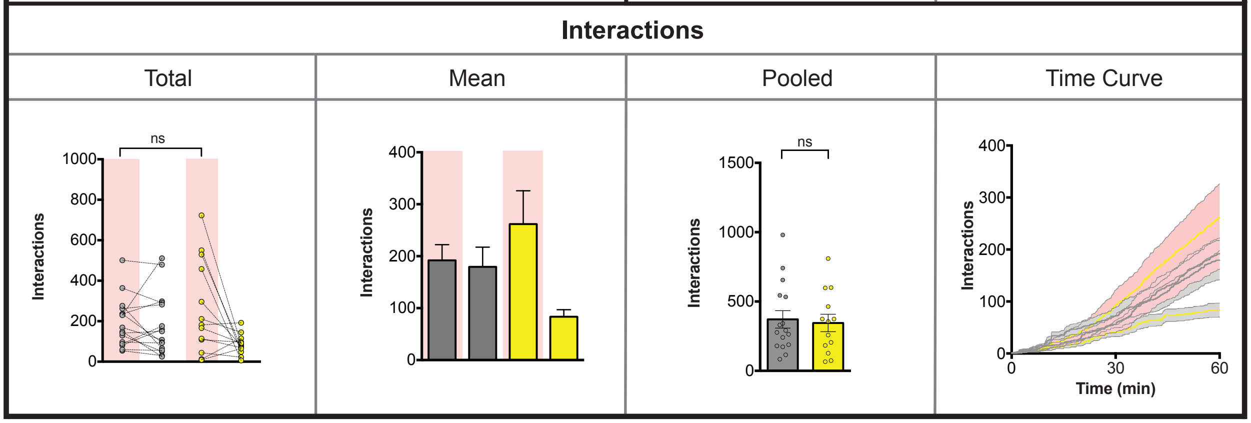

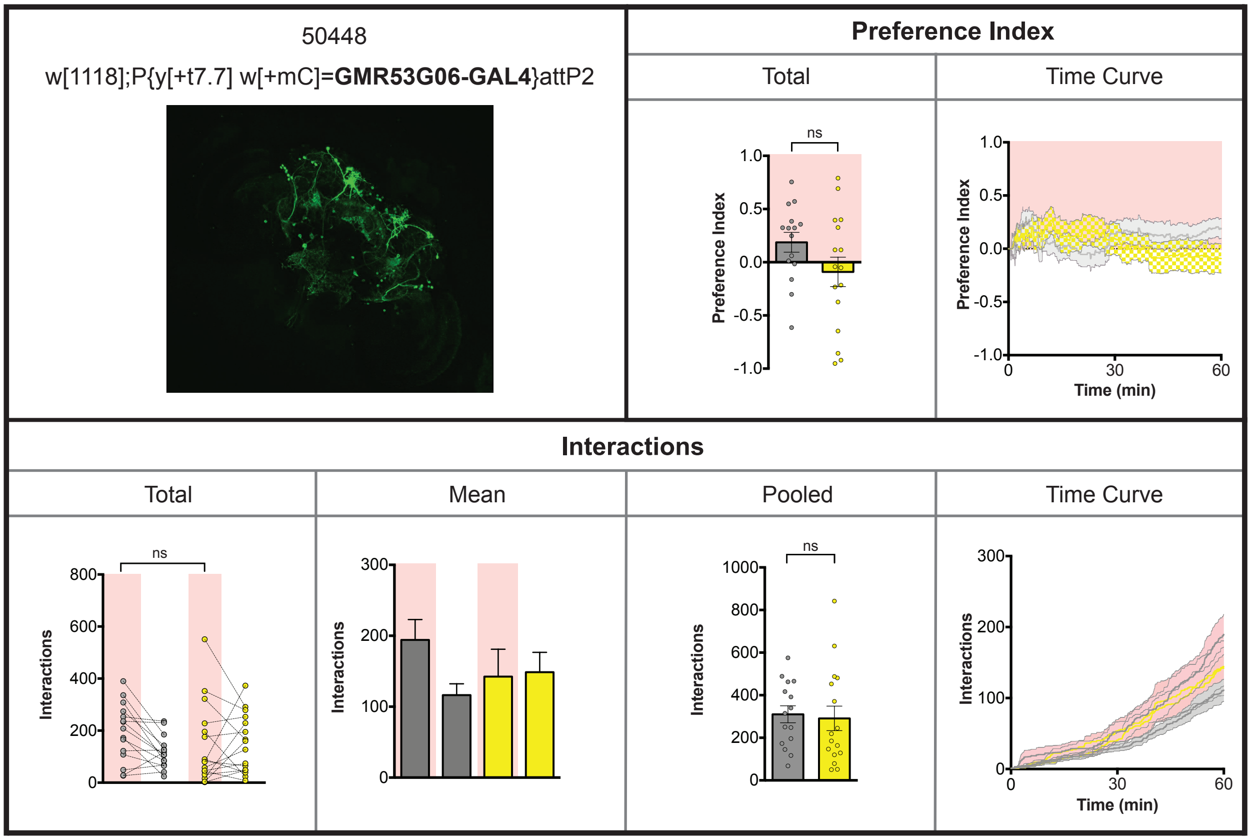
50448
w[1118];P{y[+t7.7] w[+mC]=GMR53G06-GAL4}attP2
Preference Index
Total
Time Curve
ns
Preference Index
Time (min)
Interactions
Total
Mean
Pooled
Time Curve
ns
Interactions
Time (min)

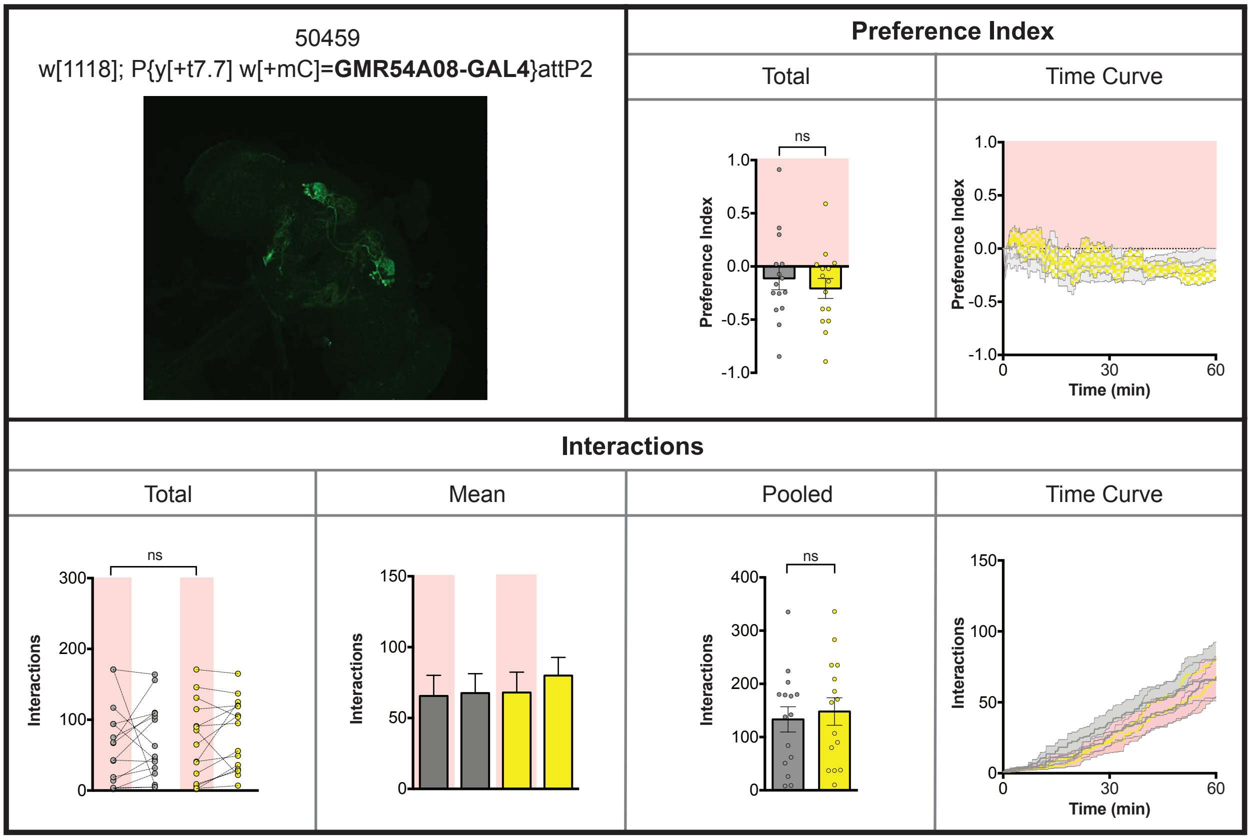
50459
w[1118]; P{y[+t7.7] w[+mC]=GMR54A08-GAL4}attP2
Preference Index
Total
Time Curve
ns
Preference Index
Time (min)
Interactions
Total
Mean
Pooled
Time Curve
ns
Interactions
Time (min)

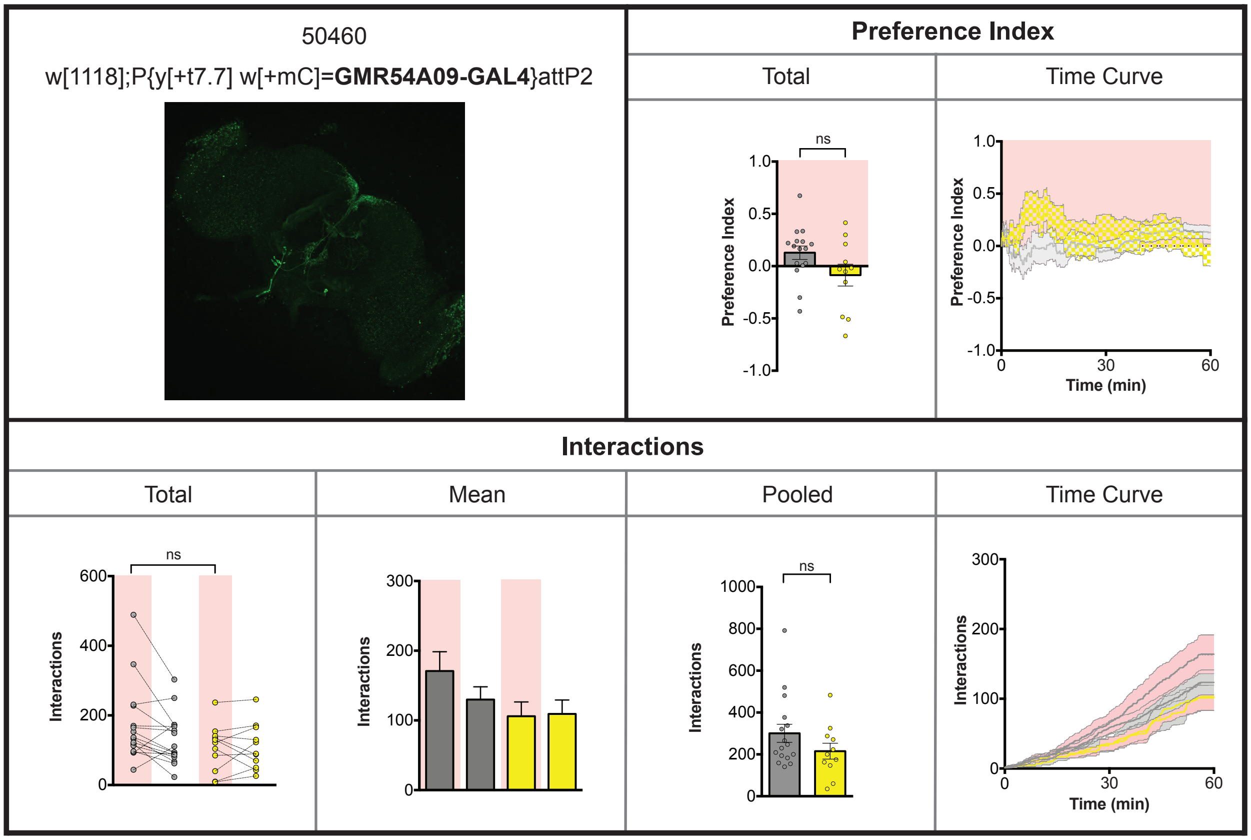
50460
w[1118];P{y[+t7.7] w[+mC]=GMR54A09-GAL4}attP2
Preference Index
Total
Time Curve
ns
Preference Index
Time (min)
Interactions
Total
Mean
Pooled
Time Curve
ns
ns
Interactions
Time (min)

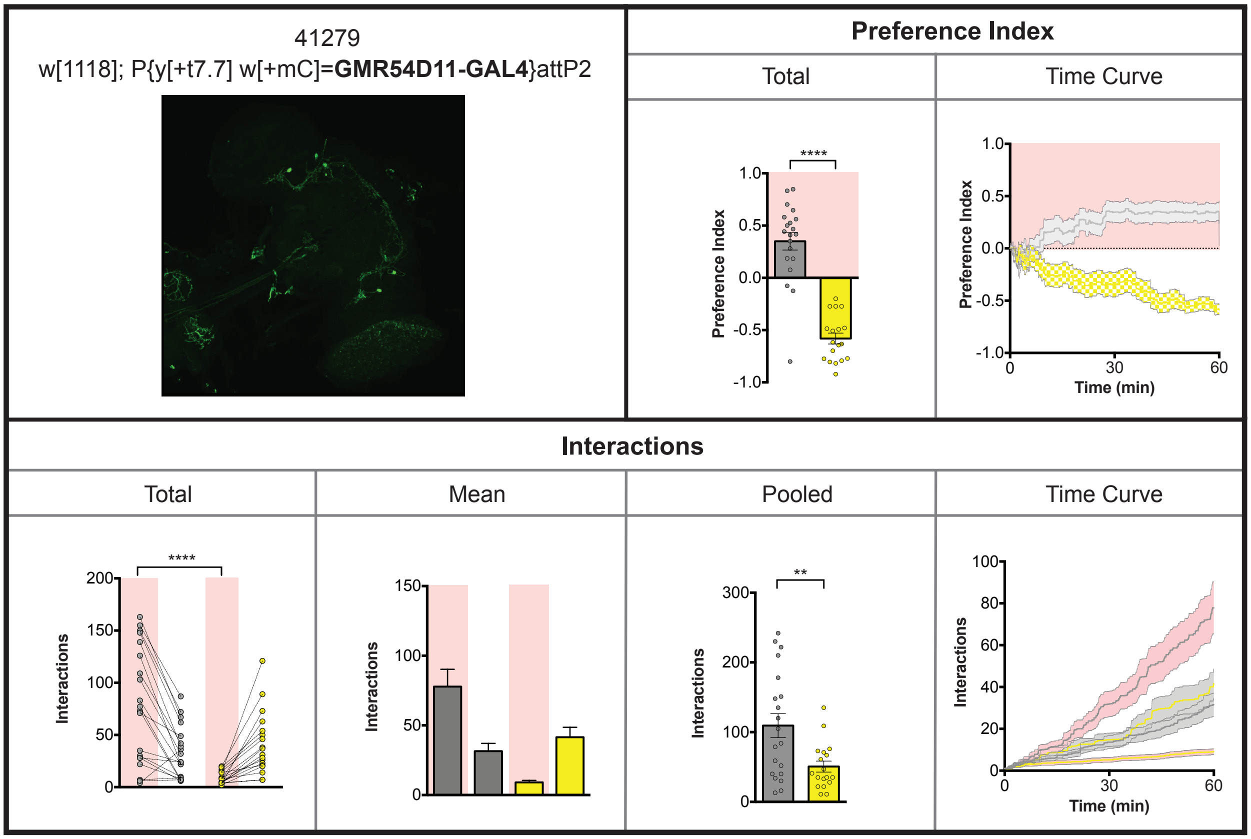
41279
w[1118]; P{y[+t7.7] w[+mC]=GMR54D11-GAL4}attP2
Preference Index
Total
Time Curve
****
Preference Index
Time (min)
Interactions
Total
Mean
Pooled
Time Curve
****
**
Interactions
Time (min)

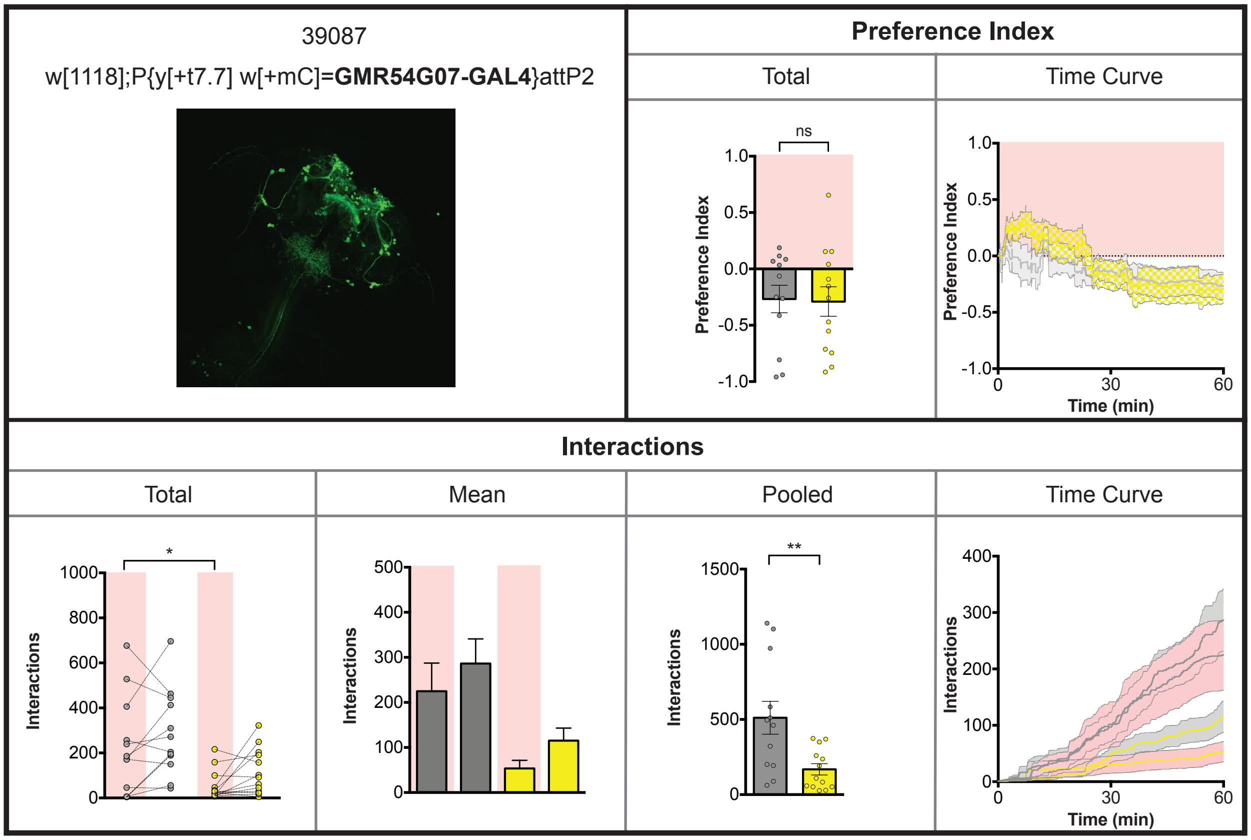
39087
w[1118];P{y[+t7.7] w[+mC]=GMR54G07-GAL4}attP2
Preference Index
Total
Time Curve
ns
1.0
0.5
0.0
-0.5
-1.0
Preference Index
0
30
60
Time (min)
Interactions
Total
Mean
Pooled
Time Curve
*
1000
800
600
400
200
0
500
400
300
200
100
**
1500
1000
500
Interactions

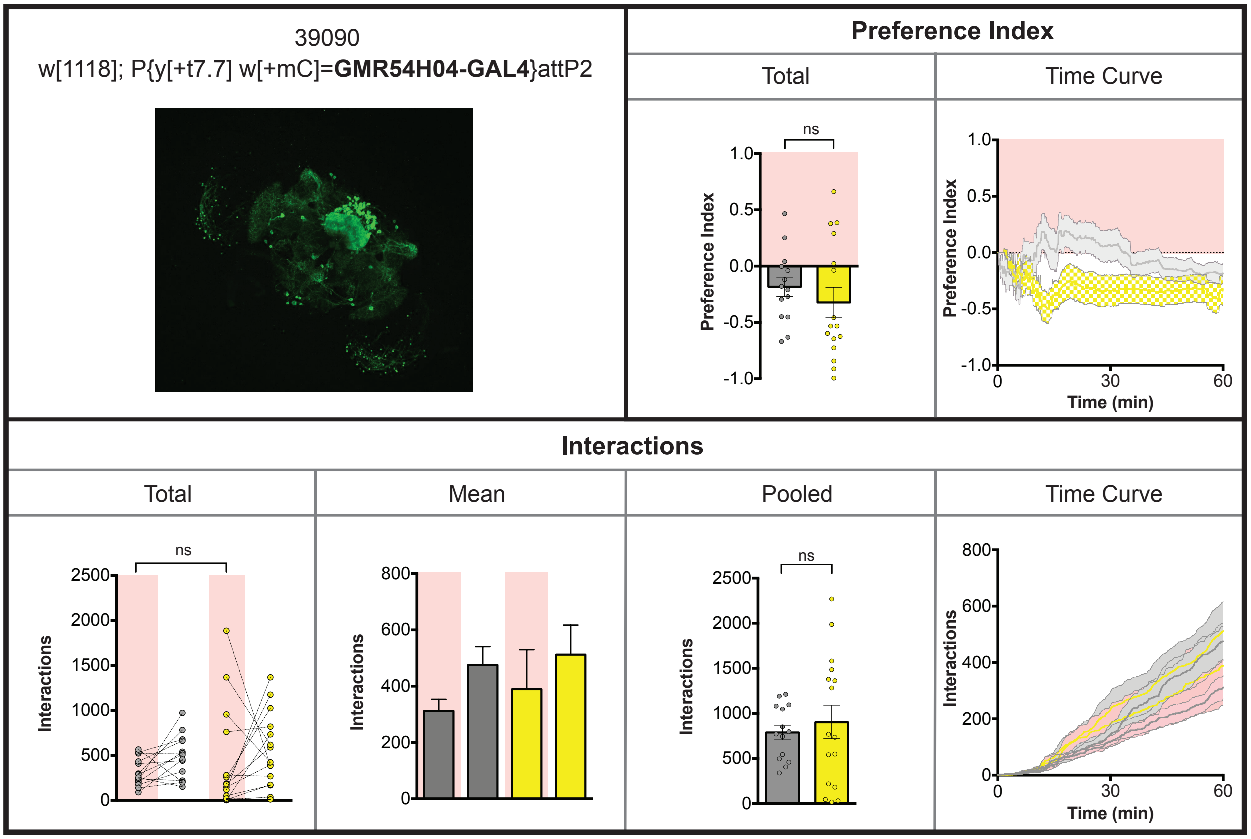
39090
w[1118]; P{y[+t7.7] w[+mC]=GMR54H04-GAL4}attP2
Preference Index
Total
Time Curve
ns
1.0
0.5
0.0
-0.5
-1.0
Preference Index
0
30
60
Time (min)
Interactions
Total
Mean
Pooled
Time Curve
ns
2500
2000
1500
1000
500
0
800
600
400
200
Interactions

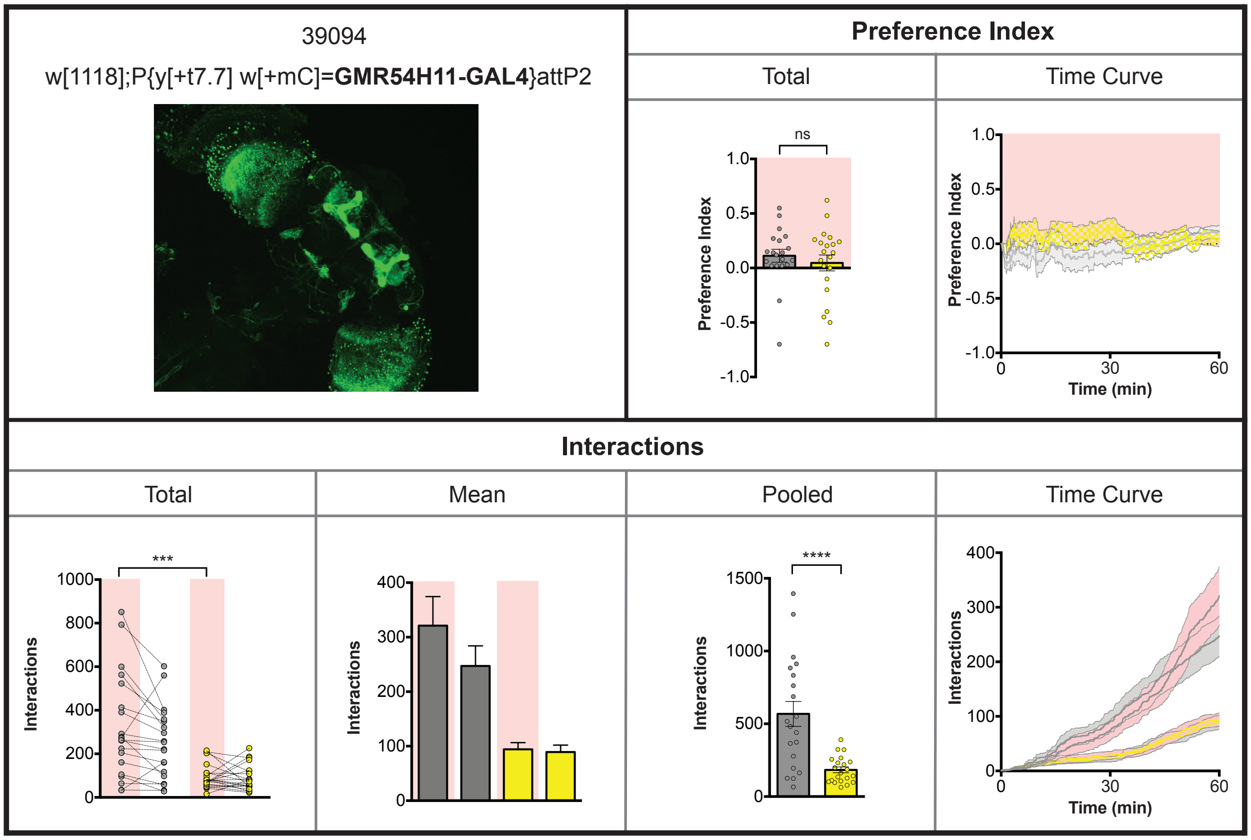
39094
w[1118];P{y[+t7.7] w[+mC]=**GMR54H11-GAL4**}attP2
Preference Index
Total
Time Curve
ns
Preference Index
1.0
0.5
0.0
-0.5
-1.0
0
30
60
Time (min)
Interactions
Total
Mean
Pooled
Time Curve
***
****
Interactions
1000
800
600
400
200
0
400
300
200
100
1500
500
Time (min)

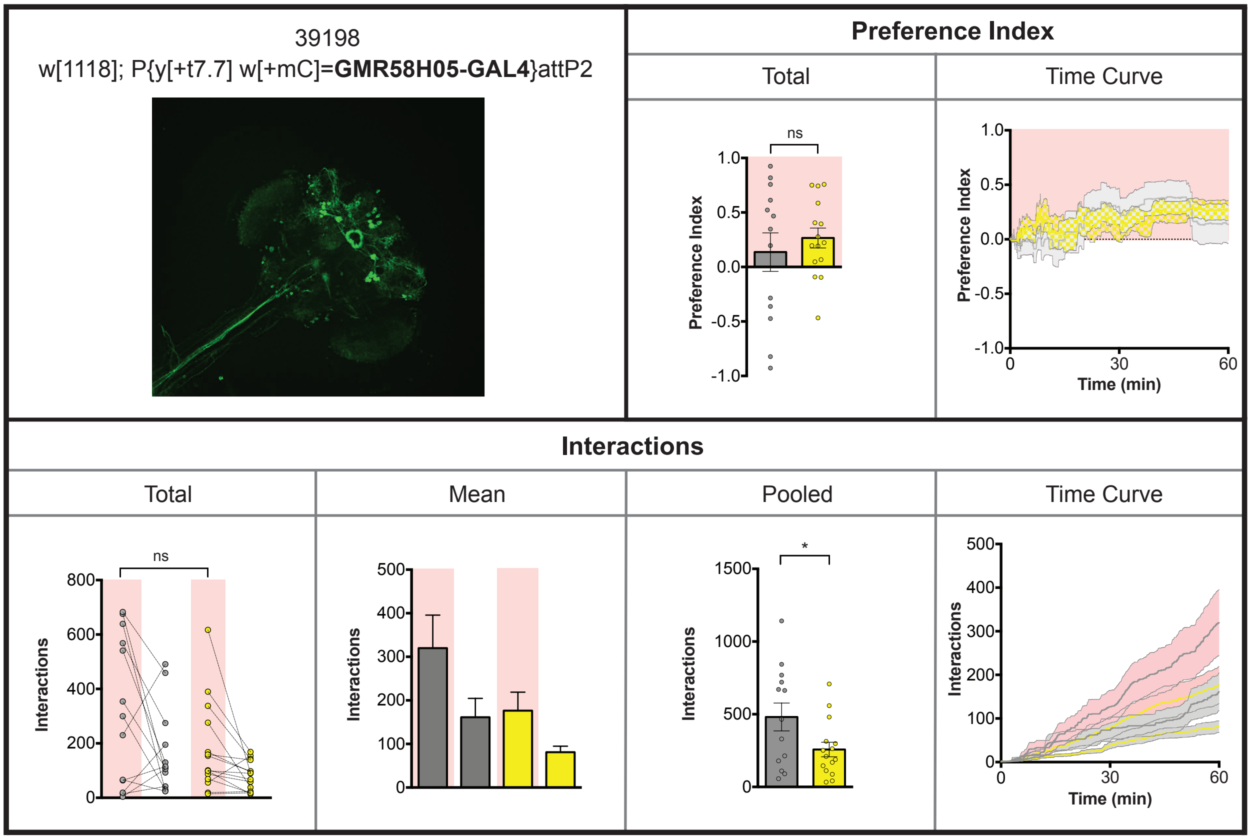
39198
w[1118]; P{y[+t7.7] w[+mC]=**GMR58H05-GAL4**}attP2
Preference Index
Total
Time Curve
ns
Preference Index
1.0
0.5
0.0
-0.5
-1.0
0
30
60
Time (min)
Interactions
Total
Mean
Pooled
Time Curve
ns
*
Interactions
800
600
400
200
0
500
400
300
200
100
1500
1000
Time (min)

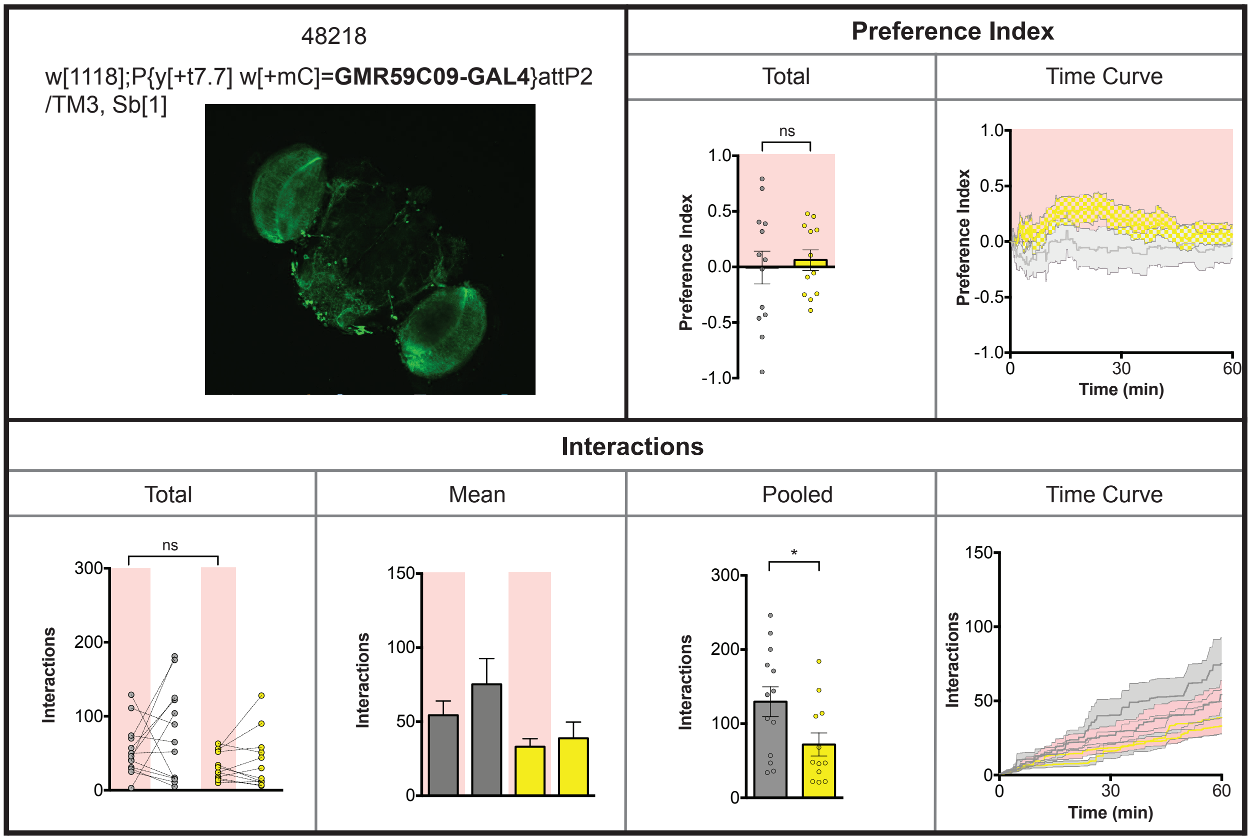
48218
w[1118];P{y[+t7.7] w[+mC]=**GMR59C09-GAL4**}attP2
/TM3, Sb[1]
Preference Index
Total
Time Curve
ns
Preference Index
Time (min)
Interactions
Total
Mean
Pooled
Time Curve
ns
*
Interactions
Time (min)

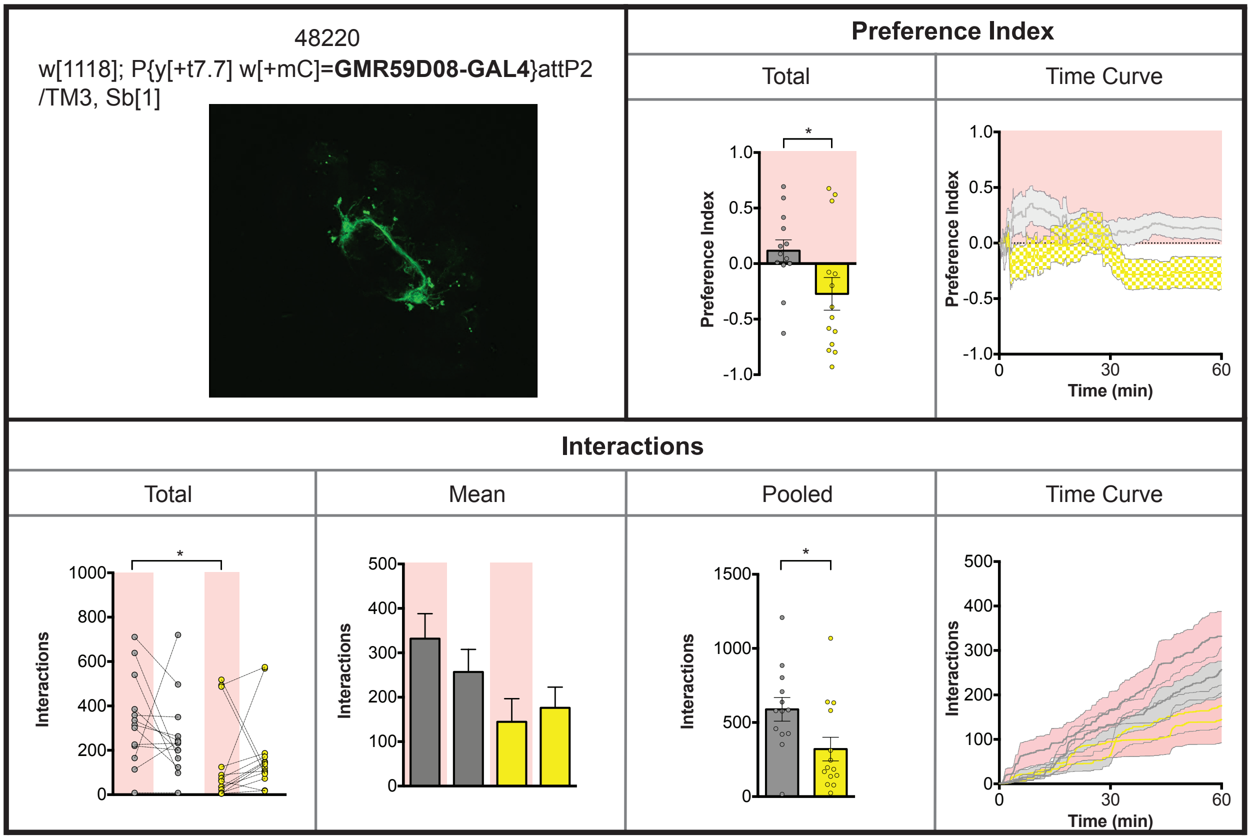
48220
w[1118]; P{y[+t7.7] w[+mC]=**GMR59D08-GAL4**}attP2
/TM3, Sb[1]
Preference Index
Total
Time Curve
*
Preference Index
Time (min)
Interactions
Total
Mean
Pooled
Time Curve
*
*
Interactions
Time (min)

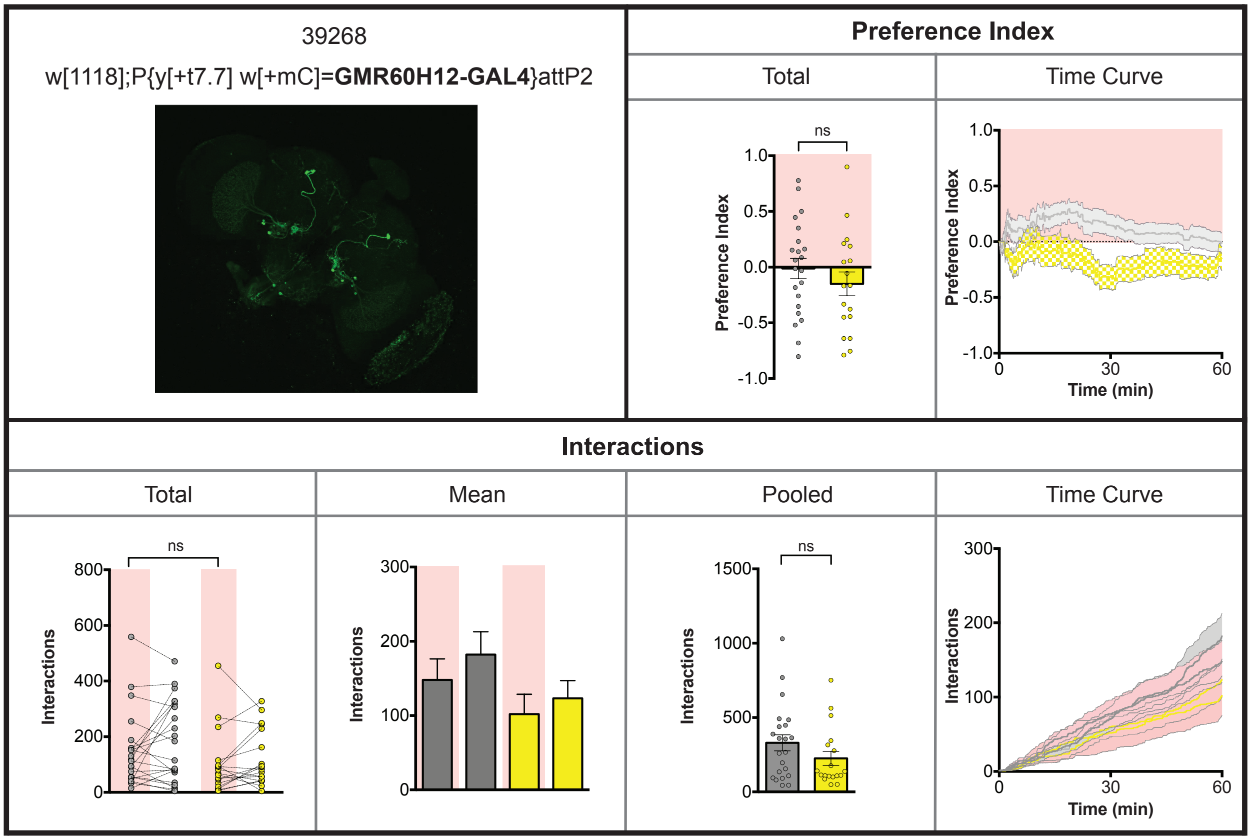
39268
w[1118];P{y[+t7.7] w[+mC]=GMR60H12-GAL4}attP2
Preference Index
Total
Time Curve
ns
1.0
0.5
0.0
-0.5
-1.0
Preference Index
0
30
60
Time (min)
Interactions
Total
Mean
Pooled
Time Curve
ns
800
600
400
200
0
Interactions
300
200
100
0
ns
1500
1000
500
0
Time (min)

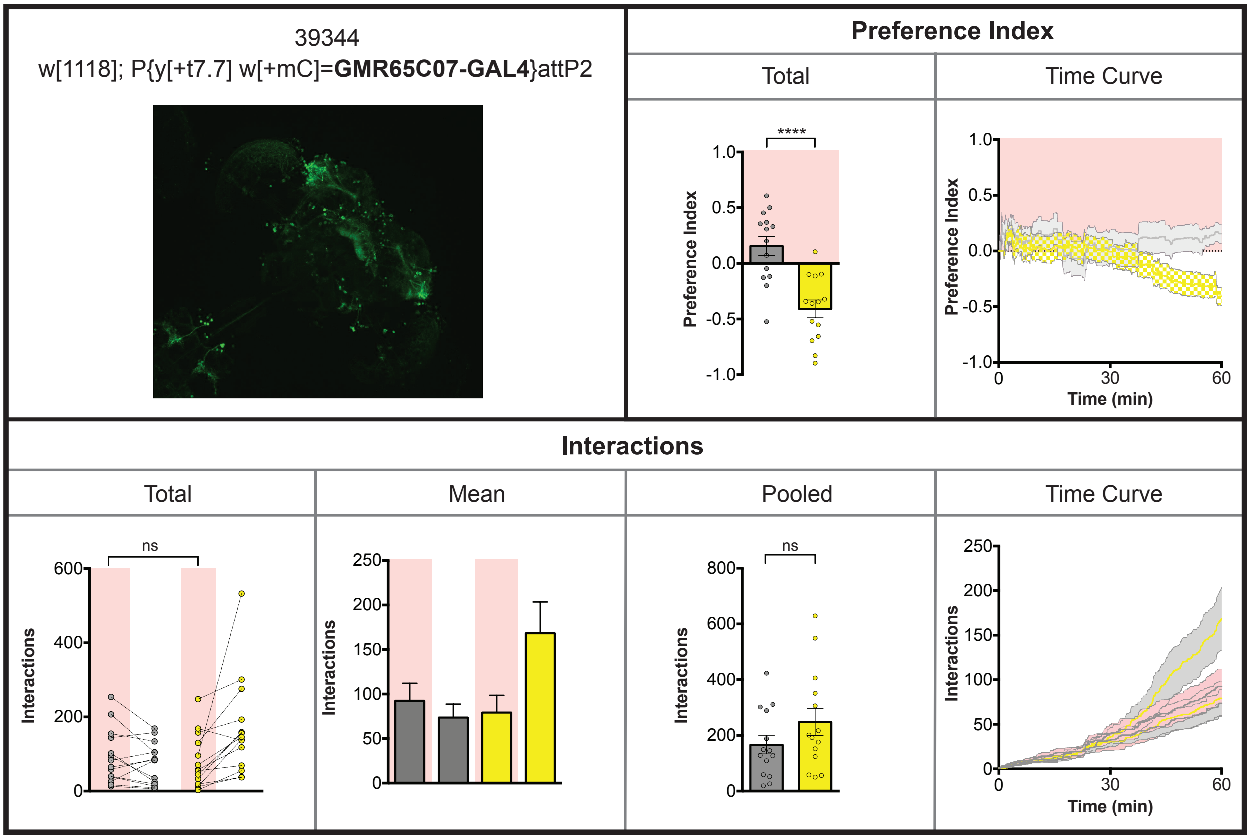
39344
w[1118]; P{y[+t7.7] w[+mC]=GMR65C07-GAL4}attP2
Preference Index
Total
Time Curve
****
1.0
0.5
0.0
-0.5
-1.0
Preference Index
0
30
60
Time (min)
Interactions
Total
Mean
Pooled
Time Curve
ns
600
400
200
0
Interactions
250
200
150
100
50
0
ns
800
600
400
200
Time (min)

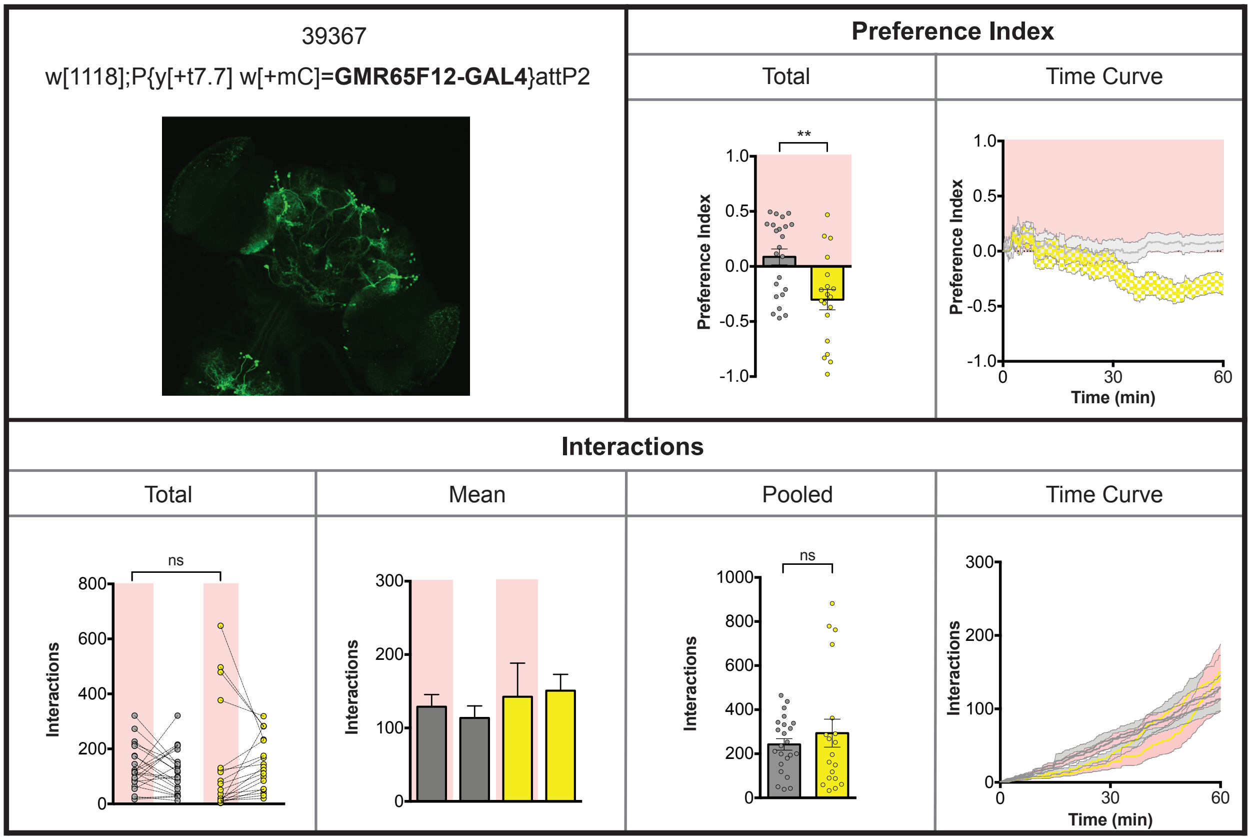
39367
w[1118];P{y[+t7.7] w[+mC]=GMR65F12-GAL4}attP2
Preference Index
Total
Time Curve
**
Preference Index
1.0
0.5
0.0
-0.5
-1.0
0
30
60
Time (min)
Interactions
Total
Mean
Pooled
Time Curve
ns
ns
800
600
400
200
0
300
200
100
1000
Time (min)

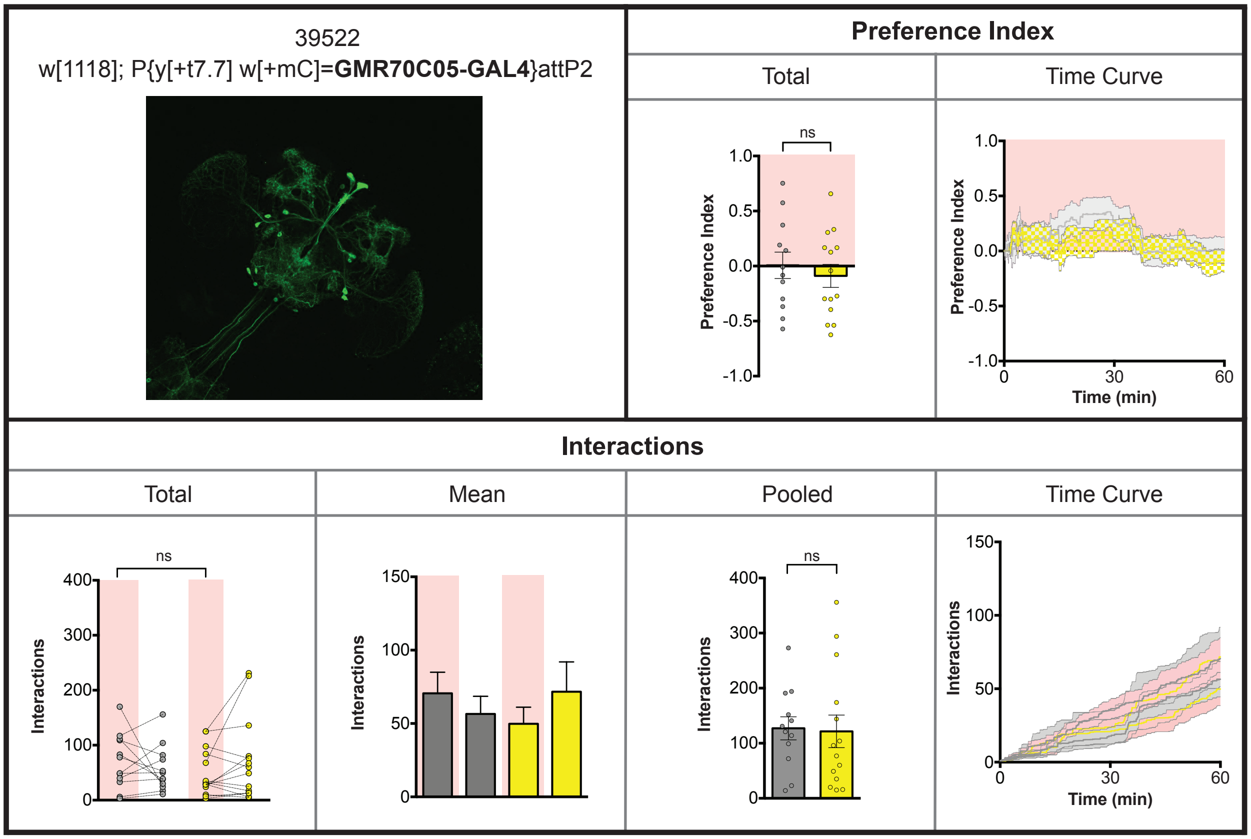
39522
w[1118]; P{y[+t7.7] w[+mC]=GMR70C05-GAL4}attP2
Preference Index
Total
Time Curve
ns
Preference Index
1.0
0.5
0.0
-0.5
-1.0
0
30
60
Time (min)
Interactions
Total
Mean
Pooled
Time Curve
ns
ns
400
300
200
100
0
150
50
Time (min)

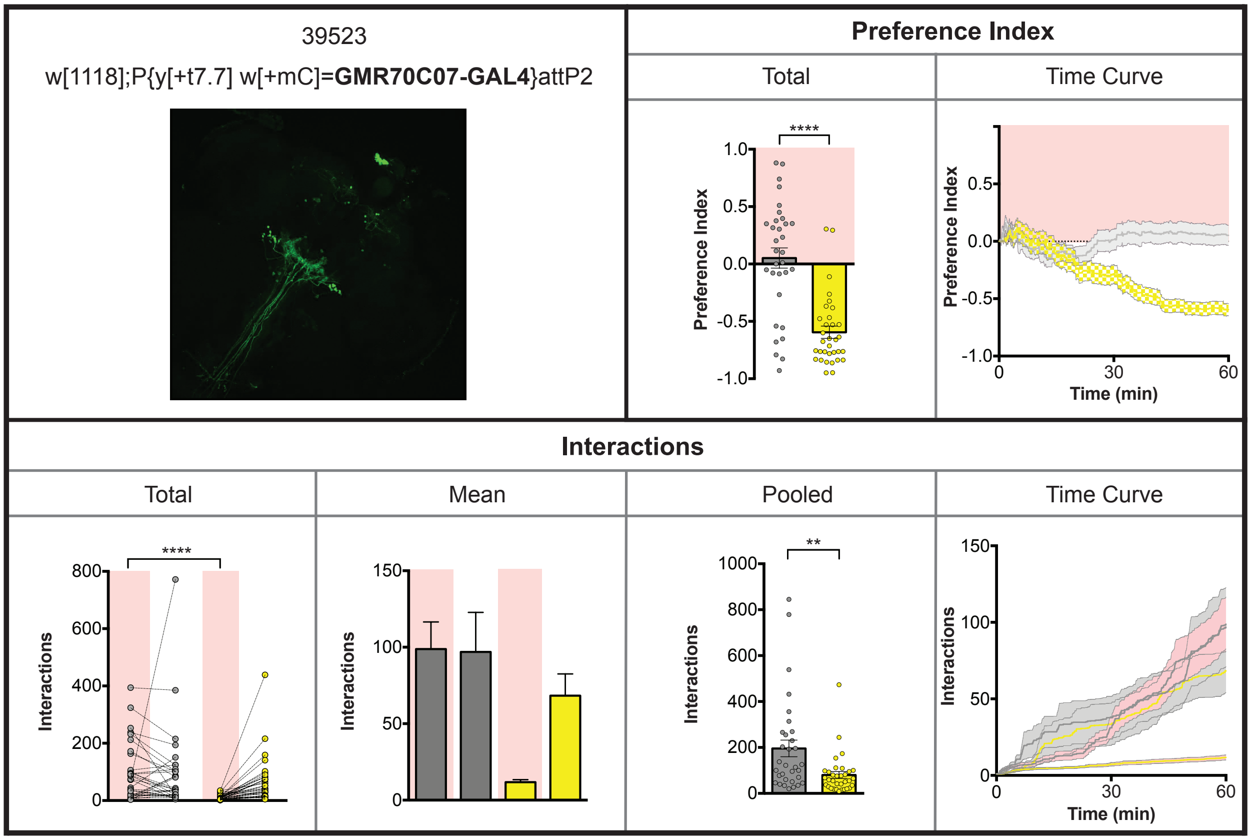
39523
w[1118];P{y[+t7.7] w[+mC]=GMR70C07-GAL4}attP2
Preference Index
Total
Time Curve
Preference Index
****
Time (min)
Interactions
Total
Mean
Pooled
Time Curve
Interactions
****
**
Time (min)

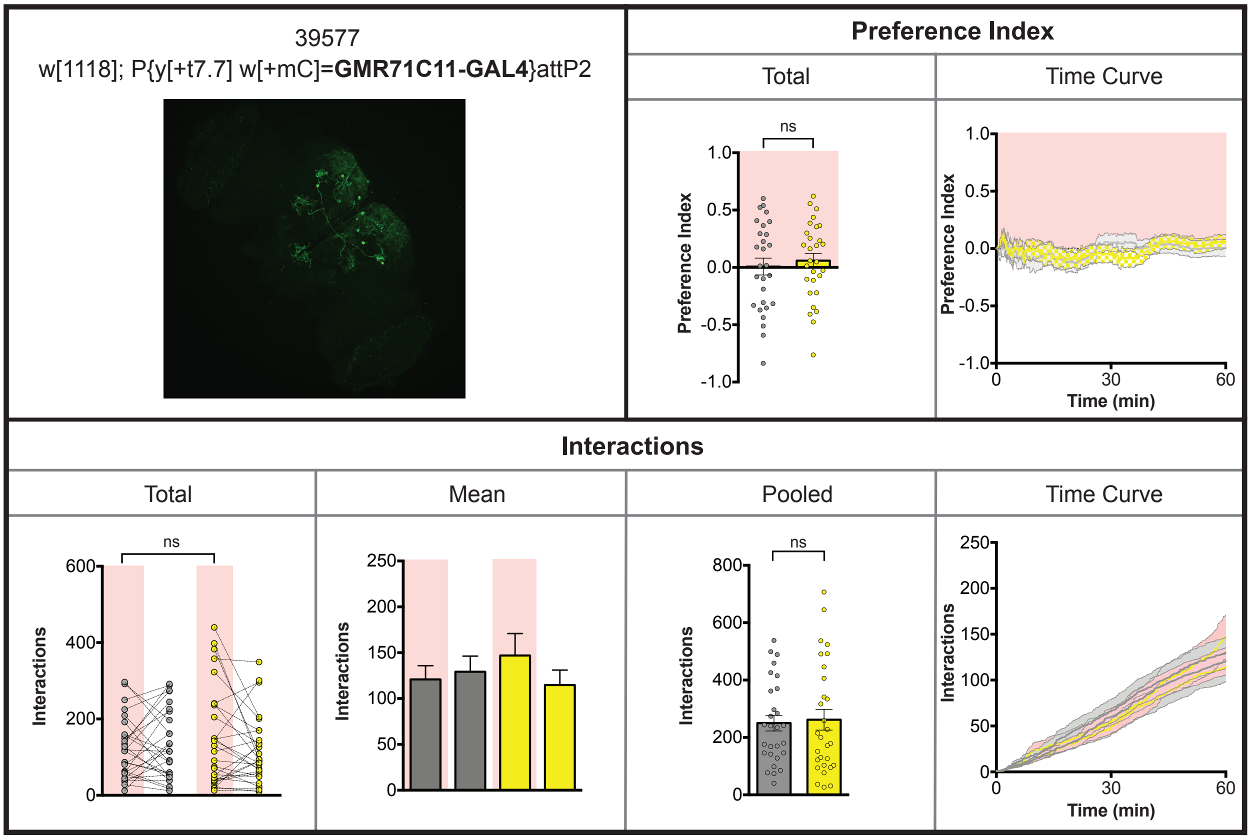
39577
w[1118]; P{y[+t7.7] w[+mC]=GMR71C11-GAL4}attP2
Preference Index
Total
Time Curve
Preference Index
ns
Time (min)
Interactions
Total
Mean
Pooled
Time Curve
Interactions
ns
ns
Time (min)

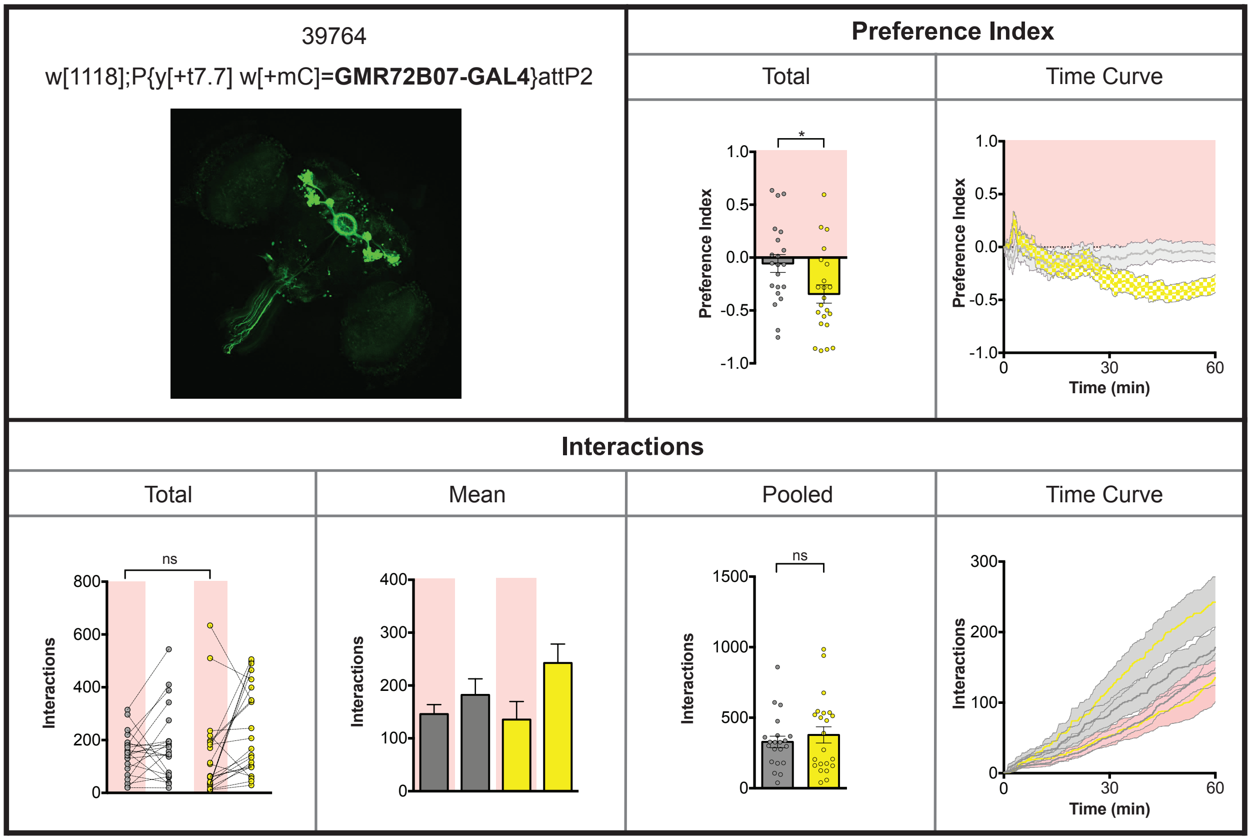
39764
w[1118];P{y[+t7.7] w[+mC]=**GMR72B07-GAL4**}attP2
Preference Index
Total
Time Curve
Preference Index
Time (min)
Interactions
Total
Mean
Pooled
Time Curve
ns
Interactions
Time (min)

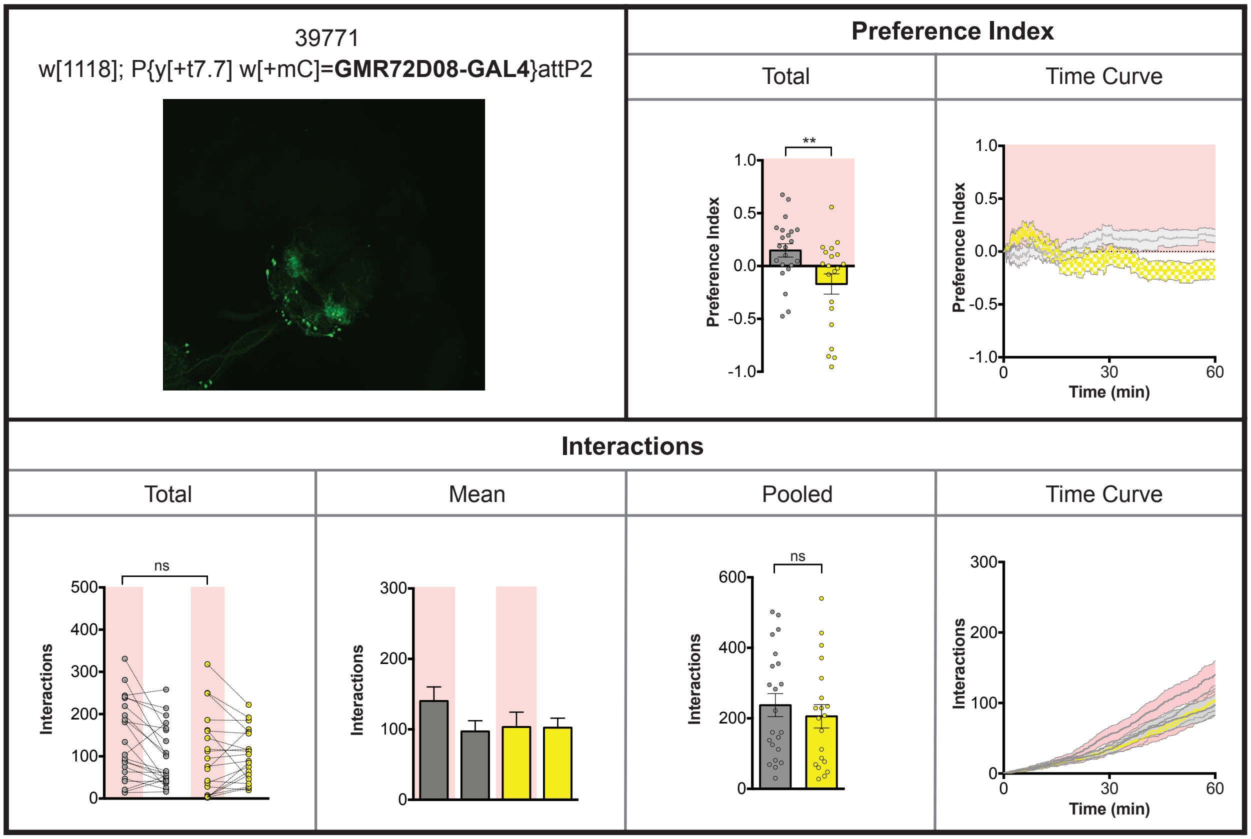
39771
w[1118]; P{y[+t7.7] w[+mC]=**GMR72D08-GAL4**}attP2
Preference Index
Total
Time Curve
Preference Index
Time (min)
Interactions
Total
Mean
Pooled
Time Curve
ns
Interactions
Time (min)

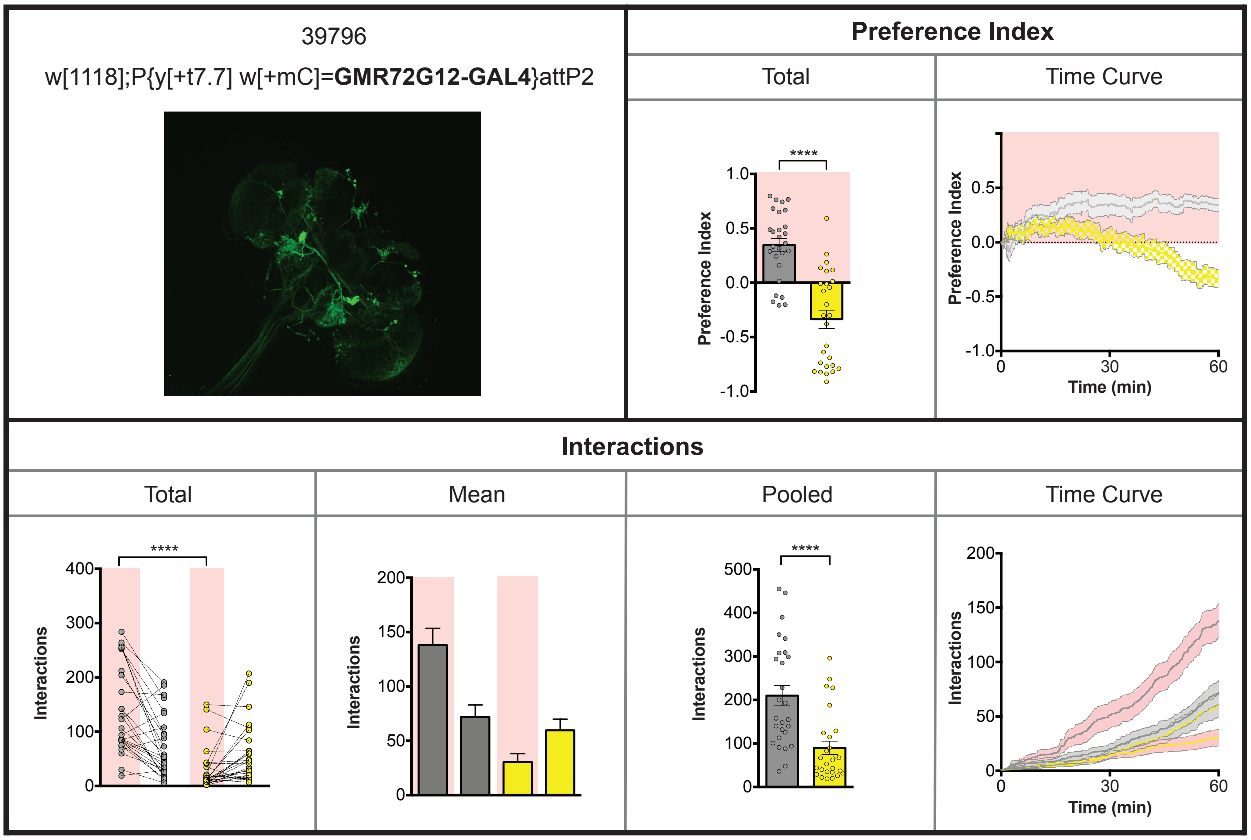
39796
w[1118];P{y[+t7.7] w[+mC]=GMR72G12-GAL4}attP2
Preference Index
Total
Time Curve
Preference Index
Time (min)
Interactions
Total
Mean
Pooled
Time Curve
Interactions
Time (min)
****

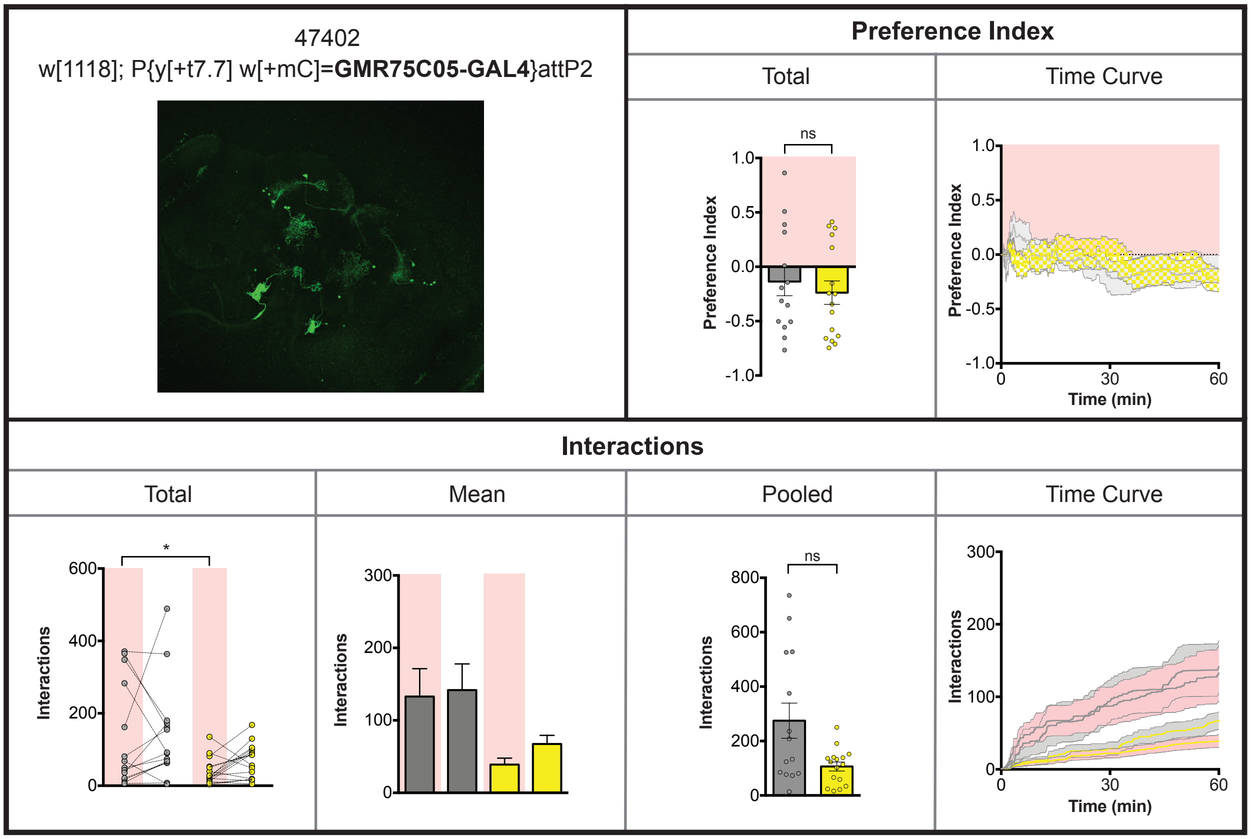
47402
w[1118]; P{y[+t7.7] w[+mC]=GMR75C05-GAL4}attP2
Preference Index
Total
Time Curve
Preference Index
Time (min)
ns
Interactions
Total
Mean
Pooled
Time Curve
Interactions
Time (min)
*

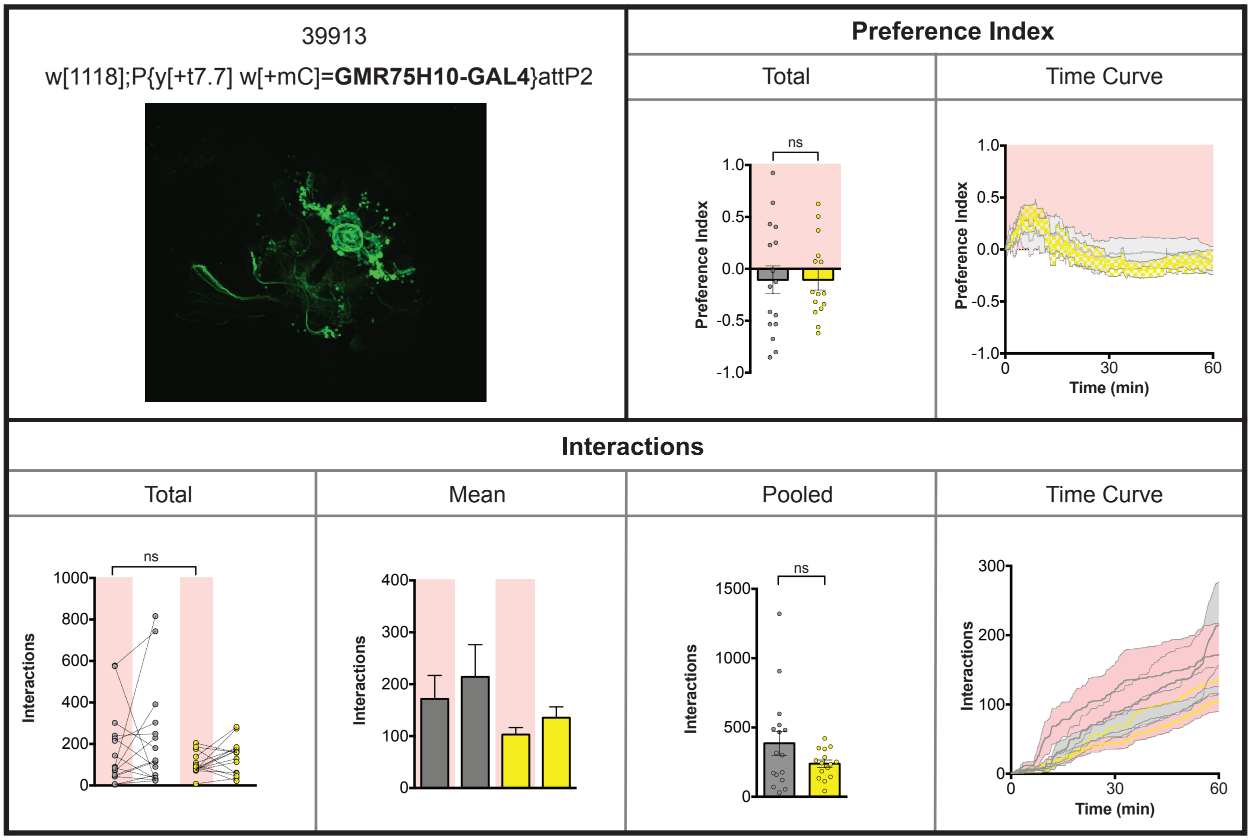
39913
w[1118];P{y[+t7.7] w[+mC]=GMR75H10-GAL4}attP2
Preference Index
Total
Time Curve
ns
Preference Index
1.0
0.5
0.0
-0.5
-1.0
0
30
60
Time (min)
Interactions
Total
Mean
Pooled
Time Curve
ns
Interactions
1000
800
600
400
200
0
400
300
200
100
1500
1000
500
300
Time (min)

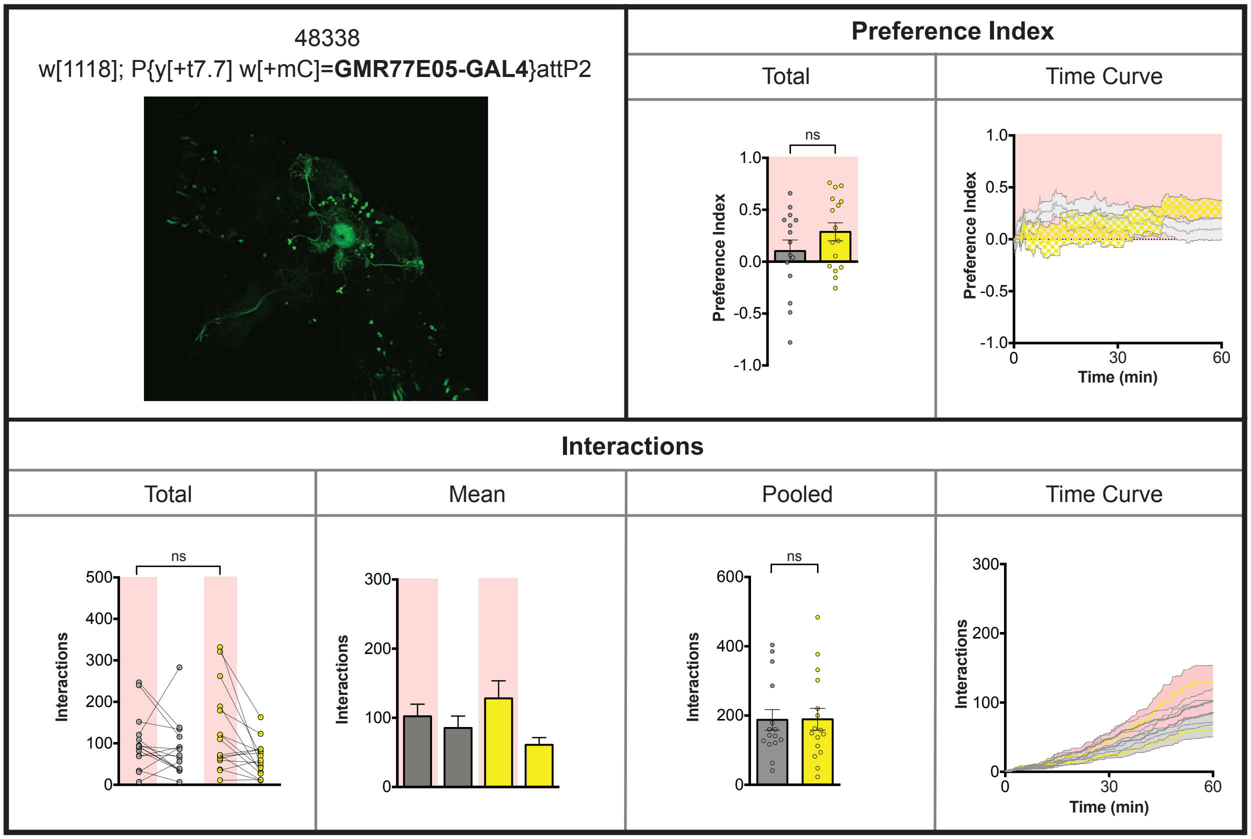
48338
w[1118]; P{y[+t7.7] w[+mC]=GMR77E05-GAL4}attP2
Preference Index
Total
Time Curve
ns
Preference Index
1.0
0.5
0.0
-0.5
-1.0
0
30
60
Time (min)
Interactions
Total
Mean
Pooled
Time Curve
ns
Interactions
500
400
300
200
100
0
600
Time (min)

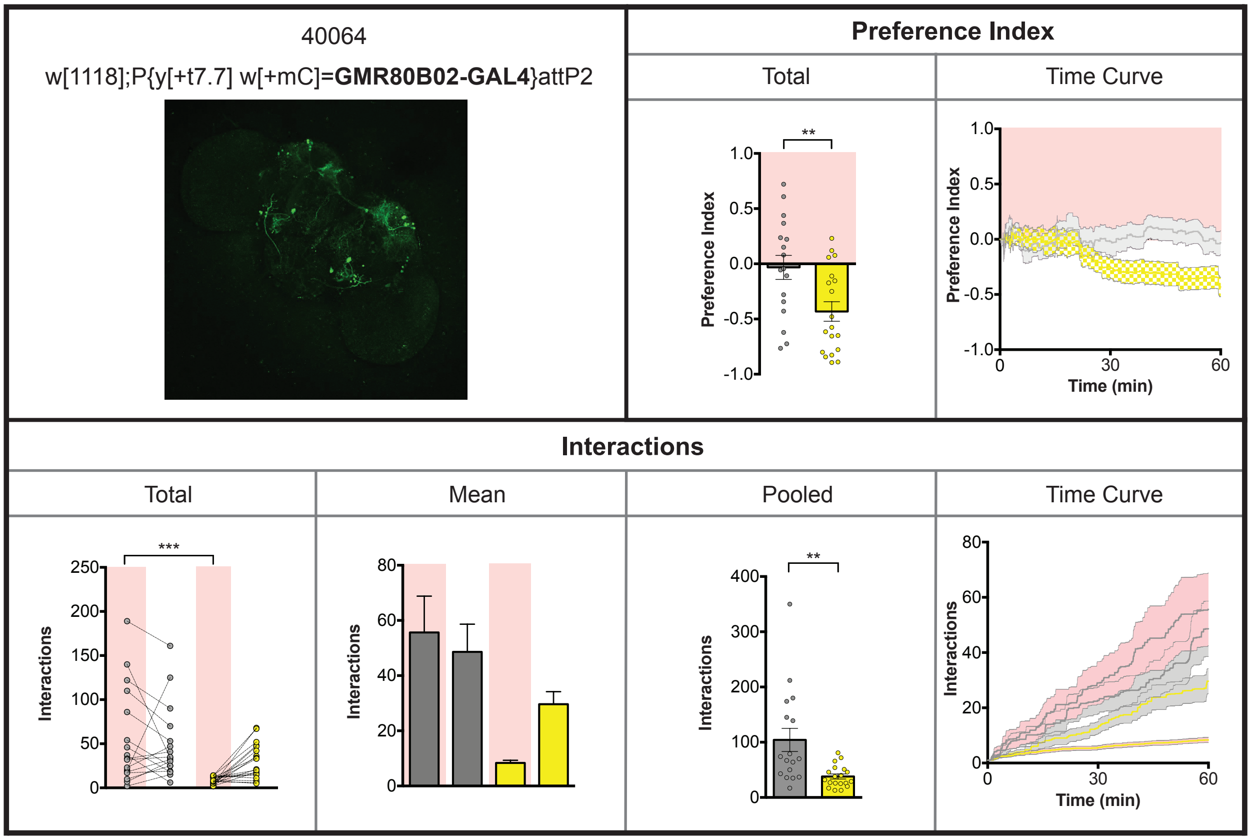
40064
w[1118];P{y[+t7.7] w[+mC]=**GMR80B02-GAL4**}attP2
Preference Index
Total
Time Curve
**
Preference Index
1.0
0.5
0.0
-0.5
-1.0
Time (min)
0
30
60
Interactions
Total
Mean
Pooled
Time Curve
***
250
200
150
100
50
0
80
60
40
20
0
**
400
300
200
100
0
Interactions

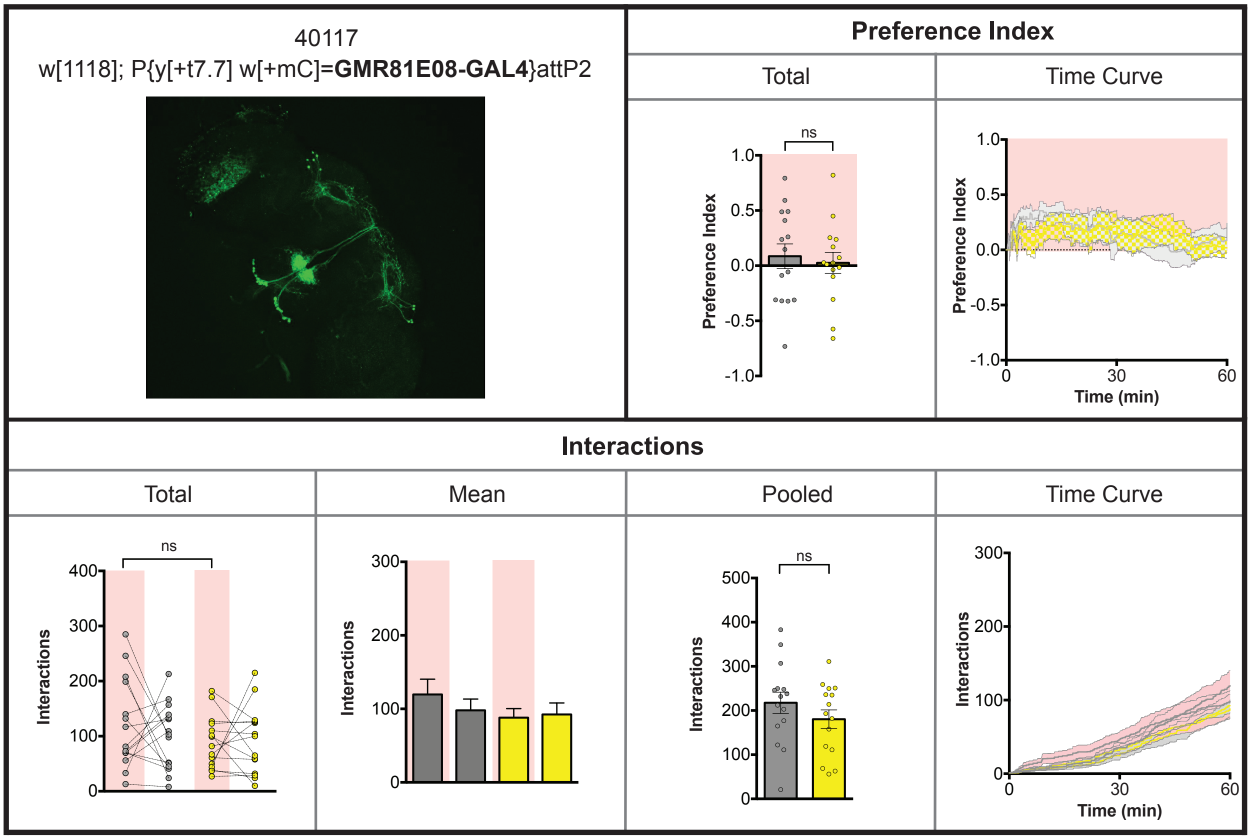
40117
w[1118]; P{y[+t7.7] w[+mC]=**GMR81E08-GAL4**}attP2
Preference Index
Total
Time Curve
ns
Preference Index
1.0
0.5
0.0
-0.5
-1.0
Time (min)
0
30
60
Interactions
Total
Mean
Pooled
Time Curve
ns
400
300
200
100
0
500
Interactions

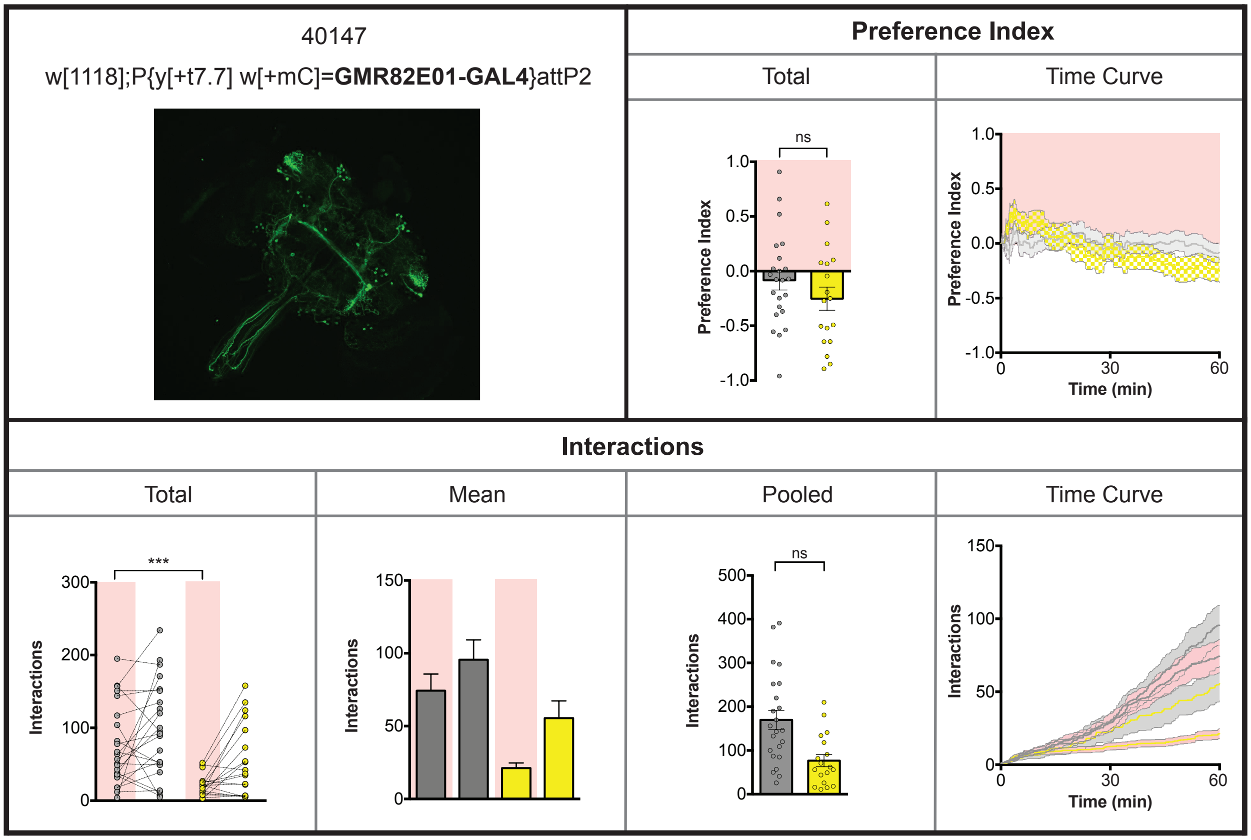
40147
w[1118];P{y[+t7.7] w[+mC]=GMR82E01-GAL4}attP2
Preference Index
Total
Time Curve
ns
Preference Index
Time (min)
Interactions
Total
Mean
Pooled
Time Curve
***
Interactions
ns
Time (min)

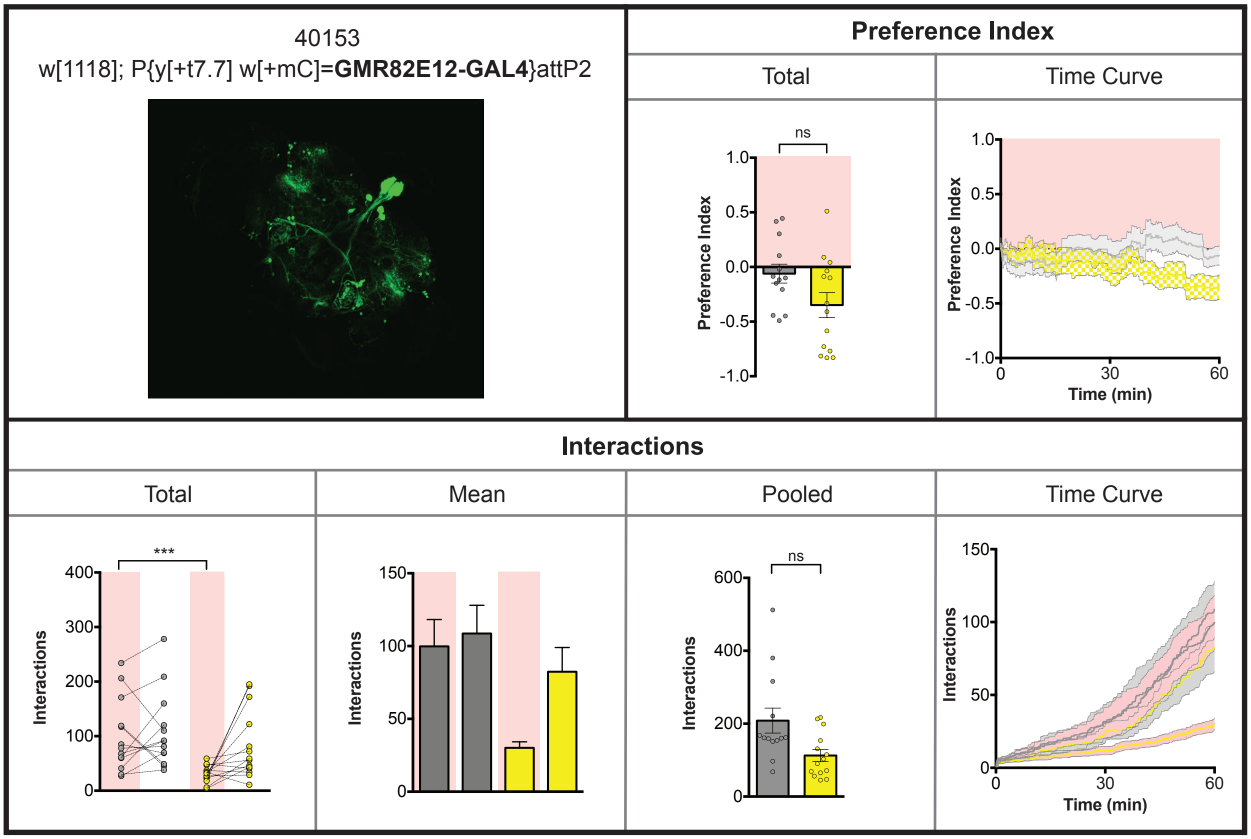
40153
w[1118]; P{y[+t7.7] w[+mC]=GMR82E12-GAL4}attP2
Preference Index
Total
Time Curve
ns
Preference Index
Time (min)
Interactions
Total
Mean
Pooled
Time Curve
***
Interactions
ns
Time (min)

40360

w[1118];P{y[+t7.7] w[+mC]=**GMR83D12-GAL4**}attP2

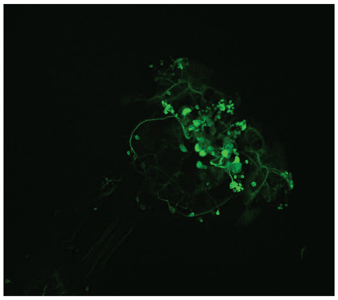

### Preference Index

Total

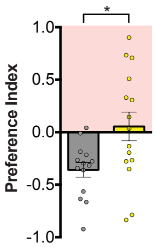


Time Curve

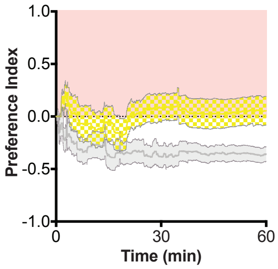


### Interactions

Total

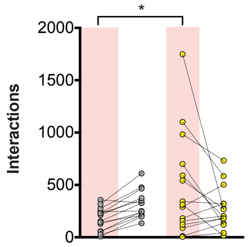


Mean

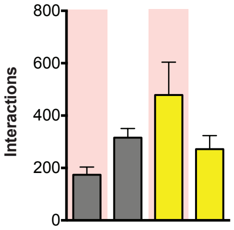


Pooled

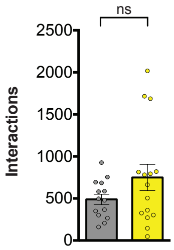


Time Curve

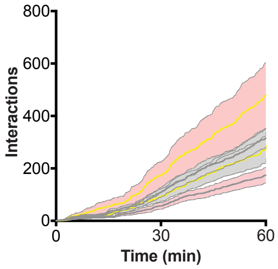


48378

w[1118]; P{y[+t7.7] w[+mC]=**GMR84C10-GAL4**}attP2 /TM3, Sb[1]

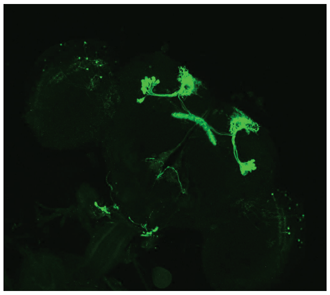

### Preference Index

Total

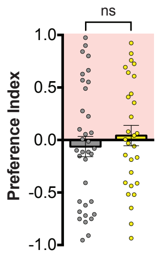


Time Curve

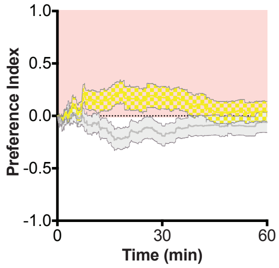


### Interactions

Total

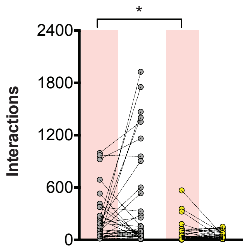


Mean

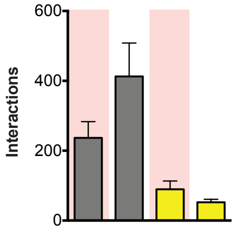


Pooled

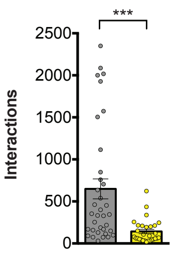


Time Curve

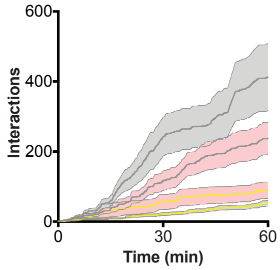

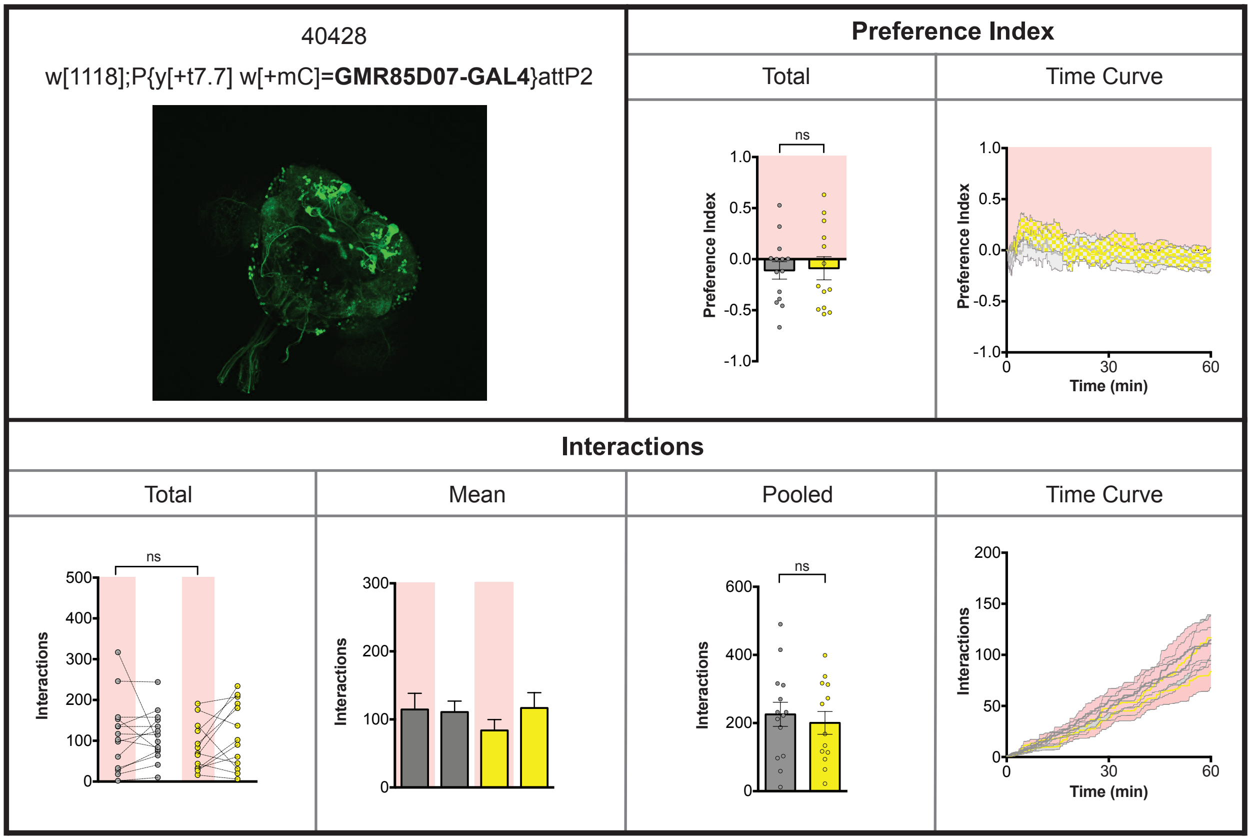
40428
w[1118];P{y[+t7.7] w[+mC]=**GMR85D07-GAL4**}attP2
Preference Index
Total
Time Curve
ns
Preference Index
Time (min)
Interactions
Total
Mean
Pooled
Time Curve
ns
Interactions
Time (min)

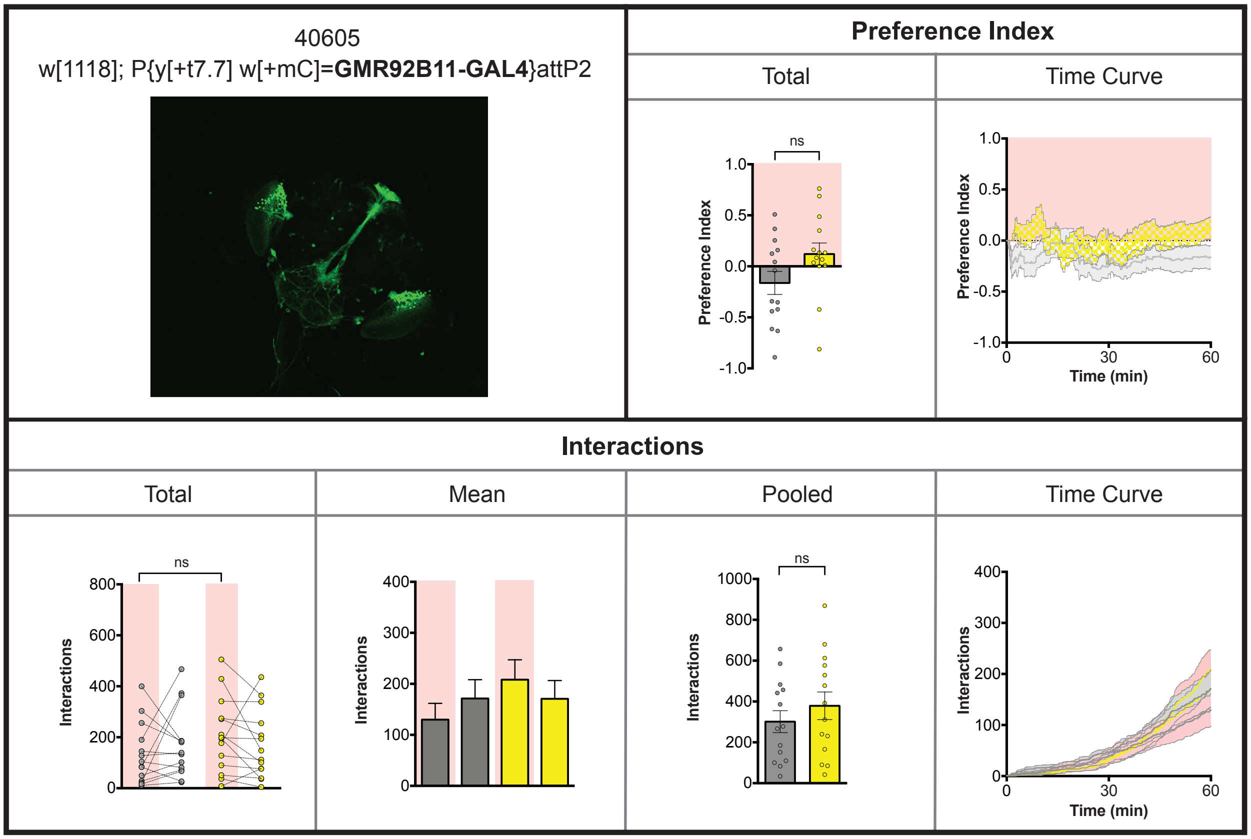
40605
w[1118]; P{y[+t7.7] w[+mC]=**GMR92B11-GAL4**}attP2
Preference Index
Total
Time Curve
ns
Preference Index
Time (min)
Interactions
Total
Mean
Pooled
Time Curve
ns
Interactions
Time (min)

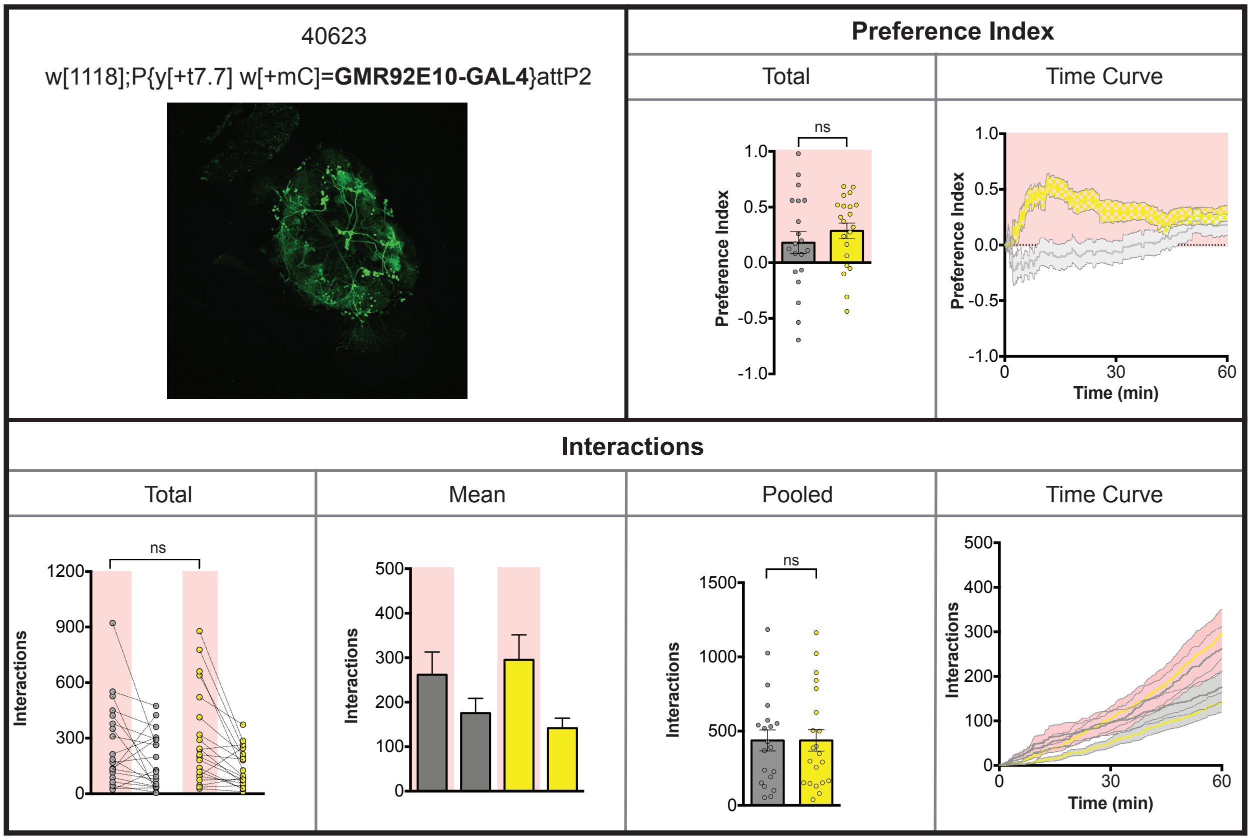
40623
w[1118];P{y[+t7.7] w[+mC]=GMR92E10-GAL4}attP2
Preference Index
Total
Time Curve
ns
Preference Index
1.0
0.5
0.0
-0.5
-1.0
0
30
60
Time (min)
Interactions
Total
Mean
Pooled
Time Curve
ns
Interactions
1200
900
600
300
0
500
400
300
200
100
1500
1000
500
ns
Time (min)

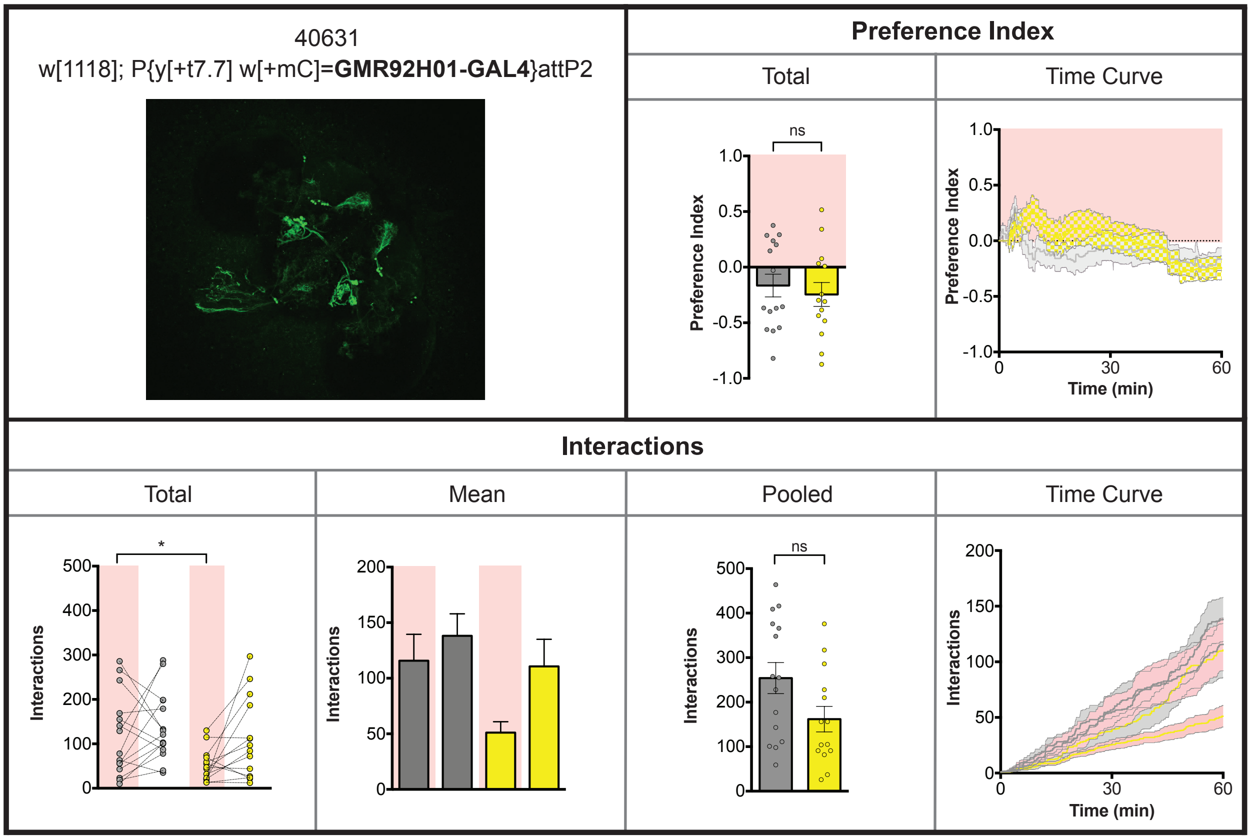
40631
w[1118]; P{y[+t7.7] w[+mC]=GMR92H01-GAL4}attP2
Preference Index
Total
Time Curve
ns
Preference Index
1.0
0.5
0.0
-0.5
-1.0
0
30
60
Time (min)
Interactions
Total
Mean
Pooled
Time Curve
*
Interactions
500
400
300
200
100
0
200
150
50
ns
Time (min)

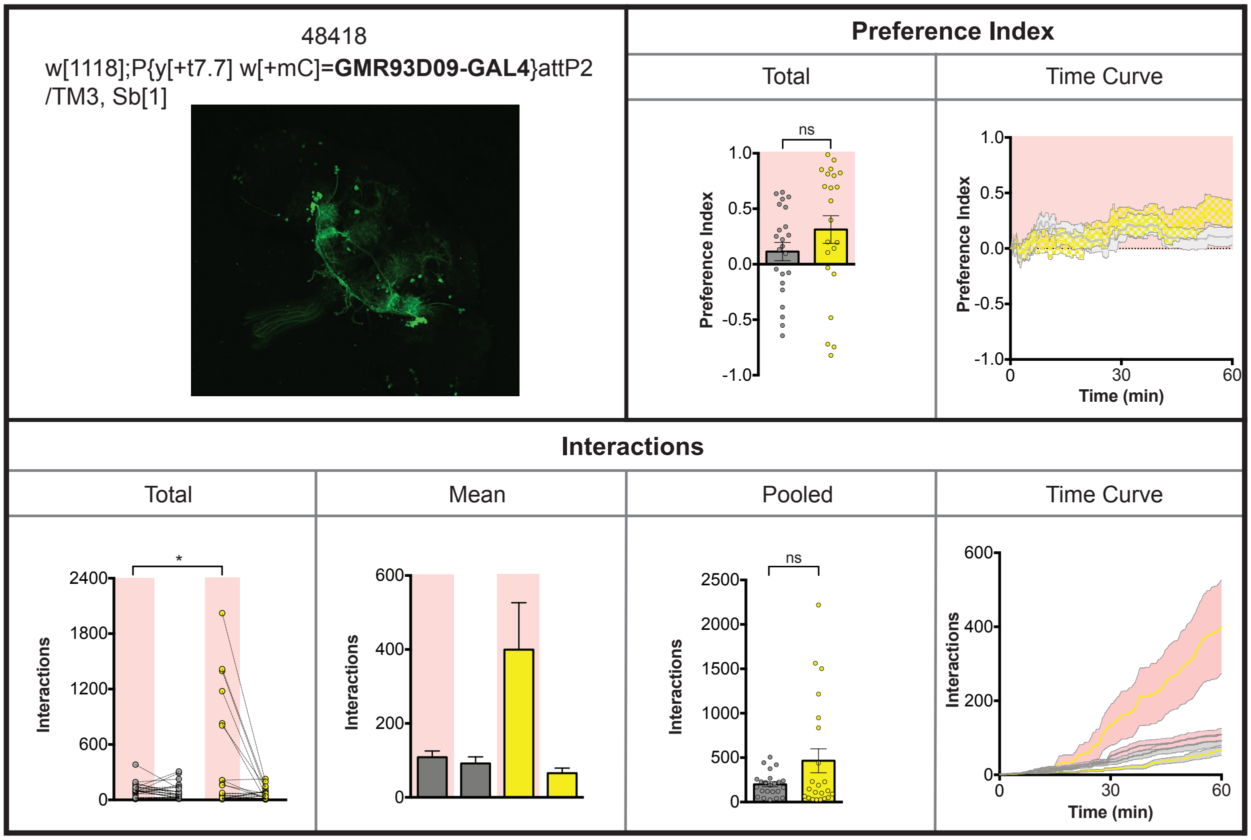
48418
w[1118];P{y[+t7.7] w[+mC]=GMR93D09-GAL4}attP2
/TM3, Sb[1]
Preference Index
Total
Time Curve
ns
Preference Index
1.0
0.5
0.0
-0.5
-1.0
0
30
60
Time (min)
Interactions
Total
Mean
Pooled
Time Curve
*
2400
1800
1200
600
0
600
400
200
ns
2500
2000
1500
1000
500
Interactions

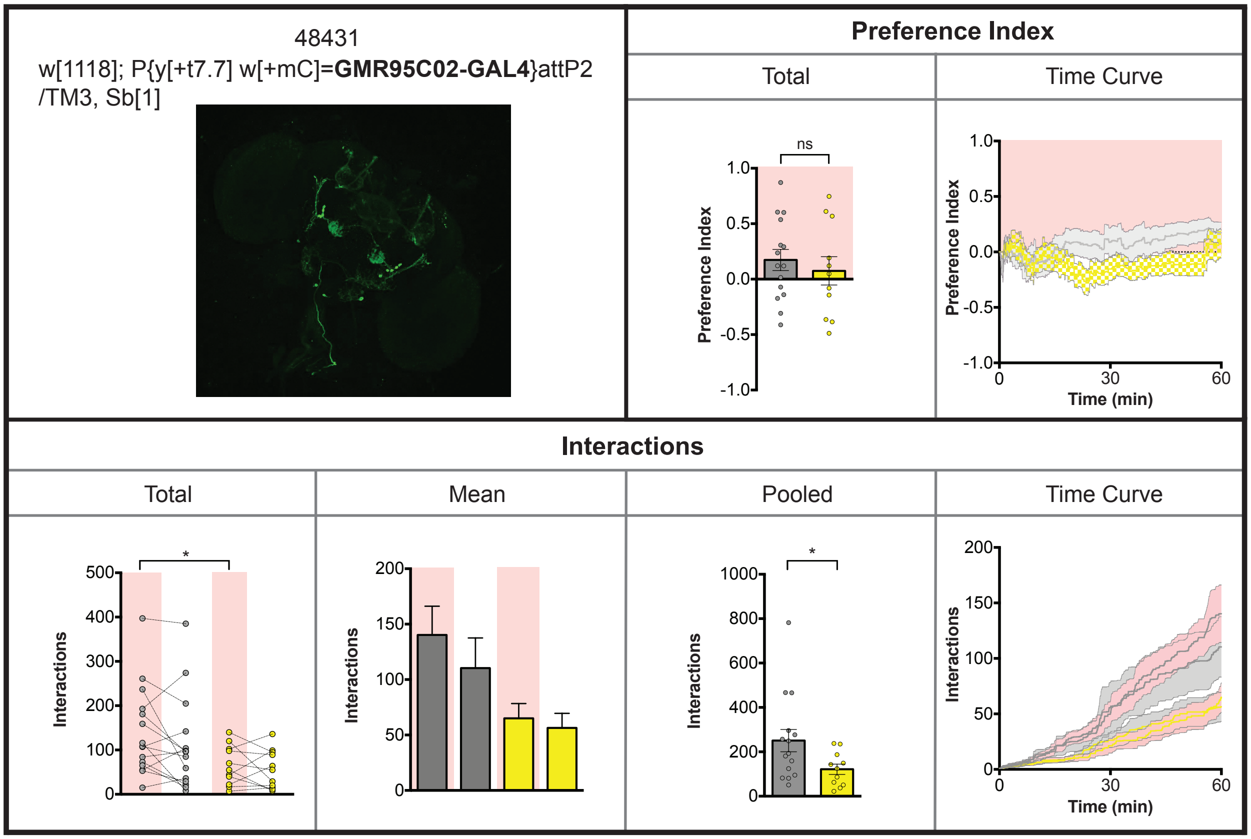
48431
w[1118]; P{y[+t7.7] w[+mC]=GMR95C02-GAL4}attP2
/TM3, Sb[1]
Preference Index
Total
Time Curve
ns
Preference Index
1.0
0.5
0.0
-0.5
-1.0
0
30
60
Time (min)
Interactions
Total
Mean
Pooled
Time Curve
*
500
400
300
200
100
0
150
50
1000
800
600
Interactions
